# The receptor binding properties of H5Nx influenza A viruses have evolved to promiscuously bind to avian-type mucin-like O-glycans

**DOI:** 10.1101/2025.04.24.650378

**Authors:** Julia Weber, Niels L.D. Ponse, Xueyong Zhu, María Ríos Carrasco, Alvin X. Han, Mathis Funk, Ting-Hui Lin, Alba Gabarroca García, Cindy M. Spruit, Ding Zhang, Wenli Yu, Ian A. Wilson, Mathilde Richard, Geert-Jan Boons, Robert P. de Vries

## Abstract

Highly pathogenic H5Nx influenza A viruses are causing unprecedented, season-independent outbreaks across avian and mammalian species, including dairy cattle, a novel reservoir. The sialoside-binding properties of influenza A hemagglutinin (HA) are strongly related to its ability to infect and transmit between hosts. Mucin-like O-glycans, omnipresent in respiratory tracts, have been understudied as viral receptors due to their complexity. To address this, we synthesized 25 O-linked glycans with diverse sialosides, including modifications by fucosides and sulfates. Our findings reveal that H5Nx 2.3.4.4b viruses uniquely bind core 3 sialyl-Lewisx and Sia-Gal-β3GalNAc, glycans not recognized by classical H5 or other avian viruses. By determining its crystal structure, we resolved the structural features of both structures in an H5 hemagglutinin (HA) from a 2016 2.3.4.4b virus. While these viruses do not bind human-type receptors, their promiscuous receptor specificity enhances binding to human tracheal tissues, suggesting that O-glycan recognition contributes to their zoonotic potential.

## Introduction

Influenza A viruses (IAV) are maintained in wild waterfowl, regularly spill-over to poultry and, occasionally to mammalian species, including humans^1^. Only a few lineages of influenza A viruses exist in mammalian hosts, such as H3N8 viruses in horses, H1 swine viruses, and pandemic H1N1, H2N2, and H3N2 viruses in humans^2^.

In 2021, highly pathogenic H5N1 viruses belonging to the 2.3.4.4b hemagglutinin (HA) phylogenetic clade appeared ^3,4^, which are now circulating year-round from the northern to the southern hemisphere. These viruses cause high mortality in wild birds^5,6^, and regularly infect poultry. They have been transmitted to several species of mammals^7–9^. Sustained mammal-to-mammal transmission in fur farms in Europe and South American marine mammals was a first for H5N1 viruses^10^. A recent zoonotic transmission into dairy cattle represents an exceptional novel viral reservoir in important livestock with occasional transmission to dairy farm workers^11^. With humans being immunologically naïve to H5N1 viruses, it is vital to establish molecular determinants important for the increased zoonotic potential of these viruses.

A widely accepted paradigm is that avian viruses require terminal α2,3-linked sialosides (SIAs) as receptors for cell binding and entry, while human IAVs employ α2,6-linked SIAs as receptors ^12,13^. Recent studies have indicated that this notion is an oversimplification and other features, such as fucosylation, sulfation, and the length of *N*-acetyl lactosamine (LacNAc, galactose−β1-4-N-acetylglucosamine) chains, can determine or modulate receptor specificity and may represent barriers to cross-species infection^14,15^. Only one sialyl transferase (ST6Gal1) can transfer terminal α2,6-linked SIAs onto oligosaccharide chains, and this enzyme preferentially modifies LacNAc moieties of *N*-glycans^16,17^. On the other hand, several sialyl transferases can add α2,3-linked SIAs on *N*- and *O*-linked glycans and glycolipids, resulting in a far greater structural diversity^18^. A range of *N*-linked glycans has been synthesized and employed for glycan array development, which allowed the investigation of how *N*-glycan structural complexity can influence receptor specificity^13,19–22^. It has, for example, been found that an asymmetric *N*-glycan having an α2,6-sialoside at a di-LacNAc moiety is a commonly employed receptor by human influenza A viruses^23^.

Several sialyl transferases can catalyze the addition of α2,3-linked SIAs on *O*-glycans, and this type of sialoside is particularly abundant on mucins^24,25^. IAVs have evolved an ability to attach and penetrate the mucus layer of respiratory and enteric tissues^26,27^ and several reports indicate that IAVs can bind core 2, 3, and 4 *O*-glycan structures^28,29^. There are, however, no systematic investigations of binding specificities of IAVs to biologically relevant *O*-glycans.

To address this deficiency, we developed a chemoenzymatic approach providing a library of 25 *O*-linked glycopeptides with different cores and terminal epitopes. These compounds were printed as a glycan microarray, which was employed to probe receptor specificities of evolutionary distinct H5 influenza viruses. We focused on H5 influenza viruses of the 2.3.4.4b clade, including a virus representing the HA of a human isolate of the cattle outbreak^30^, and examined differences in receptor specificities from their predecessors. Virus HAs of the 2.3.4.4b clade have undergone major antigenic changes due to mutations in and around the receptor binding site, including S133aL, ST137-138AA, NST158-160NDA, K/R193N, Q196K, K222Q and S227R (H3 numbering throughout) (Fig. S1)^31,32^. The micro-array binding data revealed that the currently circulating 2.3.4.4b H5 viruses are unique in their ability to bind sialyl Lewis^x^ (SLe^x^) on a core 3 *O*-glycan and the sialylated β3 arm of a core 2 *O*-linked glycan. Classical H5 HAs isolated at the beginning of this century have a more restricted receptor binding profile and preferentially bound sialylated sulfated LacNAc structures. Notably, the broader receptor specificity of 2.3.4.4b viruses coincided with binding to human upper respiratory tract tissues while binding to chicken trachea was maintained. We determined crystal structures of a representative 2.3.4.4b HA with specific mucin-like O-glycans and demonstrated how the fucoside is accommodated in 2.3.4.4b viruses by HA residues Q222 and G225. We also demonstrated that the promiscuous receptor specificity is due to positions 133a and 227 and is unique, as no other subtype displayed such receptor binding properties. The broader receptor repertoire of H5 of the 2.3.4.4b clade for *O*-glycans may be relevant for their broad host range.

## Results

### Chemoenzymatic synthesis of a library of *O*-linked glycopeptides

We embarked on the chemoenzymatic synthesis of a panel of *O*-linked glycopeptides displaying different core structures and terminal modifications (Fig. 1). O-glycan biosynthesis is initiated by adding a single GalNAc residue to Ser/Thr to form a so-called Tn-antigen ^33^. The latter moiety can be extended by a β1,3Gal moiety to form core 1, which subsequently can be extended to core 2 by adding β1,6GlcNAc to the GalNAc residue. Core 3 has a β1,3GlcNAc linked to the Tn-antigen, and core 4 is further branched by the addition of β1-6GlcNAc. The core structures can be modified by various epitopes, including sialosides, blood groups, and Lewis antigens. Core 1 and 2 containing structures are commonly found on glycoproteins and mucins of many different cell types, whereas core 3 and 4 display is more restricted, and these structures are mainly found on gastrointestinal tissues^33^.

**Figure 1.**
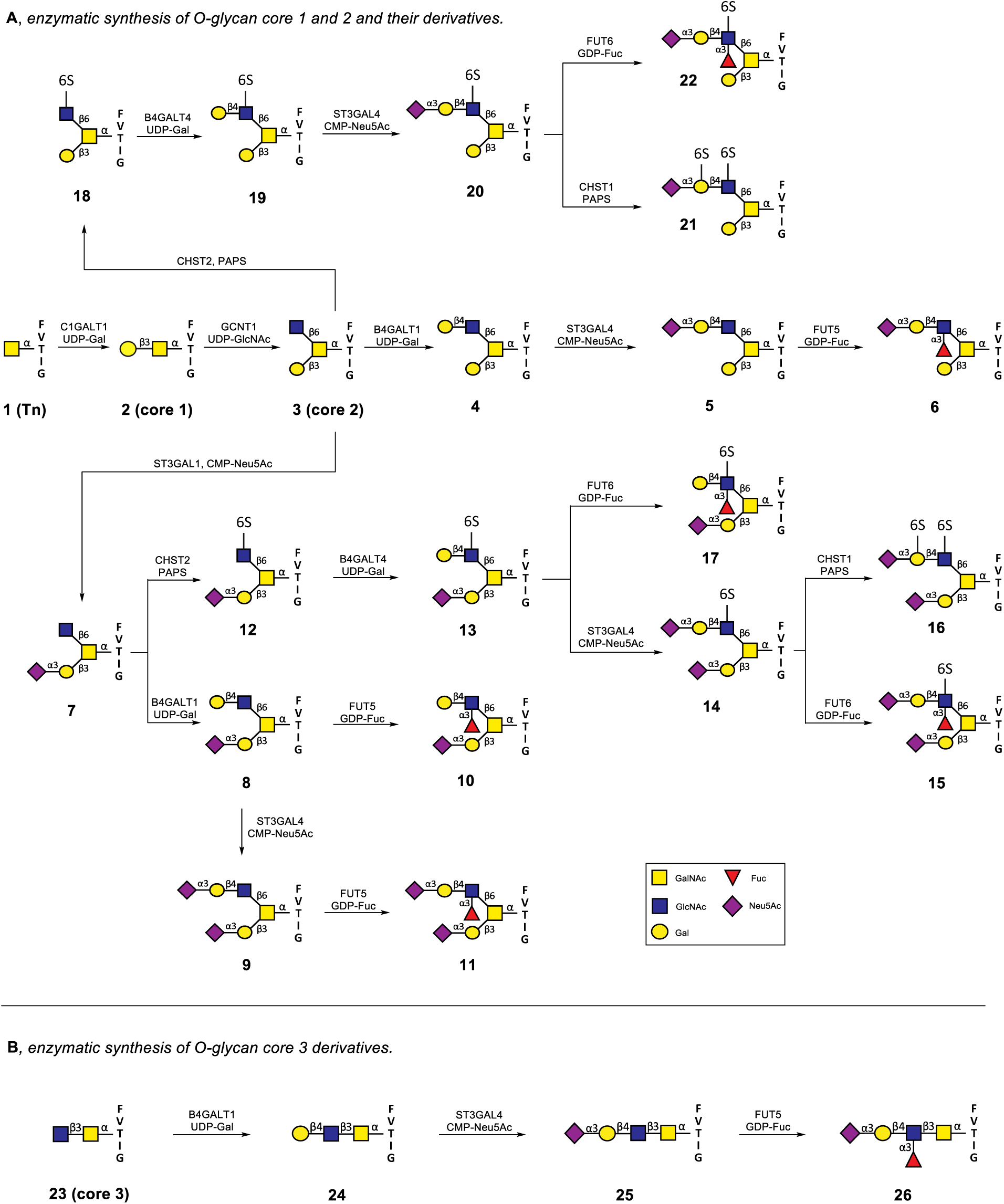
Chemoenzymatic synthesis of mucin-like O-glycans. (A) Enzymatic synthesis of core 1 (**2**), core 2 (**3**) and extended core 2 O-glycan structures (**4-22**) (B) Enzymatic synthesis of extended core 3 O-glycans (**23- 26**)

Compounds **2**, **3,** and **23** represent core 1, 2, and 3 structures, respectively (Fig. 1). We targeted compounds with terminal epitopes that may function as receptors of IAV. It includes 2,3-linked sialosides and derivatives with additional fucosylation and/or sulfation to give sialyl Lewis^x^ (SLe^x^) and 6-sulfo-SLe^x^ and 6-sulfo-sialyl-LacNAc derivatives. There are data to support that IAV can employ such structures as receptors^15,32,34,35^ and, therefore, were included as target compounds. The attraction of this series of glycopeptides is that these epitopes are presented on different core structures, making it possible to examine whether these differences influence binding.

The synthesis of the core 1 and 2 structures started with the assembly of glycopeptide **1** by solid phase peptide synthesis (SPPS) using a Rink amide AM resin, Fmoc protected amino acids, and Fmoc-Thr-(3,4,6-tri-*O*-acetyl-α-D-GalNAc) (Scheme S1). After assembly, the glycopeptide was cleaved and purified by HPLC employing a reverse-phase C18 column. The *N*-terminal α-amine of the glycopeptides was left free to facilitate immobilization onto NHS ester-activated microarray slides for glycan array development.

The enzyme glycoprotein-*N*-acetyl galactosamine 3-β-galactosyltransferase 1 (C1GalT1) can convert a Tn antigen into a core 1 structure (**2**). It requires an underlying peptide chain, and earlier studies have shown that the peptide sequence FVTIG is appropriate for this enzyme ^36^. As anticipated, treatment of **1** with uridine-5’-diphospho-galactose (UDP-Gal) in the presence of recombinant C1GalT1 resulted in the facile formation of glycopeptide **2**. The latter compound was converted into core 2 by β-1,3-galactosyl-*O*-glycosyl-glycoprotein β-1,6-N-acetylglucosaminyltransferase (GCNT1) and uridine-5’-diphospho-glucosamine (UDP-GlcNAc) (Fig. 1A). The resulting glycopeptide **3** was the starting point for the installation of various terminal epitopes. For example, the GlcNAc moiety of **3** could be extended by a β(1,4)-galactoside to give LacNAc-containing derivative **4** by treatment with β-1,4-galactosyltransferase 4 (B4GalT4) and UDP-Gal. The resulting terminal Gal moiety of glycopeptide **4** could selectively be modified as an α2,3-sialoside by using α2,3-sialyltransferase 4 (ST3Gal4) and cytidine-5′-monophospho-*N*-acetylneuraminic acid (CMP-Neu5Ac) to form glycopeptide **5**. SLe^x^ containing glycopeptide **6** was formed by treatment of **5** with either α1,3-fucosyltransferase 5 (FUT5) or 6 (FUT6) and guanosine-5’-diphospho-β-L-fucose (GDP-Fuc).

In an alternative sequence of reactions, the β1,3-galactoside of core 2 glycopeptide **3** was selectively α2,3-sialylated to give **7** by treatment with α2,3-sialyltransferase 1 (ST3Gal1) in the presence of CMP-Neu5Ac. This compound could be converted into a variety of structures, and for example, the enzyme B4GalT1 could convert the terminal GlcNAc moiety into LacNAc **8,** which could be further sialylated by ST3Gal4 to give sialylated glycopeptide **9**. Treatment of **8** or **9** with FUT5 or FUT6 resulted in the formation of Lewis^x^ (Le^x^) and SLe^x^ containing compounds **10** and **11**, respectively.

The C-6 hydroxyl of the GlcNAc moiety of core 2 structures can be sulfated by GlcNAc-6-*O*-sulfotransferases 2 and 6 (CHST2 and 6)^37^. This enzyme only modifies terminal GlcNAc moieties, and as anticipated, treatment of **7** with 3′-phosphoadenosine-5′-phosphosulfate (PAPS) in the presence of CHST2 resulted in the facile formation of sulfated glycopeptide **12**. The galactosyl transferase B4GalT4 can attach a 1,4-linked galactoside to 6-sulfo-GlcNAc residues and, as expected, could readily convert **12** into **13,** which is an appropriate substrate for ST3Gal4 to give glycopeptide **14** having 6-sulfo-sialyl-LacNAc moiety. The latter compound could be fucosylated with FUT6 to provide 6-sulfo-SLe^x^ containing glycopeptide **15**. A sulfo-sialyl-LacNAc moiety is an appropriate substrate for keratan sulfate galactose 6-sulfotransferase (KSGal6ST or CHST1), which in the presence of PAPS can sulfate the C-6 hydroxyl of the galactoside and this enzyme could readily convert **14** into the di-sulfate **16**. Compound 13 was also fucosylated with FUT6 to provide 6-sulfo-Le^x^ containing glycopeptide **17**. A similar sequence of reactions starting from **18**, which was synthesized by sulfation of **3** with CHST2 and PAPS, provided compounds **19**, **20**, **22,** and **21**.

Acetylgalactosaminyl-O-glycosyl-glycoprotein β-1,3-N-acetylglucosaminyltransferase (B3GNT6) is the enzyme that can convert core 1 (**2**) into a core 3 structure **23** (Fig. 1). This enzyme has limited *in vitro* activity ^38^ and could not successfully be expressed; therefore, we chemically synthesized appropriately protected GlcNAc (S1 &S3), GalNAc-Thr, which was employed in glycopeptide assembly to give the core 3 containing structure **23** (Scheme S1). This compound could be extended by galactosylation by B4GalT1 in the presence of UDP-Gal to form glycopeptide **24**. α2,3-Sialylation of **24** was achieved by using ST3Gal4 and CMP-Neu5Ac to obtain **25**. Treatment of **25** with FUT5 and GDP-Fuc resulted in the formation of **26** (Fig. 1B). An attempt to sulfate core 3 (**7**) with PAPS and CHST2 or CHST6 failed, and probably these compounds are not appropriate substrates for these enzymes.

### Glycan microarray development and screening

The library of 25 O-glycans having various terminal epitopes was printed on amine-reactive NHS-activated glass slides (Fig. 2A), resulting in an *O*-glycopeptide microarray. The resulting array slides were probed by several plant lectins ^39^, which confirmed spot integrity and proper substrate deposition and revealed fine specificities of glycan-binding proteins examined for *O*-glycans (Fig. S2). The lectins MAL-I and MAL-II have well-known affinity for α2,3-sialylated glycans and are commonly used for detecting avian IAV receptors on tissues ^40^. On the array, MAL-I showed robust binding to α2,3-sialylated LacNAc structures on O-glycan cores 2 and 3 (**5**, **9**, **25**) and tolerated a 6-sulfate on GlcNAc (**14**, **20**) (Fig. S2). MAL-II exhibited a more expanded responsiveness to sulfated core 2 O-glycans with an α2,3-linked SIA on the β3-branch (**12**, **13**) and to sulfated core 2 di-sialosides (**14-16**), and the results highlight that the specificities of MAL-I and -II are significantly different. A mouse IgM antibody (CD15s, clone CSLEX1)) was used to detect SLe^x^ epitopes, which are present on structures **6** and **11** (Fig. S2B). Notably, glycan **26** was not recognized by this IgM. The lectin ECA detected non-sialylated epitopes^41^ only binding to terminal type 2 LacNAc structures. It tolerated a 6-sulfate but not fucose on GlcNAc.

**Figure 2.**
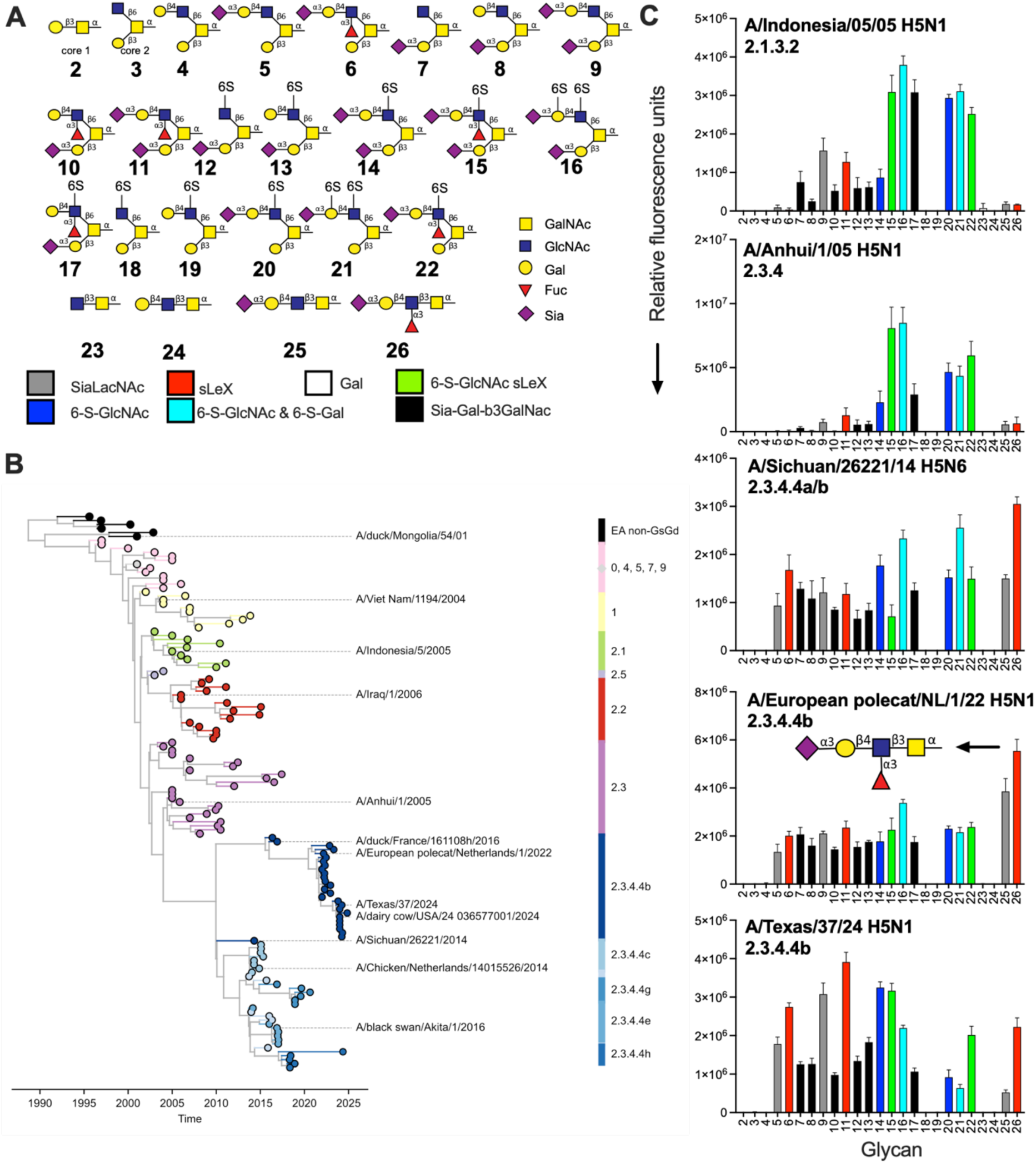
Mucin-like O-glycan binding analyses of H5 viruses. (A) The shown O-glycans were printed on a glycan microarray and the terminal epitopes are indicated in different colors. Monosacharides nomenclature is shown. (B) phylogenetic tree of H5 viruses used in this work, reconstructed temporally using TreeTime (C) The specificity of several influenza A H5 viruses, with relative fluorescence units on the y-axis and glycan numbering on the X-axis as in (A). Shown is a representative of two independent assays, standard deviation from 6 technical replicates with the highest and lowest data points removed.

Additionally, lectins Jacalin and PNA were used to analyze non-sialylated epitopes, specifically terminal Tn-antigen^42,43^ (Fig. S2C). Interestingly, both lectins recognized different O-glycan structures since Jacalin did not tolerate 6-sulfation on the GlcNAc and recognized **2, 23**, and **24**. In contrast, PNA more selectively recognized all terminal T-antigen epitopes. Ricin was used to analyze type 2 LacNAc^44^, which bound the unmodified terminal type 2 LacNAc. AAL was used to detect fucosylated O-glycans^45^ and indeed recognized all fucosylated structures.

### H5Nx 2.3.4.4 HAs have a promiscuous yet specific receptor-binding repertoire

Next, we focused on receptor binding selectivity of different H5Nx viruses (Fig. 2B). Clade 2.3.4.4b viruses have accumulated significant changes in the receptor binding site (RBS) in the 130, 150, 220-loops, and 190-helix (3) (Fig. S1). Specifically, positions 132, 133a, 137, 158-160, 193, 196, 222, and 227 in the receptor binding site have changed between classical (viruses isolate before 2010) and current H5 viruses and can affect receptor binding properties^12^; however, amino acid changes conferring human-type receptor binding are absent.

Recombinant viruses that contained the HA (without a multibasic cleavage site) of interest in the background of the attenuated vaccine strain A/Puerto Rico/8/1934 (PR/8) were handled under biosafety level (BSL) 2 conditions in agreement with national regulations on genetically modified viruses (GMO-99-090 and IG 15-160_IIk). All viruses in this study are thus recombinant, but we do refer to them with the full virus name throughout the manuscript. We used two pre-2.3.4.4 viruses, A/Indonesia/05/2005 (2.1.3.2) and 2.3.4 A/Anhui/1/2005 (Fig. 2B), it is known that such viruses bind terminal α2,3-linked SIAs on N-glycans^46,47^; however, detailed binding specificities of *O*-glycans have not been determined.

The A/Indonesia/05/2005 virus bound to core 2 O-glycans with a sialylated LacNAc containing a 6-sulfated GlcNAc (**15-17** and **20-22**), with slight preference if these structures displayed 2 sialic acids, compound **9** over **5**, **11** over **6**, **8** over **9** and **11** over **10** (Fig. 2C). The direct precursor 2.3.4 virus (A/Anhui/1/2005) also displays a conventional avian-type (α2,3-linked SIA) receptor binding profile, very similar to A/Indonesia/05/2005, and is not able to bind the unmodified sialylated LacNAc (**5**, **9**, and **25**). The A/Anhui/1/2005 virus preferred sialylated LacNAc in which the GlcNAc moiety is sulfated regardless of whether the LacNAc is fucosylated (**15, 16, 20-22**). Notably, the non-sulfated SLe^x^ displays minimal responsiveness (**6**, **11**, and **26**). Interestingly, the only Sia-Gal-β3GalNAc structures bound contain a sulfated Le^x^ (**17**).

Sulfated structures are less critical for 2.3.4.4b H5 viruses, as compounds **5-11** are equally bound. Both 2.3.4.4 viruses, A/Sichuan/26221/14 H5N6 and A/European polecat/NL/1/22 H5N1, bind every available α2,3-linked SIA (Fig. 2C). A/Sichuan/26221/14 H5N6 represents an older 2.3.4.4a subclade (Fig. S3), while the A/European polecat/Netherlands/1/2022 is a 2.3.4.4b virus. The differentiating features between non-2.3.4.4 and 2.3.4.4 viruses include simple mucin-like core 2 *O*-linked structures displaying Sia-Gal-β3GalNAc epitopes (**7, 8-10, 12, and 13**) and core 3 O-glycans, with the non-modified core 3 structure **25** efficiently bound as well as **26**. Finally, we tested a representative virus from the current bovine outbreak in the US, A/Texas/37/2024. Compared to the classical A/Indonesia/5/2005 and A/Anhui/1/2005 viruses, the binding profile is significantly widened, with high responsiveness to SLe^x^ epitopes (**6**, **11,** and **26**). However, compared to the early and the European 2.3.4.4b viruses, the A/Texas/37/2024 did not bind modified epitopes equally well when the Gal-β3GalNAc was not sialylated (**20** and **21**) and hardly bound **25**. Conclusively, 2.3.4.4b viruses have significantly broadened receptor repertoire compared to their predecessors, exemplifying receptor binding evolution and possibly explaining the increased zoonotic capabilities of these viruses.

### H5Nx 2.3.4.4 recombinant HAs display near-identical receptor binding specificities compared to viruses

Recombinant HAs are valuable tools for quickly and safely analyzing receptor binding properties of potential pandemic zoonotic viruses, as they are surrogates for whole viruses. Previously, we have shown that avian-like receptor binding is very similar between recombinant HAs and viruses, with the multivalency effect leading to higher relative fluorescent intensities detected when using viruses^46^.

To compare the data obtained using recombinant viruses’ data with those obtained with recombinant protein and confirm the broadened receptor binding specificities, we tested recombinant classical and 2.3.4.4.b HA H5 hemagglutinins. We employed the HAs from A/duck/Mongolia/54/01 H5N2^32^, A/Vietnam/1203/04 H5N1 (clade 1)^46^, and A/Indonesia/05/05 H5N1. The latter is identical at the amino-acid level to the recombinant virus HA, which is commonly used in receptor-binding studies^31^. Furthermore, we investigated two HA proteins from the 2.3.4.4 subclade, A/chicken/Kumamoto/1-7/2014 is a representative strain for subclade 2.3.4.4c, and we used a 2.3.4.4b subclade (A/duck/France/161108h/2016, H5FR), in which the multibasic cleavage site was removed^48^, which represents currently circulating panzootic viruses in Europe. The relation between the viruses used in this work is shown in the phylogenetic trees in Fig. 2B.

We observed that the HAs of the classical H5s preferentially bound core 2 O-glycans with sialylated LacNAc containing a 6-sulfated GlcNAc (**14-16, 20 and 22**) (Figs 3A and 3B). Indeed, the literature indicates that many avian viruses, including H5, can bind 6-sulfate GlcNAc-containing sialosides^15,48^. Interestingly, structures only displaying Sia-Gal-GalNAc (black bars) are hardly bound by these three HAs; similarly, compounds **25** and **26** are not bound. The extra SIA on the β3-branch is required for binding since glycan **9** is bound, in contrast to **5**.

**Figure 3.**
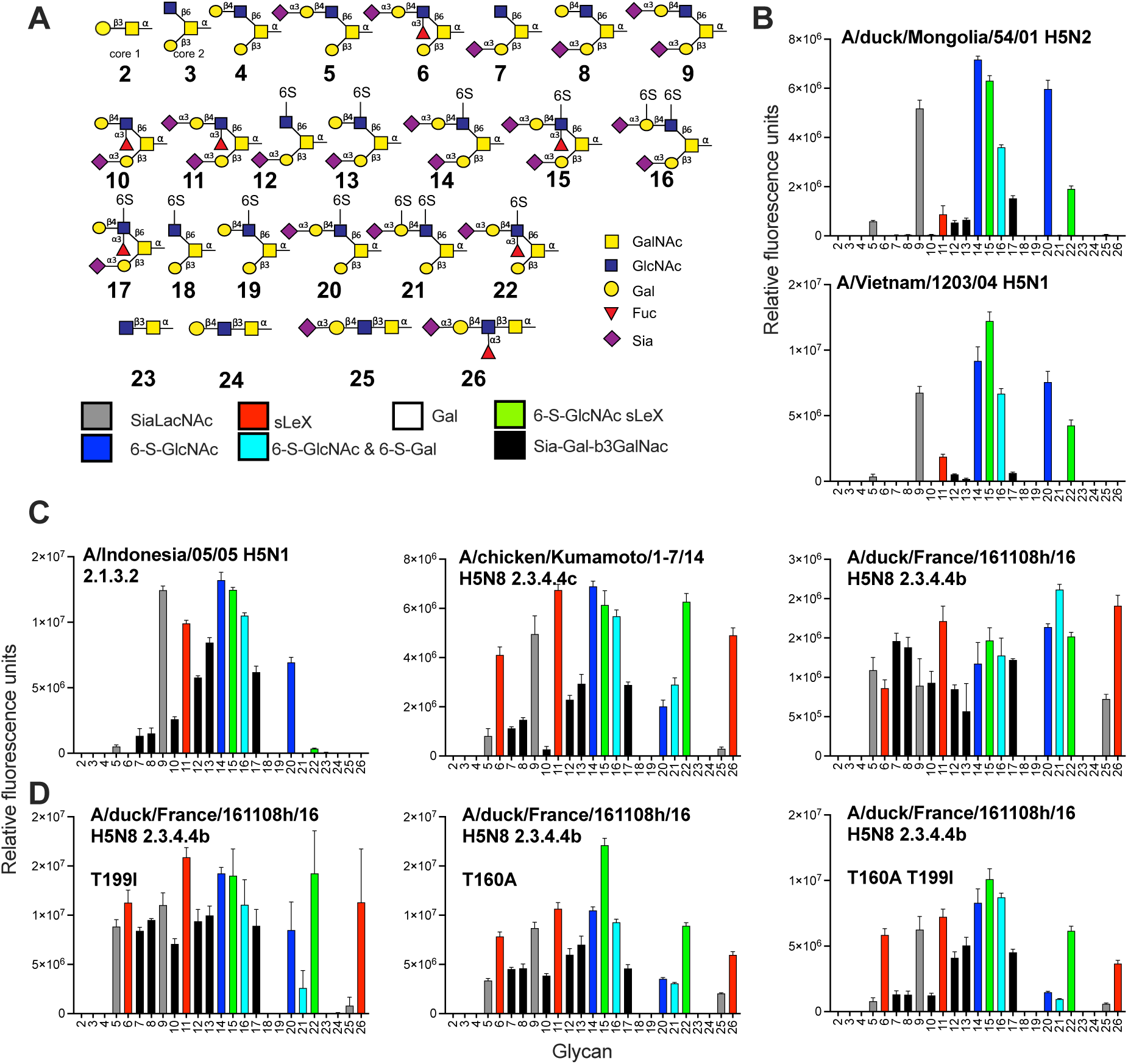
Mucin-like O-glycan binding analyses of H5 hemagglutinins. (A) The shown O-glycans were printed on a glycan microarray and the terminal epitopes are indicated in different colors. (B) The specificity of classical H5 HAs, (C) A/Indonesia/05/05 and two 2.3.4.4b H5 proteins and (D) H5FR mutant proteins. Relative fluorescence units are on the y-axis and glycan numbering on the X-axis as in (A). Shown is a representative of two independent assays, standard deviation from 6 technical replicates with the highest and lowest data points removed.

The A/Indonesia/5/05 protein (Fig. 3C) is identical on the amino acid level compared to the recombinant virus in figure 2C. The main difference is the lack of binding to **21** and **22**, while their counterparts (**15** & **16**) with an additional SIA are bound by both the virus and protein. The presence of an extra SIA on the β3-branch also allows binding to **9** and **11**. **11** has a sLe^x^ epitope on the β6-branch, while the presence of solely the SIA on the β3-branch (**7, 8,** and **10**) or the sLe^x^ on the β6-branch (**5**) does not allow for binding for classical H5s. Interestingly, structures **13** and **17** contain a 6-sulfate on the GlcNAc and showed increased binding compared to **7** and **8**, indicating that this charged structure is vital for binding. At the same time, this arm lies outside of the RBS as the position of SIA in the RBS is conserved.

The binding preference of HAs of 2.3.4.4 viruses, A/chicken/Kumamoto/1-7/2014 and H5FR is remarkably different. Substantial changes in amino acids are present in the HA1 receptor binding domains of these viruses compared to earlier classical H5s, including A132T, S133aL, S137A, NST158-160NDA, K/R193N, Q196K, K222Q, and S227R (Fig. S1)^31,32^. However, none of the canonical changes for α2,3-linked *vs.* α2,6-linked SIA specificities are present at positions 226 and 228^12^, and the H5FR indeed does not bind human-type receptors^50^. The representative H5 HAs from subclade 2.3.4.4 displayed a broader receptor binding repertoire compared to other H5 HAs, with a nearly identical trend compared to the viruses (Fig. 3C). The separating features between classical and 2.3.4.4b H5s include binding to **7**, **8**, **25** and **26**, representing Sia-Gal-β3GalNAc and SLe^x^ epitopes. These epitopes have been identified in different cells and organs from various species ^29,50,51^. We titrated an H5FR HA protein and a virus, A/European polecat/NL/1/22, on the array to check if we could identify preferred receptors, but none were found (Fig. S4). Thus, the broad host range of these viruses might be explained by the promiscuous binding specificity to the Sia-Gal-β3GalNAc structure or O-glycan Core 3 sLe^x^.

### Bovine-specific mutations T160A and T199I do not affect receptor binding specificities when introduced in H5FR HA

Clearly, classical and currently circulating H5N1 viruses have different receptor binding profiles. Indeed, several recent reports demonstrated that contemporary H5N1 viruses have a broadened receptor binding profile^52–54^. Moreover, one report suggested that this broader receptor binding profile compared to their European counterparts is due to T199I^56^. We, however, do not observe differences between the A/Texas/37/2024 virus and its European, A/European polecat/NL/1/22 H5N1, nor its 10-year-old Asian, A/Sichuan/26221/14 H5N6, predecessors. We introduced T199I into H5FR to analyze this discrepancy and determine if this mutant would further increase the binding repertoire (Fig 3D). The mutants displayed a receptor-binding repertoire very similar to the wild type. While circulating in cattle, small groups of viruses have been identified containing A160T, an antigenic change that introduces an N-glycosylation site but also affects human-type receptor binding in human-adapted H5N1 viruses^57^. When we introduced the A160T mutations in both the wildtype and T199I mutant, we observed some decrease in binding but no substantial change in binding specificity on the *O*-glycan array (Fig. 3D). Notable diminished binding was observed for compounds **5**, **7**, **8**, **10**, **20**, **21**, and **25**, but binding to **6** and **26** is maintained. We conclude that the receptor binding specificity of 2.3.4.4b HAs is conserved.

### The broad *O*-glycan specificity results in binding to human respiratory tract tissues

To test the hypothesis that the promiscuous or broad receptor binding profile of 2.3.4.4 H5Nx HAs can explain its remarkable zoonotic properties, we tested their ability to bind chicken and human tracheal tissues as representatives of upper respiratory tract organs. As controls, we analyzed a human H3N2 H3 protein (A/Netherlands/109/2003) that indeed only bound the human trachea tissue (Fig. 4). The H5FR HA protein bound to chicken with high responsiveness, and human tracheal tissue epithelial cells with moderate responsiveness in a SIA-dependent manner. The H5FR fails to bind α2,6-linked SIA containing-receptors^55^; thus, the human respiratory tract might display O-glycans that H5FR can bind. None of the classical H5 proteins could bind human tracheal epithelial cells. They did, however, efficiently bind to the chicken trachea in a SIA-dependent manner as neuraminidase treatment abrogated binding (Fig. 4).

**Figure 4.**
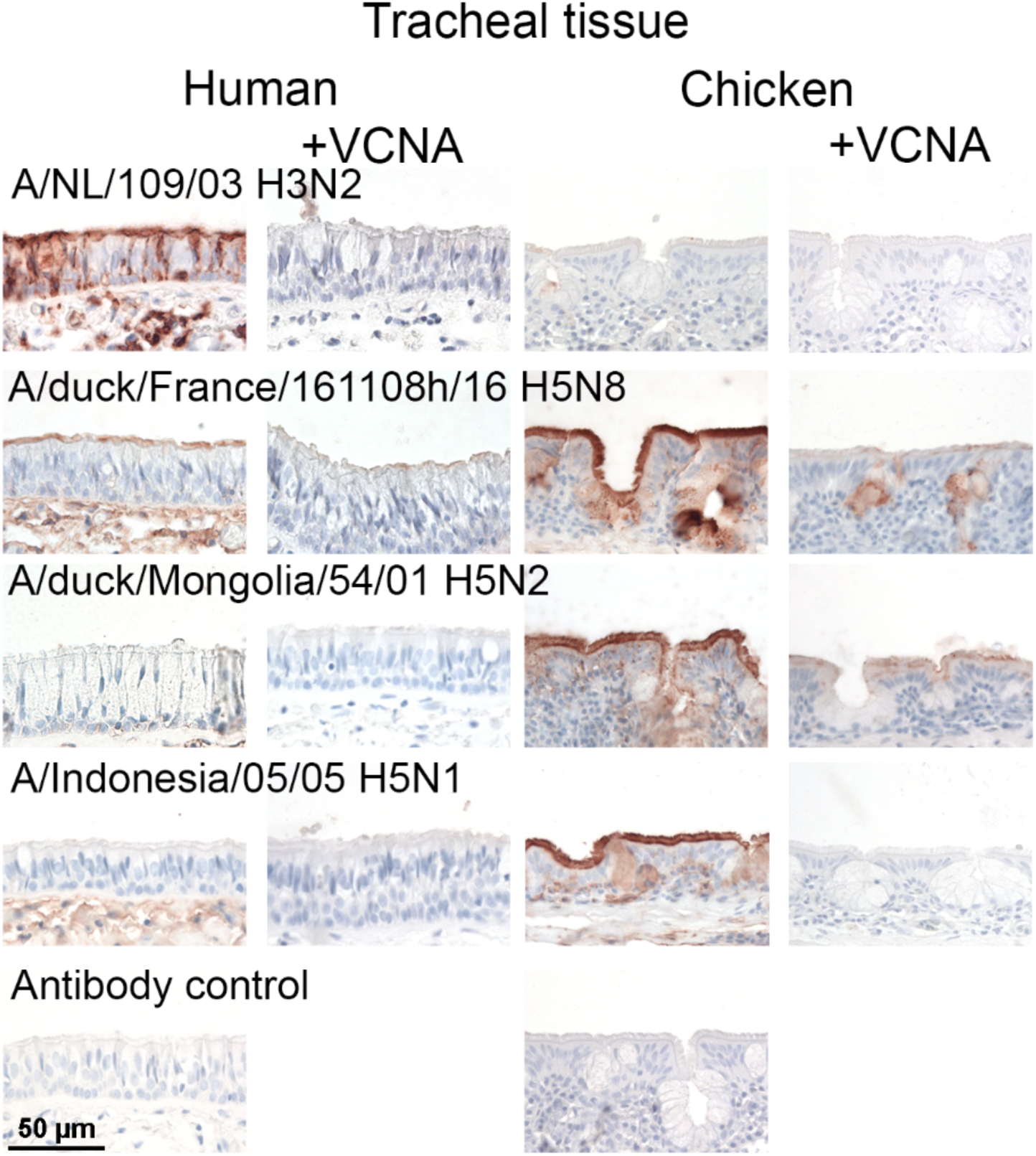
Tissue binding properties of A/duck/France/161108h/2016 (H5N8) are distinct from other H5 hemagglutinins. The binding to human and chicken tracheal tissue was investigated for different influenza A H5 HAs. The HA from A/NL/109/2003 was used as a positive control for human trachea binding. Sialic acid dependency was shown using tissue slides that were pre-treated with *Vibrio cholerae* neuraminidase (VCNA). AEC staining was used to visualize tissue binding.

To analyze the presence of SLe^x^ in these tissues, we employed the anti-SLe^x^ CD15s antibody (Fig. S5). Importantly, this antibody does not accommodate sulfated SLe^x^ structures (Fig. S2) and thus should be an appropriate control for the 2.3.4.4 H5Nx proteins to differentiate between SLe^x^ and their sulfated counterpart bound by non-2.3.4.4 HA proteins. The CD15s antibody bound efficiently to the chicken trachea^32^ but failed to attach to the human trachea. Thus, the human trachea does not display canonical SLe^x^ epitopes, although structure **26** (core 3 SLe^x^) is not recognized by the CD15s antibody (Fig. S2) and can still be present. To further confirm our findings, we analyzed a variety of other H5 HAs (Fig. S5) and used human trachea from another donor (Fig. S6). Again, only the human H3 control and the H5FR bound human tracheal epithelial tissues.

### Crystal structures of 2.3.4.4b H5FR H5 HA in complex with core 3 compounds 25 and 26, core 2 compound 7, as well as avian receptor analog LSTa

To understand the expanded receptor binding repertoire of H5 2.3.4.4b HA, we determined crystal structures of H5FR H5 HA in apo-form (1.94 Å resolution) and in complex with *O*-linked glycan compounds **26** (2.50 Å)**, 25** (1.98 Å) and **7** (2.40 Å), as well as avian receptor analog LSTa (NeuAcα2-3Galβ1-3GlcNAcβ1-3Galβ1-4Glc) (2.90 Å) (Figs. 5, S7, and Table S1).

**Figure 5.**
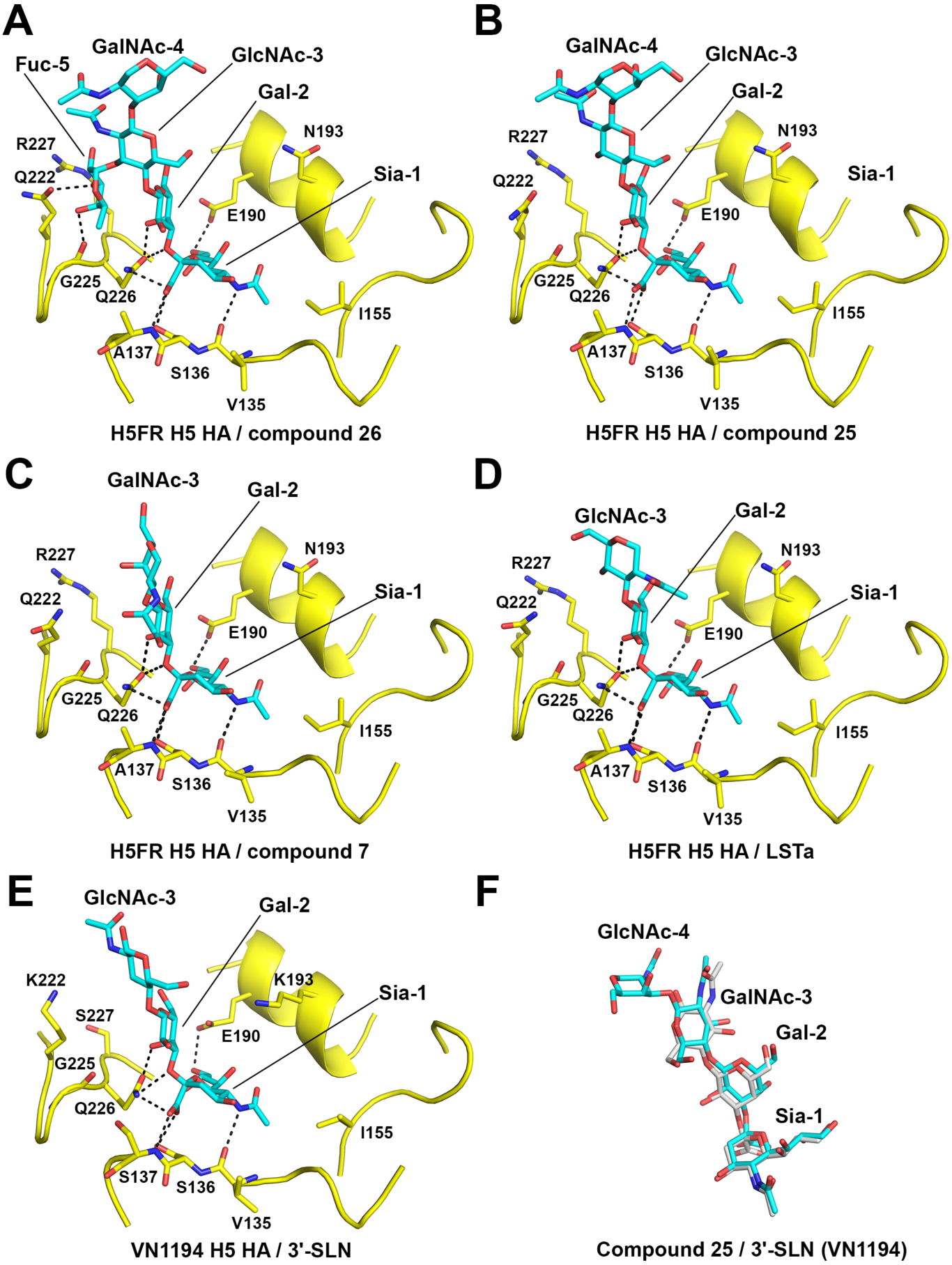
Crystal structures of H5FR H5 HA in complex with glycan compounds 26, 25, and 7, as well as LSTa. (A) core 3 compound **26** bound to H5FR HA. (B) core 3 compound **25** bound to H5FR HA. (C) core 2 compound **7** bound to H5FR HA. (D) Avian receptor analog LSTa bound to H5FR HA. (E) Avian receptor analog 3’-SLN bound to VN1194 H5 HA from A/Vietnam/1194/2004 (H5N1) (PDB ID 4BGY). (F) Superimposition of compound **25** (cyan) and 3’-SLN (grey, from PDB ID 4BGY) in the HA RBS.

**Figure 6.**
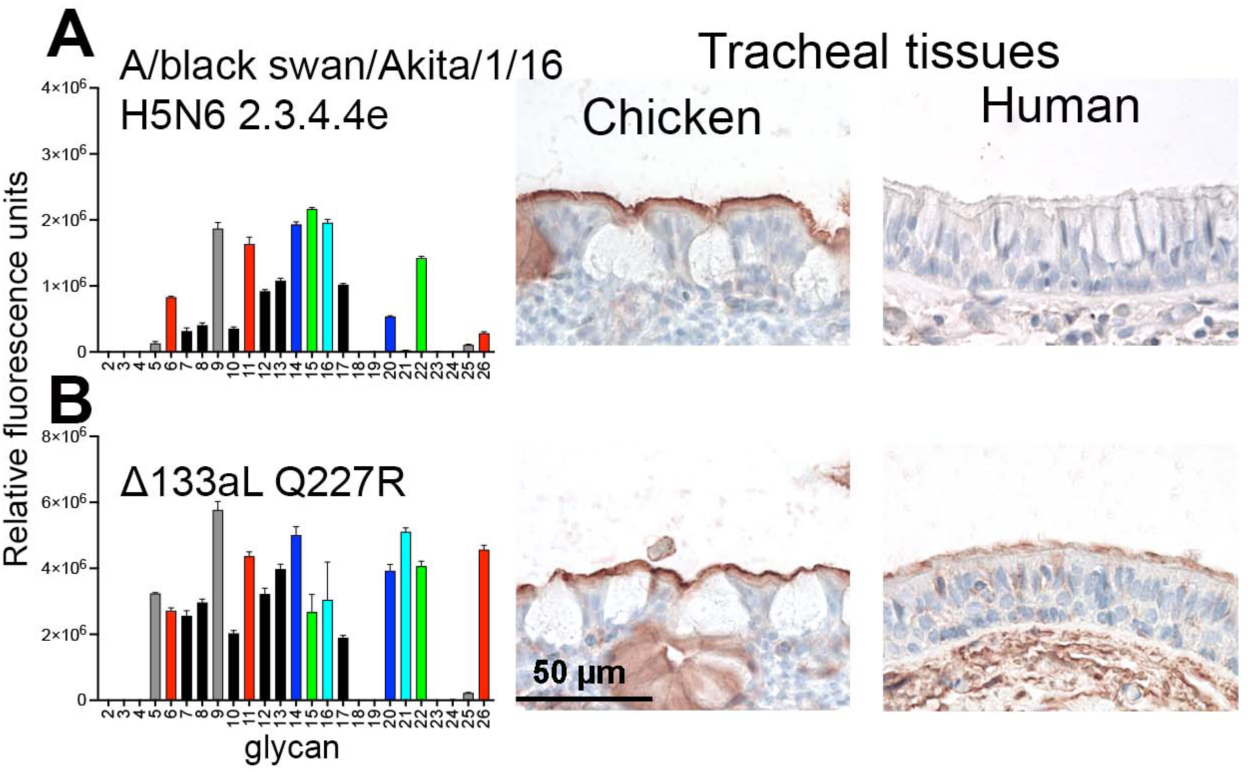
Two amino acid mutations in an H5N6 HA confer binding to human tracheal tissues. (A) Binding of the A/black swan/Akita/1/16 to the O-glycan array as shown in Fig. 2A was investigated. Relative fluorescence units are on the y-axis and glycan numbering on the X-axis as in figure 3A. Shown is a representative of two independent assays, standard deviation from 6 technical replicates with the highest and lowest data points removed. Binding of this HA to chicken and human tracheal tissue was also investigated. AEC staining was used to visualize tissue binding. (B) The same assays were performed with the Δ133aL and Q227R mutant of A/black swan/Akita/1/16.

**Figure 7.**
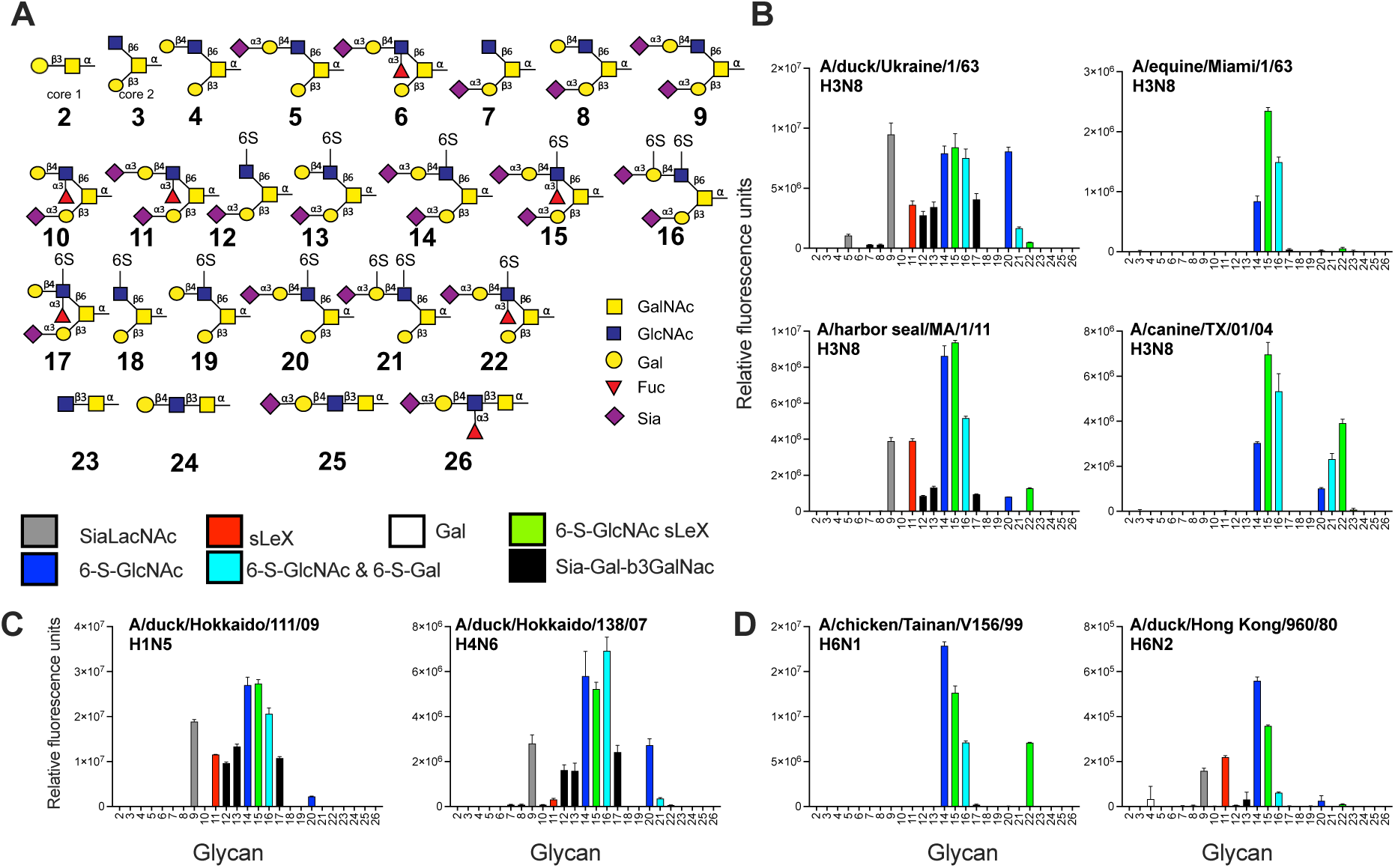
Extended analyses of a variety of influenza A HA subtypes on the *O*-linked glycan array. (A) The shown O-glycans were printed on a glycan microarray and the terminal epitopes are indicated in different colors. (B) The binding of several H3 HAs, (C) H1, H4, and (D) H6 HAs were determined. Relative fluorescence units are on the y-axis and glycan numbering on the X-axis as in (A). Shown is a representative of two independent assays, standard deviation from 6 technical replicates with the highest and lowest data points removed.

To resolve the molecular determinants of SLe^x^ specificity for 2.3.4.4b viruses, we started with compound **26**. Compound **26** shares the same structure with **25** but with an additional fucose 5 (Fuc-5) attached to GlcNAc-3 (Fig. 1, Fig. S7). Indeed, Fuc-5 mediates additional hydrogen bonds with Q222 and the main-chain carbonyl of G225 (Fig. 5A) but not with R227, although it was proposed that Q222 and R227 confer sLe^x^ specificity^32,57^. While G225 is conserved between H5FR and H5VN H5 HAs, H5VN HA has K222 (Fig. S1). These additional interactions between Fuc-5 and H5FR HA truly enhance the binding of **26**; we even observed weak density for the threonine (T3) in peptide linker moiety of the glycopeptide (Fig. S7A).

H5FR H5 HA undergoes little conformational changes upon binding to other glycans; for example, when **25** binds, only very slight movements or rotamer differences of side chains, such as Gln226 and Leu194, are observed in the RBS (Fig. S8). The *O*-linked glycans **25** and **26** share a common terminal sequence SIA 1 (Sia-1), galactose 2 (Gal-2), and N-acetylglucosamine 3 (GlcNAc-3) with the avian receptor analog 3’-SLN (NeuAcα2-3Galβ1-4GlcNAc), while compound **7** shares the first two monosaccharides Sia-1 and Gal-2 with 3’-SLN (Fig. 1). For **25** in complex with H5FR HA, the conformations of Sia-1, Gal-2 and GlcNAc-3 are very similar to 3’-SLN in its complex with VN1194 H5 HA from A/Vietnam/1194/2004 (H5N1) (Fig. 5). VN1194 H5 HA and H5VN HA would share the same receptor specificity as the only amino acid difference between these two HAs is located in the HA stem, and is HA1 T46 (H3 numbering) for VN1194 H5 HA and K46 for H5VN HA^58^. In the H5FR HA–**25** complex, **25** adopts a trans configuration in the Siaα2-3Gal linkage (Fig. 5B), similar to 3’-SLN in its complex with VN1194 H5 HA^59^ (Figs. 5E and 5F). Compound **25** Sia-1 and Gal-2 form hydrogen bonds with RBS residues A136, A137 and E190, while the N-acetylgalactosamine 4 (GalNAc-4 that connects to Thr or Ser in an *O*-linked glycan) projects out of the RBS and the peptide linker residues FVTIG (Phe-Val-Thr-Ile-Gly) are not visible in the electron density map (Fig. 5B, Fig. S7B). Previously, we determined a crystal structure of human Sh2 H7 HA from A/Shanghai/2/2013 (H7N9) with a similar avian *O*-linked glycan, glycan **#21** (NeuAcα2-3Galβ1-4GlcNAcβ1-3GalNAcα-Thr), which shares the same glycan structure with **25** but with Thr only for the peptide linker (Fig. S9)^50^. In contrast, glycan **#21** adopts a cis configuration in the Siaα2-3Gal linkage with Sh2 H7 HA (Fig. S9A), and the receptor exits over the 220 loop, facilitating contacts between GlcNAc-3 and HA Q222 of Sh2 H7 HA (Fig. S9A). The different binding modes of the *O*-linked glycan (Fig. 9B) highlight the diversity in HA-glycan recognition, which is not predictable without determining crystal structures.

Compound **7** is a branched glycan (Fig. 1) with one arm containing Sia-1, Gal-2, and GalNAc-3, and the other arm has only GlcNAc, which is not modeled due to poor electron density (Fig. S7C). Similar to **26** and **25**, the overall binding of **7** to H5FR HA is through Sia-1 and Gal-2, with no substantial glycan-HA interactions visualized for GalNAc-3 (Fig. 5C).

As a positive control, LSTa without GalNAc, a commonly used avian receptor analog for structural and functional studies, was investigated (Fig. 5D). Sia-1, Gal-2 and GlcNAc-3 were ordered in LSTa-H5FR HA complex structure (Figs. 5D & S7D), while in the LSTa complex with VN1194 H5 HA (PDB 3ZP0, 2.51 Å resolution), only Sia-1 and Gal-2 were visible^60^. Structural comparison of H5FR HA with LSTa versus H5FR HA in complex with **26**, **25,** and **7** reveals the similar key interactions between glycan and H5FR HA through Sia-1 and Gal-2 (Fig. 5).

In summary, Fuc-5 of compound **26** directly interacts with the H5FR HA, facilitating stronger binding than compound **25** (Figs 5A and 5B). Compounds **25** and **7** bind H5FR HA mainly through interactions of Sia-1 and Gal-2 (Figs 5B and 5C). RBS residue differences between different H5 HA strains, such as residue 222, as well as residues 137, 193, and 227 that differ between 2.3.4.4 H5 strains and other classical H5 strains (Fig. S1), need to be further interrogated to determine their roles in binding specificity of these O-glycans.

### The peptide linker FVTIG of glycopeptides binds to the H5FR H5 HA stem domain and head domain in the crystal structures

Unexpectedly, when we determined crystal structures of H5FR HA with glycan compounds **25**, **26,** and **7** by soaking H5FR HA crystals in these glycans at 5 mM for 10 minutes, strong additional density was observed in the H5FR HA stem domain for all three glycan complexes and in the head domain with the **25** complex (Fig. S10). The substantial densities were from the binding of these glycopeptides to the H5 HA (Fig. S10) via their peptide linker FVTIG moiety (Fig. 1). The peptide FVTIG in **25**, **26,** and **7** was found to bind H5FR HA stem region with the same constellation of HA residues in each structure (Fig. S10). Among key peptide-HA interactions, the peptide V2 main chain forms two hydrogen bonds with the side chain of HA2 N53, and peptide G5 hydrogen bonds to the side chain of HA1 H38 (Fig. S10). Peptide residue V2 also interacts with HA1 M292 and HA2 V52 and I56, and the peptide I4 makes hydrophobic interactions with HA2 W21, I45 and V48 (Fig. S10). The H5FR HA contacting residues to peptide FVTIG are fully conserved in H5VN H5 HA and overlap with the CR9114 epitope in H5VN H5 HA except for HA2 N53, which does not contact CR9114^61^.

The peptide linker FVTIG in **25** was also found to bind H5FR HA1 head residues K165 to T171 primarily through main-chain hydrogen bond interactions of peptide V2 with HA1 K165 and S167, peptide I4 with HA1 N169, as well as peptide G5 with the HA1 side chain T171 (Fig. S10B). The lack of apparent electron density for peptide FVTIG from **26** and **7** binding to H5FR HA head may be attributed to the weaker binding affinity of the peptide to the HA head than to the HA stem. The potential impact on HA function by peptide binding to HA1 fragment 165-171 needs to be further investigated (Figs. 3 and S10).

In glycan array analysis, the peptide moiety of FVTIG might not directly bind HA due to the method of glycan immobilization. The peptide with a primary amine on one end was immobilized onto NHS ester-activated microarray slides in the glycan array and was not freely available for HA binding. In HA crystal soaking, the FVTIG moieties of glycan compounds are accessible, as we observe in the complex structures (Fig. S10). To estimate the binding affinity of the peptide FVTIG to HA, we conducted surface plasmon resonance (SPR) binding analysis of H5FR HA to core 1 glycan compound **2**. Glycopeptide **2** contains the FVTIG moiety but lacks SIA (Fig. 1), which renders no binding to H5FR HA in our glycan array studies (Fig. 2). In our SPR analysis in which the FVTIG moiety of **2** is accessible, the binding of **2** to H5FR HA was weak as the estimated apparent *K*_D_ value was greater than 650 μM (Fig. S11).

### A/black swan/Akita/1/2016 2.3.4.4e needs two amino acid mutations to confer binding to human tracheal epithelial cells

In the previous glycan array and tissue binding experiments, we employed an H5 HA of the 2.3.4.4e virus A/black swan/Akita/1/16. This virus contains a representative RBS for 2.3.4.4e viruses, with a clade-specific deletion at position L133aΔ and an R227Q mutation (Fig. S1 and S3). This HA did not bind promiscuously as it did not recognize Sia-Gal-β3GalNAc and core 3 SLe^x^ structures efficiently (Fig. 6A) and failed to bind human tracheal epithelial cells. At the same time, it was biologically active, as evidenced by its classical avian-type receptor binding profile on the glycan array and binding to the chicken trachea. The clade-specific L133aΔ and R227Q mutations are directly in the RBS, and we considered whether a double mutant towards the H5FR sequence (Δ133aL / Q227R) would convert to promiscuous glycan binding properties and result in binding to human tracheal tissues. Indeed, the double mutant, Δ133aL / Q227R, was able to bind Sia-Gal-β3GalNAc and core 3 SLe^x^, similar to 2.3.4.4b HA proteins and viruses, and bound human tracheal tissues (Fig. 6B). Thus, distinct amino acids on position 133a and 227 are essential for the receptor binding phenotypes of 2.3.4.4b H5 viruses.

### Sia-Gal-6-S-GlcNAc as a universal receptor for avian HAs

Next, we focused on the *O*-glycan specificities of a wide variety of IAV HAs, as only a handful of studies have previously investigated the involvement of O-glycans as receptors^26^. H3N8 is a subtype that has been able to infect many avian and mammalian species, including humans ^62,63^. We therefore compared the glycan binding specificities of H3N8 isolates from different species: A/duck/Ukraine/1/1963, A/harbor seal/Massachusetts/1/2011, A/equine/Miami/1/1963, and A/canine/Texas/1/2004^22,46^ (Fig. 7B). Again the 6-sulfate on GlcNAc is important as the di-sialosides **14-16** are bound by all H3 proteins (Fig. 7B). Interestingly, both the duck and seal HAs had the broadest receptor binding profile bound to some Sia-Gal-β3GalNAc epitopes, if a sulfate was present (**12**, **13**, and **17**), but with low responsiveness. Both HAs also displayed binding to a non-modified Sia-LacNAc (**9**) and SLe^x^ (**11**). The equine and canine strains displayed more restricted binding profiles by not binding any Sia-Gal-β3GalNAc epitope. The canine isolate is more promiscuous than the equine as it can bind structures **20-22,** which are the same as **14-16,** except for the additional Sia in the Sia-Gal-β3GalNAc epitope. Analyzing two other IAV HAs of the H1 (A/duck/Hokkaido/111/2009 H1N5) and H4 (A/duck/Hokkaido/138/2007 H4N6) subtypes further consolidate the preference for 6S-GlcNAc containing structures, such as **14-16,** as these displayed the highest responsiveness (Fig. 6C). These receptor binding profiles are more similar to classical H5s and A/duck/Ukraine/1/1963, as some Sia-Gal-β3GalNAc epitopes are bound with a sulfated non-sialylated β6-arm (**12, 13**, and **17**). Furthermore, non-modified Sia-LacNAc (**9**) was bound as well. Next, we focused on two H6 isolates described with differential specificity to 6-sulfated GlcNAc structures^35^. The A/chicken/Tainan/V156/1999 H6N1 HA entirely depended on the presentation of sulfated sLe^x^ in di-sialosides (Fig. 6D), while the A/duck/Hong Kong/960/1980 H6N2 HA was able to bind non-modified Sia-LacNAc (**9**) and SLe^x^ (**11**). In conclusion, the receptor binding properties of 2.3.4.4 H5 viruses are unique in their promiscuity.

## Discussion

In this study, we created an O-glycan array by combining chemical and enzymatic synthesis. We synthesized core 1, 2, and 3 O-glycans encompassing various sialylated LacNAc structures with modifications such as sulfates and fucosides. We discovered that the Sia-Gal-β3GalNAc epitope in core 2 O-glycans and core 3 SLe^x^ are bound by contemporary 2.3.4.4b H5Nx viruses, whereas all other HAs tested failed to do so. The distinct *O*-glycan specificity of the widespread 2.3.4.4 H5Nx viruses may be relevant for the increased host range and the observed gain of binding to human tracheal tissues.

Our findings on the binding of distinct *O*-glycans emphasize the need to develop new methods to create *O*-glycans. Reported enzymes for extending O-glycans are often of microbial origin as they are generally easier to express than mammalian enzymes. However, microbial glycosyltransferases that operate early in the O antigen pathway require sugar-diphosphate-lipids as acceptors^38^, which were unsuitable in our synthesis since we aimed to synthesize glycopeptides that were printable on the glycan microarray slides. For instance, the microbial transferase Wbwc can add a galactose on a GalNAc-α-diphosphate-lipid acceptor but not on GalNAc-α-peptides, and we could not utilize this enzyme. Instead, we used the corresponding mammalian enzyme C1GalT1, which acts specifically on GalNAc-α-peptides.

O-glycans are present in several glycan arrays that evaluate IAV receptor binding^47,57,64,65^. However, these are often not discussed in detail, and it is difficult to get a proper overview of IAV receptor binding specificities to these glycan cores^29^. Here, we show the importance of investigating O-glycans as potential receptors for IAV. For contemporary circulating H5N1 viruses, we not only demonstrated increased binding to O-glycans on the array, but we also observed binding of H5 HAs to human tracheal tissue, not only of H5FR but also for H5AK with mutations Δ133aL and Q227R. Similar human respiratory tissue binding properties have been described previously^54,66^. Indicating that binding to human tracheal epithelial cells can also be due to binding α2,3 sialosides on O-glycans.

The Sia-Gal-β3GalNAc structures, which we show to be bound by 2.3.4.4b viruses, are uniquely found in *O*-linked glycans^29,50,51^, whereas SLe^x^ epitopes are also found in *N*-linked structures. The ability of some H5 viruses to bind sLe^x^ has been shown previously, but it was thought not to be essential^57^. Here, we demonstrate that core 3 O-glycans (**25** and **26**) are unique as they are only bound by 2.3.4.4b H5Nx viruses, indicating that the β3-linkage to the core GalNAc is essential. H5FR H5 HA from clade 2.3.4.4b H5N8 virus binds to glycans **25** and LSTa similarly to VN1194 HA or H5VN HA from clade 1 H5N1 viruses, and subtle amino acid differences around RBS and beyond may determine the binding specificity to O-glycans, such as Q222 in H5FR HA that might be critical for its enhanced binding to compound **26** (Fig. 3). This specific O-glycan is present in the human respiratory tract^68^, but how it precisely relates to the zoonotic properties of 2.3.4.4b H5Nx viruses remains unknown.

There have been few changes in the RBS of 2.3.4.4b H5Nx viruses over the past years. Notably, several hallmark mutations that are described that confer avian- to human-type receptor specificity with E190D and G225D for H1 and Q226L and G228S for H3^68^, are not present in contemporary H5 viruses. However, the introduction of the H3N2 pandemic came with additional amino acid mutations^69^, and literature suggests that H5 proteins also need three or more amino acid mutations to bind human-type receptors^56,70–72^. We recently showed that 2.3.4.4e only needs the single Q226L mutation to bind human-type receptors on linear structures^74^. It has also been recently demonstrated that the A/Texas/37/2024 virus HA protein, identical to the virus used in this study (except for the multibasic cleavage site), only requires the single Q226L mutant to bind human-type receptors presented on complex N-glycans^74^. Human-type receptor binding was specific to branched poly-LacNAc containing symmetrical N-glycans, rare in the human upper respiratory tract ^29,75^. Significantly, however, all previous studies have been limited in their use of biologically relevant O-glycans.

In conclusion, synthesizing biologically relevant O-glycans with diverse LacNAc modifications led us to the discovery that the current 2.3.4.4 H5Nx influenza A viruses tolerate a variety of sialylated glycans. Thus, while these viruses maintain an overall avian-type receptor specificity, their apparent zoonotic properties in mammalian species can be explained by their novel O-glycan specificities.

## Materials and methods

### Synthesis of glycopeptides

The synthesis of the O-glycans is described in the supplementary file.

### Expression and purification of trimeric influenza A hemagglutinins

Recombinant trimeric IAV hemagglutinin (HA) ectodomain proteins were cloned into the pCD5 expression vector (an example is Addgene plasmid #182546^46^) in frame with a GCN4 trimerization motif (KQIEDKIEEIESKQKKIENEIARIKK), a super folder GFP^76^ or mOrange2^78^ and the Twin-Strep-tag (WSHPQFEKGGGSGGGSWSHPQFEK); IBA, Germany). Mutations in HAs were generated by site-directed mutagenesis. The trimeric HAs were expressed in HEK293S GnTI(-) cells with polyethyleneimine I (PEI) in a 1:8 ratio (µg DNA:µg PEI) for the HAs as previously described^79^. The transfection mix was replaced after 6 hours by 293 SFM II suspension medium (Invitrogen, 11686029), supplemented with sodium bicarbonate (3.7 g/L), Primatone RL-UF (3.0 g/L, Kerry, NY, USA), glucose (2.0 g/L), glutaMAX (1%, Gibco), valproic acid (0.4 g/L) and DMSO (1.5%). According to the manufacturer’s instructions, culture supernatants were harvested five days post-transfection and purified with sepharose strep-tactin beads (IBA Life Sciences, Germany).

### Glycan microarray binding studies

HAs (50 µg/ml) were pre-complexed with human anti-streptag and goat anti-human-Alexa555 (#A21433, Thermo Fisher Scientific) antibodies in a 4:2:1 molar ratio, respectively, in 50 µL PBS with 0.1% Tween-20. Biotinylated lectins (5 µg/ml) were pre-complexed with streptavidin-Alexa555 (#S32355, Thermo Fisher Scientific) in a 5:1 weight ratio. The following biotinylated lectins from Vector Laboratories were used: MAL-I (B-1315-2), MAL-II (B-1265-1), ECA (B-1145-5), Jacalin (B-1155-5), PNA (B-1075-5), Ricin (B-1085-5), and AAL (B-1395-1). Siglec-9 (50µg/mL) was precomplexed with pA-LS at a 1:1 mass ratio. The mixtures were incubated on ice for 15 minutes and then on the array surface for 90 minutes in a humidified chamber. Then, slides were rinsed successively with PBS-T (0.1% Tween-20), PBS, and deionized water. For the anti-sialyl-Lewis^x^ antibody CD15S (clone CSLEX1, #551344, BD Biosciences), a solution of 50 µg/ml in 40 µL PBS with 0.1% Tween-20 was first incubated on the slide for 90 min in a humidified chamber. After washing successively with PBS-T (0.1% Tween-20), PBS, and deionized water, a mixture of 10 µg/ml goat anti-mouse IgM-HRP (#1021-05, Southern Biotech) and 5 µg/ml donkey anti-goat IgG-Alexa555 (#A21432, Thermo Fisher Scientific) in 40 µL PBS with 0.1% Tween-20 was incubated on the slide for 90 min in a humidified chamber. Afterward, the slides were rinsed successively with PBS-T (0.1% Tween-20), PBS, and deionized water. The arrays were dried by centrifugation and immediately scanned as described previously^46^. Processing of the six replicates was performed by removing the highest and lowest replicates and subsequently calculating the mean value and standard deviation over the four remaining replicates.

### Expression, crystallization, and crystal structure determination of H5FR H5 HA and its glycan complexes

The ectodomain of H5FR H5 HA was expressed in a baculovirus system essentially as previously described^79,80^. Briefly, the cDNA corresponding to residues 11-327 of HA1 and 1-174 of HA2 (H3 numbering) of wildtype H5FR H5 HA from A/duck/France/161108h/2016 (H5N8) (GISAID accession number EPI869809) was incorporated into a baculovirus vector, pFastbacHT-A (Invitrogen) with an N-terminal gp67 signal peptide, a C-terminal foldon trimerization domain, and a His_6_-tag, as well as a thrombin cleavage site between the HA ectodomain and the foldon trimerization domain. The constructed plasmid was used to transform DH10bac competent bacterial cells, and the purified recombinant bacmids were used to transfect Sf9 insect cells for overexpression. HA protein was produced in suspension cultures of High Five insect cells. Soluble HA protein was harvested in supernatants and purified by metal affinity chromatography using Ni-nitrilotriacetic acid (Ni-NAT) resin (Qiagen). For crystallization, the foldon trimerization domain and His_6_-tag were cleaved from the HA ectodomain with thrombin, and the HA ectodomain was purified further by size-exclusion chromatography on a Hiload 16/90 Superdex 200 column (GE Healthcare) in 20 mM Tris, pH 8.0, 150 mM NaCl and 0.02% NaN_3_.

Crystallization experiments were set up using the sitting drop vapor diffusion method on our robotic CrystalMation system (Rigaku). For structure determination of the apo-form H5FR H5 HA and its complex with glycan compounds **7** and **25**, purified protein at 11.1 mg/ml was crystallized in 0.1 M Tris, pH 7.0, 0.11 M lithium sulfate, and 2.0 M ammonium sulfate at 20 °C. For structure determination of the H5FR HA with compound **26**, the purified HA at 11.1 mg/ml was crystallized in 0.1 M sodium citrate, pH 5.6, 0.2 M potassium sodium tartrate, and 2.0 M ammonium sulfate at 20 °C. For structure determination of the H5FR HA in complex with compound LSTa (NeuAcα2-3Galβ1-3GlcNAcβ1-3Galβ1-4Glc), purified H5FR HA at 11.1 mg/ml was crystallized in 1.6 M sodium citrate, pH 6.5 at 20 °C. H5FR HA-ligand complexes were obtained by soaking HA crystals in the well solution containing ligands and cryo-protectants. The final concentrations of compounds **7**, **25**, **26,** and LSTa were all 5 mM, and the soaking times were all about 10 min. The crystals were flash-cooled at 100K in mother liquid, and 15% ethylene glycol (v/v) was added as a cryoprotectant for apo-form H5FR HA, and its complex with compounds **7** and **25**, and 20% glycerol (v/v) was added as a cryoprotectant for H5FR HA in complex with compounds **26** and LSTa.

Diffraction data were collected at Stanford Synchrotron Radiation Lightsource (SSRL) beamline 12-1 (**Table S1**). Data were integrated and scaled with HKL3000 ^82^. Data and refinement statistics are summarized in **Table S1**.

The crystal structure of H5FR H5 HA was determined by molecular replacement (MR) using the program Phaser^83^ with the crystal structure of H5 HA from A/duck/Eastern China/L0230/2010 (H5N2) (PDB code 7DEB) as input MR model. Initial rigid body refinement was performed in REFMAC5^84^, and further restrained refinement, including TLS refinement, was carried out in Phenix^84^. The model building between rounds of refinement was carried out with Coot^86^. The final statistics are summarized in **Table S1**. The structure quality was analyzed using the JCSG validation suite (qc-check.usc.edu/QC/qc_check.pl) and MolProbity^86^. All structural figures were generated with PyMol (www.pymol.org).

### Surface Plasmon Resonance (SPR) binding analysis of H5FR H5 HA and glycan 2

SPR was conducted on BiaCore S200 instrument (Cytiva Inc.). High-density poly-NTA HC hydrogel chips (NiHC1500M, XanTec) were used to capture polyhistidine-tagged HA protein. The protein immobilization level reached more than 20,000 RU in the sample flow cell. All experiments were performed at 25 °C, with PBS, pH 7.4 as the running buffer. 1 mM of glycan **2** solution was prepared by more than 100 times dilution from stock solution with running buffer. Further sample concentrations were prepared by serial dilution four times from a 1 mM solution. Samples were injected over the reference paired 2 flow cells for 120 s followed by 240 s dissociation time. Five sample concentrations were injected, with 3 cycles of zero concentration in between to remove residual samples left on the chip surface or protein pocket. All signals were reference corrected, and the negative sample was subtracted before analysis. Signals from 5 s before injection stop were used for steady-state analysis.

### Protein histochemical tissue staining

Sections of formalin-fixed, paraffin-embedded chicken (*Gallus gallus domesticus*) were obtained from the Division of Pathology, Department of Biomolecular Health Sciences, Faculty of Veterinary Medicine of Utrecht University, the Netherlands. Sections of formalin-fixed, paraffin-embedded human tissues were obtained from the UMC Utrecht, Department of Pathology, Utrecht, the Netherlands (TCBio-number 22-599). In the figures, representative images of at least two individual experiments are shown. Protein histochemistry was performed as previously described^87,88^. In short, tissue sections of 4 µm were deparaffinized and rehydrated, after which antigens were retrieved by heating the slides in 10 mM sodium citrate (pH 6.0) for 10 min. Endogenous peroxidase was inactivated using 1% hydrogen peroxide in MeOH for 30 min at RT. When neuraminidase treatment was performed, slides were incubated overnight at 37°C with neuraminidase from *Vibrio cholerae* (#11080725001, Roche) diluted 1:50 in a solution of 10mM potassium acetate and 0.1% Triton X-100 at pH 4.2. Non-treated slides in experiments with neuraminidase were incubated with only buffer. Tissues were blocked at 4°C using 3% BSA (w/v) in PBS for at least 90 minutes. For plant lectin stains, blocking was performed using carbo-free blocking solution (SP-5040-125; Vector Laboratories, Burlingame, CA, USA) instead. Subsequently, slides were stained for 90 minutes using anti-sLe^x^ antibody or pre-complexed HAs as previously described for the glycan microarray. For A/duck/France/161108h/2016, 2 µg/ml of H5 HA was used, while for the other H5 HAs, 2.5 µg/ml was used. For A/NL/109/2003, 5.0 µg/ml H3 HA were used. For antibody control images, only antibodies and no HAs have been used. The detection of sLe^x^ was performed with the anti-sLe^x^ antibody CD15S (clone CSLEX1, #551344, BD Biosciences), which was diluted 1:1000 in PBS and precomplexed with goat anti-mouse IgM-HRP (#1021-05, Southern Biotech) in a 1:100 dilution. Siglec 9 was precomplexed with an HRP-conjugated goat-anti-human antibody at a 2:1 molar ratio.

### H5 HA phylogenetic trees

We downloaded all available influenza H5Nx HA sequences and metadata that were available up until 1 February 2025 from the Global Initiative on Sharing All Influenza Data (GISAID) (https://gisaid.org/) EpiFlu database. Low-quality (>1% ambiguous nucleotide) and incomplete sequences (<95% of HA segment length) were removed. Besides the viruses studied in this work, we randomly subsampled 5-20 representative sequences per H5 clade as assigned by GISAID. We aligned sequences using MAFFT v7.520^90^, and reconstructed the maximum-likelihood phylogenetic tree with IQ-TREE v2.2.2.6^91^, using the general time reversible (GTR) model. We then reconstructed the temporally resolved phylogenies using TreeTime v0.11.4^92^. We used Baltic (https://github.com/evogytis/baltic/tree/master) python library to produce all tree visualizations.

## Supporting information

Spectra

## Data Availability Statement

The authors confirm that the data supporting the findings of this study are available within the paper. The atomic coordinates and structure factors are being deposited in the Protein Data Bank (PDB) under accession codes 9NRR, 9NRS, 9NRT, 9NRU, and 9NRV for H5FR H5 HA apo-form, and in complex with compounds **26**, **25**, **7,** and LSTa, respectively.

## Acknowledgments

JW is supported by the Austrian Science Fund (FWF, J-4260-B21). This research was made possible by funding from ICRAD, an ERA-NET co-funded under the European Union’s Horizon 2020 research and innovation program. (https://ec.europa.eu/programmes/horizon2020/en), under Grant Agreement n°862605 (Flu-Switch) to RPdV. The glycan array setup was supported by the Netherlands Organization for Scientific Research (NWO, TOP-PUNT 718.015.003 to G.-J. B.). Dr. Margreet A. Wolfert (Utrecht University) developed, printed, and validated the glycan microarray. This research was partially funded by the National Institutes of Health NIAID Centers of Excellence for Influenza Research and Response contract 75N93021C00015 / PENN CEIRR (I.A.W.)

We thank Takahiro Hiono, Masatoshi Okamatsu, Yoshihiro Sakoda (Hokkaido University), and Romain Volmer (Université de Toulouse) for generously sharing reagents.

We gratefully acknowledge all data contributors, i.e., the authors and their originating laboratories responsible for obtaining the specimens and their submitting laboratories for generating the genetic sequence and metadata and sharing via the GISAID Initiative, on which part of this research is based.

We thank Henry Tien for automated robotic crystal screening. The SSRL is a Directorate of SLAC National Accelerator Laboratory and an Office of Science User Facility operated for the U.S. Department of Energy Office of Science by Stanford University. The SSRL Structural Molecular Biology Program is supported by the DOE Office of Biological and Environmental Research, and by the National Institutes of Health, National Institute of General Medical Sciences (including P41GM103393) and the National Center for Research Resources (P41RR001209).

