## Supplemental information for "The receptor binding properties of H5Nx influenza A viruses have evolved to promiscuously bind to avian-type mucin-like O-glycans"

#### This file includes:

Supplementary tables and figures……………………………………………S2
General Remarks………………………………………………………………… S16
Chemical synthesis and schemes…………………………………………..S17
Enzymatic synthesis………………………………………………………………S27
References……………………………………………………………………… S42
Spectra………………………………………………………………………...S43

### Supplementary tables and figures

#### Table S1. Data collection and refinement statistics

| Dataset | **H5FR HA** | **H5FR HA** | **H5FR HA** |
|---|---|---|---|
| | ***apo*** | **+ compound 26** | **+ compound 25** |
| **Data Collection** | | | |
| X-ray source | SSRL 12-1 | SSRL 12-1 | SSRL 12-1 |
| Wavelength (Å) | 0.97946 | 0.97946 | 0.97946 |
| Space group | $P2_12_12_1$ | $P2_12_12_1$ | $P2_12_12_1$ |
| Unit cell (Å) | $a$ = 95.4, | $a$ = 95.3 | $a$ = 95.2 |
| | $b$ = 171.6, | $b$ = 171.9, | $b$ = 171.6, |
| | $c$ = 226.0 | $c$ = 225.6 | $c$ = 226.4 |
| Resolution (Å)[a] | 45.56-1.94 (1.97-1.94) | 49.12-2.50 (2.54-2.50) | 43.87-1.98 (2.01-1.98) |
| Unique reflections[a] | 270,800 (13,249) | 127,703 (6,285) | 254,864 (11,594) |
| Redundancy[a] | 9.8 (6.7) | 7.4 (7.6) | 6.7 (5.5) |
| Average $I/\sigma(I)$[a] | 20.9 (1.0) | 9.1 (0.9) | 16.1 (0.9) |
| Completeness (%)[a] | 99.9 (99.1) | 99.9 (100) | 98.7 (90.5) |
| $R_{sym}$[a,b] | 0.14 (>1.0) | 0.20 (>1.0) | 0.12 (>1.0) |
| $R_{pim}$[a,b] | 0.05 (0.60) | 0.08 (0.70) | 0.05 (0.67) |
| $CC_{1/2}$[a] | 0.996 (0.538) | 0.985 (0.420) | 0.995 (0.459) |
| No. molecules per ASU[c] | 3 | 3 | 3 |
| **Refinement** | | | |
| Resolution (Å)[a] | 45.56-1.94 (1.97-1.94) | 49.12-2.50 (2.53-2.50) | 43.87-1.98 (2.00-1.98) |
| Reflections in refinement | 270,603 | 127,534 | 254,641 |
| Refined residues | 1,476 | 1,497 | 1,496 |
| Refined waters | 1,810 | 665 | 1,793 |
| $R_{cryst}$[a,d] | 0.167 (0.329) | 0.174 (0.273) | 0.165 (0.308) |
| $R_{free}$[a,e] | 0.188 (0.358) | 0.208 (0.322) | 0.189 (0.313) |
| $B$-values (Å$^2$) | | | |
| Protein | 37 | 52 | 38 |
| Ligand | - | 63 | 53 |
| Water | 49 | 52 | 49 |
| Wilson $B$-values (Å$^2$) | 30 | 42 | 32 |
| Ramachandran values (%)[f] | 97.6, 0 | 96.4, 0.3 | 97.8, 0 |
| r.m.s.d. bond (Å) | 0.007 | 0.003 | 0.007 |
| r.m.s.d. angle (deg.) | 0.84 | 0.56 | 0.86 |
| PDB codes | 9NRR | 9NRS | 9NRT |

**Table S1. Data collection and refinement statistics - continued**

| Data set | **H5FR HA + compound 7** | **H5FR HA + LSTa** |
|---|---|---|
| **Data Collection** | | |
| X-ray source | SSRL 12-1 | SSRL 12-1 |
| Wavelength (Å) | 0.97946 | 0.97946 |
| Space group | $P2_12_12_1$ | $P2_12_12_1$ |
| Unit cell (Å) | $a$ = 95.7<br>$b$ = 172.0,<br>$c$ = 226.8 | $a$ = 95.0<br>$b$ = 177.0,<br>$c$ = 224.6 |
| Resolution (Å)[a] | 46.12-2.40 (2.44-2.40) | 48.97-2.90 (2.97-2.90) |
| Unique reflections[a] | 148,008 (7,295) | 85,027 (5,581) |
| Redundancy[a] | 8.5 (6.9) | 6.1 (6.1) |
| Average $I/\sigma(I)$[a] | 17.1 (1.0) | 5.5 (0.7) |
| Completeness (%)[a] | 99.9 (100) | 99.9 (99.9) |
| $R_{sym}$[a,b] | 0.16 (>1.0) | 0.27 (>1.0) |
| $R_{pim}$[a,b] | 0.06 (0.61) | 0.12 (0.80) |
| $CC_{1/2}$[a] | 0.990 (0.456) | 0.968 (0.402) |
| No. molecules per ASU[c] | 3 | 3 |
| **Refinement** | | |
| Resolution (Å)[a] | 46.12-2.40 (2.43-2.40) | 48.97-2.90 (2.92-2.90) |
| Reflections in refinement | 147,849 | 84,890 |
| Refined residues | 1,495 | 1,482 |
| Refined waters | 1,090 | 183 |
| $R_{cryst}$[a,d] | 0.182 (0.293) | 0.178 (0.291) |
| $R_{free}$[a,e] | 0.206 (0.327) | 0.218 (0.321) |
| $B$-values (Å$^2$) | | |
| Protein | 45 | 61 |
| Ligand | 80 | 66 |
| Water | 54 | 53 |
| Wilson $B$-values (Å$^2$) | 41 | 53 |
| Ramachandran values (%)[f] | 97.1, 0.2 | 94.1, 0.3 |
| r.m.s.d. bond (Å) | 0.003 | 0.009 |
| r.m.s.d. angle (deg.) | 0.54 | 1.02 |
| PDB codes | 9NRU | 9NRV |

[a] Parentheses denote outer-shell statistics.
[b] $R_{sym} = \sum_{hkl}\sum_i |I_{hkl,i} - \langle I_{hkl}\rangle| / \sum_{hkl}\sum_i I_{hkl,i}$ and $R_{pim} = \sum_{hkl}[1/(N-1)]^{1/2}\sum_i |I_{hkl,i} - \langle I_{hkl}\rangle| / \sum_{hkl}\sum_i I_{hkl,i}$, where $I_{hkl,i}$ is the scaled intensity of the i$^{th}$ measurement of reflection $h$, $k$, $l$, $\langle I_{hkl}\rangle$ is the average intensity for that reflection, and $N$ is the redundancy. $R_{pim} = \Sigma_{hkl} (1/(n-1))^{1/2} \Sigma_i | I_{hkl,i} - \langle I_{hkl}\rangle | / \Sigma_{hkl} \Sigma_i I_{hkl,i}$, where n is the redundancy
[c] No. molecules for complexes refers to number of HA protomers per asymmetric unit (ASU), i.e. an HA trimer is in the ASU.
[d] $R_{cryst} = \sum_{hkl} |F_o - F_c| / \sum_{hkl} |F_o|$, where $F_o$ and $F_c$ are the observed and calculated structure factors.
[e] $R_{free}$ was calculated as for $R_{cryst}$, but on 5% of data excluded before refinement.
[f] The values are percentage of residues in the favored and outliers regions analyzed by MolProbity [1].

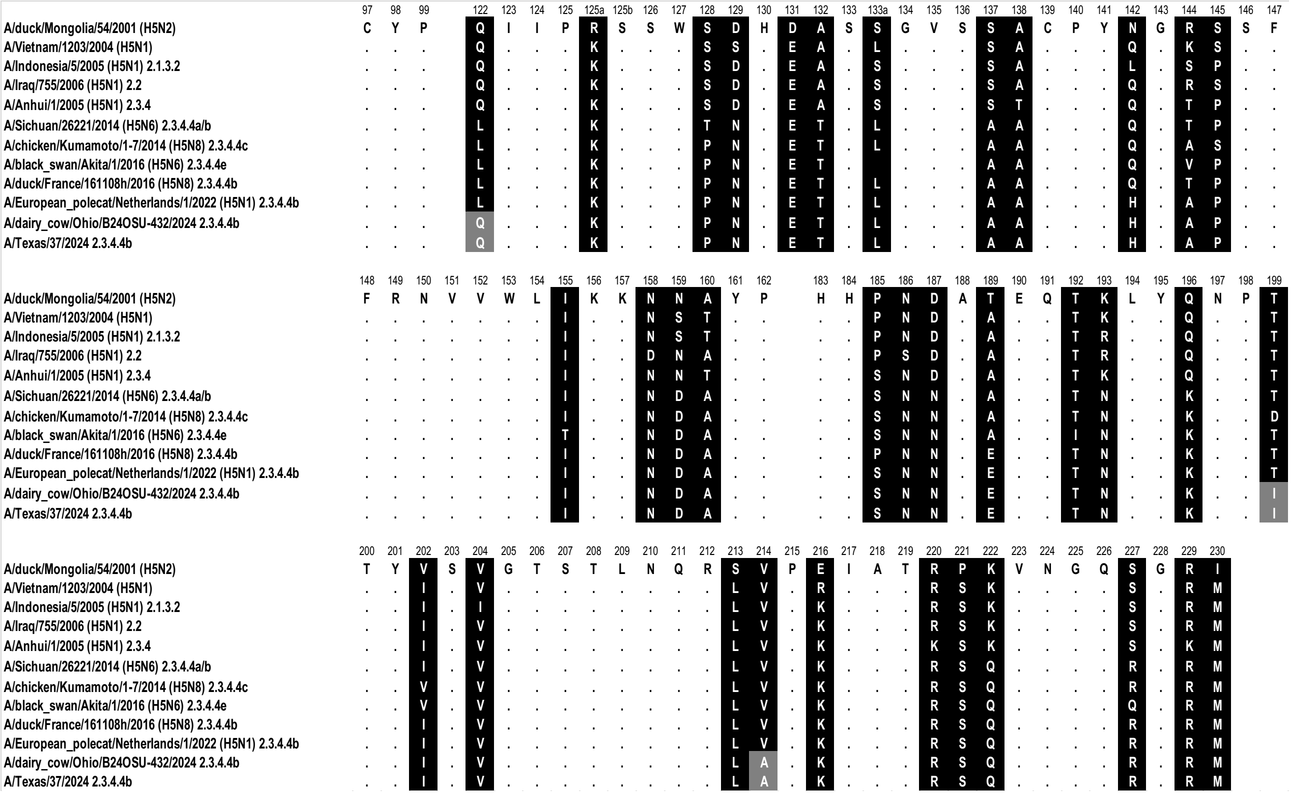

**Figure S1. HA RBS amino acid alignment of the H5 hemagglutinins used in this study.** Alignment of the RBS residues with amino acid positions (H3 numbering) indicated above the alignment, non-conserved residues highlighted in a black background, dots indicating identical amino acids. 2024, cattle outbreak specific amino acid changes, are highlighted with a gray background.

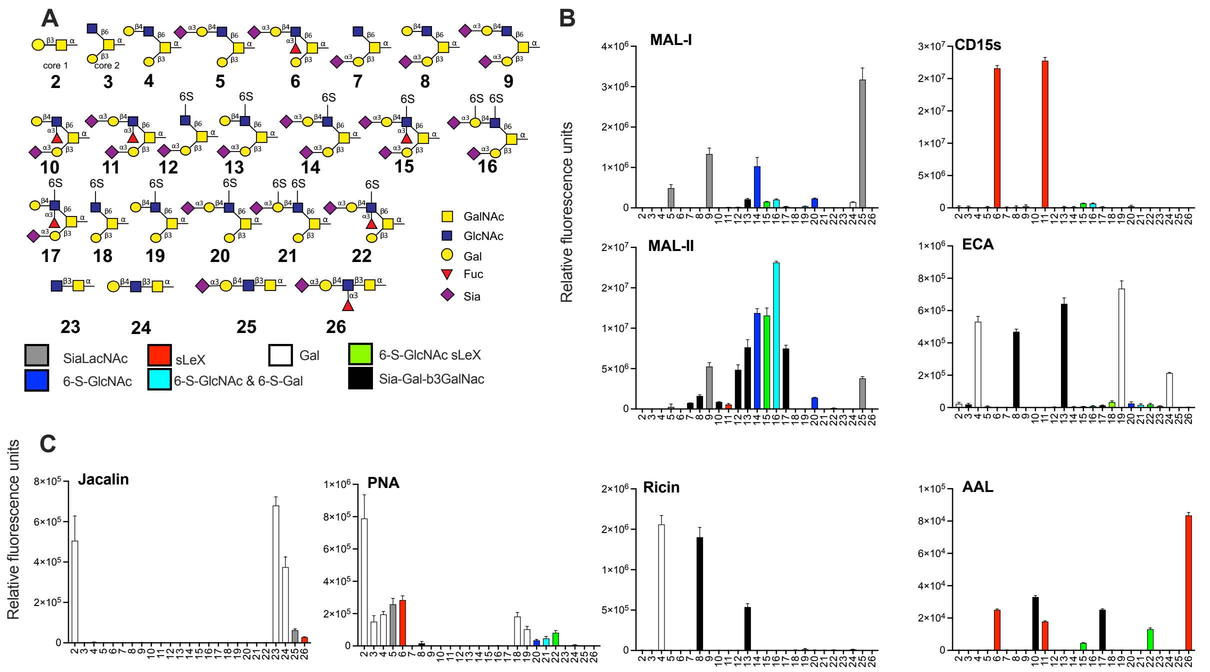

**Figure S2. O-glycan characterization using plant lectins.** (A) The O-glycans were printed on a glycan microarray, and the terminal epitopes were indicated in different colors. (B) The binding of the plant lectins Jacalin (binds to terminal T-antigen (Gal-β1,3-GalNAc), [2]), PNA (peanut agglutinin, binds to binds terminal T-antigen (Gal-β1,3-GalNAc) [3]), Ricin (binds to terminal type 2 LacNAc [4]), and AAL (*Aleuria aurantia* lectin, binds to fucose [5]).

79

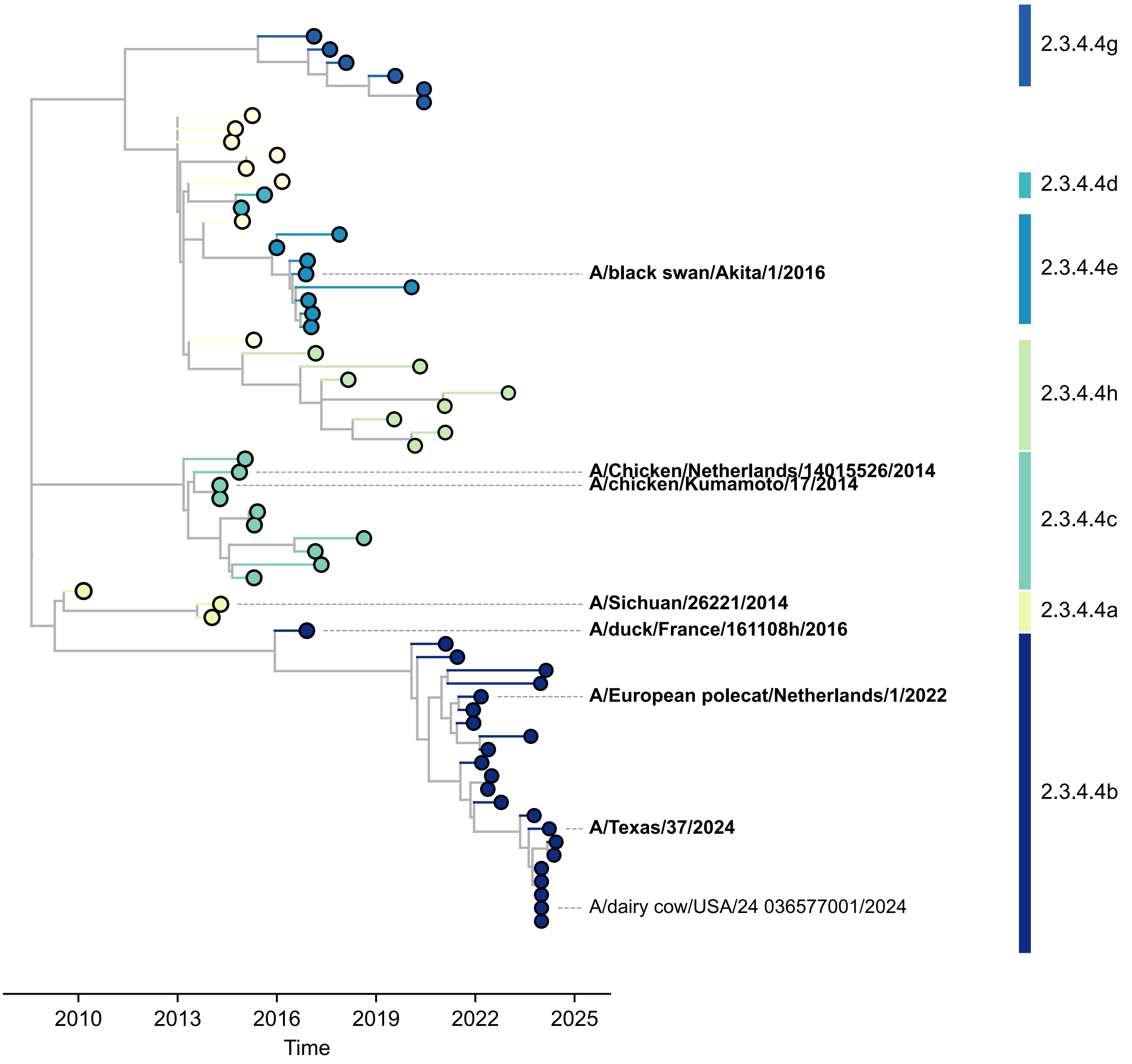

**Figure S3. Phylogenetic tree of 2.3.4.4.b H5 HAs.** The viruses that were studied in this work are indicated.

80

81

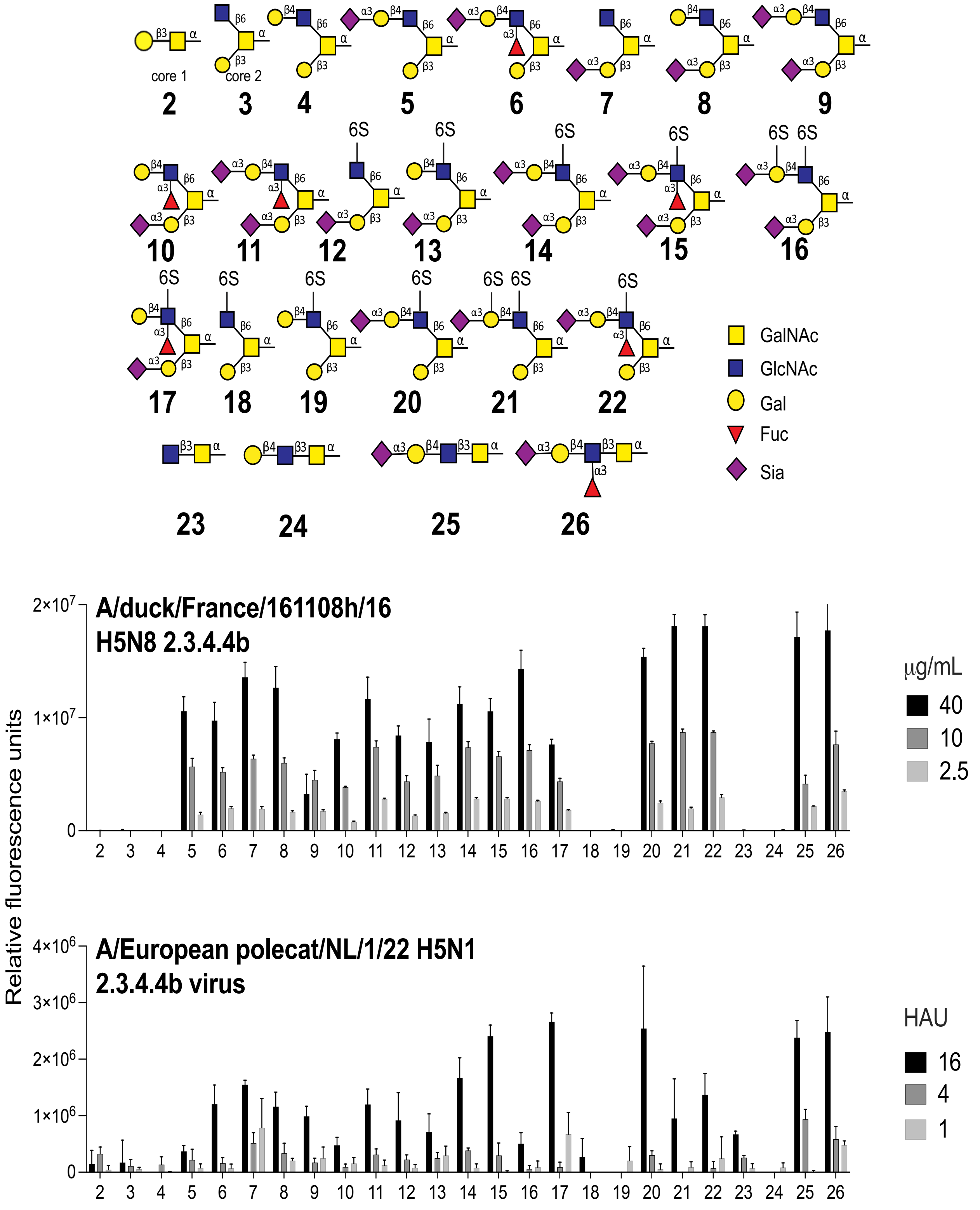

**Figure S4. Titration of HA and virus on the glycan microarray.** Top panel is the O-linked glycan ligands on the array and bottom panels are the binding of these ligands to both HA and virus titrated from 40 to 2,5 µg/mL and 16 to 1 hemagglutination units. For reference, in Fig. 2, 50 µg/mL and 32 HAU units were used respectively.

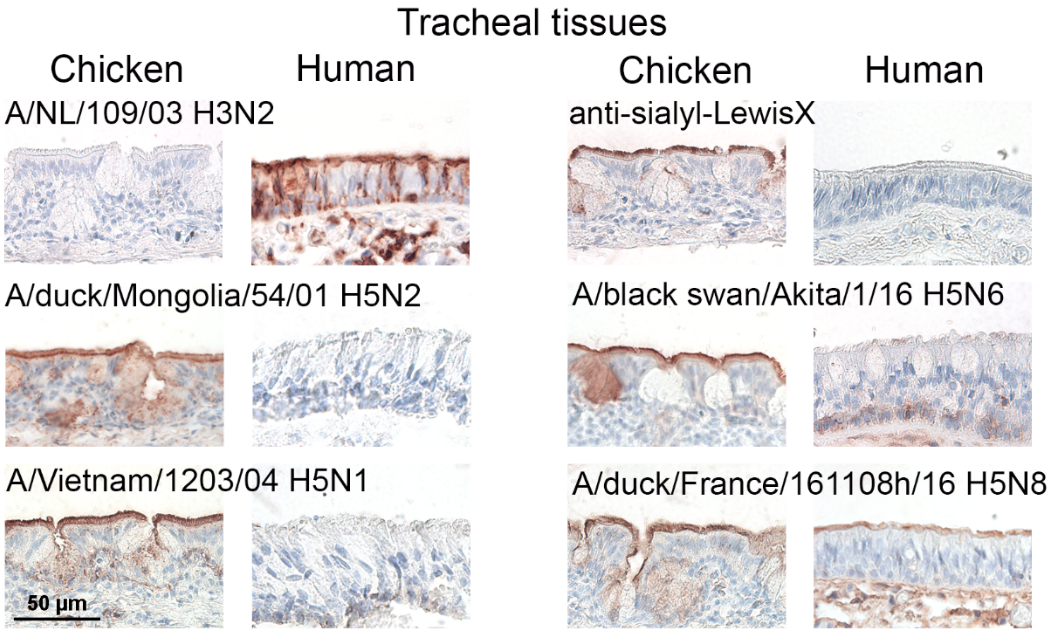

**Figure S5. The HA of A/duck/France/161108h/2016 H5N8 is the only H5 HA that binds human tracheal tissue.** The binding to human and chicken tracheal tissue was investigated for different influenza A H5 HAs. The HA from A/NL/109/2003 was used as a positive control for human trachea binding. The antibody CD15S was used to visualize sialyl-LewisX epitopes. AEC staining was used to visualize tissue binding.

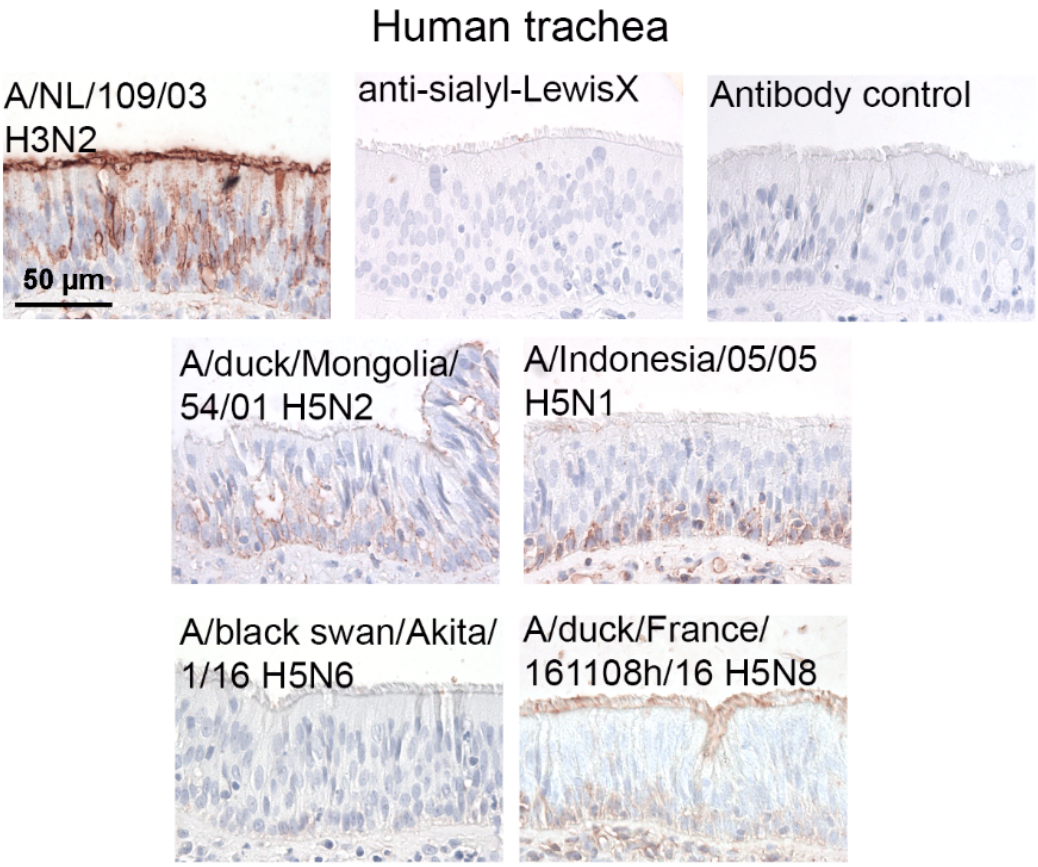

**Figure S6. Tissue binding analyses with additional human donor trachea.** Binding experiments with several influenza H5 HAs, the anti-sialyl-LewisX antibody, and human tracheal tissues (from a different patient than in Fig. 4) were performed. The H3 HA from A/NL/109/2003 was used as a positive control. AEC staining was used to visualize tissue binding.

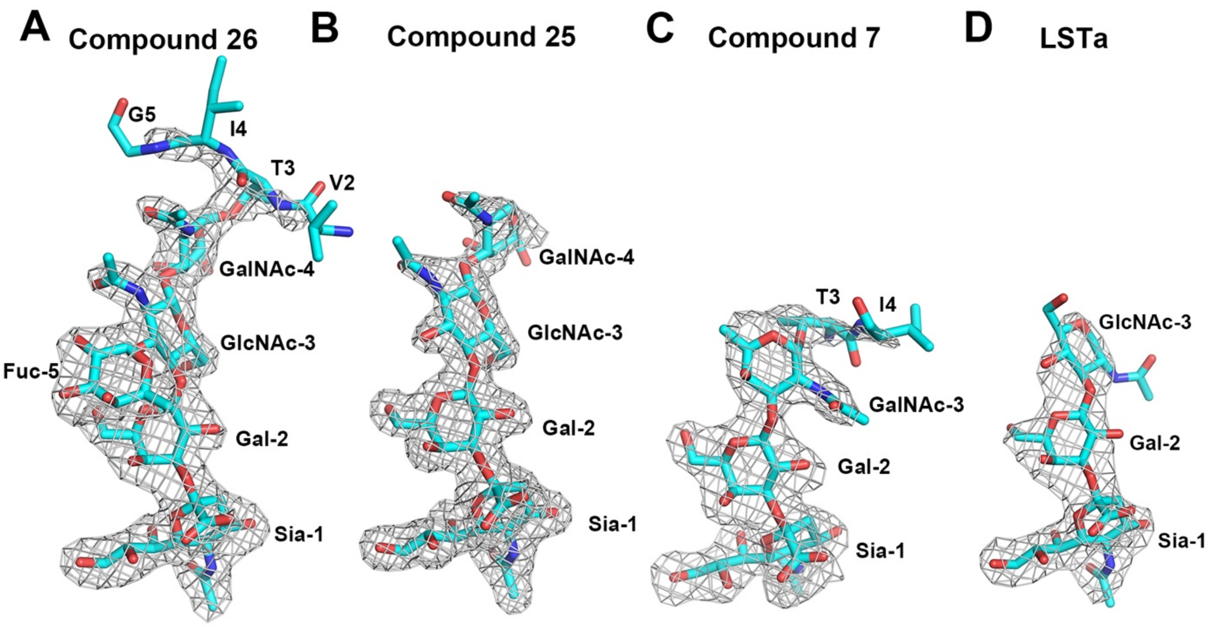

**Figure S7. Simulated annealing omit Fo-Fc electron density maps of glycan compounds bound in H5FR H5 HA RBSs.** (A) Compound **26** (2.50 Å resolution). (B) Compound **25** (1.98 Å resolution). (C) Compound **7** (2.40 Å resolution). (D) Glycan LSTa (2.90 Å resolution). Some residues from the peptide linker moeity (FVTIG) of the glycan compounds (Figure 1) could also be modeled, albeit into weak electron density. The compounds are colored with cyan carbon atoms. Simulated annealing omit *Fo-Fc* maps are represented in grey mesh and contoured at 2.5 σ.

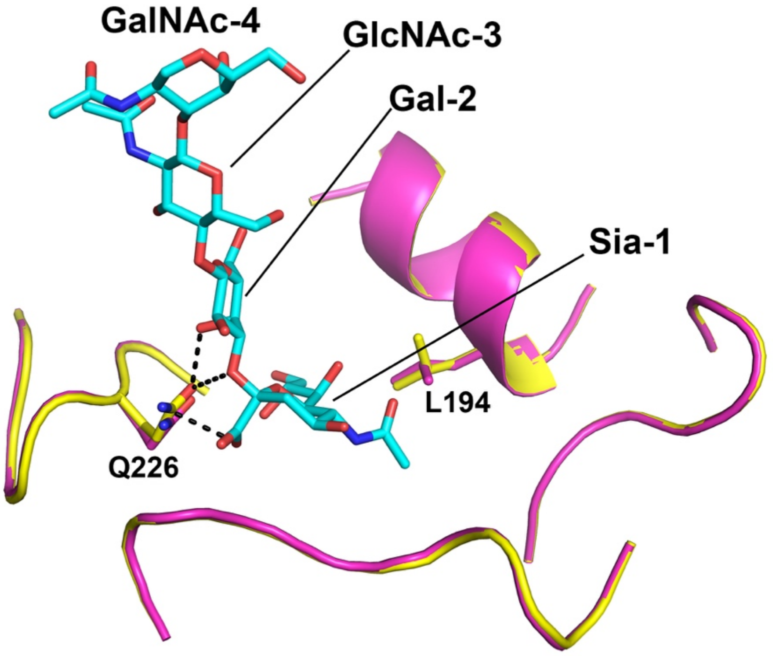

**Figure S8. Structural comparison of the apo-form H5FR H5 HA and its complex with glycan compound 25.** The apo HA is in pink carbon atoms, the complexed HA in yellow carbon atoms, and the ligand in cyan carbon atoms. The side chains of Q226 and L194 with very minor conformational changes are shown for comparison to the unaltered backbones.

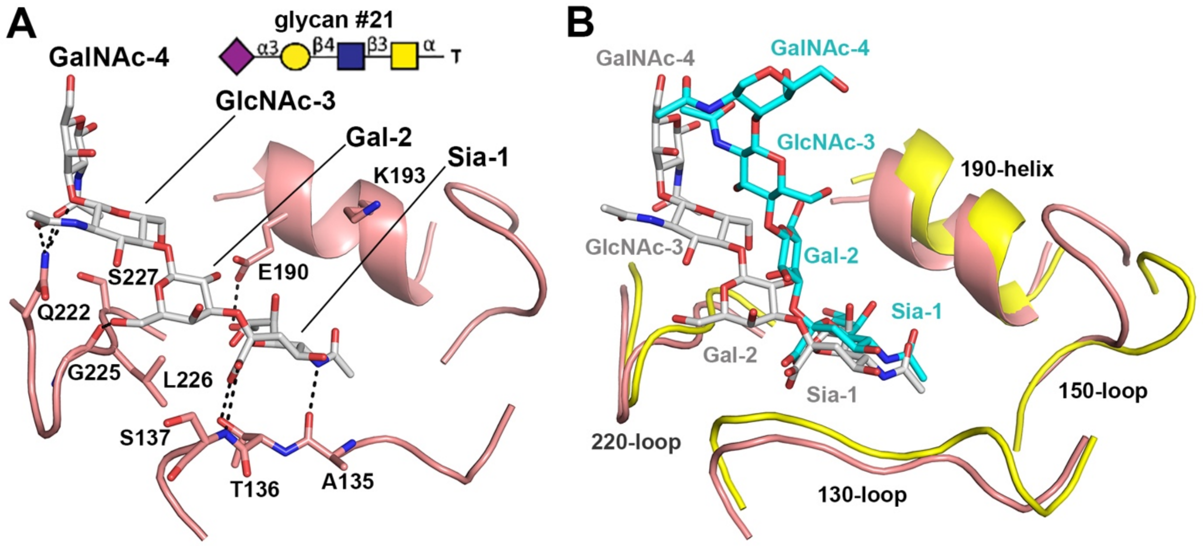

**Figure S9. Structural comparison of human Sh2 H7N9 HA in complex with avian O-linked glycan #21 with H5FR H5 HA in complex with glycan compound 25.** Avian O-linked glycan #21 (NeuAcα2-3Galβ1-4GlcNAcβ1-3GalNAcα-Thr) share the same glycan structure with compound **25,** but with Thr only in the peptide linker. (A) Glycan #21 bound to human H7 HA (PDB ID 4N63). (B) Superposition of human H7 HA in complex #21 with H5FR HA in complex compound **25**. The H7 HA is in pink carbon atoms with ligand #21 in grey carbon atoms, and the H5FR HA in yellow carbon atoms with ligand **25** in cyan carbon atoms.

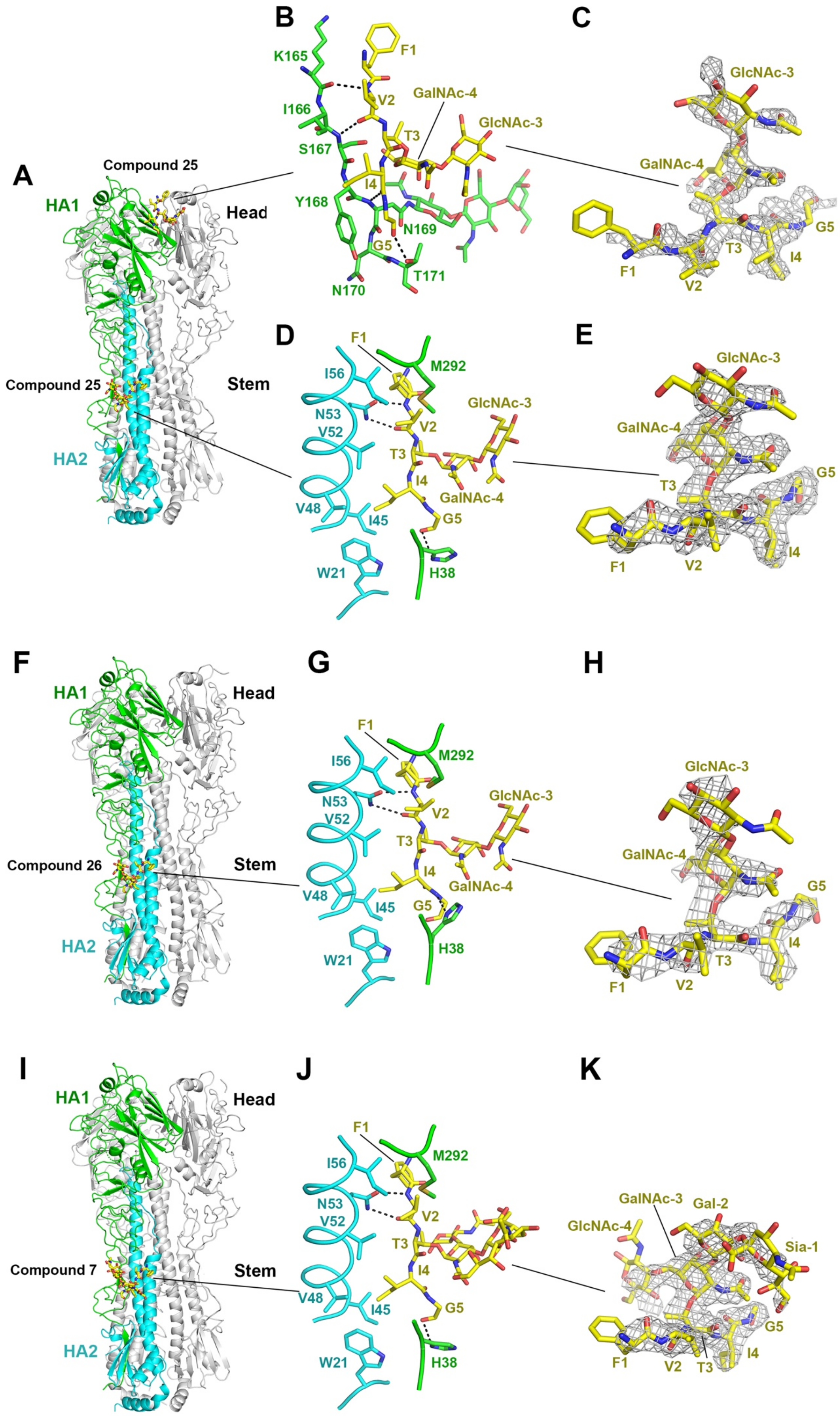
A
B
C
D
E
F
G
H
I
J
K
HA1
HA2
Head
Stem
Compound 25
Compound 26
Compound 7
K165
I166
S167
Y168
N169
N170
T171
F1
V2
T3
I4
G5
GalNAc-4
GlcNAc-3
I56
N53
V52
V48
I45
W21
M292
H38
GalNAc-3
Gal-2
Sia-1
GlcNAc-4

**Figure S10. Crystal structures of H5FR H5 HA binding sites outside of RBS for glycan compounds 25, 26 and 7 through the peptide linker moiety (FVTIG).**

(A) to (E) H5FR H5 HA in complex with compound **25**. In all these figures, compound binding to the RBS has been removed for clarity. One HA protomer is colored with HA1 in green and HA2 in cyan and the other two protomers in grey. Compound **25** is colored in yellow carbon atoms, and the electron density (simulated annealing omit Fo-Fc map) for the glycopeptide is represented in a grey mesh and contoured at 2.5 $\sigma$. The same coloring scheme is used throughout this figure. (A) Overall structure of H5FR HA with **25**. (B) Compound **25** also binds to a non-RBS site in the apex of the HA head domain. (C) Electron density map (as indicated above) for **25** in the HA head. (D) Compound **25** also binds to the HA stem domain. (E) Electron density map for **25** in the HA stem. (F) to (H) H5FR HA with compound **26** bound to the HA stem. (F) Overall structure of H5FR HA with **26**. (G) Compund **26** interaction in the HA stem. (H) Electron density map for **26**. (I) to (K) H5FR HA with compound **7** bound to the HA stem. (I) Overall structure of H5FR with **7**. (G) Compound **7** interacts with the HA stem. (H) Electron density map for **7**.

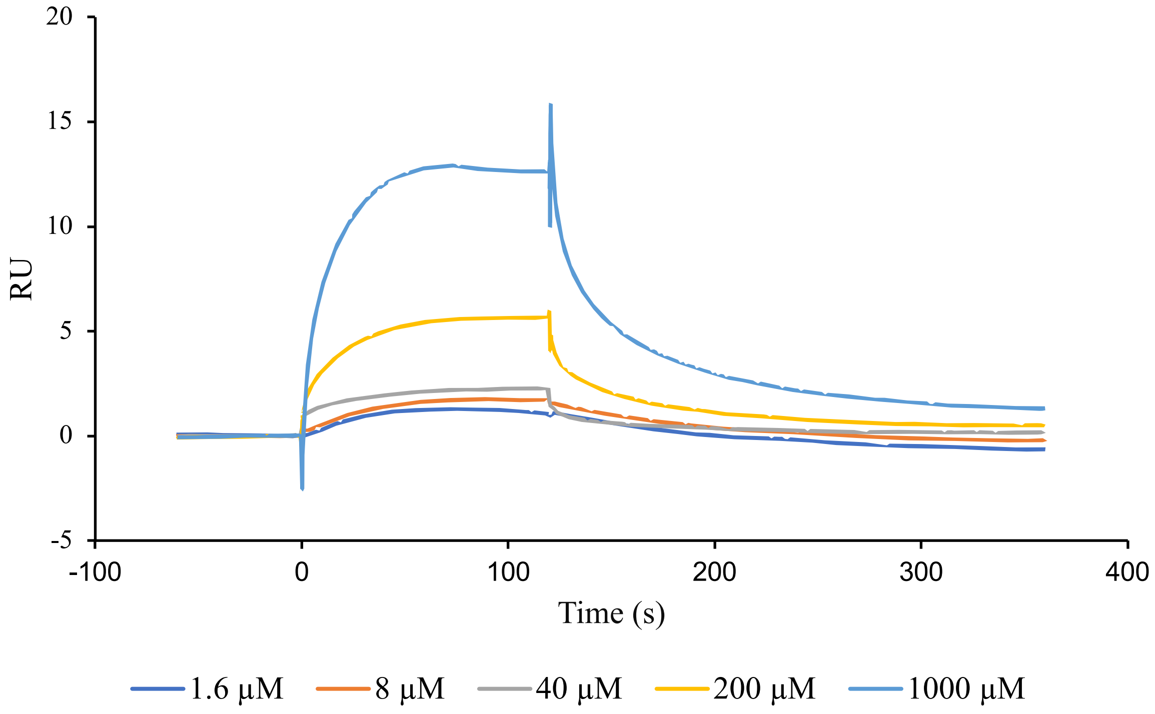

**Figure S11. SPR analysis of glycan compound 2 binding to H5FR H5 HA protein**. Signals (represented as response units, RU) were detected when serial diluted compound **2** solution flow through HA-immobilized sensor chips. Signals were recorded over time. Different colored lines represent different concentrations of **2**, as indicated in the legend. $K_D$ value was estimated from steady-state affinity analysis. Since the upper end of the fitting curve did not saturate, we can only get an estimate of the value of $K_D$>650 µM.

**General remarks**

Thin layer chromatography (TLC) was performed over silica gel 60 F 254 (Merck). The chromatograms were visualized either by UV-radiation (254 or 366 nm) or by heat staining with ceric ammonium molybdate in ethanol/sulfuric acid. HPLC analysis of compounds S2, S11, 1, and 23 were performed on a Shimadzu VP HPLC UV-System using a C18 column (ReproSIL-Pur 120 C18-AQ, 5µm particle size, 250x4.6mm). All other HPLC analysis was done on a Nexera X2® UHPLC system (Shimadzu®) comprised of LC-30AD pumps, SIL-30AC autosampler, CTO-20AC column oven and DGU-20A5/3 degasser module. Detection was done using an SPD-M20A photo diode array and an LCMS-2020 mass spectrometer (ESI/APCI). All separations on the UHPLC system were done using a Waters® XSelect® CSH™ C18 2,5 µm (3.0 x 50 mm) column XP with water/acetonitrile + 0.1% formic acid gradient elution. Preparative column chromatography was performed using a Büchi Sepacore Flash System X10 /X50. Preparative HPLC separation was carried out on a Buchi Pure Chromatography System using a Kinetex® 5 µm C18 100 Å, AXIA LC column (100 x 30.0 mm, Phenomenex) or on a Shimadzu VP Prep HPLC System using a ReproSIL-Pur 120 10µm C18-AQ LC column (250x20mm). Proton nuclear magnetic resonance ($^1$H - NMR) spectra were recorded on a Bruker Ascend 600 MHz spectrometer (Bruker, Germany). Data were recorded using TOPSPIN and evaluated using Mestrenova. Chemical shifts are reported in ppm (δ) relative to tetramethylsilane and calibrated using solvent residual peaks. Multiplicities are abbreviated as s (singlet), br s (broad singlet), d (doublet), dd (doublet of doublets), td (triplet of doublets), t (triplet), q (quartet) or m (multiplet). Spectra were assigned using gCOSY and multiplicity-edited gHSQC experiments. HRMS analysis was carried out using water solutions (concentration: 10 ppm) on an Agilent 6545 LC Q-TOF mass spectrometer equipped with an Agilent Dual AJS ESI-Source. The mass spectrometer was connected to a liquid chromatography system which contained an Agilent G7167B multi sampler, an Agilent G7120A binary pump with degasser, and an Agilent G7116B oven from Agilent Technologies, Palo Alto, CA, USA.

#### Chemical Synthesis

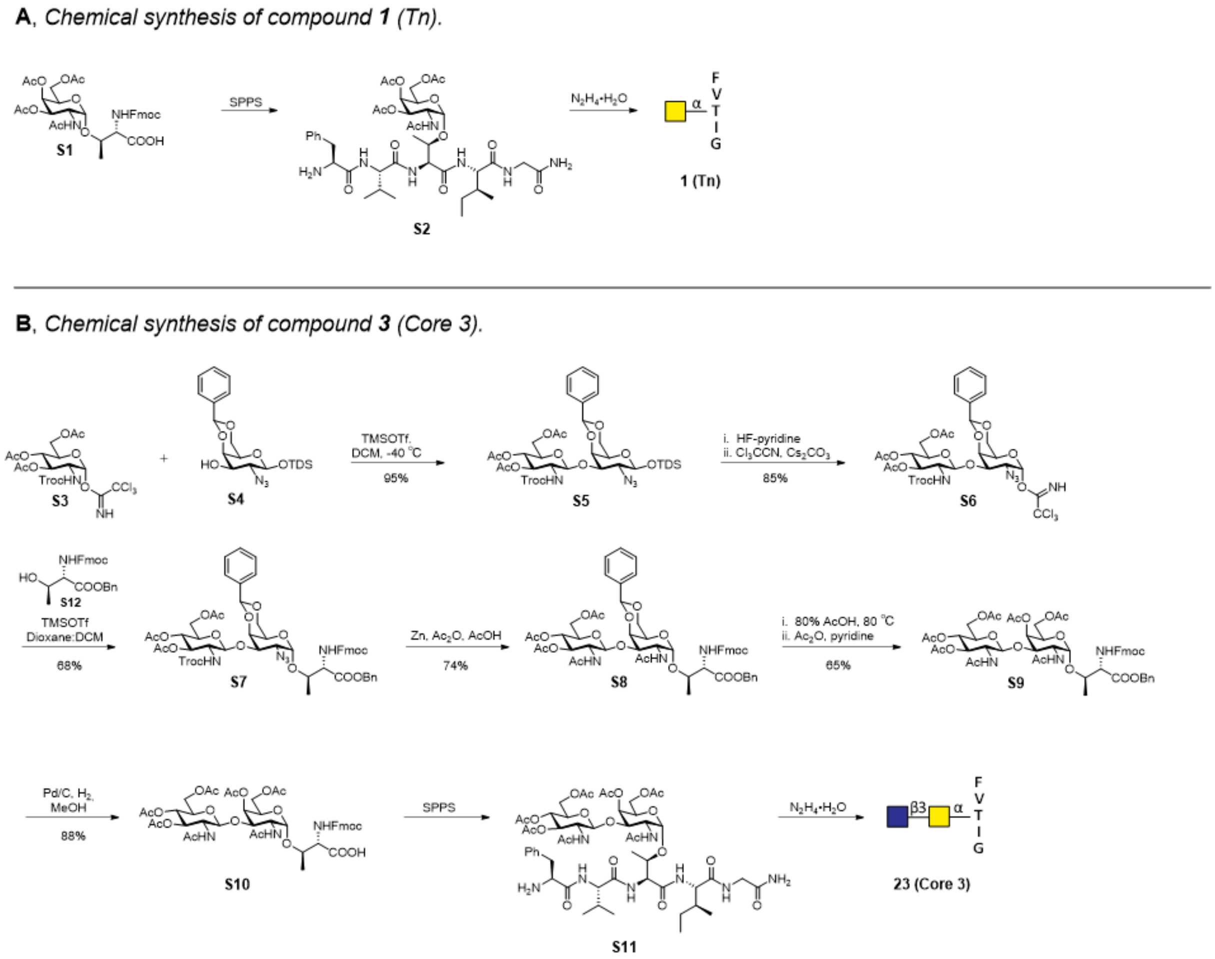

**Scheme S1** Overview of the chemical synthesis towards compounds **1** and **23.**

**N-(Fluoren-9-ylmethoxycarbonyl)-(2-acetamido-2-deoxy-3,4,6-tri-O-acetyl-α,D-glucopyranosyl)-L-threonine (S1)**

T* antigen **S1** was synthesized according to published procedures[6,7].

**2,2,2-Trichloroacetimidate 3,4,6-tri-*O*-acetyl-2-deoxy-2-[[(2,2,2-trichloroethoxy)carbonyl] amino]-α-D-glucopyranoside (S3)**

A solution of D-glucosamine HCl (20 g, 92.7 mmol, 1 eq.) and $NaHCO_3$ (23.4 g, 278 mmol, 3 eq.) in water (180 mL) was stirred vigorously at rt for 5 min. 2,2,2-trichloroethoxycarbonyl chloride (18.5 mL, 134 mmol, 1.4 eq.) was added dropwise and the reaction mixture was stirred at rt. After 4 h the suspension was filtered, washed with water, and dried under vacuum. The resulting white solid (46.5 g, 131 mmol, 1 eq.) was dissolved in 175 mL pyridine and cooled to 0 °C. Acetic anhydride (74 mL, 786 mmol, 6 eq.) was slowly added, and the mixture was stirred for 16h at rt. The mixture was concentrated and coevaporated with toluene. The residue was dissolved in EtOAc (100 mL) and washed with 1M HCl, $H_2O$ and brine. The organic phase was dried over $Na_2SO_4$ and concentrated. The resulting compound (24.4 g, 61%) was dissolved in THF (235 mL) together with DMAPA (30 mL, 233.4 mmol, 5 eq.). After stirring for 2h, the reaction mixture was diluted with DCM, washed with 1M HCl and brine. The organic phase was dried over $Na_2SO_4$, filtered and concentrated. The resulting white solid was dissolved in DCM (250 mL) and trichloroacetonitrile (46.8 mL, 83.2 mmol, 10 eq.) together with DBU (0.67 mL, 0.832 mmol, 0.1 eq) were added at 0 °C. The reaction was slowly warmed to rt and stirred for 2h. The reaction mixture was concentrated and purified by silica gel column chromatography using hexanes-EtOAc (2:1) to afford **S3** (24.0 g, 83%). Analytical data matched those reported in the literature[8].

**Dimethylthexylsilyl 2-azido-4,6-O-benzylidene-2-deoxy-β-D-galactopyranoside (S4)**

Compound **S4** was synthesized according to published procedures[8-10].

**Dimethylthexylsilyl 3,4,6-tri-O-acetyl-2-[(2,2,2-trichloroethoxy)carbonylamino]-β-D-glucopyranosyl-(1→3)-2-azido-2-deoxy-4,6-O-benzylidene-β-D-galactopyranoside (S5)**

A mixture of donor **S3** (3.5 g, 5.6 mmol, 1.5 eq.), acceptor **S4** (1.6 g, 3.7 mmol, 1 eq.) and 4 Å molecular sieves (6 g) was stirred in DCM (63 mL) for 30 min. The reaction mixture was cooled to -40°C and TMSOTf (136 µL, 0.7 mmol, 0.2 eq.) was added. In 1 h the reaction mixture was slowly warmed to -20°C and then quenched with $Et_3N$. The reaction was filtered and concentrated in vacuo. The obtained residue was purified by silica column chromatography using hexanes: EtOAc (5:1 to 3:2) as the eluent to obtain **S5** (3.2 g, 95%). $^1H$ (600MHz, $CDCl_3$): δ = 7.56-7.47 (m, 2H, $CH_{Ar}$-Phe), 7.42-7.31 (m, 3H, $CH_{Ar}$-Phe), 5.53 (s, 1H, CH-$C_6H_5$), 5.44 (t, J = 9.7 Hz, 1H, H3-GlcNAc), 5.25 (d, J = 7.5 Hz, 1H, NH), 5.06 (t, J = 9.7 Hz, 1H, H4-GlcNAc), 5.03 (d, J = 8.0 Hz, 1H, H1-GlcNAc), 4.72 (d, J = 12.1 Hz, 1H, $CH_2$-Troc), 4.68 (d, J = 12.1 Hz, 1H, $CH_2$-Troc), 4.53 (d. J = 7.7 Hz, 1H, H1-GalNAc), 4.29-4.15 (m, 4H, H4, $H6_a$-GalNAc, $H6_{a+b}$-GlcNAc), 4.04 (dd, J = 12.0, 1.8, Hz, 1H, $H6_b$-GalNAc), 3.72 (dd, J = 10.5, 7.8 Hz, 1H, H2-GalNAc), 3.71-3.67 (m, 1H, H5-GlcNAc), 3.60-3.51 (m, 1H, H2-GlcNAc), 3.45 (dd, J = 10.8, 3.7 Hz, 1H, H3-GalNAc), 3.35-3.32 (m, 1H, H5-GalNAc), 2.06 (s, 3H, OAc), 2.03 (s, 3H, OAc), 2.01 (s, 3H, OAc), 1.68 (sept, J = 6.9 Hz, 1H, CH-TDS), 0.92-0.88 (m, 12H, 4x$CH_3$-TDS), 0.21 (s, 3H, Si-$CH_3$), 0.20 (s, 3H, Si-$CH_3$). $^{13}C$ (150MHz, $CDCl_3$): δ = 170.7 (s, 1C, OAc), 170.5 (s, 1C, OAc), 169.6 (s, 1C, OAc), 154.0 (s, 1C, Troc), 137.9 (s, 1C, Phe), 129.2 (d, 1C, $CH_{Ar}$-Phe), 128.4 (d, 2C, $CH_{Ar}$-Phe), 126.5 (d, 2C, $CH_{Ar}$-Phe), 101.4 (d, 1C, C1-GlcNAc), 101.1 (d, 1C, CH-$C_6H_5$), 97.4 (d, 1C, C1-GalNAc), 95.5 (s, 1C, $CCl_3$) 79.0 (d, 1C, C3-GalNAc), 75.2 (d, 1C, C4-GalNAc), 74.6 (t, 1C, $CH_2$-Troc ),72.0 (d, 1C, C5-GlcNAc), 71.5 (d, 1C, C3-GlcNAc),69.2 (t, 1C, C6-GalNAc), 68.8 (d, 1C, C4-GlcNAc), 66.6 (d, 1C, C5-GalNAc), 64.6 (d, 1C, C2-GalNAc), 62.0 (t, 1C, C6-GlcNAc), 56.6 (d, 1C, C2-GlcNAc), 34.1 (d, 1C, CH-TDS), 25.0 (s, 1C, C-Si), 21.0 (q, 1C, OAc), 20.8 (q, 1C, OAc), 20.7 (q, 1C, OAc), 20.2 (q, 1C, $CH_3$-TDS), 20.1 (q, 1C, $CH_3$-TDS), 18.7 (q, 1C, $CH_3$-TDS), 18.6 (q, 1C, $CH_3$-TDS), -2.9 (q, 1C, Si-$CH_3$), -1.7 (q, 1C, Si-$CH_3$). ESI-MS calcd for $C_{36}H_{51}Cl_3N_4NaO_{14}Si^+$ $[M+Na]^+$ 919.2134, found 919.2150

**2,2,2-Trichloroacetimidate 3,4,6-tri-O-acetyl-2-[[(2,2,2-trichloroethoxy)carbonyl]amino]-β-D-glucopyranosyl-(1→3)- 2-azido-2-deoxy-4,6-O-benzylidene-α-D-galactopyranoside (S6)**

To a solution of disaccharide **S5** (3g, 3.3 mmol, 1eq.) in pyridine (30 mL), HF·pyridine (70% HF, 3 mL) was added. The reaction mixture was stirred in a plastic round bottom flask at rt for 16h. Afterwards it was diluted with DCM and quenched with saturated aqueous $NaHCO_3$ solution. The organic phase was washed with sat. aq. $NaHCO_3$ (2x), dried over $Na_2SO_4$, filtered and concentrated in vacuo. Quick silica column purification using hexanes:EtOAc (5:1 to 1:2) as eluent afforded the intermediate in 99% yield. The obtained intermediate (2.5 g, 3.3 mmol, 1eq.) was dissolved in DCM (45 mL) and $Cs_2CO_3$ (1.1 g, 3.3 mmol, 1eq.), followed by $Cl_3CCN$ (3.3 mL, 33 mmol, 10 eq.) was added at 0°C. The reaction mixture was warmed to rt, stirred for 2h and concentrated in vacuo. The residue was purified by silica column chromatography using hexanes: EtOAc (5:1 to 3:2) to afford the title compound **S6** (2.3 g, 85%). $^1$H (600MHz, $CDCl_3$): δ = 8.75 (s, 1H, C=NH), 7.55-7.49 (m, 2H, $CH_{Ar}$-Phe), 7.42-7.33 (m, 3H, $CH_{Ar}$-Phe), 6.55 (d, J = 3.3 Hz, 1H, H1-GalNAc), 5.57 (s, 1H, CH-$C_6H_5$), 5.47 (t, J = 9.9 Hz, 1H, H3-GlcNAc), 5.37 (d, J = 8.4 Hz, 1H, NH), 5.11-5.06 (m, 2H, H4-GlcNAc, H1-GlcNAc), 4.74 (d, J = 12.1 Hz, 1H, $CH_2$-Troc), 4.59 (d, J = 12.1 Hz, 1H, $CH_2$-Troc), 4.53 (d, J= 2.2 Hz, 1H, H4-GalNAc), 4.36 (dd, J= 12.3, 2.5, 1H, $H6_a$-GlcNAc), 4.29 (dd, J = 12.7, 1.6 Hz, 1H, $H6_a$-GalNAc), 4.21 (dd, J = 10.6, 3.4 Hz, 1H, H2-GalNAc), 4.17-4.10 (m, 2H, $H6_b$-GlcNAc, H3-GalNAc), 4.04 (dd, J = 12.7, 1.8, Hz, 1H, $H6_b$-GalNAc), 3.89-3.85 (m, 1H, H5-GalNAc), 3.81-3.75 (m, 1H, H5-GlcNAc), 3.68-3.58 (m, 1H, H2-GlcNAc), 2.04 (s, 3H, OAc), 2.02 (s, 6H, 2xOAc). $^{13}$C (150MHz, $CDCl_3$): δ = 170.6 (s, 1C, OAc), 170.5 (s, 1C, OAc), 169.7 (s, 1C, OAc), 160.6 (s, 1C, C=NH), 154.0 (s, 1C, Troc), 137.6 (s, 1C, Phe), 129.2 (d, 1C, $CH_{Ar}$-Phe), 128.4 (d, 2C, $CH_{Ar}$-Phe), 126.2 (d, 2C, $CH_{Ar}$-Phe), 101.6 (d, 1C, C1-GlcNAc), 100.9 (d, 1C, CH-$C_6H_5$), 95.7 (d, 1C, C1-GalNAc), 95.4 (s, 1C, $CCl_3$-Troc), 91.1 (s, 1C, $CCl_3$-Im), 76.8 (d, 1C, C3-GalNAc), 75.2 (d, 1C, C4-GalNAc), 74.6 (t, 1C, $CH_2$-Troc ), 72.1 (d, 1C, C5-GlcNAc), 71.5 (d, 1C, C3-GlcNAc),69.6 (t, 1C, C6-GalNAc), 68.7 (d, 1C, C4-GlcNAc), 65.5 (d, 1C, C5-GalNAc), 62.1 (t, 1C, C6-GlcNAc), 58.5 (d, 1C, C2-GalNAc), 56.6 (d, 1C, C2-GlcNAc), 21.0 (q, 1C, OAc), 20.8 (q, 1C, OAc), 20.7 (q, 1C, OAc). ESI-MS calcd for $C_{30}H_{33}Cl_6N_5NaO_{14}^+$ $[M+Na]^+$ 920.0053, found 920.0096

##### ***N*-(9*H*-Fluorene-9-yl)-methoxycarbonyl-*O*-(2-azido-2-deoxy-4,6-*O*-benzylidene-3-*O*-[ 2-deoxy-2-[[(2,2,2-trichloro-ethoxy)carbonyl]amino]-3,4,6-tri-O-acetyl-β-D-glucopyranosyl]-α-D-galactopyranosyl)-L-threonine benzyl ester (S7)**

A mixture of disaccharide donor **S6** (1 g, 1.1 mmol, 1.3 eq), acceptor **S12** (368 mg, 0.85 mmol, 1 eq.) and 4 Å molecular sieves (500 mg) was stirred in DCM: dioxane (1:1, 10 mL). After 30 min TMSOTf (14 µL, 70 µmol, 0.09 eq.) was added and the reaction was continued to be stirred for 1.5h. The reaction mixture was quenched with DIPEA (13 µL), filtered and concentrated in vacuo. The residue was purified by silica column chromatography using hexanes:EtOAc (5:1 to 3:2) as the eluent to obtain **S7** (672 mg, 68%). $^1$H (600MHz, $CDCl_3$): δ = 7.82-7.71 (m, 2H, CH-Fmoc), 7.68-7.58 (m, 2H,), 7.53-7.47 (m, 2H, CH-Fmoc), 7.45-7.27 (m, 12H, CH-Fmoc, $CH_{Ar}$-Phe, $CH_{Ar}$-Bn), 5.76 (d, J = 9.4 Hz, 1H, NH-Thr), 5.53 (s, 1H, CH-$C_6H_5$), 5.45 (t, J = 9.9 Hz, 1H, H3-GlcNAc), 5.27 (d, J = 8.4 Hz, 1H, NHTroc), 5.22 (s, 2H, $CH_2$-Bn), 5.08 (t, J = 9.6 Hz, 1H, H4-GlcNAc), 5.01 (d, J = 8.2 Hz, 1H, H1-GlcNAc), 4.92 (d, J = 3.7 Hz, 1H, H1-GalNAc), 4.69 (d, J = 12.0 Hz, 1H, $CH_2$-Troc), 4.60 (d, J = 12.0 Hz, 1H, $CH_2$-Troc), 4.52 (dd, J = 10.6, 7.0 Hz, 1H, $CH_2$-Fmoc), 4.48- 4.43 (m, 2H, $CH_{\alpha+\beta}$-Thr), 4.41 (d, J= 3.3 Hz, 1H, H4-GalNAc), 4.39-4.32 (m, 2H, $H6_a$-GlcNAc, $CH_2$-Fmoc), 4.27 (t, J = 7.2 Hz, 1H, CH-

Fmoc), 4.21 (dd, J = 12.6, 1.6 Hz, 1H, $H6_a$-GalNAc), 4.13 (dd, J = 4.4, 12.4 Hz, 1H, $H6_b$-GlcNAc), 4.02 (dd, J = 12.6, 1.6, Hz, 1H, $H6_b$-GalNAc), 4.02-3.97 (m, 1H, H3-GalNAc), 3.80 (dd, J = 10.8, 3.6 Hz, 1H, H2-GalNAc), 3.76-3.69 (m, 1H, H5-GlcNAc), 3.67-3.57 (m, 2H, H5-GalNAc, H2-GlcNAc), 2.04 (s, 3H, OAc), 2.03-193 (m, 6H, 2xOAc), 1.30 (d, J = 6.3 Hz, 3H, $CH_3$-Thr). $^{13}C$ (150MHz, $CDCl_3$): δ = 170.6 (s, 1C, OAc), 170.5 (s, 1C, OAc), 170.2 (s, 1C, OAc), 169.7 (s, 1C, COO-Thr), 156.9 (s, 1C, COO-Fmoc), 154.1 (s, 1C, Troc), 144.0 (s, 1C, Fmoc), 143.8 (s, 1C, Fmoc), 141.4 (s, 2C, Fmoc), 137.7 (s, 1C, Phe), 135.1 (s, 1C, Bn), 129.1 (d, 1C, $CH_{Ar}$), 128.8 (d, 4C, $CH_{Ar}$), 128.6 (d, 1C, $CH_{Ar}$), 128.3 (d, 2C, $CH_{Ar}$), 127.9 (d, 2C, $CH_{Ar}$), 127.3 (d, 2C, $CH_{Ar}$), 126.3 (d, 2C, $CH_{Ar}$), 125.3 (d, 2C, $CH_{Ar}$), 120.2 (d, 2C, $CH_{Ar}$), 101.5 (d, 1C, C1-GlcNAc), 100.9 (d, 1C, CH-$C_6H_5$), 99.2 (d, 1C, C1-GalNAc), 95.4 (s, 1C, $CCl_3$-Troc), 76.5 (d, 1C, C3-GalNAc), 76.2 (d, 1C, Thr), 75.5 (d, 1C, C4-GalNAc), 74.6 (t, 1C, $CH_2$-Troc ), 72.0 (d, 1C, C5-GlcNAc), 71.5 (d, 1C, C3-GlcNAc),69.2 (t, 1C, C6-GalNAc), 68.7 (d, 1C, C4-GlcNAc), 67.9 (t, 1C, Bn), 67.5 (t, 1C, Fmoc), 63.6 (d, 1C, C5-GalNAc), 61.7 (t, 1C, C6-GlcNAc), 59.2 (d, 1C, C2-GalNAc), 58.9 (d, 1C, Thr), 56.6 (d, 1C, C2-GlcNAc), 47.3 (d, 1C, Fmoc), 21.0 (q, 1C, OAc), 20.8 (q, 1C, OAc), 20.7 (q, 1C, OAc), 18.2 (q, 1C, Thr). ESI-MS calcd for $C_{54}H_{56}Cl_3N_5NaO_{18}^+$ $[M+Na]^+$ 1190.2584, found 1190.2599

**_N_-(9_H_-Fluorene-9-yl)-methoxycarbonyl-_O_-(2-acetamido-2-deoxy-4,6-_O_-benzylidene-3-_O_-[ 2-acetamido-2-deoxy-3,4,6-tri-O-acetyl-β-D-glucopyranosyl]-α-D-galactopyranosyl)-L-threonine benzyl ester (S8)**

To a solution of glycoside **S7** (810 mg, 0.69 mmol, 1 eq.) in THF: $Ac_2O$: AcOH (3:2:1, 15 mL), activated zinc dust (589 mg, 9.0 mmol, 13 eq.) was added. The reaction mixture was stirred for 2 h at rt, filtered over a pad of Celite and concentrated in vacuo. The obtained residue was purified by silica column chromatography using DCM: MeOH (1:0 to 30:1) as eluent to obtain compound **S8** (536 mg, 74%). $^1H$ (600MHz, $CDCl_3$): δ = 7.82-7.71 (m, 2H, CH-Fmoc), 7.68-7.58 (m, 2H, CH-Fmoc), 7.54-7.45 (m, 2H, CH-Fmoc), 7.44-7.38 (m, 2H, CH-Fmoc), 7.38-7.29 (m, 10H, $CH_{Ar}$-Phe, $CH_{Ar}$-Bn), 6.12 (d, J = 9.6 Hz, 1H, NH-Thr), 6.09 (d, J = 9.0 Hz, 1H, NH-GalNAc), 5.85 (d, J = 8.2 Hz, 1H, NH-GlcNAc), 5.50 (s, 1H, CH-$C_6H_5$), 5.31 (t, J = 9.8 Hz, 1H, H3-GlcNAc), 5.21 (d, J = 12.0 Hz, 1H, $CH_2$-Bn), 5.11-5.05 (m, 2H, $CH_2$-Bn, H4-GlcNAc), 5.01 (d, J = 8.4 Hz, 1H, H1-GlcNAc), 4.91 (d, J = 3.9 Hz, 1H, H1-GalNAc), 4.58-4.38 (m, 5H, $CH_2$-Fmoc, H2-GalNAc, $CH_2$-GlcNAc, $CH_α$-Thr), 4.38- 4.29 (m, 2H, $CH_β$-Thr, H4-GalNAc), 4.25 (t, J = 6.6 Hz, 1H, CH-Fmoc), 4.18 (d, J = 12.2 Hz, 1H, $H6_a$-GalNAc), 4.08 (dd, J = 4.0, 12.3 Hz, 1H, $H6_b$-GlcNAc), 4.04 (d, J = 12.3 Hz, 1H, $H6_b$-GalNAc), 3.92 (dd, J = 3.5, 11.2 Hz, 1H, H3-GalNAc), 3.77-3.60 (m, 3H, H2, H5-GlcNAc, H5-GalNAc), 2.04 (s, 3H, OAc), 2.03 (s, 6H, OAc, NHAc), 2.02 (s, 3H, OAc), 1.88 (s, 3H, NHAc), 1.25 (d, J = 6.3 Hz, 3H, $CH_3$-Thr). $^{13}C$ (150MHz, $CDCl_3$): δ = 171.3 (s, 1C, NHAc), 171.1 (s, 1C, NHAc), 170.8 (s, 1C, OAc), 170.7 (s, 1C, OAc), 170.6 (s, 1C, OAc), 169.5 (s, 1C, COO-Thr), 157.0 (s, 1C, COO-Fmoc), 143.9 (s, 1C, Fmoc), 143.8 (s, 1C, Fmoc), 141.5 (s, 2C, Fmoc), 137.9 (s, 1C, Phe), 134.8 (s, 1C, Bn), 128.9 (d, 3C, $CH_{Ar}$), 128.7 (d, 2C, $CH_{Ar}$), 128.3 (d, 3C, $CH_{Ar}$), 127.9 (d, 2C, $CH_{Ar}$), 127.2 (d, 2C, $CH_{Ar}$), 126.4 (d, 2C, $CH_{Ar}$), 125.1 (d, 2C, $CH_{Ar}$), 120.1 (d, 2C, $CH_{Ar}$), 101.0 (d, 1C, CH-$C_6H_5$), 100.1 (d, 1C, C1-GalNAc), 99.4 (d, 1C, C1-GlcNAc), 76.5 (d, 1C, C3-GalNAc), 75.8 (d, 1C, Thr), 75.2 (d, 1C, C4-GalNAc), 72.5 (d, 1C, C5-GlcNAc), 72.3 (d, 1C, C3-GlcNAc), 69.3 (t, 1C, C6-GalNAc), 68.8 (d, 1C, C4-GlcNAc), 67.8 (t, 1C, Bn), 67.1 (t, 1C, Fmoc), 63.5 (d, 1C, C5-GalNAc),

61.8 (t, 1C, C6-GlcNAc), 58.9 (d, 1C, Thr), 55.2 (d, 1C, C2-GlcNAc), 48.3 (d, 1C, C2-GalNAc), 47.3 (d, 1C, Fmoc), 23.5 (q, 2C), 20.9 (q, 1C, OAc), 20.8 (q, 2C, OAc), 19.1 (q, 1C, Thr). ESI-MS calcd for $C_{55}H_{61}N_3NaO_{18}^+$ $[M+Na]^+$ 1074.3848, found 1074.3841.

**_N_-(9_H_-Fluorene-9-yl)-methoxycarbonyl-_O_-(2-acetamido-2-deoxy-4,6-di-_O_-acetyl-3-_O_-[2-acetamido-2-deoxy-3,4,6-tri-O-acetyl-β-D-glucopyranosyl]-α-D-galactopyranosyl)-L-threonine benzyl ester (S9)**

A solution of compound **S8** (478 mg, 0.45 mmol, 1 eq.) in 80% aqu. acetic acid (4.6 mL) was heated to 80°C for 3h. The reaction mixture was allowed to cool to rt and concentrated in vacuo. After coevaporation with toluene, the residue was purified by a quick column chromatography using DCM: MeOH (1:0 to 20:1) to afford the intermediate (328 mg, 75%). The obtained intermediate (328 mg, 0.34 mmol, 1 eq.) was dissolved in pyridine (3.3 mL) and acetic anhydride (1.7 mL) was added. After stirring the reaction mixture for 16h at rt, it was concentrated in vacuo. The crude product was purified by column chromatography using DCM:MeOH (1:0 to 11:1) to yield the title compound **S9** (306 mg, 86%). $^1H$ (600MHz, $CDCl_3$): δ = 7.82-7.71 (m, 2H, CH-Fmoc), 7.67-7.58 (m, 2H, CH-Fmoc), 7.45-7.40 (m, 2H, CH-Fmoc), 7.39-7.29 (m, 7H, CH-Fmoc, $CH_{Ar}$-Bn), 6.28 (d, J = 8.7 Hz, 1H, NH-GalNAc), 5.97 (d, J = 9.6 Hz, 1H, NH-Thr), 5.91 (d, J = 7.9 Hz, 1H, NH-GlcNAc), 5.39 (t, J = 9.9 Hz, 1H, H3-GlcNAc), 5.34(d, J = 3.2 Hz, 1H, H4-GalNAc), 5.19 (d, J = 12.0 Hz, 1H, $CH_2$-Bn), 5.09 (d, J = 12.0 Hz, 1H, $CH_2$-Bn), 5.06 (t, J = 9.8 Hz, 1H, H4-GlcNAc), 4.98 (d, J = 8.2 Hz, 1H, H1-GlcNAc), 4.81 (d, J = 3.8 Hz, 1H, H1-GalNAc), 4.56 (dd, J = 6.6, 10.8 Hz, 1H, $CH_2$-Fmoc), 4.48 (dd, J = 6.6, 10.8 Hz, 1H, $CH_2$-Fmoc), 4.43-4.23 (m, 5H, $H6_a$-GlcNAc, CH-Fmoc, H2-GalNAc, $CH_{\alpha+\beta}$-Thr), 4.16- 4.03 (m, 3H, H5-$H6_a$-GalNAc, $H6_b$-GlcNAc), 4.01-3.93 (m, 1H, $H6_b$-GalNAc), 3.88 (dd, J = 3.3, 11.0 Hz, 1H, H3-GalNAc), 3.77-3.70 (m, 1H, H5-GlcNAc), 3.56-3.48 (m, 1H, H2- GlcNAc), 2.11 (s, 3H, OAc), 2.06 (s, 3H, OAc), 2.04 (s, 6H, OAc), 2.02 (s, 3H, NHAc), 2.00 (s, 3H, OAc), 1.95 (s, 3H, NHAc), 1.27 (d, J = 6.3 Hz, 3H, $CH_3$-Thr). $^{13}C$ (150MHz, $CDCl_3$): δ = 171.4 (s, 1C, NHAc), 171.3 (s, 1C, NHAc), 170.9 (s, 1C, OAc), 170.8 (s, 2C, OAc), 170.6 (s, 1C, OAc), 170.2 (s, 1C, OAc), 169.5 (s, 1C, COO-Thr), 157.0 (s, 1C, COO-Fmoc), 143.9 (s, 1C, Fmoc), 143.8 (s, 1C, Fmoc), 141.5 (s, 2C, Fmoc), 134.7 (s, 1C, Bn), 128.9 (d, 3C, $CH_{Ar}$), 128.7 (d, 2C, $CH_{Ar}$), 127.9 (d, 2C, $CH_{Ar}$), 127.3 (d, 2C, $CH_{Ar}$), 125.1 (d, 2C, $CH_{Ar}$), 120.2 (d, 2C, $CH_{Ar}$), 99.7 (d, 1C, C1-GalNAc), 99.2 (d, 1C, C1-GlcNAc), 76.3 (d, 1C, Thr), 75.2 (d, 1C, C4-GalNAc), 72.3 (d, 1C, C5-GlcNAc), 72.2 (d, 1C, C3-GalNAc), 72.1 (d, 1C, C3-GlcNAc), 68.5 (d, 1C, C4-GlcNAc), 67.9 (t, 1C, Bn), 67.8 (d, 1C, H5-GalNAc), 67.1 (t, 1C, Fmoc), 62.8 (t, 1C, C6-GalNAc), 61.7 (t, 1C, C6-GlcNAc), 58.9 (d, 1C, Thr), 55.5 (d, 1C, C2-GlcNAc), 48.9 (d, 1C, C2-GalNAc), 47.3 (d, 1C, Fmoc), 23.5 (q, 2C, NHAc), 20.9 (q, 1C, OAc), 20.8 (q, 4C, OAc), 18.7 (q, 1C, Thr). ESI-MS calcd for $C_{52}H_{61}N_3NaO_{20}^+$ $[M+Na]^+$ 1070.3746, found 1070.3743

**_N_-(9_H_-Fluorene-9-yl)-methoxycarbonyl-_O_-(2-acetamido-2-deoxy-4,6-di-_O_-acetyl-3-_O_-[ 2-acetamido-2-deoxy-3,4,6-tri-O-acetyl-β-D-glucopyranosyl]-α-D-galactopyranosyl)-L-threonine (S10)**

To a solution of compound **S9** (170 mg, 0.16 mmol, 1 eq.) in MeOH, Pd/C (30 mg) was added under argon atmosphere. The argon balloon was changed for a $H_2$-balloon and the reaction mixture was stirred for 1.5h. The reaction mixture was filtered through a syringe PET filter and the filtrate was concentrated. The crude product was purified by column chromatography using DCM+0.1% AcOH: MeOH+0.1% AcOH (1:0 to 11:1) to afford compound **S10** (135 mg, 88%). $^{1}$H (600MHz, MeOD): δ = 7.84-7.79 (m, 2H, Fmoc), 7.73-7.76 (m, 2H, Fmoc), 7.45-7.37 (m, 2H, Fmoc), 7.36-7.31 (m, 2H, Fmoc), 5.44 (dd, J = 9.2, 10.6 Hz, 1H, H3-GlcNAc), 5.38 (d, J = 4.4 Hz, 1H, H4-GalNAc), 4.98 (d, J = 8.2 Hz, 1H, H1-GlcNAc), 4.95 (t, J = 9.7 Hz, 1H, H4-GlcNAc), 4.86 (d, J = 3.9 Hz, 1H, H1-GalNAc)*, 4.58 (dd, J = 6.1, 10.8 Hz, 1H, $CH_2$-Fmoc), 4.54 (dd, J = 6.1, 10.8 Hz, 1H, $CH_2$-Fmoc), 4.37 (qd, J = 2.1, 6.5 Hz, 1H, $CH_β$-Thr), 4.32-4.27 (m, 3H, H2, $H6_a$-GalNAc, CH-Fmoc), 4.25 (d, J = 2.1 Hz, 1H, $CH_α$-Thr), 4.21-4.17 (m, 1H, H5-GalNAc), 4.15-4.09 (m, 2H, $H6_b$-GalNAc, $H6_a$-GlcNAc), 3.99 (dd, J = 7.6, 11.5 Hz, 1H, $H6_b$-GlcNAc), 3.87 (dd, J = 3.4, 11.1 Hz, 1H, H3-GalNAc), 3.71-3.65 (m, 1H, H5-GlcNAc), 3.33 (dd, J = 8.1, 11.4 Hz, 1H, H2-GlcNAc), 2.11 (s, 3H, OAc), 2.07 (s, 3H, OAc), 2.06 (s, 3H, OAc), 2.01 (s, 3H, OAc), 1.98 (s, 3H, OAc), 1.97 (s, 3H, NHAc), 1.87 (s, 3H, NHAc), 1.24 (d, J = 6.4 Hz, 3H, $CH_3$-Thr). $^{13}$C (150MHz, MeOD): δ = 173.5 (q, 1C, COOH), 173.4 (q, 2C, NHAc), 172.5 (q, 1C, OAc), 172.3 (q, 1C, OAc), 171.9 (q, 1C, OAc), 171.8 (q, 1C, OAc), 171.2 (q, 1C, OAc), 159.1 (q, 1C, Fmoc), 145.4 (q, 1C, Fmoc), 145.1 (q, 1C, Fmoc), 142,7 (q, 2C, Fmoc), 128.9 (d, 1C, Fmoc), 128.8 (d, 1C, Fmoc), 128.2 (d, 2C, Fmoc), 126.1 (d, 1C, Fmoc), 126.0 (d, 1C, Fmoc), 121.0 (d, 2C, Fmoc), 101.3 (d, 1C, C1-GlcNAc), 100.9 (d, 1C, C1-GalNAc), 77.3 (d, 1C, Thr), 74.6 (d, 1C, C3-GalNAc), 72.9 (d, 1C, C3-GlcNAc), 72.8 (d, 1C, C5-GlcNAc), 71.4 (d, 1C, C4-GalNAc), 70.3 (d, 1C, C4-GlcNAc), 68.7 (d, 1C, C5-GalNAc), 67.6 (t, 1C, Fmoc), 64.1 (t, 1C, C6-GalNAc), 62.6 (t, 1C, C6-GlcNAc), 59.9 (d, 1C, Thr), 56.9 (d, 1C, C2-GlcNAc), 49.8 (d, 1C, C2-GalNAc), 48.6 (d, 1C, Fmoc), 23.3 (q, 1C, NHAc), 23.0 (q, 1C, NHAc), 20.8 (q, 2C, OAc), 20.7 (q, 1C, OAc), 20.6 (q, 2C, OAc), 19.1 (q, 1C, Thr). ESI-MS calcd for $C_{45}H_{55}N_3NaO_{20}^+$ $[M+Na]^+$ 980.3277, found 980.3273

**Solid Phase Peptide Synthesis (SPPS) of the glycopeptide FVT*IG (*= $Ac_3$GalNAc (S2) or $Ac_3$GlcNAc-β1,3-$Ac_2$GalNAc (S11))**

The peptide FVT*IG was synthesized manually using a peptide synthesis glass vessel. Agitation was done by nitrogen bubbling and vacuum was used to remove the solvents. Rink resin (0.23mmol/g) was used as the solid support. The N-terminus Fmoc group was deprotected using 20% piperidine in DMF. The couplings were performed using HOBt/HBTU as activators. To a solution of the Fmoc-protected amino acid (4 eq.), HBTU (4 eq.) and HOBt (4 eq.) in DMF, DIPEA (8eq.) was added. The solution was quickly mixed and immediately added to the resin. The suspension was agitated for 45 min. A Kaiser test was performed after each coupling to control the coupling conversion. The coupling step with the Fmoc- and acetyl-protected Tn-antigen **S1** or compound **S10** was done using less equivalents of the T* (2eq.), HBTU (2eq.), HOBt (2eq.) and DIPEA (4 eq.). The coupling time was extended to 16h. An acetyl capping step using $Ac_2O$, DIPEA and HOBt in DMF was performed after each coupling. After the last coupling step with Phe, the Fmoc group was deprotected to obtain a free amino group at the end of the peptide. The peptide was cleaved off the resin by stirring the resin in 95% TFA solution in water for 2h. The peptide was precipitated from a cold mixture of ether/hexanes (1:1). The glycopeptide was purified by preparative RP-C18 HPLC (ReproSIL-Pur 120 C18-AQ, 10µm ps, 250x20mm, $H_2O$:ACN with 0.1% TFA 5-50%, 12.5 mL/min, 120 min). **S2**: ESI-MS calcd for $C_{40}H_{62}N_7O_{14}^+$ $[M+H]^+$ 864.4355, found 864.4359. **S11**: ESI-MS calcd for $C_{52}H_{78}N_8NaO_{21}^+$ $[M+Na]^+$ 1173.5179, found 1173.5181. A UV-spectra was measured for characterization at 214 nm using a

ReproSIL-Pur 120 C18-AQ, 5µm ps, 250x4.6mm column ($H_20$:ACN with 0.1% TFA 5-95%, 1 mL/min, 60 min).

##### ***N*-(9*H*-Fluorene-9-yl)-methoxycarbonyl L-threonine benzyl ester (S12)**

The protected threonine **S12** was synthesized as described before [9,11].

**HPLC-UV chromatogram at 214nm of compound S2**

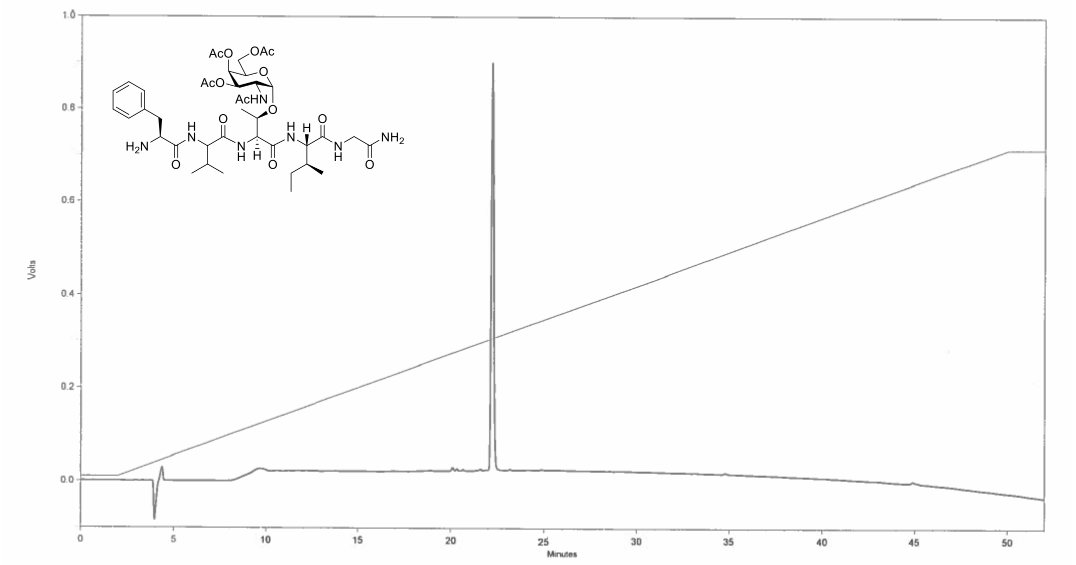

**HPLC-UV chromatogram at 214nm of compound S11**

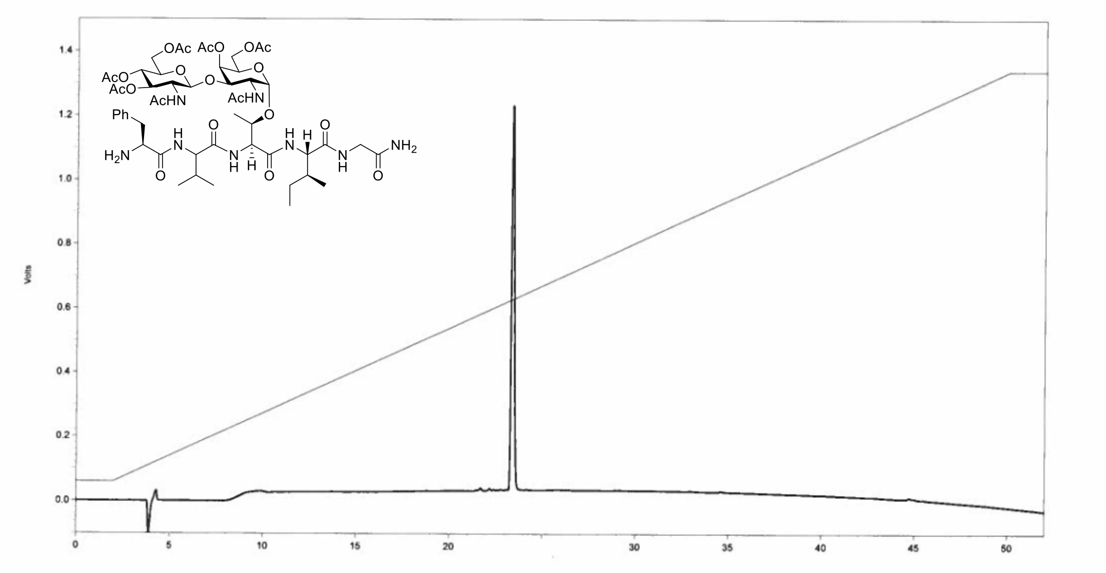

##### **Deacetylation of the glycopeptide FVT*IG (1)**

Glycopeptide **S2** (45 mg, 0.05 mmol, 1eq.) was dissolved in a 5% aqueous hydrazine hydrate solution (4 mL). The reaction mixture was stirred at rt. After 16 h the reaction solution was directly loaded on a preparative RP-C18 HPLC-column (ReproSIL-Pur

120 C18-AQ, 10µm ps, 250x20mm, $H_20$:ACN with 0.1% TFA 5-35%, 12.5 mL/min, 120 min) to obtain the deprotected glycopeptide **1** (29 mg, 75%). ESI-MS calcd for $C_{34}H_{56}N_7O_{11}^+$ [M+H]$^+$ 738.4038, found 738.4045. A UV-spectra was measured for characterization at 214 nm using a ReproSIL-Pur 120 C18-AQ, 5µm ps, 250x4.6mm column ($H_2O$:ACN with 0.1% TFA 5-95%, 1 mL/min, 60 min). Deacetylation of glycopeptide **S11** was performed in a similar way to obtain compound **23**.

**HPLC-UV chromatogram at 214nm of compound 1**

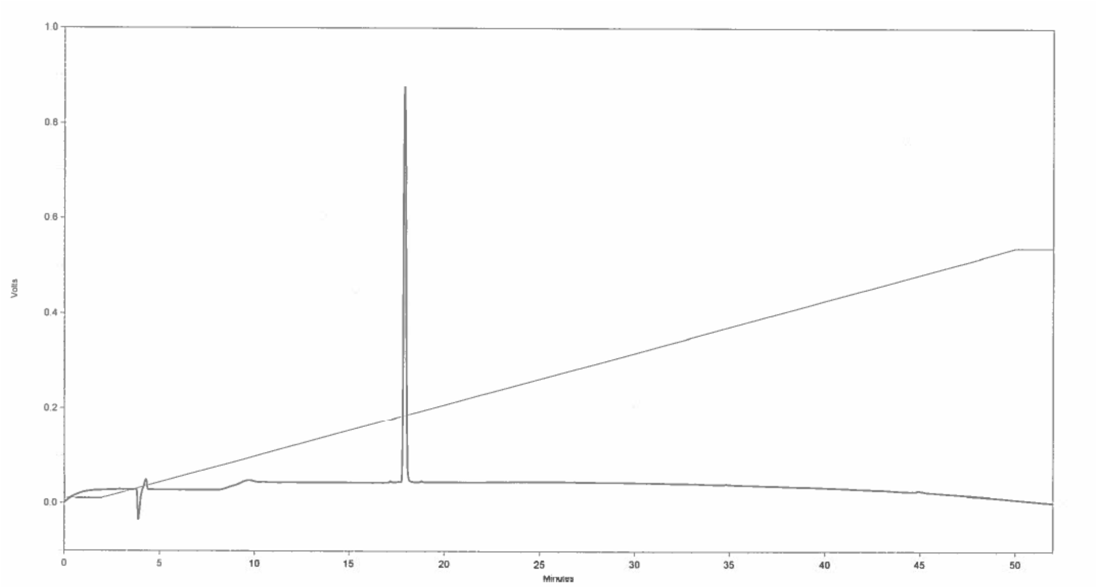

**HPLC-UV chromatogram at 214nm of compound 23**

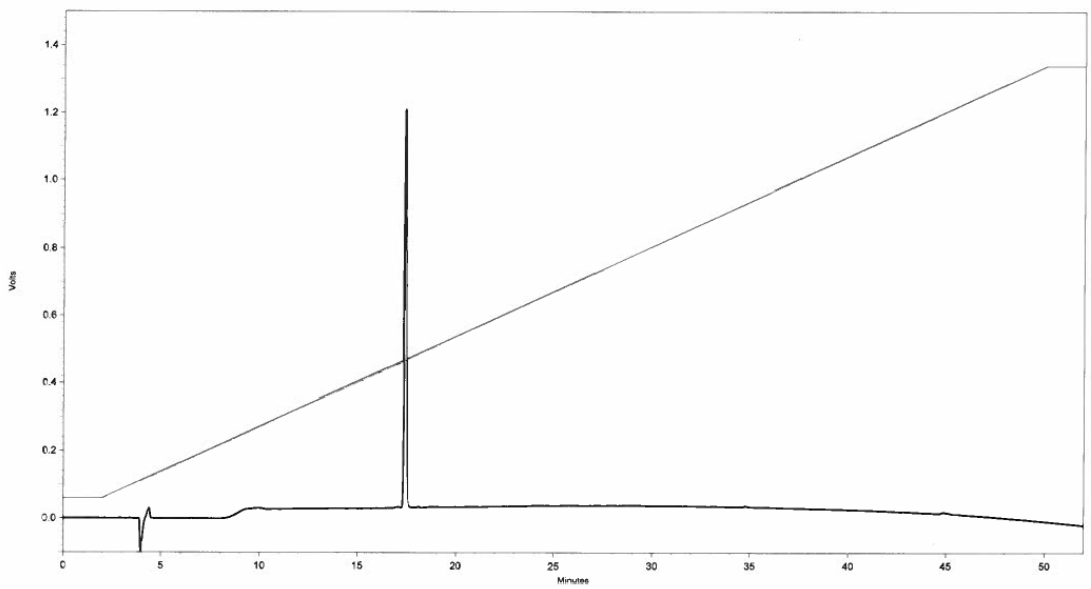

#### General Materials and Methods for Enzymatic Synthesis

Enzymatic reactions were incubated overnight at 37 °C with gentle shaking. The progress of the reactions was monitored by LCMS. In case of incomplete conversion after 18h, additional enzyme was added and incubated overnight. Enzymes and BSA were removed from the finished reactions by centrifugation using Vivaspin 2 centrifugal concentrators (PES, 10kDa cut off). The filtrate was lyophilized and purified by RP-C18 HPLC ($H_2O$+0.1%HCOOH:ACN+0.1%HCOOH, 5-25%, 25ml/min, 30min). (for sulfates $H_2O$+$NH_4COOH$ (pH 8.5): ACN 5-25%, 25 ml/min, 30 min)

**General Procedure I - Installation of β1,3-Gal using C1GalT1**

FVT*IG (**1,** 1 eq) and UDP-Gal (1.5 eq) were dissolved at a final FVT*IG (**1**) concentration of 10mM in a TRIS buffer (100mM, pH 7.5) containing $MnCl_2$ (10mM). To this mixture calf intestine alkaline phosphatase (CIAP, 1 U/µL, 1 U per µmol of added nucleotide) and C1GalT1 (12 µg per µmol acceptor) were added and incubated at 37 °C overnight with gentle shaking. The progress of the reaction was monitored by LCMS. The finished reaction was lyophilized, spin filtered and purified by preparative RP-C18 HPLC giving the O-glycan core 1 (**2**).

**General Procedure II - Installation of β1,6-GlcNAc using GCNT1**

O-Glycan core 1 (**2**, 1 eq) and UDP-GlcNAc (1.5 eq) were dissolved at a final O-glycan core 1 (**2**) concentration of 10mM in TRIS buffer (100mM, pH 7.5). To this mixture calf intestine alkaline phosphatase (CIAP, 1 U/µL, 1 U per µmol of added nucleotide), BSA (0.1% wt/wt) and GCNT1 (10 µg per µmol acceptor) were added and incubated at 37 °C overnight. Following purification, O-glycan core 2 (**3**) was obtained as a white solid.

**General Procedure III - Installation of β1,4-GlcNAc using B4GalT1**

The acceptor (1 eq) and UDP-Gal (1.5 eq) were dissolved at a final acceptor concentration of 10mM in a TRIS buffer (100mM, pH 7.5) containing $MnCl_2$ (10mM). To this mixture calf intestine alkaline phosphatase (CIAP, 1 U/µL, 1 U per µmol of added nucleotide), BSA (0.1% wt/wt) and B4GalT1 (10 µg per µmol acceptor) were added and incubated at 37 °C overnight followed by purification.

**General Procedure IV - Installation of α2,3-Neu5Ac using ST3Gal1**

The acceptor (1 eq) and CMP-NANA (1.5 eq) were dissolved at a final acceptor concentration of 10mM in a TRIS buffer (100mM, pH 7). To this mixture calf intestine alkaline phosphatase (CIAP, 1 U/µL, 1 U per µmol of added nucleotide), BSA (0.1% wt/wt) and ST3Gal1 (10 µg per µmol acceptor) were added and incubated at 37 °C overnight followed by purification.

**General Procedure V - Installation of α2,3-Neu5Ac using ST3Gal4**

The acceptor (1 eq) and CMP-NANA (1.5 eq) were dissolved at a final acceptor concentration of 10mM in a TRIS buffer (100mM, pH 7). To this mixture calf intestine alkaline phosphatase (CIAP, 1 U/µL, 1 U per µmol of added nucleotide), BSA (0.1% wt/wt) and ST3Gal4 (10 µg per µmol acceptor) were added and incubated at 37 °C overnight followed by purification.

**General Procedure VI - Installation of α1,3-Fuc using FUT5 or FUT6**

The acceptor (1 eq) and GDP-Fuc (1.5 eq.) were dissolved at a final acceptor concentration of 10mM in a TRIS buffer (50mM, pH 7.5) containing $MnCl_2$ (10mM). To this mixture calf intestine alkaline phosphatase (CIAP, 1 U/µL, 1 U per µmol of added nucleotide), BSA (0.1% wt/wt) and FUT5 or FUT6 (20 µg per µmol acceptor) were added and incubated at 37 °C overnight followed by purification.

**General Procedure VII - Installation of a $SO_3$-group on the 6-position of GlcNAc using CHST2**

The acceptor (1 eq) and PAPS (1.6 eq.) were dissolved at a final acceptor concentration of 10mM in a TRIS buffer (50mM, pH 7.0) containing $MgCl_2$ (10mM). To this mixture CHST2 (50 µg per µmol acceptor) was added and incubated at 37 °C overnight followed by purification.

**General Procedure VIII - Installation of β1,4-GlcNAc using B4GalT4**

The acceptor (1 eq) and UDP-Gal (1.5 eq) were dissolved at a final acceptor concentration of 10mM in a TRIS buffer (100mM, pH 7.5) containing $MnCl_2$ (10mM). To this mixture calf intestine alkaline phosphatase (CIAP, 1 U/µL, 1 U per µmol of added nucleotide), BSA (0.1% wt/wt) and B4GalT4 (10 µg per µmol acceptor) were added and incubated at 37 °C overnight followed by purification.

**General Procedure IX - Installation of a $SO_3$-group on the 6-position of Gal using CHST1**

The acceptor (1 eq) and PAPS (2 eq.) were dissolved at a final acceptor concentration of 10mM in a TRIS buffer (50mM, pH 7.0) containing $MgCl_2$ (10mM). To this mixture CHST1 (100 µg per µmol acceptor) was added and incubated at 37 °C overnight followed by purification.

**NMR Nomenclature**

Glycan assignments were made by numbering each monosaccharide starting from the reducing terminus and continuing in sequential order. The lower branch was counted before residues on the upper branch.

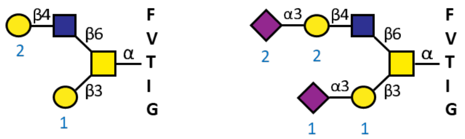

Indication of numbers dedicated to Gal, NeuAc, Fuc and GalNAc

**Enzymatic synthesis of the Core Structures and their derivatives**

**Compound 2 (Core 1)**

Starting from Tn-antigen (**1**, 12 mg, 16 µmol) following general procedure **I**, O-glycan core 1 (**2**) was obtained as a white solid (11.8 mg, 81%). $^1$H (600MHz, $D_2O$): δ = 7.40-7.35 (m, 2H, $CH_{Ar}$-Phe), 7.34-7.30 (m, 1H, $CH_{Ar}$-Phe), 7.26-7.21 (m, 2H, $CH_{Ar}$-Phe), 4.84 (d, J = 3.9 Hz, 1H, H1-GalNAc), 4.62-4.59 (m, 1H, $CH_{\alpha}$-Thr), 4.42 (d, J = 7.8 Hz, 1H, H1-Gal), 4.33- 4.24 (m, 3H, $CH_{\beta}$-Thr, $CH_{\alpha}$-Val, H2-GalNAc), 4.23-4.20 (m, 2H, H4-GalNAc, $CH_{\alpha}$-Ile), 4.09-4.05 (m, 1H, H5-GalNAc), 4.01 (dd, J = 2.0, 11.1 Hz, 1H, H3-GalNAc), 3.97-3.92 (m, 2H, $CH_{\alpha}$-Phe, $CH_2$-Gly), 3.90 (d, J = 3.4 Hz, 1H, H4-Gal), 3.84 (d, J = 16.8 Hz, 1H, $CH_2$-Gly), 3.80-3.71 (m, 4H, $H6_{a,b}$-GalNAc, $H6_{a,b}$-Gal), 3.64 (dd, J = 4.3, 7.8 Hz, 1H, H5-Gal), 3.60 (dd, J = 3.3, 9.8 Hz, 1H, H3-Gal), 3.53-3.49 (m, 1H, H2-Gal), 3.06 (dd, J = 6.6, 13.9 Hz, 1H, $CH_2$-Phe), 3.01 (dd, J = 6.6, 13.9 Hz, 1H, $CH_2$-Phe), 2.06- 1.98 (m, 1H, $CH_{\beta}$-Val), 2.01 (s, 3H, NHAc), 1.90-1.82 (m, 1H, $CH_{\beta}$-Ile), 1.56-1.45 8m, 1H, $CH_2$-Ile), 1.30 (d, J = 6.5 Hz, 3H, $CH_3$-Thr), 1.25-1.15 (m, 1H, $CH_2$-Ile), 0.98-0.91 (m, 9H, 2x$CH_3$-Val, $CH_3$-Ile), 0.87 (t, J = 7.4 Hz, 3H, $CH_3$-Ile). $^{13}$C from HSQC (150MHz, $D_2O$): δ = 129.4 (d, 2C), 128.7 (d, 2C), 127.3 (d, 1C), 104.7 (d, 1C), 98.9 (d, 1C), 77.7 (d, 1C), 76.4 8 (d, 1C), 75.2 (d, 1C),72.8 (d, 1C), 71.0 (d, 1C), 71.2

(d, 1C), 69.0 (d, 1C), 68.6 (d, 1C),61.3 (t, 2C), 59.4 (d, 1C), 58.3 (d, 1C), 57.08 (d, 1C), 55.2 (d, 1C) 48.5 (d, 1C), 42.0 (t, 1C), 39.4 (t, 1C), 36.4 (d, 1C), 30.3 (d, 1C), 24.7 (t, 1C), 22.5 (q, 1C), 18.8 (q, 2C), 18.2 (q, 1C), 14.8 (q, 1C), 10.3 (q, 1C). ESI-MS calcd for $C_{40}H_{66}N_7O_{16}^+$ $[M+H]^+$ 900.4561, found 900.4556.

**Compound 3 (Core 2)**

Starting from core 1 (**2**, 10 mg, 11 μmol) following general procedure **II**, O-glycan core 2 (**3**) was obtained as a white solid (11.7 mg, 96%). $^1H$ (600MHz, $D_2O$): δ = 7.44-7.31 (m, 3H, $CH_{Ar}$-Phe), 7.28-7.20 (m, 2H, $CH_{Ar}$-Phe), 4.82 (d, J = 3.9 Hz, 1H, H1-GalNAc), 4.60 (d, J = 2.6 Hz, 1H, $CH_\alpha$-Thr), 4.55 (d, J = 8.5 Hz, 1H, H1-GlcNAc), 4.41 (d, J = 7.7 Hz, 1H, H1-Gal), 4.27 (d, J = 8.0 Hz, 1H, $CH_\alpha$-Val), 4.24 (d, J = 3.8 Hz, 1H, H2-GalNAc), 4.23 – 4.19 (m, 3H, $CH_\alpha$-Ile, $CH_\beta$-Thr, H4-GalNAc), 4.16 (dd, J = 3.2, 8.5 Hz, 1H, H5-GalNAc), 4.10 (dd, J = 3.2, 10.8 Hz, 1H, H6a-GalNAc), 4.00 (dd, J = 3.1, 11.0 Hz, 1H, H3-GalNAc), 3.99-3.93 (m, 2H, $CH_\alpha$-Phe, $H6_a$-GlcNAc), 3.93-3.89 (m, 2H, $CH_2$-Gly, H4-Gal), 3.85-3.80 (m, 1H, $CH_2$-Gly), 3.80-3.68 (m, 5H, H2, $H6_b$-GlcNAc, $H6_{a,b}$-Gal, $H6_b$-GalNAc), 3.63 (m, 1H, H5-Gal), 3.60 (dd, J = 3.5, 9.9 Hz, 1H, H3-Gal), 3.55-3.46 (m, 3H, H3-GlcNAc, H2-Gal, H5-GlcNAc), 3.45-3.40 (m, 1H, H4-GlcNAc), 3.07 (dd, J = 6.6, 13.9 Hz, 1H, $CH_2$-Phe), 3.03 (dd, J = 6.6, 13.9 Hz, 1H, $CH_2$-Phe), 2.06 (s, 3H, NHAc), 2.02 (s, 3H, NHAc), 2.05-1.98 (m, 1H, $CH_\beta$-Val), 1.89-1.81 (m, 1H, $CH_\beta$-Ile), 1.54-1.45 (m, 1H, $CH_2$-Ile), 1.26 (d, J = 6.4 Hz, 3H, $CH_3$-Thr), 1.24-1.16 (m, 1H, $CH_2$-Ile), 0.99-0.91 (m, 9H, 2x$CH_3$-Val, $CH_3$-Ile), 0.87 (t, J = 7.4 Hz, 3H, $CH_3$-Ile). $^{13}C$ from HSQC (150MHz, $D_2O$): δ = 129.5 (d, 2C), 128.9 (d, 2C), 127.2 (d, 1C), 104.6 (d, 1C), 101.1 (d, 1C), 99.4 (d, 1C), 76.9 (d, 1C), 76.7 (d, 1C), 76.0 (d, 1C), 75.1 (d, 1C), 74.3 (d, 1C), 72.3 (d, 1C), 70.5 (d, 1C), 70.2 (t, 1C), 69.7 (d, 1C), 69.4 (d, 1C), 69.1 (d, 1C), 68.4 (d, 1C), 60.8 (t, 2C), 59.1 (d, 1C), 58.2 (d, 1C), 56.7 (d, 1C), 55.1 (d, 1C), 54.9 (d, 1C), 48.1 (d, 1C), 41.9 (t, 1C), 39.0 (t, 1C), 36.4 (d, 1C), 30.8 (d, 1C), 24.4 (t, 1C), 22.2 (q, 1C), 22.1 (q, 1C), 18.2 (q, 1C), 18.1 (q, 2C), 14.3 (q, 1C), 10.1 (q, 1C). ESI-MS calcd for $C_{48}H_{79}N_8O_{21}^+$ $[M+H]^+$ 1103.5355, found 1103.5363.

**Compound 4**

Staring from core 2 (**3**, 1.1 mg, 1.0 μmol) following general procedure **III** gave **4** (1.1 mg, 90%) as a white solid. $^1H$ (600MHz, $D_2O$): δ = 7.45-7.35 (m, 3H, $CH_{Ar}$-Phe), 7.29-7.24 (m, 2H, $CH_{Ar}$-Phe), 4.89 (d, J = 3.8 Hz, 1H, H1-GalNAc), 4.61 (d, J = 1.8 Hz, 1H, $CH_\alpha$-Thr), 4.58 (d, J = 8.5 Hz, 1H, H1-GlcNAc), 4.44 (d, J = 7.9 Hz, 1H, H1-Gal-2), 4.38 (d, J = 7.7 Hz, 1H, H1-Gal-1), 4.37-4.33 (m, 2H, $CH_\alpha$-Val, $CH_\alpha$-Phe), 4.29-4.18 (m, 4H, H2, H4-GalNAc, $CH_\alpha$-Ile, $CH_\beta$-Thr), 4.17 (dd, J = 3.2, 8.4 Hz, 1H, H5-GalNAc), 4.11 (dd, J = 3.2, 10.8 Hz, 1H, $H6_a$-GalNAc), 4.03-3.97 (m, 2H, H3-GalNAc, $H6_a$-GlcNAc), 3.92 (d, J= 3.5 Hz, 1H, H4-Gal-2), 3.90 (d, J= 3.5 Hz, 1H, H4-Gal-1), 3.86-3.47 (m, 18H, H2-$H6_b$- GlcNAc, $CH_2$-Gly, H2-H3, H5-$H6_{a,b}$-Gal-1, H2-H3, H5-$H6_{a,b}$-Gal-2, $H6_b$-GalNAc), 3.26 (dd, J = 6.8, 14.3 Hz, 1H, $CH_2$-Phe), 3.20 (dd, J = 6.8, 14.3 Hz, 1H, $CH_2$-Phe), 2.07-2.01 (m, 1H, $CH_\beta$-Val), 2.06 (s, 3H, NHAc), 2.00 (s, 3H, NHAc), 1.92-1.84 (m, 1H, $CH_\beta$-Ile), 1.54-1.45 (m, 1H, $CH_2$-Ile), 1.30 (d, J = 6.3 Hz, 3H, $CH_3$-Thr), 1.24-1.16 (m, 1H, $CH_2$-Ile), 1.02-0.92 (m, 9H, 2x$CH_3$-Val, $CH_3$-Ile), 0.87 (t, J = 7.4 Hz, 3H, $CH_3$-Ile). $^{13}C$ from HSQC (150MHz, $D_2O$): δ = 129.3 (d, 2C), 128.9 (d, 3C), 104.4 (d, 1C), 102.8 (d, 1C), 101.0 (d, 1C), 99.3 (d, 1C), 78.3 (d, 1C), 77.1 (d, 1C), 76.7 (d, 1C), 75.4 (d, 1C), 74.8 (d, 2C), 72.6 (d, 2C), 72.3 (d, 1C), 70.8 (d, 1C), 70.4 (d, 1C), 69.9 (t, 1C), 69.5 (d, 1C), 69.3 (d, 1C), 68.9 (d, 2C), 60.9 (t, 2C), 60.0 (t, 1C), 59.0 (d, 1C),

58.2 (d, 1C), 57.0 (d, 1C), 54.9 (d, 1C), 53.9 (d, 1C), 47.9 (d, 1C), 43.1 (t, 1C), 36.8 (t, 1C), 36.6 (d, 1C), 30.4 (d, 1C), 24.2 (t, 1C), 22.4 (q, 1C), 22.3 (q, 1C), 18.3 (q, 1C), 18.0 (q, 2C), 14.5 (q, 1C), 9.9 (q, 1C). ESI-MS calcd for $C_{54}H_{87}N_8O_{26}^-$ [M-H]$^-$ 1263.5737, found 1263.5711.

**Compound 5**

Starting from compound **4** (1 mg, 0.8 µmol) following general procedure V, compound **5** (0.9 mg, 73%) was obtained as a white solid. $^1$H (600MHz, $D_2O$): δ = 7.43-7.34 (m, 3H, $CH_{Ar}$-Phe), 7.29-7.21 (m, 2H, $CH_{Ar}$-Phe), 4.83 (d, J = 3.1 Hz, 1H, H1-GalNAc), 4.62 (d, J = 2.3 Hz, 1H, $CH_\alpha$-Thr), 4.57 (d, J = 8.5 Hz, 1H, H1-GlcNAc), 4.49 (d, J = 7.9 Hz, 1H, H1-Gal-2), 4.40 (d, J = 7.8 Hz, 1H, H1-Gal-1), 4.33 (d, J = 7.7 Hz, 1H, $CH_\alpha$-Val), 4.26-4.14 (m, 5H, H2, H4-H5-GalNAc, $CH_\alpha$-Ile, $CH_\beta$-Thr), 4.11-4.06 (m, 2H, $H6_a$-GalNAc, H3-Gal-2), 4.04-3.98 (m, 2H, H3-GalNAc, $H6_a$-GlcNAc), 3.96 -3.54 (m, 26, $CH_\alpha$-Phe, H2, H4-$H6_{a,b}$-Gal-2, H3-$H6_{a,b}$ -Gal-1, $CH_2$-Gly, H4-$H9_{a,b}$ -NeuAc, H2-$H6_b$-GlcNAc, $H6_b$-GalNAc), 3.50 (dd, J = 7.8, 9.9 Hz, 1H, H2-Gal-1), 3.25-3.04 (m, 2H, $CH_2$-Phe), 2.76 (dd, J = 4.6, 12.34 Hz, 1H, $H3_a$-NeuAc), 2.07-2.01 (m, 1H, $CH_\beta$-Val), 2.05 (s, 3H, NHAc), 2.03 (s, 3H, NHAc), 2.00 (s, 3H, NHAc), 1.92-1.84 (m, 1H, $CH_\beta$-Ile), 1.80 (t, J = 12.1 Hz, 1H, $H3_b$-NeuAc), 1.54-1.45 (m, 1H, $CH_2$-Ile), 1.29 (d, J = 6.3 Hz, 3H, $CH_3$-Thr), 1.24-1.16 (m, 1H, $CH_2$-Ile), 1.02-0.92 (m, 9H, 2x$CH_3$-Val, $CH_3$-Ile), 0.87 (t, J = 7.4 Hz, 3H, $CH_3$-Ile). $^{13}$C from HSQC (150MHz, $D_2O$): δ = 129.5 (d, 2C), 128.9 (d, 2C), 127.7 (d, 1C), 104.8 (d, 1C), 102.6 (d, 1C), 101.0 (d, 1C), 99.1 (d, 1C), 78.8 (d, 1C), 78.4 (d, 1C), 76.8 (d, 1C), 75.3 (d, 1C), 75.1 (d, 1C), 74.8 (d, 1C), 73.1 (d, 1C), 72.7 (d, 2C), 72.3 (d, 1C), 71.5 (d, 1C), 70.4 (d, 1C), 69.8 (t, 1C), 69.7 (d, 1C), 69.4 (d, 1C), 69.0 (d, 1C), 68.6 (d, 1C), 68.3 (d, 1C), 68.1 (d, 1C), 67.5 (d, 1C), 62.4 (t, 1C), 61.0 (t, 2C), 60.1 (t, 1C), 59.1 (d, 1C), 58.1 (d, 1C), 57.0 (d, 1C), 55.0 (d, 1C), 51.4 (d, 1C), 48.3 (d, 1C), 41.9 (t, 1C), 39.6 (t, 1C), 36.5 (d, 1C), 30.5 (d, 1C), 24.3 (t, 1C), 22.1 (q, 3C), 18.4 (q, 1C), 18.1 (q, 2C), 14.6 (q, 1C), 10.1 (q, 1C)*. ESI-MS calcd for $C_{65}H_{104}N_9O_{34}^-$ [M-H]$^-$ 1554.6691, found 1554.6681.

* signal of $CH_2$-Phe and $CH_\alpha$-Phe are not visible in HSQC

**Compound 6**

Starting from compound **5** (660 µg, 0.4 µmol) following general procedure VI (FUT5) compound **6** (670 µg, 93%) was obtained as a white solid. $^1$H (600MHz, $D_2O$): δ = 7.42-7.30 (m, 3H, $CH_{Ar}$-Phe), 7.20-7.19 (m, 2H, $CH_{Ar}$-Phe), 5.09 (d, J = 3.9 Hz, 1H, H1-Fuc), 4.84-4.80 (m, 2H, H1-GalNAc, H5-Fuc), 4.60 (d, J = 2.2 Hz, 1H, $CH_\alpha$-Thr), 4.57 (d, J = 8.4 Hz, 1H, H1-GlcNAc), 4.48 (d, J = 7.8 Hz, 1H, H1-Gal-2), 4.42 (d, J = 7.7 Hz, 1H, H1-Gal-1), 4.27 (d, J = 8.1 Hz, 1H, $CH_\alpha$-Val), 4.25-4.13 (m, 5H, H2, H4-H5-GalNAc, $CH_\alpha$-Ile, $CH_\beta$-Thr), 4.11-4.05 (m, 2H, $H6_a$-GalNAc, H3-Gal-2), 4.04-3.98 (m, 2H, H3-GalNAc, $H6_a$-GlcNAc), 3.96 -3.58 (m, 30, $CH_\alpha$-Phe, H2, H4-$H6_{a,b}$-Gal-2, H2-$H6_{a,b}$ -Gal-1, $CH_2$-Gly, H4-$H9_{a,b}$ -NeuAc, H2-$H6_b$-GlcNAc, $H6_b$-GalNAc, H2-H4-Fuc), 3.09-2.95 (m, 2H, $CH_2$-Phe), 2.77 (dd, J = 4.6, 12.4 Hz, 1H, $H3_a$-NeuAc), 2.07-1.97 (m, 1H, $CH_\beta$-Val), 2.05 (s, 3H, NHAc), 2.03 (s, 3H, NHAc), 2.00 (s, 3H, NHAc), 1.88-1.82 (m, 1H, $CH_\beta$-Ile), 1.79 (t, J = 12.1 Hz, 1H, $H3_b$-NeuAc), 1.54-1.45 (m, 1H, $CH_2$-Ile), 1.25 (d, J = 6.4 Hz, 3H, $CH_3$-Thr), 1.24-1.16 (m, 1H, $CH_2$-Ile), 1.16 (d, J = 6.3 Hz, 3H, $CH_3$-Fuc), 0.98-0.91 (m, 9H, 2x$CH_3$-Val, $CH_3$-Ile), 0.87 (t, J = 7.4 Hz, 3H, $CH_3$-Ile). $^{13}$C from HSQC (150MHz, $D_2O$): δ = 129.4 (d, 2C), 129.1 (d, 2C), 127.8 (d, 1C), 104.6 (d, 1C), 101.6 (d, 1C), 101.0 (d, 1C), 99.1 (d, 1C), 98.6 (d, 1C), 76.9 (d, 2C), 75.6 (d, 1C), 75.5 (d,

1C), 75.1 (d, 1C), 74.9 (d, 1C), 74.8 (d, 1C), 73.4 (d, 1C), 72.7 (d, 2C), 72.4 (d, 1C), 71.7 (d, 1C), 71.5 (d, 1C), 70.5 (d, 1C), 70.1 (t, 1C), 69.4 (d, 2C), 69.5 (d, 1C), 69.2 (d, 1C), 68.5 (d, 1C), 68.1 (d, 1C), 67.9 (d, 1C), 67.4 (d, 2C), 66.7 (d, 1C), 62.4 (t, 1C), 60.6 (t, 2C), 59.6 (t, 1C), 59.2 (d, 1C), 57.9 (d, 1C), 56.9 (d, 1C), 55.5 (d, 1C), 51.5 (d, 1C), 48.2 (d, 1C), 41.6 (t, 1C), 39.8 (t, 1C), 36.6 (d, 1C), 30.5 (d, 1C), 24.5 (t, 1C), 22.3 (q, 2C), 22.0 (q, 1C), 18.6 (q, 1C), 18.4 (q, 2C), 15.0 (q, 1C), 14.7 (q, 1C), 9.9 (q, 1C)*. ESI-MS calcd for $C_{71}H_{116}N_9O_{38}^+$ $[M+H]^+$ 1702.7421, found 1702.7414.

* Signal of $CH_2$-Phe and $CH_\alpha$-Phe are not visible in HSQC

**Compound 7**

Starting from core 2 (**3**, 4 mg, 3.6 μmol) following general procedure IV, compound **7** (4.3 mg, 85%) was obtained. $^1H$ (600MHz, $D_2O$): δ = 7.44-7.31 (m, 3H, $CH_{Ar}$-Phe), 7.28-7.20 (m, 2H, $CH_{Ar}$-Phe), 4.83 (d, J = 3.6 Hz, 1H, H1-GalNAc), 4.64-4.61 (m, 1H, $CH_\alpha$-Thr), 4.56 (d, J = 8.6 Hz, 1H, H1-GlcNAc), 4.47 (d, J = 7.8 Hz, 1H, H1-Gal), 4.39-4.34 (m, 2H, $CH_\alpha$-Val, $CH_\alpha$-Phe), 4.27-4.18 (m, 4H, H2-GalNAc, $CH_\alpha$-Ile, $CH_\beta$-Thr, H4-GalNAc), 4.16 (dd, J = 2.9, 8.5 Hz, 1H, H5-GalNAc), 4.13-4.09 (m, 1H, $H6_a$-GalNAc), 4.05 (dd, J = 2.9, 9.8 Hz, 1H, H3-Gal), 4.02 (dd, J = 3.0, 11.0 Hz, 1H, H3-GalNAc), 3.97-3.92 (m, 3H, H4-Gal, $H6_a$-GlcNAc, $CH_2$-Gly), 3.91-3.80 (m, 4H, H5-,H8-, $H9_a$-NeuAc, $CH_2$-Gly), 3.77-3.67 (m, 6H, $H6_b$-GlcNAc, $H6_{a,b}$-Gal, H2-GlcNAc, $H6_b$-GalNAc, H4-NeuAc), 3.66-3.58 (m, 4H, H6-, H7-, $H9_b$-NeuAc, H5-Gal), 3.56-3.46 (m, 3H, H3, H5-GlcNAc, H2-Gal), 3.45-3.40 (m, 1H, H4-GlcNAc), 3.26 (dd, J = 6.7, 14.3 Hz, 1H, $CH_2$-Phe), 3.20 (dd, J = 6.7, 14.3 Hz, 1H, $CH_2$-Phe), 2.76 (dd, J = 4.7, 12.5, 1H, $H3_a$-NeuAc), 2.06 (s, 3H, NHAc), 2.04 (s, 3H, NHAc), 2.02 (s, 3H, NHAc), 2.07-2.01 (m, 1H, $CH_\beta$-Val), 1.90-1.82 (m, 1H, $CH_\beta$-Ile), 1.79 (t, J = 12.1, 1H, $H3_b$-NeuAc), 1.54-1.45 (m, 1H, $CH_2$-Ile), 1.28 (d, J = 6.3 Hz, 3H, $CH_3$-Thr), 1.24-1.16 (m, 1H, $CH_2$-Ile), 0.99-0.91 (m, 9H, 2x$CH_3$-Val, $CH_3$-Ile), 0.87 (t, J = 7.4 Hz, 3H, $CH_3$-Ile). $^{13}C$ from HSQC (150MHz, $D_2O$): δ = 129.4 (d, 2C), 129.2 (d, 3C), 104.6 (d, 1C), 101.1 (d, 1C), 99.2 (d, 1C), 77.2 (d, 1C), 76.9 (d, 1C), 76.0 (d, 1C), 75.5 (d, 1C), 74.4 (d, 1C), 74.1 (d, 1C), 72.8 (d, 1C), 71.8 (d, 1C), 70.3 (t, 1C), 69.7 (d, 1C), 69.6 (d, 1C), 68.9 (d, 1C), 68.8 (d, 1C), 68.4 (d, 2C), 67.2 (d, 1C), 62.2 (t, 1C), 60.5 (t, 2C), 58.6 (d, 1C), 57.8 (d, 1C), 56.6 (d, 1C), 55.4 (d, 1C), 53.6 (d, 1C), 51.40 (d, 1C), 47.8 (d, 1C), 42.1 (t, 1C), 39.8 (t, 1C), 36.4 (t, 1C), 36.3 (d, 1C), 30.3 (d, 1C), 24.4 (t, 1C), 22.3 (q, 3C), 18.3 (q, 1C), 18.0 (q, 2C), 14.8 (q, 1C), 10.3 (q, 1C). ESI-MS calcd for $C_{59}H_{96}N_9O_{29}^+$ $[M+H]^+$ 1394.6309, found 1394.6312.

**Compound 8**

Starting from compound **7** (3.7 mg, 2.7 μmol) following general procedure **III** gave **8** (2.8 mg, 68%). $^1H$ (600MHz, $D_2O$): δ = 7.46-7.37 (m, 3H, $CH_{Ar}$-Phe), 7.31-7.24 (m, 2H, $CH_{Ar}$-Phe), 4.83 (d, J = 3.8 Hz, 1H, H1-GalNAc), 4.63 (d, J = 2.2 Hz, 1H, $CH_\alpha$-Thr), 4.57 (d, J = 8.5 Hz, 1H, H1-GlcNAc), 4.47 (d, J = 7.8 Hz, 1H, H1-Gal-1), 4.42 (d, J = 7.9 Hz, 1H, H1-Gal-2), 4.39-4.33 (m, 2H, $CH_\alpha$-Val, $CH_\alpha$-Phe), 4.28-4.18 (m, 4H, H2, H4-GalNAc, $CH_\alpha$-Ile, $CH_\beta$-Thr), 4.17 (dd, J = 2.5, 8.9 Hz, 1H, H5-GalNAc), 4.11 (dd, J = 2.9, 10.9 Hz, 1H, $H6_a$-GalNAc), 4.05 (dd, J = 3.2, 9.9 Hz, 1H, H3-Gal-1), 4.03-3.97 (m, 2H, H3-GalNAc, $H6_a$-GlcNAc), 3.96-3.87 (m, 4H, H4-Gal-1, H4-Gal-2, $CH_2$-Gly, H8-NeuAc), 3.86-3.58 (m, 20H, H4-H7, $H9_{a,b}$-NeuAc, $CH_2$-Gly, H2-$H6_b$- GlcNAc, H5-$H6_{a,b}$-Gal-1, H3, H5-$H6_{a,b}$-Gal-2, $H6_b$-GalNAc), 3.56-3.50 (m, 2H, H2-Gal-1, H2-Gal-2), 3.26 (dd, J = 6.8, 14.3 Hz, 1H, $CH_2$-Phe), 3.20 (dd, J = 6.8, 14.3 Hz, 1H, $CH_2$-Phe),

2.76 (dd, J = 4.7, 12.5, 1H, H3$_a$-NeuAc), 2.07 (s, 3H, NHAc), 2.03 (s, 3H, NHAc), 2.01 (s, 3H, NHAc), 2.07-2.01 (m, 1H, CH$_\beta$-Val), 1.89-1.82 (m, 1H, CH$_\beta$-Ile), 1.79 (t, J = 12.1, 1H, H3$_b$-NeuAc), 1.54-1.45 (m, 1H, CH$_2$-Ile), 1.30 (d, J = 6.4 Hz, 3H, CH$_3$-Thr), 1.24-1.16 (m, 1H, CH$_2$-Ile), 1.02-0.92 (m, 9H, 2xCH$_3$-Val, CH$_3$-Ile), 0.87 (t, J = 7.4 Hz, 3H, CH$_3$-Ile). $^{13}$C from HSQC (150MHz, D$_2$O): δ = 129.4 (d, 2C), 128.9 (d, 3C), 104.2 (d, 1C), 103.0 (d, 1C), 101.4 (d, 1C), 99.08 (d, 1C), 78.2 (d, 1C), 77.1 (d, 1C), 76.6 (d, 1C), 75.8 (d, 1C), 75.4 (d, 1C), 74.7 (d, 2C), 72.7 (d, 1C), 72.5 (d, 2C), 72.0 (d, 1C), 70.9 (d, 1C), 70.0 (t, 1C), 69.3 (d, 1C), 68.8 (d, 2C), 68.2 (d, 1C), 68.1 (d, 1C), 66.8 (d, 1C), 66.4 (d, 1C), 62.5 (t, 1C), 60.9 (t, 2C), 59.8 (t, 1C), 59.2 (d, 1C), 57.6 (d, 1C), 57.0 (d, 1C), 54.7 (d, 1C), 54.0 (d, 1C), 51.7 (d, 1C), 48.1 (d, 1C), 41.88 (t, 1C), 39.6 (t, 1C), 36.62 (t, 1C), 36.3 (d, 1C), 30.4 (d, 1C), 24.5 (t, 1C), 22.3 (q, 2C), 22.1 (q, 1C), 18.5 (q, 1C), 17.9 (q, 2C), 14.6 (q, 1C), 10.0 (q, 1C). ESI-MS calcd for $C_{65}H_{106}N_9O_{34}^+$ [M+H]$^+$ 1556.6837, found 1556.6826.

**Compound 9**

Starting from compound **8** (1.5 mg, 0.96 μmol) following general procedure V compound **9** (1.7 mg, 97%) was obtained. $^1$H (600MHz, D$_2$O): δ = 7.49-7.38 (m, 3H, CH$_{Ar}$-Phe), 7.30-7.24 (m, 2H, CH$_{Ar}$-Phe), 4.83 (d, J = 3.9 Hz, 1H, H1-GalNAc), 4.63 (d, J = 2.1 Hz, 1H, CH$_\alpha$-Thr), 4.57 (d, J = 8.4 Hz, 1H, H1-GlcNAc), 4.48 (d, J = 7.8 Hz, 1H, H1-Gal-2), 4.47 (d, J = 7.7 Hz, 1H, H1-Gal-1), 4.39-4.34 (m, 2H, CH$_\alpha$-Val, CH$_\alpha$-Phe), 4.29-4.15 (m, 5H, H2, H4-H5-GalNAc, CH$_\alpha$-Ile, CH$_\beta$-Thr,), 4.13- 3.98 (m, 5H, H3, H6$_a$-GalNAc, H3-Gal-1, H3-Gal-2, H6$_a$-GlcNAc), 3.97-3.92 (m, 3H, CH$_2$-Gly, H4-Gal-1, H4-Gal-2), 3.91-3.81 (m, 8H, H5, H8-H9$_a$-NeuAc-1, H5, H8-H9$_a$-NeuAc-2, H6$_b$-GlcNAc, CH$_2$-Gly), 3.80-3.50 (m, 21H, H4, H6-H7, H9$_b$-NeuAc-1, H4, H6-H7, H9$_b$-NeuAc-2, H2-H5- GlcNAc, H2, H5-H6$_{a,b}$-Gal-1, H2, H5-H6$_{a,b}$-Gal-2, H6$_b$-GalNAc), 3.26 (dd, J = 6.8, 14.3 Hz, 1H, CH$_2$-Phe), 3.20 (dd, J = 6.8, 14.3 Hz, 1H, CH$_2$-Phe), 2.76 (dd, J = 4.7, 12.5, 2H, H3-NeuAc-1, H3-NeuAc-2), 2.06 (s, 3H, NHAc), 2.04 (s, 3H, NHAc), 2.03 (s, 3H, NHAc), 2.01 (s, 3H, NHAc), 2.07-2.01 (m, 1H, CH$_\beta$-Val), 1.89-1.82 (m, 1H, CH$_\beta$-Ile), 1.82-1.76 (m, 2H, H3-NeuAc-1, H3-NeuAc-2), 1.54-1.45 (m, 1H, CH$_2$-Ile), 1.30 (d, J = 6.4 Hz, 3H, CH$_3$-Thr), 1.24-1.16 (m, 1H, CH$_2$-Ile), 1.02-0.92 (m, 9H, 2xCH$_3$-Val, CH$_3$-Ile), 0.87 (t, J = 7.4 Hz, 3H, CH$_3$-Ile). $^{13}$C from HSQC (150MHz, D$_2$O): δ = 129.4 (d, 2C), 128.9 (d, 3C), 104.7 (d, 1C), 102.7 (d, 1C), 101.4 (d, 1C), 99.2 (d, 1C), 78.2 (d, 1C), 77.2 (d, 1C), 77.1 (d, 1C), 75.6 (d, 1C), 75.5 (d, 1C), 75.0 (d, 3C), 73.3 (d, 3C), 71.9 (d, 2C), 70.4 (t, 1C), 69.3 (d, 1C), 69.2 (d, 1C), 69.0 (d, 1C), 68.9 (d, 1C), 68.8 (d, 2C), 68.0 (d, 2C), 67.4 (d, 2C), 62.6 (t, 2C), 61.2 (t, 2C), 60.3 (t, 1C), 59.3 (d, 1C), 57.9 (d, 1C), 57.1 (d, 1C), 54.8 (d, 1C), 53.9 (d, 1C), 52.0 (d, 2C) 48.2 (d, 1C), 42.03 (t, 1C), 39.4 (t, 2C), 37.1 (t, 1C), 36.6 (d, 1C), 30.4 (d, 1C), 24.5 (t, 1C), 22.1 (q, 4C), 18.4 (q, 1C), 18.1 (q, 2C), 14.8 (q, 1C), 10.0 (q, 1C). ESI-MS calcd for $C_{76}H_{122}N_{10}O_{42}^-$ [M-H]$^-$ 1845.7645, found 1845.7626.

**Compound 10**

Starting from compound **8** (500μg, 0.3 μmol) following general procedure VI (FUT5) gave compound **10** (370 μg, 68%) as a white solid. $^1$H (600MHz, D$_2$O): δ = 7.46-7.37 (m, 3H, CH$_{Ar}$-Phe), 7.31-7.24 (m, 2H, CH$_{Ar}$-Phe), 5.10 (d, J = 4.1 Hz, 1H, H1-Fuc), 4.83 (d, 1H, H5-Fuc)*, 4.81 (d, 1H, H1-GalNAc)*, 4.60 (d, J = 2.2 Hz, 1H, CH$_\alpha$-Thr), 4.57 (d, J = 8.3 Hz, 1H, H1-GlcNAc), 4.49 (d, J = 8.0 Hz, 1H, H1-Gal-1), 4.42 (d, J = 7.8 Hz, 1H, H1-Gal-2), 4.30 (d, J = 8.1 Hz, 1H, CH$_\alpha$-Val), 4.25-4.18 (m, 4H, H2, H4-GalNAc,

$CH_{\alpha}$-Ile, $CH_{\beta}$-Thr), 4.17-4.12 (m, 1H, H5-GalNAc), 4.11-3.99 (m, 4H, H3,H6$_a$-GalNAc, H3-Gal-1, H6$_a$-GlcNAc), 3.96-3.77 (m, 14H, $CH_{\alpha}$-Phe, H4-Gal-1, H4-Gal-2, $CH_2$-Gly, H5, H8, H9$_a$ -NeuAc, H2-H4, H6$_b$-GlcNAc, H3-H4 Fuc), 3.75-3.47 (m, 16H, H4,H6-H7, H9$_b$-NeuAc, H5- GlcNAc, H2,H5-H6$_{a,b}$-Gal-1, H2-H3, H5-H6$_{a,b}$-Gal-2, H6$_b$-GalNAc, H2-Fuc), 3.14-2.98 (m, 2H, $CH_2$-Phe), 2.76 (dd, J = 4.6, 12.3, 1H, H3$_a$-NeuAc), 2.05 (s, 3H, NHAc), 2.03 (s, 3H, NHAc), 2.02 (s, 3H, NHAc), 2.07-2.01 (m, 1H, $CH_{\beta}$-Val), 1.89-1.82 (m, 1H, $CH_{\beta}$-Ile), 1.79 (t, J = 12.1, 1H, H3$_b$-NeuAc), 1.54-1.45 (m, 1H, $CH_2$-Ile), 1.26 (d, J = 6.4 Hz, 3H, $CH_3$-Thr), 1.24-1.16 (m, 1H, $CH_2$-Ile), 1.17 (d, J = 6.7 Hz, 3H, $CH_3$-Fuc), 1.02-0.92 (m, 9H, 2x$CH_3$-Val, $CH_3$-Ile), 0.87 (t, J = 7.4 Hz, 3H, $CH_3$-Ile). $^{13}$C from HSQC (150MHz, $D_2O$): δ = 129.4 (d, 2C), 128.9 (d, 3C), 104.6 (d, 1C), 102.0 (d, 1C), 100.8 (d, 1C), 99.3 (d,1C), 98.5 (d, 1C), 76.9 (d, 1C), 76.6 (d, 1C), 75.6 (d, 1C), 75.4 (d, 2C), 75.1 (d, 2C), 74.7 (d, 1C), 73.9 (d, 1C), 72.9 (d, 1C), 71.9 (d, 2C), 71.1 (d, 1C), 69.9 (t, 1C), 69.6 (d, 1C), 69.1 (d, 3C), 68.2 (d, 2C), 67.9 (d, 1C), 67.4 (d, 1C), 66.6 (d, 1C), 62.3 (t, 1C), 61.4 (d, 1C), 61.1 (t, 2C), 59.6 (t, 1C), 58.8 (d, 1C), 58.0 (d, 1C), 56.5 (d, 1C), 55.3 (d, 1C), 51.5 (d, 1C), 48.3 (d, 1C), 42.0 (t, 1C), 39.7 (t, 1C), 36.4 (d, 1C), 30.6 (d, 1C), 24.5 (t, 1C), 22.3 (q, 2C), 22.0 (q, 1C), 18.6 (q, 1C), 18.3 (q, 2C), 15.2 (q, 1C), 14.5 (q, 1C), 10.0 (q, 1C)**. ESI-MS calcd for $C_{71}H_{114}N_9O_{38}^-$ $[M-H]^-$ 1700.7270, found 1700.7251.

* Signal lies under $D_2O$ peak, ** signal of $CH_2$-Phe and $CH_{\alpha}$-Phe are not visible in HSQC

**Compound 11**

Starting compound **9** (500 µg, 0.27 µmol) following general procedure VI (FUT6) compound **11** (430 µg, 80%) was obtained. $^1$H (600MHz, $D_2O$): δ = 7.45-7.38 (m, 3H, $CH_{Ar}$-Phe), 7.30-7.24 (m, 2H, $CH_{Ar}$-Phe), 5.09 (d, J = 3.5 Hz, 1H, H1-Fuc), 4.81 (m, J = 171 Hz*, 2H, H1-GalNAc, H5-Fuc), 4.64-4.62 (m, 1H, $CH_{\alpha}$-Thr), 4.56 (d, J = 8.3 Hz, 1H, H1-GlcNAc), 4.49-4.45 (m, 2H, H1-Gal-1, H1-Gal-2), 4.39-4.33 (m, 2H, $CH_{\alpha}$-Val, $CH_{\alpha}$-Phe), 4.28-4.14 (m, 5H, $CH_{\alpha}$-Ile, $CH_{\beta}$-Thr, H2, H4-H5-GalNAc), 4.12-3.99 (m, 5H, H3, H6$_a$-GalNAc, H3-Gal-1, H3-Gal-2, H6$_a$-GlcNAc), 3.97-3.78 (m, 15H, H2-H4, H6$_b$ GlcNAc, $CH_2$-Gly, H4-Gal-1, H4-Gal-2, H5, H8-H9$_a$-NeuAc-1, H5, H8-H9$_a$-NeuAc-2, H3-Fuc), 3.77-3.46 (m, 20H, H2, H4-Fuc, H4, H6-H7, H9$_b$ NeuAc-1, H4, H6-H7, H9$_b$ NeuAc-2, H5-GlcNAc, H2, H5-H6$_{a,b}$-Gal-1, H2, H5-H6$_{a,b}$-Gal-2, H6$_b$-GalNAc), 3.23 (dd, J = 6.8, 14.3 Hz, 1H, $CH_2$-Phe), 3.20 (dd, J = 6.8, 14.3 Hz, 1H, $CH_2$-Phe), 2.75 (dd, J = 4.7, 12.5, 2H, H3$_a$-NeuAc-1, H3$_a$-NeuAc-2), 2.05 (s, 3H, NHAc), 2.04 (s, 3H, NHAc), 2.03 (s, 3H, NHAc), 2.01 (s, 3H, NHAc), 2.07-2.01 (m, 1H, $CH_{\beta}$-Val), 1.89-1.82 (m, 1H, $CH_{\beta}$-Ile), 1.82-1.76 (m, 2H, H3$_b$-NeuAc-1, H3$_b$-NeuAc-2), 1.54-1.45 (m, 1H, $CH_2$-Ile), 1.28 (d, J = 6.4 Hz, 3H, $CH_3$-Thr), 1.24-1.16 (m, 1H, $CH_2$-Ile), 1.15 (d, J= 6.6Hz, 3H, $CH_3$-Fuc), 1.02-0.92 (m, 9H, 2x$CH_3$-Val, $CH_3$-Ile), 0.87 (t, J = 7.4 Hz, 3H, $CH_3$-Ile). $^{13}$C from HSQC (150MHz, $D_2O$): δ = 129.6 (d, 2C), 128.8 (d, 3C), 104.2 (d, 1C), 101.7 (d, 1C), 101.1 (d, 1C), 99.3 (d, 1C), 98.6 (d, 1C), 76.9 (d, 2C), 75.4 (d, 1C), 75.1 (d, 3C), 75.0 (d, 1C), 74.8 (d, 1C), 73.6 (d, 1C), 73.0 (d, 2C), 72.1 (d, 2C), 71.8 (d, 2C), 70.3 (t, 1C), 69.7 (d, 1C), 69.1 (d, 1C), 68.9 (d, 3C), 68.1 (d, 2C), 67.7 (d, 2C), 67.2 (d, 2C), 66.6 (d, 1C), 62.5 (d, 2C), 61.1 (t, 2C) 59.8 (t, 1C), 59.4 (d, 1C), 59.2 (d, 1C), 57.8 (d, 1C), 56,9 (d, 1C), 55.5 (d, 1C), 53.7 (d, 1C), 51.8 (d, 2C), 48.1 (d,1C), 42.1 (t, 1C), 39.9 (t, 1C), 36.6 (t, 1C), 36.5 (d, 1C), 30.4 (d, 1C), 24.4 (t, 1C), 21.4 (q, 4C), 18.9 (q, 1C), 17.8 (q, 2C), 15.0 (q, 1C), 14.4 (q, 1C), 10.0 (q, 1C). ESI-MS calcd for $C_{82}H_{133}N_{10}O_{46}^+$ $[M+2H]^{2+}$ 997.4227, found 997.4200.

* Signal lies under $D_2O$ peak. The J-coupling was measured in a decoupled HSQC spectra.

**Compound 12**

Starting from compound **7** (2.9 mg, 2.1 µmol) following general procedure VII compound **12** (3.1 g, 90%) was obtained as a white solid. $^1$H (600MHz, $D_2O$): δ = 7.46-7.37 (m, 3H, $CH_{Ar}$-Phe), 7.31-7.23 (m, 2H, $CH_{Ar}$-Phe), 4.82 (d, 1H, H1-GalNAc)*, 4.64 (d, J = 2.1 Hz, 1H, $CH_\alpha$-Thr), 4.56 (d, J = 8.5 Hz, 1H, H1-GlcNAc), 4.47 (d, J = 7.8 Hz, 1H, H1-Gal), 4.40-4.31 (m, 3H, $CH_\alpha$-Val, $CH_\alpha$-Phe, $H6_a$-GlcNAc), 4.27-4.17 (m, 6H, H2, H4-H5-GalNAc, $CH_\alpha$-Ile, $CH_\beta$-Thr, $H6_b$-GlcNAc), 4.11-4.06 (m, 1H, $H6_a$-GalNAc), 4.05 (dd, J = 3.1, 9.8 Hz, 1H, H3-Gal), 4.02 (dd, J = 3.0, 11.0 Hz, 1H, H3-GalNAc), 3.96-3.58 (m, 16H, H4-$H6_{a,b}$ -Gal, H5-GlcNAc, $CH_2$-Gly, H4-$H9_{a,b}$-NeuAc, H2-GlcNAc, $H6_b$-GalNAc), 3.55-3.47 (m, 3H, H3-H4-GlcNAc, H2-Gal), 3.27 (dd, J = 6.7, 14.3 Hz, 1H, $CH_2$-Phe), 3.20 (dd, J = 6.7, 14.3 Hz, 1H, $CH_2$-Phe), 2.76 (dd, J = 4.6, 12.4, 1H, $H3_a$-NeuAc), 2.07 (s, 3H, NHAc), 2.05 (s, 3H, NHAc), 2.03 (s, 3H, NHAc), 2.07-2.01 (m, 1H, $CH_\beta$-Val), 1.90-1.82 (m, 1H, $CH_\beta$-Ile), 1.78 (t, J = 12.1, 1H, $H3_b$-NeuAc), 1.54-1.45 (m, 1H, $CH_2$-Ile), 1.31 (d, J = 6.3 Hz, 3H, $CH_3$-Thr), 1.24-1.16 (m, 1H, $CH_2$-Ile), 0.99-0.91 (m, 9H, 2x$CH_3$-Val, $CH_3$-Ile), 0.87 (t, J = 7.4 Hz, 3H, $CH_3$-Ile). $^{13}$C from HSQC (150MHz, $D_2O$): δ = 129.8 (d, 2C), 129.0 (d, 3C), 104.6 (d, 1C), 101.4 (d, 1C), 99.2 (d, 1C), 77.4 (d, 1C), 77.2 (d, 1C), 75.6 (d, 1C), 74.7 (d, 1C), 74.2 (d, 2C), 73.9 (d, 1C), 72.8 (d, 1C), 72.0 (d, 1C), 70.6 (t, 1C), 69.2 (d, 3C), 68.3 (d, 1C), 68.1 (d, 1C), 67.2 (d, 1C), 67.0 (t, 1C), 62.5 (t, 1C), 61.0 (t, 1C), 59.2 (d, 1C), 57.9 (d, 1C), 57.0 (d, 1C), 55.5 (d, 1C), 53.7 (d, 1C), 51.6 (d, 1C), 48.2 (d, 1C), 41.9 (t, 1C), 39.6 (t, 1C), 36.7 (t, 1C), 36.4 (d, 1C), 30.3 (d, 1C), 24.4 (t, 1C), 22.2 (q, 2C), 22.1 (q, 1C), 18.7 (q, 1C), 18.4 (q, 2C), 14.4 (q, 1C), 10.2 (q, 1C). ESI-MS calcd for $C_{59}H_{94}N_9O_{32}S^-$ $[M-H]^-$ 1472.5726, found 1472.5718.

* Signal lies under $D_2O$ peak

**Compound 13**

Starting from compound **12** (2.8 mg, 1.9 µmol) following general procedure VIII, compound **13** (2.8 mg, 90%) was obtained as a white solid. $^1$H (600MHz, $D_2O$): δ = 7.46-7.40 (m, 3H, $CH_{Ar}$-Phe), 7.31-7.23 (m, 2H, $CH_{Ar}$-Phe), 4.81 (d, 1H, H1-GalNAc)*, 4.64 (d, J = 2.0 Hz, 1H, $CH_\alpha$-Thr), 4.59 (d, J = 8.5 Hz, 1H, H1-GlcNAc), 4.49 (d, J = 7.8 Hz, 1H, H1-Gal-2), 4.47 (d, J = 7.9 Hz, 1H, H1-Gal-1), 4.42-4.30 (m, 4H, $CH_\alpha$-Val, $CH_\alpha$-Phe, $H6_{a,b}$-GlcNAc), 4.27-4.17 (m, 5H, H2, H4-H5-GalNAc, $CH_\alpha$-Ile, $CH_\beta$-Thr), 4.11-3.99 (m, 3H, H3, $H6_a$-GalNAc, H3-Gal-1), 3.97-3.57 (m, 23H, H3-$H6_{a,b}$-Gal-2, H4-$H6_{a,b}$ -Gal-1, H2-H5-GlcNAc, $CH_2$-Gly, H4-$H9_{a,b}$-NeuAc, $H6_b$-GalNAc), 3.55-3.47 (m, 2H, H2-Gal-1, H2-Gal-2), 3.26 (dd, J = 6.5, 14.4 Hz, 1H, $CH_2$-Phe), 3.19 (dd, J = 6.8, 14.3 Hz, 1H, $CH_2$-Phe), 2.76 (dd, J = 4.6, 12.4, 1H, $H3_a$-NeuAc), 2.06 (s, 3H, NHAc), 2.03 (s, 3H, NHAc), 2.01 (s, 3H, NHAc), 2.07-2.01 (m, 1H, $CH_\beta$-Val), 1.90-1.82 (m, 1H, $CH_\beta$-Ile), 1.78 (t, J = 12.1, 1H, $H3_b$-NeuAc), 1.54-1.45 (m, 1H, $CH_2$-Ile), 1.32 (d, J = 6.3 Hz, 3H, $CH_3$-Thr), 1.24-1.16 (m, 1H, $CH_2$-Ile), 0.99-0.91 (m, 9H, 2x$CH_3$-Val, $CH_3$-Ile), 0.87 (t, J = 7.4 Hz, 3H, $CH_3$-Ile). $^{13}$C from HSQC (150MHz, $D_2O$): δ = 129.5 (d, 2C), 129.3 (d, 2C), 128.2 (d, 1C), 104.5 (d, 1C), 102.5 (d, 1C), 101.3 (d, 1C), 99.2 (d, 1C), 77.3 (d, 1C), 77.2 (d, 1C), 75.5 (d, 1C), 75.4 (d, 1C), 74.6 (d, 1C), 72.9 (d, 2C), 72.6 (d, 3C), 71.8 (d, 1C), 70.7 (d, 1C), 70.7 (t, 1C), 70.0 (d, 1C), 69.0 (d, 2C), 68.7 (d, 1C), 68.4 (d, 1C), 68.1 (d, 1C), 67.3 (d, 1C), 66.1 (t, 1C), 62.4 (t, 1C), 61.1 (t, 2C), 59.2 (d, 1C), 58.0

(d, 1C), 56.9 (d, 1C), 54.5 (d, 1C), 53.8 (d, 1C), 51.6 (d, 1C), 48.3 (d, 1C), 42.0 (t, 1C), 39.6 (t, 1C), 37.0 (t, 1C), 36.6 (d, 1C), 30.5 (d, 1C), 24.5 (t, 1C), 22.4 (q, 2C), 22.0 (q, 1C), 18.7 (q, 1C), 18.2 (q, 2C), 14.8 (q, 1C), 10.2 (q, 1C). ESI-MS calcd for $C_{65}H_{104}N_9O_{37}S^-$ $[M+H]^-$ 1634.6259, found 1634.6227

* Signal lies under $D_2O$ peak

**Compound 14**

Starting from compound **13** (3.1 mg, 1.9 µmol) following general procedure V compound **14** (2.3 mg, 63%) was obtained as a white solid. $^1H$ (600MHz, $D_2O$): δ = 7.49-7.38 (m, 3H, $CH_{Ar}$-Phe), 7.30-7.24 (m, 2H, $CH_{Ar}$-Phe), 4.81 (d, 1H, H1-GalNAc)*, 4.65 (d, J = 2.0 Hz, 1H, $CH_{\alpha}$-Thr), 4.58 (d, J = 8.4 Hz, 1H, H1-GlcNAc), 4.54 (d, J = 7.8 Hz, 1H, H1-Gal-2), 4.48 (d, J = 7.7 Hz, 1H, H1-Gal-1), 4.42-4.30 (m, 4H, $CH_{\alpha}$-Val, $CH_{\alpha}$-Phe, $H6_{a,b}$-GlcNAc), 4.29-4.16 (m, 5H, H2, H4-H5-GalNAc, $CH_{\alpha}$-Ile, $CH_{\beta}$-Thr), 4.12-3.99 (m, 4H, H3, $H6_a$-GalNAc, H3-Gal-2, H3-Gal-1), 3.97-3.57 (m, 29H, $CH_2$-Gly, H4-$H6_{a,b}$-Gal-1, H4-$H6_{a,b}$-Gal-2, H4-$H9_{a,b}$-NeuAc-1, H4-$H9_{a,b}$-NeuAc-2, H2-H5-GlcNAc, $H6_b$-GalNAc), 3.54 (dd, J = 7.8, 9.8 Hz, 1H, H2-Gal-1), 3.51 (dd, J = 7.8, 9.8 Hz, 1H, H2-Gal-2), 3.27 (dd, J = 6.8, 14.3 Hz, 1H, $CH_2$-Phe), 3.20 (dd, J = 6.8, 14.3 Hz, 1H, $CH_2$-Phe), 2.79-2.72 (m, 2H, $H3_a$-NeuAc-1, $H3_a$-NeuAc-2), 2.06 (s, 3H, NHAc), 2.03 (s, 6H, NHAc), 2.01 (s, 3H, NHAc), 2.07-2.01 (m, 1H, $CH_{\beta}$-Val), 1.89-1.82 (m, 1H, $CH_{\beta}$-Ile), 1.82-1.76 (m, 2H, $H3_b$-NeuAc-1, $H3_b$-NeuAc-2), 1.54-1.45 (m, 1H, $CH_2$-Ile), 1.33 (d, J = 6.4 Hz, 3H, $CH_3$-Thr), 1.24-1.16 (m, 1H, $CH_2$-Ile), 1.02-0.92 (m, 9H, 2x$CH_3$-Val, $CH_3$-Ile), 0.87 (t, J = 7.4 Hz, 3H, $CH_3$-Ile). $^{13}C$ from HSQC (150MHz, $D_2O$): δ = 129.6 (d, 2C), 129.4 (d, 2C), 128.2 (d, 1C), 104.3 (d, 1C), 102.1 (d, 1C), 100.8 (d, 1C), 99.2 (d, 1C), 77.2 (d, 3C), 75.5 (d, 2C), 75.0 (d, 3C), 72.9 (d, 2C), 72.4 (d, 1C), 71.8 (d, 2C), 71.4 (d, 1C), 70.7 (t, 1C), 70.1 (d, 1C), 69.3 (d, 1C), 69.1 (d, 1C), 68.9 (d, 1C), 68.4 (d, 2C), 68.1 (d, 2C), 67.4 (d, 1C), 66.2 (t, 1C), 62.5 (t, 2C), 60.8 (t, 2C), 59.2 (d, 1C), 58.1 (d, 1C), 56.6 (d, 1C), 54.7 (d, 1C), 53.9 (d, 1C), 51.5 (d, 2C), 48.2 (d, 1C), 41.8 (t, 1C), 39.5 (t, 2C), 36.8 (t, 1C), 36.6 (d, 1C), 30.5 (d, 1C), 24.4 (t, 1C), 22.2 (q, 2C), 22.0 (q, 2C), 18.7 (q, 1C), 18.2 (q, 2C), 14.6 (q, 1C), 10.1 (q, 1C). ESI-MS calcd for $C_{76}H_{121}N_{10}O_{45}S^-$ $[M-2H]^{2-}$ 962.3565, found 962.3553.

* Signal lies under $D_2O$-peak

**Compound 15**

Starting from compound **14** (920 µg, 0.5 µmol) following general procedure VI (FUT6) compound **15** (780 µg, 79%) was obtained as a white solid. $^1H$ (600MHz, $D_2O$): δ = 7.49-7.39 (m, 3H, $CH_{Ar}$-Phe), 7.32-7.23 (m, 2H, $CH_{Ar}$-Phe), 5.07 (d, J = 4.1 Hz, 1H), 4.83-4.78 (m, 2H, H1-GalNAc, H5-Fuc)*, 4.68-4.64 (m, 1H, $CH_{\alpha}$-Thr), 4.59 (d, J = 8.0 Hz, 2H, H1-GlcNAc, H1-Gal-2), 4.47 (d, J = 7.8 Hz, 1H, H1-Gal-1), 4.43-4.31 (m, 4H, $CH_{\alpha}$-Val, $CH_{\alpha}$-Phe, $H6_{a,b}$-GlcNAc), 4.29-4.15 (m, 5H, H2, H4-H5-GalNAc, $CH_{\alpha}$-Ile, $CH_{\beta}$-Thr), 4.12- 3.58 (m, 35H, H3, $H6_{a,b}$-GalNAc, H3-$H6_{a,b}$-Gal-1, H3-H4, $H6_{a,b}$-Gal-2, $CH_2$-Gly, H4-$H9_{a,b}$-NeuAc-1, H4-$H9_{a,b}$-NeuAc-2, H2-H5-GlcNAc, H2-H4 Fuc), 3.54-3.46 (m, 3H, H2-Gal-1, H2, H5-Gal-2), 3.27 (dd, J = 6.8, 14.3 Hz, 1H, $CH_2$-Phe), 3.20 (dd, J = 6.8, 14.3 Hz, 1H, $CH_2$-Phe), 2.79-2.72 (m, 2H, $H3_a$-NeuAc-1, $H3_a$-NeuAc-2), 2.05 (s, 3H, NHAc), 2.03 (s, 6H, NHAc), 2.01 (s, 3H, NHAc), 2.07-2.01 (m, 1H, $CH_{\beta}$-Val), 1.89-1.82 (m, 1H, $CH_{\beta}$-Ile), 1.82-1.76 (m, 2H, $H3_b$-NeuAc-1, $H3_b$-NeuAc-2), 1.54-1.45 (m, 1H, $CH_2$-Ile), 1.31 (d, J = 6.3 Hz, 3H, $CH_3$-Thr), 1.24-1.16 (m, 1H, $CH_2$-Ile),

1.16 (d, J = 6.6 Hz, 3H, $CH_3$-Fuc), 1.02-0.92 (m, 9H, 2x$CH_3$-Val, $CH_3$-Ile), 0.87 (t, J = 7.4 Hz, 3H, $CH_3$-Ile). $^{13}C$ from HSQC (150MHz, $D_2O$): δ = 129.5 (d, 2C), 129.3 (d, 2C), 128.2 (d, 1C), 104.6 (d, 1C), 101.1 (d, 2C), 99.3 (d, 1C), 98.6 (d, 1C), 77.4 (d, 2C), 75.6 (d, 1C), 75.4 (d, 1C), 75.1 (d, 1C), 74.7 (d, 1C), 74.6 (d, 1C), 73.0 (d, 1C), 72.9 (d, 1C), 72.7 (d, 1C), 71.8 (d, 1C), 71.3 (d, 1C), 70.7 (t, 1C), 70.0 (d, 1C), 69.1 (d, 1C), 69.0 (d, 2C), 68.4 (d, 2C), 68.1 (d, 2C), 67.7 (d, 2C), 66.7 (d, 1C), 65.8 (t, 1C), 62.5 (t, 2C), 61.1 (t, 1C), 60.8 (t, 1C), 59.4 (d, 1C), 58.0 (d, 1C), 56.9 (d, 1C), 55.4 (d, 1C), 54.0 (d, 1C), 51.6 (d, 2C), 48.2 (d, 1C), 42.0 (t, 1C), 39.7 (t, 2C), 36.6 (t, 1C), 36.6 (d, 1C), 30.4 (d, 1C), 24.4 (t, 1C), 22.4 (q, 1C), 22.2 (q, 1C), 22.1 (q, 1C), 18.7 (q, 1C), 18.1 (q, 2C), 15.2 (q, 1C), 14.7 (q, 1C), 10.1 (q, 1C). ESI-MS calcd for $C_{82}H_{131}N_{10}O_{49}S^-$ $[M-2H]^{2-}$ 1035.3855, found 1035.3839.

**Compound 16**

Starting from compound **14** (920 µg, 0.5 µmol) following general procedure IX compound **16** (710 µg, 74%) was obtained as a white solid. $^1H$ (600MHz, $D_2O$): δ = 7.47-7.42 (m, 3H, $CH_{Ar}$-Phe), 7.32-7.27 (m, 2H, $CH_{Ar}$-Phe), 4.82 (d, 1H, H1-GalNAc)*, 4.66 (d, J = 1.9 Hz, 1H, $CH_α$-Thr), 4.58 (d, J = 8.4 Hz, 1H, H1-GlcNAc), 4.49 (d, J = 7.9 Hz, 1H, H1-Gal-2), 4.43 (d, J = 7.9 Hz, 1H, H1-Gal-1), 4.42-4.36 (m, 2H, $CH_α$-Val, $H6_a$-GlcNAc), 4.33 (t, J = 6.7 Hz, 1H, $CH_α$-Phe), 4.31-4.19 (m, 6H, $CH_β$-Thr, $H6_b$-GlcNAc, H2, H4-H5-GalNAc, $CH_α$-Ile), 4.15- 4.00 (m, 6H, H3, $H6_{a,b}$-Gal-2, H3, $H6_a$-GalNAc, H3-Gal-1), 3.99-3.56 (m, 27H, $CH_2$-Gly, H4-$H6_{a,b}$-Gal-1, H4-H5-Gal-2, H4-$H9_{a,b}$-NeuAc-1, H4-$H9_{a,b}$-NeuAc-2, H2-H5-GlcNAc, $H6_b$-GalNAc), 3.55-3.48 (m, 2H, H2-Gal-1, H2-Gal-2), 3.28 (dd, J = 6.8, 14.3 Hz, 1H, $CH_2$-Phe), 3.21 (dd, J = 6.7, 14.2 Hz, 1H, $CH_2$-Phe), 2.79-2.72 (m, 2H, $H3_a$-NeuAc-1, $H3_a$-NeuAc-2), 2.05 (s, 3H, NHAc), 2.03 (s, 6H, NHAc), 2.01 (s, 3H, NHAc), 2.07-2.01 (m, 1H, $CH_β$-Val), 1.89-1.82 (m, 1H, $CH_β$-Ile), 1.82-1.76 (m, 2H, $H3_b$-NeuAc-1, $H3_b$-NeuAc-2), 1.53-1.45 (m, 1H, $CH_2$-Ile), 1.37 (d, J = 6.3 Hz, 3H, $CH_3$-Thr), 1.24-1.18 (m, 1H, $CH_2$-Ile), 1.01-0.93 (m, 9H, 2x$CH_3$-Val, $CH_3$-Ile), 0.87 (t, J = 7.4 Hz, 3H, $CH_3$-Ile). $^{13}C$ from HSQC (150MHz, $D_2O$): δ = 129.6 (d, 2C), 129.5 (d, 2C), 128.2 (d, 1C), 104.5 (d, 1C), 102.2 (d, 1C), 101.2 (d, 1C), 99.3 (d, 1C), 78.7 (d, 1C), 77.7 (d, 1C), 77.3 (d, 1C), 75.3 (d, 2C), 74.6 (d, 1C), 72.8 (d, 2C), 72.5 (d, 2C), 72.4 (d, 2C), 71.8 (d, 2C), 70.6 (t, 1C), 70.4 (d, 1C), 69.1 (d, 3C), 68.5 (d, 1C), 68.1 (d, 2C), 67.2 (d, 1C), 67.0 (d, 1C), 66.9 (t, 1C), 66.6 (t, 1C), 62.6 (t, 2C), 60.8 (t, 1C), 59.2 (d, 1C), 57.9 (d, 1C), 57.0 (d, 1C), 54.9 (d, 1C), 54.0 (d, 1C), 51.5 (d, 2C), 48.4 (d, 1C), 41.9 (t, 1C), 39.6 (t, 2C), 36.8 (t, 1C), 36.6 (d, 1C), 30.4 (d, 1C), 24.4 (t, 1C), 22.2 (q, 2C), 22.0 (q, 2C), 18.9 (q, 1C), 18.3 (q, 2C), 14.7 (q, 1C), 10.1 (q, 1C). ESI-MS calcd for $C_{76}H_{121}N_{10}O_{48}S_2^-$ $[M-2H]^{2-}$ 1002.3349, found 1002.3331.

**Compound 17**

Starting from compound **13** (500 µg, 0.3 µmol) following general procedure VI (FUT6) compound **17** (400 µg, 73%) was obtained as a white solid. $^1H$ (600MHz, $D_2O$): δ = 7.47-7.35 (m, 3H, $CH_{Ar}$-Phe), 7.31-7.21 (m, 2H, $CH_{Ar}$-Phe), 5.09 (d, J = 4.0 Hz, 1H, H1-Fuc), 4.85-4.76 (m, 2H, H5-Fuc, H1-GalNAc)*, 4.63 (d, J = 2.0 Hz, 1H, $CH_α$-Thr), 4.59 (d, J = 8.3 Hz, 1H, H1-GlcNAc), 4.54 (d, J = 7.8 Hz, 1H, H1-Gal-2), 4.48 (d, J = 7.8 Hz, 1H, H1-Gal-1), 4.41-4.32 (m, 3H, $CH_α$-Val, $H6_{a,b}$-GlcNAc), 4.26-4.15 (m, 5H, H2, H4-H5-GalNAc, $CH_α$-Ile, $CH_β$-Thr), 4.12-4.07 (m, 1H, $H6_a$-GalNAc), 4.05 (dd, J = 3.1, 9.9 Hz, 1H, H3-Gal-1), 4.02 (dd, J = 2.9, 11.0 Hz, 1H, H3-GalNAc), 3.99-3.54 (m, 27H, $CH_α$-Phe, H4-$H6_{a,b}$-Gal-1, H3-$H6_{a,b}$-Gal-2, H2-H5-GlcNAc, $CH_2$-Gly, H4-$H9_{a,b}$-

NeuAc, H6$_b$-GalNAc, H2-H4-Fuc), 3.54-3.45 (m, 2H, H2-Gal-1, H2-Gal-2), 3.23-3.05 (m, 2H, $CH_2$-Phe), 2.76 (dd, J = 4.6, 12.4, 1H, H3$_a$-NeuAc), 2.05 (s, 3H, NHAc), 2.03 (s, 3H, NHAc), 2.01 (s, 3H, NHAc), 2.07-2.01 (m, 1H, $CH_\beta$-Val), 1.90-1.82 (m, 1H, $CH_\beta$-Ile), 1.78 (t, J = 12.1, 1H, H3$_b$-NeuAc), 1.54-1.45 (m, 1H, $CH_2$-Ile), 1.29 (d, J = 6.4 Hz, 3H, $CH_3$-Thr), 1.24-1.16 (m, 1H, $CH_2$-Ile), 1.17 (d, J = 6.6 Hz, 3H, $CH_3$-Fuc), 0.99-0.91 (m, 9H, 2x$CH_3$-Val, $CH_3$-Ile), 0.87 (t, J = 7.4 Hz, 3H, $CH_3$-Ile). $^{13}$C from HSQC (150MHz, $D_2O$): δ = 129.5 (d, 2C), 129.2 (d, 2C), 127.9 (d, 1C), 104.6 (d, 1C), 101.5 (d, 1C), 101.1 (d, 1C), 99.2 (d, 1C), 98.6 (d, 1C), 77.2 (d, 2C), 75.4 (d, 1C), 75.1 (d, 1C), 75.0 (d, 1C), 74.7 (d, 1C), 73.1 (d, 1C), 72.9 (d, 1C), 72.8 (d, 1C), 72.5 (d, 1C), 71.9 (d, 1C), 71.7 (d, 1C), 70.8 (d, 1C), 70.7 (t, 1C), 70.0 (d, 1C), 69.3 (d, 1C), 69.0 (d, 1C), 68.8 (d, 1C), 68.5 (d, 1C), 68.4 (d, 1C), 67.9 (d, 1C), 67.8 (d, 1C), 67.3 (d, 1C), 66.6 (d, 1C), 65.9 (t, 1C), 62.3 (t, 1C), 61.0 (t, 2C), 59.2 (d, 1C), 57.9 (d, 1C), 56.7 (d, 1C), 55.4 (d, 2C), 51.6 (d, 1C), 48.4 (d, 1C), 41.9 (t, 1C), 39.7 (t, 1C), 36.5 (d, 1C), 30.4 (d, 1C), 24.5 (t, 1C), 22.3 (q, 2C), 22.0 (q, 1C), 18.7 (q, 1C), 18.2 (q, 2C), 15.2 (q, 1C), 14.7 (q, 1C), 10.1 (q, 1C). ESI-MS calcd for $C_{71}H_{114}N_9O_{41}S^-$ [M-2H]$^{2-}$ 889.8377, found 889.8361.

* Signal lies under $D_2O$ peak ** signal of $CH_2$-Phe is not visible in HSQC

**Compound 18**

Starting from compound **3** (3.5 mg, 3.2 µmol) following general procedure VII compound **18** (2.8 mg, 75%) was obtained as a white solid. $^1$H (600MHz, $D_2O$): δ = 7.45-7.36 (m, 3H, $CH_{Ar}$-Phe), 7.29-7.22 (m, 2H, $CH_{Ar}$-Phe), 4.82 (d, J = 4.0 Hz, 1H, H1-GalNAc), 4.62 (d, J = 2.2 Hz, 1H, $CH_\alpha$-Thr), 4.57 (d, J = 8.5 Hz, 1H, H1-GlcNAc), 4.39 (d, J = 7.8 Hz, 1H, H1-Gal), 4.36-4.33 (m, 2H, $CH_\alpha$-Val, H6$_a$-GlcNAc), 4.28-4.18 (m, 7H, $CH_\alpha$-Phe, $CH_\alpha$-Thr, $CH_\alpha$-Ile, H6$_b$-GlcNAc, H2, H4-H5-GalNAc), 4.07 (dd, J = 2.5, 11.1 Hz, 1H, H6$_a$-GalNAc), 4.00 (dd, J = 3.1, 11.0 Hz, 1H, H3-GalNAc), 3.96-3.88 (m, 2H, $CH_2$-Gly, H4-Gal), 3.83-3.70 (m, 5H, $CH_2$-Gly, H6$_{a,b}$-Gal, H2-GlcNAc, H6$_b$-GalNAc), 3.68-3.64 (m, 1H, H5-GlcNAc), 3.63 (dd, J = 4.5, 7.8 Hz, 1H, H5-Gal), 3.57 (dd, J = 3.3, 9.9 Hz, 1H, H3-Gal), 3.55-3.46 (m, 3H, H3-H4-GlcNAc, H2-Gal), 3.22 (dd, J = 6.8, 14.2 Hz, 1H, $CH_2$-Phe), 3.15 (dd, J = 6.8, 14.2 Hz, 1H, $CH_2$-Phe), 2.06 (s, 3H, NHAc), 2.02 (s, 3H, NHAc), 2.08-2.00 (m, 1H, $CH_\beta$-Val), 1.92-1.82 (m, 1H, $CH_\beta$-Ile), 1.54-1.45 (m, 1H, $CH_2$-Ile), 1.32 (d, J = 6.4 Hz, 3H, $CH_3$-Thr), 1.24-1.16 (m, 1H, $CH_2$-Ile), 0.99-0.91 (m, 9H, 2x$CH_3$-Val, $CH_3$-Ile), 0.87 (t, J = 7.4 Hz, 3H, $CH_3$-Ile). $^{13}$C from HSQC (150MHz, $D_2O$): δ = 129.5 (d, 2C), 129.2 (d, 2C), 127.9 (d, 1C), 104.5 (d, 1C), 101.4 (d, 1C), 99.3 (d, 1C), 77.2 (d, 1C), 77.1 (d, 1C), 75.0 (d, 1C), 74.2 (d, 1C), 73.6 (d, 1C), 72.7 (d, 1C), 70.5 (d, 1C), 70.5 (t, 1C), 70.0 (d, 1C), 69.6 (d, 1C), 69.2 (d, 1C), 68.5 (d, 1C), 67.0 (t, 1C), 60.9 (t, 1C), 59.1 (d, 1C), 58.0 (d, 1C), 57.0 (d, 1C), 55.3 (d, 1C), 54.1 (d, 1C), 48.2 (d, 1C), 41.9 (t, 1C), 37.4 (t, 1C), 36.4 (d, 1C), 30.5 (d, 1C), 24.4 (t, 1C), 22.3 (q, 1C), 22.2 (q, 1C), 18.7 (q, 1C), 18.2 (q, 2C), 14.7 (q, 1C), 10.1 (q, 1C). ESI-MS calcd for $C_{48}H_{77}N_8O_{24}S^-$ [M-H]$^-$ 1181.4771, found 1181.4778.

**Compound 19**

Starting from compound **18** (2.4 mg, 2.0 µmol) following general procedure VIII compound **19** (2.3 mg, 85%) was obtained as a white solid. $^1$H (600MHz, $D_2O$): δ = 7.47-7.39 (m, 3H, $CH_{Ar}$-Phe), 7.31-7.24 (m, 2H, $CH_{Ar}$-Phe), 4.83 (d, 1H, H1-GalNAc)*, 4.64 (d, J = 2.2 Hz, 1H, $CH_\alpha$-Thr), 4.59 (d, J = 8.4 Hz, 1H, H1-GlcNAc), 4.48 (d, J = 7.8 Hz, 1H, H1-Gal-2), 4.42-4.30 (m, 5H, H1-Gal-1, H6$_{a,b}$-GlcNAc, $CH_\alpha$-Val, $CH_\alpha$-Phe),

4.28-4.18 (m, 5H, $CH_{\alpha}$-Thr, $CH_{\alpha}$-Ile, H2, H4-H5-GalNAc), 4.08 (dd, J = 2.4, 11.3 Hz, 1H, $H6_a$-GalNAc), 4.00 (dd, J = 3.0, 11.0 Hz, 1H, H3-GalNAc), 3.97-3.88 (m, 3H, $CH_2$-Gly, H4-Gal-1, H4-Gal-2), 3.85-3.60 (m, 13H, $CH_2$-Gly, H5-$H6_{a,b}$-Gal-1, H3, H5-$H6_{a,b}$-Gal-2, H2-H5-GlcNAc, $H6_b$-GalNAc), 3.57 (dd, J = 3.4, 9.9 Hz, 1H, H3-Gal-1), 3.54-3.46 (m, 2H, H2-Gal-1, H2-Gal-2), 3.27 (dd, J = 6.6, 14.3 Hz, 1H, $CH_2$-Phe), 3.20 (dd, J = 6.9, 14.3 Hz, 1H, $CH_2$-Phe), 2.06 (s, 3H, NHAc), 2.01 (s, 3H, NHAc), 2.08-2.00 (m, 1H, $CH_{\beta}$-Val), 1.92-1.82 (m, 1H, $CH_{\beta}$-Ile), 1.54-1.45 (m, 1H, $CH_2$-Ile), 1.32 (d, J = 6.3 Hz, 3H, $CH_3$-Thr), 1.26-1.16 (m, 1H, $CH_2$-Ile), 1.02-0.92 (m, 9H, 2x$CH_3$-Val, $CH_3$-Ile), 0.87 (t, J = 7.4 Hz, 3H, $CH_3$-Ile). $^{13}C$ from HSQC (150MHz, $D_2O$): δ = 129.6 (d, 2C), 129.3 (d, 2C), 128.1 (d, 1C), 104.7 (d, 1C), 102.5 (d, 1C), 101.2 (d, 1C), 99.3 (d, 1C), 77.4 (d, 1C), 77.3 (d, 2C), 75.5 (d, 1C), 74.8 (d, 1C), 72.6 (d, 1C), 72.5 (d, 3C), 70.9 (d, 1C), 70.5 (d, 1C), 70.5 (t, 1C), 70.1 (d, 1C), 69.2 (d, 1C), 68.6 (d, 2C), 66.2 (t, 1C), 61.0 (t, 2C), 60.0 (d, 1C), 59.2 (d, 1C), 57.1 (d, 1C), 54.9 (d, 1C), 54.0 (d, 1C), 48.3 (d, 1C), 42.0 (t, 1C), 36.8 (t, 1C), 36.5 (d, 1C), 30.5 (d, 1C), 24.4 (t, 1C), 22.3 (q, 1C), 22.2 (q, 1C), 18.6 (q, 1C), 18.1 (q, 2C), 14.7 (q, 1C), 10.1 (q, 1C). ESI-MS calcd for $C_{54}H_{87}N_8O_{29}S^-$ $[M-H]^-$ 1343.5300, found 1343.5281.

* Signal under $D_2O$ peak

**Compound 20**

Starting from compound **19** (2.0 mg, 1.5 µmol) following general procedure V compound **20** (1.9 mg, 76%) was obtained as a white solid. $^1H$ (600MHz, $D_2O$): δ = 7.47-7.39 (m, 3H, $CH_{Ar}$-Phe), 7.31-7.24 (m, 2H, $CH_{Ar}$-Phe), 4.82 (d, 1H, H1-GalNAc)*, 4.64 (d, J = 2.1 Hz, 1H, $CH_{\alpha}$-Thr), 4.59 (d, J = 8.5 Hz, 1H, H1-GlcNAc), 4.51 (d, J = 7.8 Hz, 1H, H1-Gal-2), 4.43-4.30 (m, 5H, H1-Gal-1, $H6_{a,b}$-GlcNAc, $CH_{\alpha}$-Val, $CH_{\alpha}$-Phe), 4.29-4.19 (m, 5H, $CH_{\beta}$-Thr, $CH_{\alpha}$-Ile, H2, H4-H5-GalNAc), 4.11-4.03 (m, 2H, $H6_a$-GalNAc, H3-Gal-2), 4.01 (dd, J = 3.0, 11.0 Hz, 1H, H3-GalNAc), 3.97-3.84 (m, 6H, $CH_2$-Gly, H4-Gal-1, H4-Gal-2, H5, H8-$H9_a$-NeuAc), 3.83-3.51 (m, 18H, $CH_2$-Gly, H3, H5-$H6_{a,b}$-Gal-1, H2, H5-$H6_{a,b}$-Gal-2, H2-H5-GlcNAc, $H6_b$-GalNAc, H4, H6-H7, $H9_b$-NeuAc), 3.49 (dd, J = 7.7, 9.9 Hz, 1H, H2-Gal-1), 3.28 (dd, J = 6.7, 14.3 Hz, 1H, $CH_2$-Phe), 3.21 (dd, J = 6.7, 14.3 Hz, 1H, $CH_2$-Phe), 2.75 (dd, J = 4.7, 12.4 Hz, 1H, $H3_a$-NeuAc), 2.05 (s, 3H, NHAc), 2.04 (s, 3H, NHAc), 2.01 (s, 3H, NHAc), 2.08-2.00 (m, 1H, $CH_{\beta}$-Val), 1.92-1.82 (m, 1H, $CH_{\beta}$-Ile), 1.80 (t, J = 12.1 Hz, 1H, $H3_b$-NeuAc), 1.54-1.45 (m, 1H, $CH_2$-Ile), 1.35 (d, J = 6.3 Hz, 3H, $CH_3$-Thr), 1.26-1.16 (m, 1H, $CH_2$-Ile), 1.02-0.92 (m, 9H, 2x$CH_3$-Val, $CH_3$-Ile), 0.87 (t, J = 7.4 Hz, 3H, $CH_3$-Ile). $^{13}C$ from HSQC (150MHz, $D_2O$): δ = 129.6 (d, 2C), 129.4 (d, 2C), 128.2 (d, 1C), 104.6 (d, 1C), 102.2 (d, 1C), 101.3 (d, 1C), 99.3 (d, 1C), 77.3 (d, 3C), 75.3 (d, 1C), 75.0 (d, 2C), 72.7 (d, 2C), 72.6 (d, 1C), 72.4(d, 1C), 71.5 (d, 1C), 70.5 (d, 1C), 70.5 (t, 1C), 70.2 (d, 1C), 69.3 (d, 2C), 68.5 (d, 2C), 67.9 (d, 1C), 67.4 (d, 1C), 66.3 (t, 1C), 62.5 (t, 1C), 60.8 (t, 2C), 59.3 (d, 1C), 58.0 (d, 1C), 57.0 (d, 1C), 54.8 (d, 1C), 53.9 (d, 1C), 51.7 (d, 1C), 48.4 (d, 1C), 41.8 (t, 1C), 39.5 (t, 1C), 36.7 (t, 1C), 36.4 (d, 1C), 30.5 (d, 1C), 24.4 (t, 1C), 22.3 (q, 1C), 22.2 (q, 1C), 22.0 (q, 1C), 18.6 (q, 1C), 18.1 (q, 2C), 14.8 (q, 1C), 10.1 (q, 1C). ESI-MS calcd for $C_{65}H_{104}N_9O_{37}S^-$ $[M-H]^-$ 1634.6254, found 1634.6233.

**Compound 21**

Starting from compound **20** (810 µg, 0.5 µmol) following general procedure IX compound **21** (670 µg, 79%) was obtained as a white solid. $^1H$ (600MHz, $D_2O$): δ = 7.47-7.39 (m, 3H, $CH_{Ar}$-Phe), 7.33-7.25 (m, 2H, $CH_{Ar}$-Phe), 4.83 (d, 1H, H1-GalNAc)*,

4.64-4.62 (m, 1H, $CH_{\alpha}$-Thr), 4.59 (d, J = 8.3 Hz, 1H, H1-GlcNAc), 4.45-4.35 (m, 4H, H1-Gal-1, H1-Gal-2, $H6_a$-GlcNAc, $CH_{\alpha}$-Val), 4.32-4.18 (m, 7H, $CH_{\alpha}$-Phe, $CH_{\beta}$-Thr, $CH_{\alpha}$-Ile, H2, H4-H5-GalNAc, $H6_b$-GlcNAc), 4.12 (dd, J = 4.6, 10.6 Hz, 1H, H6a-Gal-2), 4.09-3.75 (m, 15H, $CH_2$-Gly, H3-$H6_b$-Gal-2, H3, $H6_{a,b}$-GalNAc, H2-GlcNAc, H5, H8-$H9_a$-NeuAc, H4, $H6_a$-Gal-1), 3.75-3.55 (m, 10H, H3, H5-$H6_b$-Gal-1, H3-H5 GlcNAc, H4, H6-H7, $H9_b$-NeuAc), 3.53 (dd, J = 7.8, 9.9 Hz, 1H, H2-Gal-2), 3.49 (dd, J = 7.8, 9.9 Hz, 1H, H2-Gal-1), 3.30-3.14 (m, 2H, $CH_2$-Phe), 2.75 (dd, J = 4.7, 12.5 Hz, 1H, $H3_a$-NeuAc), 2.04 (s, 3H, NHAc), 2.03 (s, 3H, NHAc), 2.01 (s, 3H, NHAc), 2.08-2.00 (m, 1H, $CH_{\beta}$-Val), 1.92-1.84 (m, 1H, $CH_{\beta}$-Ile), 1.80 (t, J = 12.4 Hz, 1H, $H3_b$-NeuAc), 1.54-1.45 (m, 1H, $CH_2$-Ile), 1.38 (d, J = 6.3 Hz, 3H, $CH_3$-Thr), 1.26-1.16 (m, 1H, $CH_2$-Ile), 1.02-0.92 (m, 9H, 2x$CH_3$-Val, $CH_3$-Ile), 0.87 (t, J = 7.4 Hz, 3H, $CH_3$-Ile). $^{13}C$ from HSQC (150MHz, $D_2O$): δ = 129.5 (d, 2C), 129.3 (d, 2C), 128.2 (d, 1C), 104.4 (d, 1C), 102.4 (d, 1C), 101.3 (d, 1C), 99.0 (d, 1C), 78.8 (d, 1C), 77.6 (d, 1C), 77.0 (d, 1C), 75.0 (d, 1C), 74.6 (d, 1C), 72.9 (d, 1C), 72.5 (d, 1C), 72.4(d, 3C), 71.5 (d, 1C), 70.5 (d, 2C), 70.5 (t, 1C), 69.2 (d, 1C), 69.0 (d, 1C), 68.6 (d, 1C), 68.5 (d, 1C), 68.2 (d, 1C), 67.1 (d, 1C), 66.9 (t, 1C), 66.5 (t, 1C), 62.5 (t, 1C), 60.8 (t, 1C), 59.1 (d, 1C), 58.0 (d, 1C), 57.0 (d, 1C), 55.1 (d, 1C), 54.5 (d, 1C), 51.6 (d, 1C), 48.4 (d, 1C), 42.0 (t, 1C), 39.5 (t, 1C), 36.5 (d, 1C), 30.4 (d, 1C), 24.3 (t, 1C), 22.3 (q, 1C), 22.2 (q, 2C), 18.7 (q, 1C), 17.9 (q, 2C), 14.6 (q, 1C), 9.9 (q, 1C).** ESI-MS calcd for $C_{65}H_{104}N_9O_{40}S_2^-$ $[M-2H]^{2-}$ 856.7872, found 856.7867.

* Signal lies under $D_2O$ peak, **signal of $CH_2$-Phe is not visible in HSQC

**Compound 22**

Starting from compound **20** (750 µg, 0.5 µmol) following general procedure VI (FUT6) compound **22** (650 µg, 80%) was obtained as a white solid. $^1H$ (600MHz, $D_2O$): δ = 7.47-7.39 (m, 3H, $CH_{Ar}$-Phe), 7.31-7.24 (m, 2H, $CH_{Ar}$-Phe), 5.08 (d, 1H, H1-Fuc), 4.82-4.75 (m, 2H, H1-GalNAc, H5-Fuc)*, 4.63 (d, J = 2.1 Hz, 1H, $CH_{\alpha}$-Thr), 4.60 (d, J = 8.3 Hz, 1H, H1-GlcNAc), 4.56 (d, J = 7.8 Hz, 1H, H1-Gal-2), 4.42-4.33 (m, 4H, H1-Gal-1, $H6_{a,b}$-GlcNAc, $CH_{\alpha}$-Val), 4.32-4.18 (m, 6H, $CH_{\alpha}$-Phe, $CH_{\beta}$-Thr, $CH_{\alpha}$-Ile, H2, H4-H5-GalNAc), 4.11-4.03 (m, 2H, $H6_a$-GalNAc, H3-Gal-2), 4.01 (dd, J = 3.0, 11.0 Hz, 1H, H3-GalNAc), 3.98-3.59 (m, 24H, $CH_2$-Gly, H2-H4-Fuc, H4, H5-$H6_{a,b}$-Gal-1, H4, $H6_{a,b}$-Gal-2, H2-H5-GlcNAc, H4-$H9_{a,b}$-NeuAc, $H6_b$-GalNAc), 3.57 (dd, J = 3.4, 10.0 Hz, 1H, H3-Gal-1), 3.53-3.46 (m, 3H, H2-Gal-1, H2-Gal-2, H5-Gal-2), 3.29- 3.13 (m, 2H, $CH_2$-Phe), 2.75 (dd, J = 4.7, 12.4 Hz, 1H, $H3_a$-NeuAc), 2.04 (s, 3H, NHAc), 2.03 (s, 3H, NHAc), 2.01 (s, 3H, NHAc), 2.08-2.00 (m, 1H, $CH_{\beta}$-Val), 1.92-1.82 (m, 1H, $CH_{\beta}$-Ile), 1.80 (t, J = 12.1 Hz, 1H, $H3_b$-NeuAc), 1.54-1.45 (m, 1H, $CH_2$-Ile), 1.33 (d, J = 6.3 Hz, 3H, $CH_3$-Thr), 1.26-1.16 (m, 1H, $CH_2$-Ile), ), 1.16 (d, J = 6.6 Hz, 3H, $CH_3$-Fuc), 1.02-0.92 (m, 9H, 2x$CH_3$-Val, $CH_3$-Ile), 0.87 (t, J = 7.4 Hz, 3H, $CH_3$-Ile). $^{13}C$ from HSQC (150MHz, $D_2O$): δ = 129.5 (d, 2C), 129.3 (d, 2C), 128.0 (d, 1C), 104.7 (d, 1C), 101.3 (d, 1C), 101.0 (d, 1C), 99.2 (d, 1C), 98.5 (d, 1C), 77.3 (d, 1C), 77.1 (d, 1C), 75.4 (d, 1C), 75.0 (d, 1C), 74.8 (d, 1C), 74.5 (d, 1C), 73.1 (d, 1C), 73.0 (d, 1C), 72.9 (d, 1C), 72.4 (d, 1C) 71.9(d, 1C), 71.4 (d, 1C), 70.4 (d, 1C), 70.6 (t, 1C), 70.6 (d, 1C), 69.3 (d, 1C), 69.2 (d, 2C), 68.4 (d, 2C), 68.0 (d, 1C), 67.8 (d, 1C), 67.2 (d, 1C), 66.7 (d, 1C), 65.9 (t, 1C), 62.4 (t, 1C), 60.9 (t, 2C), 59.1 (d, 1C), 58.1 (d, 1C), 56.9 (d, 1C), 55.5 (d, 1C), 51.6 (d, 1C), 48.3 (d, 1C), 41.9 (t, 1C), 39.7 (t, 1C), 36.6 (d, 1C), 30.4 (d, 1C), 24.4 (t, 1C), 22.2 (q, 2C), 22.0 (q, 1C), 18.8 (q, 1C), 18.2 (q, 2C), 15.2 (q, 1C), 14.8 (q,

1C), 10.0 (q, 1C). ** ESI-MS calcd for $C_{71}H_{114}N_9O_{41}S^-$ $[M-2H]^{2-}$ 889.8377, found 889.8365.

* Signals lie under $D_2O$ peak **signal of $CH_2$-Phe and $CH_\alpha$-Phe is not visible in HSQC

**Compound 23 (Core 3)**

Starting from **S11** and following the deacetylation protocol of compound **S2** O-glycan core 3 (**23**) was obtained. $^1$H (600MHz, $D_2O$): δ = 7.42-7.34 (m, 3H, $CH_{Ar}$-Phe), 7.28-7.24 (m, 2H, $CH_{Ar}$-Phe), 4.78 (d, J = 171.9 Hz, 1H, H1-GalNAc)*, 4.61 (d, J = 2.2 Hz, 1H, $CH_\alpha$-Thr), 4.59 (d, J = 8.4 Hz, 1H, H1-GlcNAc), 4.36 (d, J = 7.9 Hz, 1H, $CH_\alpha$-Val), 4.34 (t, J = 6.8 Hz, 1H, $CH_\alpha$-Phe), 4.33-4.28 (m, 1H, $CH_\beta$-Thr), 4.22 (d, J = 7.9 Hz, 1H, $CH_\alpha$-Ile), 4.21-4.17 (m, 2H, H2, H4-GalNAc), 4.05 (dd, J = 4.5, 7.8 Hz, 1H, H5-GalNAc), 3.98 (dd, J = 3.0, 11.1 Hz, 1H, H3-GalNAc), 3.94 (d, J = 16.9 Hz, 1H, $CH_2$-Gly), 3.90 (dd, J = 1.6, 12.3 Hz, 1H, $H6_a$-GlcNAc), 3.83 (d, J = 16.8 Hz, 1H, $CH_2$-Gly), 3.80- 3.72 (m, 3H, $H6_b$-GlcNAc, $H6_{a,b}$-GalNAc), 3.67 (dd, J = 8.6, 10.2 Hz, 1H, H2-GlcNAc), 3.58-3.53 (m, 1H, H3-GlcNAc), 3.46 (t, J = 9.2 Hz, 1H), 3.43-3.38 (m, 1H, H5-GlcNAc), 3.24 (dd, J = 6.7, 14.3 Hz, 1H, $CH_2$-Phe), 3.18 (dd, J = 6.7, 14.3 Hz, 1H, $CH_2$-Phe), 2.09-2.03 (m, 1H, $CH_\beta$-Val), 2.05 (s, 3H, NHAc), 2.03 (s, 3H, NHAc), 1.90-1.82 (m, 1H, $CH_\beta$-Ile), 1.55- 1.46 (m, 1H, $CH_2$-Ile), 1.32 (d, J = 6.3 Hz, 3H, $CH_3$-Thr), 1.25- 1.15 (m, 1H, $CH_2$-Ile), 1.00-0.93 (m, 9H, 2x$CH_3$-Val, $CH_3$ Ile), 0.87 (t, J = 7.4 Hz, 3H, $CH_3$ Ile). $^{13}$C from HSQC (150MHz, $D_2O$): δ = 129.4 (d, 2C), 129.0 (d, 3C), 102.3 (d, 1C), 98.9 (d, 1C), 76.2 (d, 1C), 76.02 (d, 1C), 75.5 (d, 1C), 73.3 (d, 1C), 70.9 (d, 1C), 69.8 (d, 1C), 68.9 (d, 1C), 60.9 (d, 1C), 60.2 (t, 1C), 59.0 (d, 1C), 58.0 (d, 1C), 56.9 (d, 1C), 55.3 (d, 1C), 54.0 (d, 1C), 48.0 (d, 1C), 41.8 (t, 1C), 36.8 (t, 1C), 36.5 (d, 1C), 30.5 (d, 1C), 24.3 (t, 1C), 22.3 (q, 2C), 18.1 (q, 3C), 14.7 (q, 3C), 9.9 (q, 3C). ESI-MS calcd for $C_{42}H_{69}N_8O_{16}^+$ $[M+H]^+$ 941.4826, found 941.4823

* Signal is under $D_2O$-peak. The J-coupling was determined by a decoupled HSQC.

**Compound 24**

Starting from compound **23** (6.0 mg, 6.4 µmol) following general procedure III compound **24** (6.2 mg, 88%) was obtained as a white solid. $^1$H (600MHz, $D_2O$): δ = 7.41-7.36 (m, 3H, $CH_{Ar}$-Phe), 7.27-7.21 (m, 2H, $CH_{Ar}$-Phe), 4,78 (d, J = 173.8 Hz, 1H, H1-GalNAc)*, 4.62-4.58 (m, 2H, $CH_\alpha$-Thr, H1-GlcNAc), 4.48 (d, J = 7.9 Hz, 1H, H1-Gal), 4.32-4.26 (m, 2H, $CH_\alpha$-Val, $CH_\beta$-Thr), 4.24-4.16 (m, 3H, $CH_\alpha$-Ile, H2, H4-GalNAc), 4.12-4.07 (m, 1H, $CH_\alpha$-Phe), 4.06-4.02 (m, 1H, H5-GalNAc), 4.00-3.90 (m, 4H, H3-GalNAc, $H6_a$-GlcNAc, H4-Gal, $CH_2$-Gly), 3.88-3.81 (m, 2H, $H6_b$-GlcNAc, $CH_2$-Gly), 3.80-3.70 (m, 8H, $H6_{a,b}$-GalNAc, H5-$H6_{a,b}$-Gal, H2-H4-GlcNAc), 3.69-3.66 (m, 1H, H3-Gal), 3.58-3.51 (m, 2H, H5-GlcNAc, H2-Gal), 3.12 (dd, J = 6.6, 13.9 Hz, 1H, $CH_2$-Phe), 3.07 (dd, J = 6.6, 13.9 Hz, 1H, $CH_2$-Phe), 2.04 (s, 3H, NHAc), 2.03 (s, 3H, NHAc), 2.05-1.98 (m, 1H, $CH_\beta$-Val), 1.90-1.82 (m, 1H, $CH_\beta$-Ile), 1.54-1.44 (m, 1H, $CH_2$-Ile), 1.30 (d, J = 6.3 Hz, 3H, $CH_3$-Thr), 1.25-1.15 (m, 1H, $CH_2$-Ile), 0.99-0.92 (m, 9H, 2x$CH_3$-Val, $CH_3$-Ile), 0.87 (t, J = 7.3 Hz, 3H, $CH_3$-Ile). $^{13}$C from HSQC (150MHz, $D_2O$): δ = 129.7 (d, 2C), 129.0 (d, 2C), 127.5 (d, 1C), 102.4 (d, 1C), 102.1 (d, 1C), 98.8 (d, 1C), 78.1 (d, 1C), 76.0 (d, 2C), 74.9 (d, 1C), 74.6 (d, 1C), 72.5 (d, 1C), 71.9 (d, 1C), 70.8 (d, 1C) 70.6 (d, 1C), 69.1 (d, 1C), 68.3 (d, 1C), 61.0 (t, 2C), 59.6 (t, 1C), 59.2 (d, 1C), 58.0 (d, 1C), 56.6 (d, 1C), 54.7 (d, 1C), 54.4 (d, 1C), 47.8 (d, 1C), 41.5 (t, 1C), 38.5 (t, 1C), 36.3 (d, 1C), 30.7 (d, 1C), 24.5 (t, 1C), 22.6 (q, 2C), 18.4 (q, 1C), 18.0 (q, 2C), 14.2 (q,

1C), 10.0 (q, 1C). ESI-MS calcd for $C_{48}H_{78}N_8NaO_{21}^+$ $[M+Na]^+$ 1125.5174, found 1125.5173.

* Signal is under $D_2O$-peak. The J-coupling was determined by a decoupled HSQC.

**Compound 25**

Starting from compound **24** (1.5 mg, 1.4 µmol) following general procedure V compound **25** (1.87 mg, 99%) was obtained as a white solid. $^1$H (600MHz, $D_2O$): δ = 7.42-7.35 (m, 3H, $CH_{Ar}$-Phe), 7.29-7.24 (m, 2H, $CH_{Ar}$-Phe), 4,77 (d, J = 175 Hz, 1H, H1-GalNAc), 4.62 (d, J = 2.0 Hz, 1H, $CH_\alpha$-Thr), 4.58 (d, J = 7.8 Hz, 1H, H1-GlcNAc), 4.56 (d, J = 7.9 Hz, 1H, H1-Gal), 4.39-4.34 (m, 2H, $CH_\alpha$-Val, $CH_\alpha$-Phe), 4.33- 4.27 (m, 1H, $CH_\beta$-Thr), 4.22 (d, J = 7.9 Hz, 1H, $CH_\alpha$-Ile), 4.20-4.17 (m, 2H, H2, H4-GalNAc), 4.12 (dd, J = 3.1, 9.9 Hz, 1H, H3-Gal), 4.05 (dd, J = 5.2, 7.2 Hz, 1H, H5-GalNAc), 4.01-3.92 (m, 4H, H3-GalNAc, $H6_a$-GlcNAc, H4-Gal, $CH_2$-Gly), 3.91-3.81 (m, 5H, H5, H8-$H9_a$-$Neu_5Ac$, $H6_b$-GlcNAc, $CH_2$-Gly), 3.80-3.52 (m, 14H, $H6_{a,b}$-GalNAc, $H6_{a,b}$-Gal, H2-H5-GlcNAc, H2, H5-Gal, H4, H6-H7, $H9_b$-NeuAc), 3.26 (dd, J = 6.6, 14.3 Hz, 1H, $CH_2$-Phe), 3.18 (dd, J = 6.6, 14.3 Hz, 1H, $CH_2$-Phe), 2.76 (dd, J= 4.6, 12.4 Hz, 1H, $H3_a$ NeuAc), 2.10-2.03 (m, 1H, $CH_\beta$-Val), 2.05 (s, 3H, NHAc), 2.04 (s, 3H, NHAc), 2.02 (s, 3H, NHAc), 1.90-1.82 (m, 1H, $CH_\beta$-Ile), 1.80 (t, J = 12.1, 1H, $H3_b$-NeuAc), 1.54-1.45 (m, 1H, $CH_2$-Ile), 1.30 (d, J = 6.3 Hz, 3H, $CH_3$-Thr), 1.25-1.15 (m, 1H, $CH_2$-Ile), 1.00-0.92 (m, 9H, 2x$CH_3$-Val, $CH_3$-Ile), 0.87 (t, J = 7.4 Hz, 3H, $CH_3$-Ile). $^{13}$C from HSQC (150MHz, $D_2O$): δ =129.3 (d, 2C), 128.8 (d, 3H), 102.4 (d, 1C), 102.3 (d, 1C), 98.7 (d, 1C), 78.1 (d, 1C), 76.0 (d, 1C), 75.8 (d, 1C), 75.3 (d, 1C), 75.0 (d, 1C), 74.8 (d, 1C), 72.7 (d, 1C), 72.2 (d, 1C), 71.5 (d, 1C), 70.7 (d, 1C), 69.4 (d, 1C), 68.7 (d, 1C), 68.4 (d, 1C), 67.9 (d, 1C), 67.2 (d, 1C), 62.3 (t, 1C), 60.6 (t, 2C), 59.7 (t, 1C), 59.01 (d, 1C), 57.5 (d, 1C), 57.0 (d, 1C), 55.0 (d, 1C), 53.9 (d, 1C), 51.4 (d, 1C), 48.1 (d, 1C), 41.8 (t, 1C), 39.7 (t, 1C), 36.8 (t, 1C), 36.2 (d, 1C), 30.4 (d, 1C), 24.3 (t, 1C), 22.2 (q, 3C), 18.2 (q, 2C), 17.9 (q, 1C), 14.4 (q, 1C), 10.0 (q, 1C). ESI-MS calcd for $C_{59}H_{94}N_9O_{29}^-$ $[M-H]^-$ 1392.6163, found 1392.6156.

**Compound 26**

Starting from compound **25** (790 µg, 0.6 µmol) following general procedure VI (FUT5) compound **26** (0.6 mg, 63%) was obtained as a white solid. $^1$H (600MHz, $D_2O$): δ = 7.44-7.34 (m, 3H, $CH_{Ar}$-Phe), 7.29-7.21 (m, 2H, $CH_{Ar}$-Phe), 5.10 (d, J = 3.9 Hz, 1H, H1-Fuc), 4.87-4.71 (m, 2H, H5-Fuc, H1-GalNAc)*, 4.62 (d, J = 7.0 Hz, 1H, H1-GlcNAc), 4.61 (d, J = 2.1 Hz, 1H, $CH_\alpha$-Thr), 4.53 (d, J = 7.8 Hz, 1H, H1-Gal), 4.39-4.34 (m, 2H, $CH_\alpha$-Val, $CH_\alpha$-Phe), 4.33- 4.27 (m, 1H, $CH_\beta$-Thr), 4.24-4.16 (m, 3H, $CH_\alpha$-Ile, H2, H4-GalNAc), 4.09 (dd, J = 3.2, 10.0 Hz, 1H, H3-Gal), 4.06-4.02 (m, 1H, H5-GalNAc), 4.00-3.79 (m, 13H, H3-GalNAc, H2-H4, $H6_{a,b}$-GlcNAc, H4-Gal, $CH_2$-Gly, H3-Fuc, H5, H8-$H9_a$-$Neu_5Ac$), 3.82-3.49 (m, 13H, H2, H4-Fuc, $H6_{a,b}$-GalNAc, $H6_{a,b}$-Gal, H5-GlcNAc, H2, H5-Gal, H4, H6-H7, $H9_b$-NeuAc), 3.26 (dd, J = 6.6, 14.2 Hz, 1H, $CH_2$-Phe), 3.18 (dd, J = 6.6, 14.2 Hz, 1H, $CH_2$-Phe), 2.77 (dd, J= 4.6, 12.6 Hz, 1H, $H3_a$ NeuAc), 2.10-2.03 (m, 1H, $CH_\beta$-Val), 2.05 (s, 3H, NHAc), 2.04 (s, 3H, NHAc), 2.02 (s, 3H, NHAc), 1.90-1.82 (m, 1H, $CH_\beta$-Ile), 1.79 (t, J = 12.2, 1H, $H3_b$-NeuAc), 1.54-1.45 (m, 1H, $CH_2$-Ile), 1.32 (d, J = 6.2 Hz, 3H, $CH_3$-Thr), 1.25-1.15 (m, 1H, $CH_2$-Ile), 1.17 (d, J = 6.3 Hz, 3H, $CH_3$-Fuc), 1.00-0.92 (m, 9H, 2x$CH_3$-Val, $CH_3$-Ile), 0.87 (t, J = 7.5 Hz, 3H, $CH_3$-Ile). $^{13}$C from HSQC (150MHz, $D_2O$): δ = 129.5 (d, 2C), 128.8 (d, 3C), 101.9 (d, 1C), 101.3 (d, 1C), 98.7 (d, 1C), 98.6 (d, 1C), 76.3 (d, 1C),76.0 (d, 2C), 75.4(d, 2C), 74.7 (d, 1C),

73.1 (d, 1C), 72.8 (d, 1C), 72.2 (d, 1C), 71.9 (d, 1C), 70.8 (d, 1C), 69.2 (d, 2C), 69.0 (d, 1C), 67.8 (d, 1C), 67.6 (d, 2C), 66.92 (d, 2C), 62.6 (t, 2C), 61.5 (t, 1C), 59.4 (t, 1C), 59.2 (d, 1C), 57.9 (d, 1C), 56.9 (d, 1C), 55.8 (d, 1C), 53.9 (d, 1C), 51.4 (d, 1C), 47.8 (d, 1C), 42.1 (t, 1C), 39.4 (t, 1C), 37.1 (t, 1C), 36.5 (d, 1C), 30.2 (d, 1C), 24.4(t, 1C), 22.07 (q, 3C), 18.3 (q, 3C), 15.5 (q, 1C), 14.7 (q, 1C), 10.1 (q, 1C). ESI-MS calcd for $C_{65}H_{106}N_9O_{33}^+$ [M+H]$^+$ 1540.6888, found 1540.6884.

* Signal lies under $D_2O$-peak.
