## Supplementary material for "The receptor binding properties of H5Nx influenza A viruses have evolved to promiscuously bind to avian-type mucin-like O-glycans": Spectra

### Compound spectra for

#### This file includes:

Spectra…………………………………………………………………………...

#### NMRs of chemically synthesized products

##### Compound S5

##### $^1$H-NMR ($CDCl_3$, 600MHz)

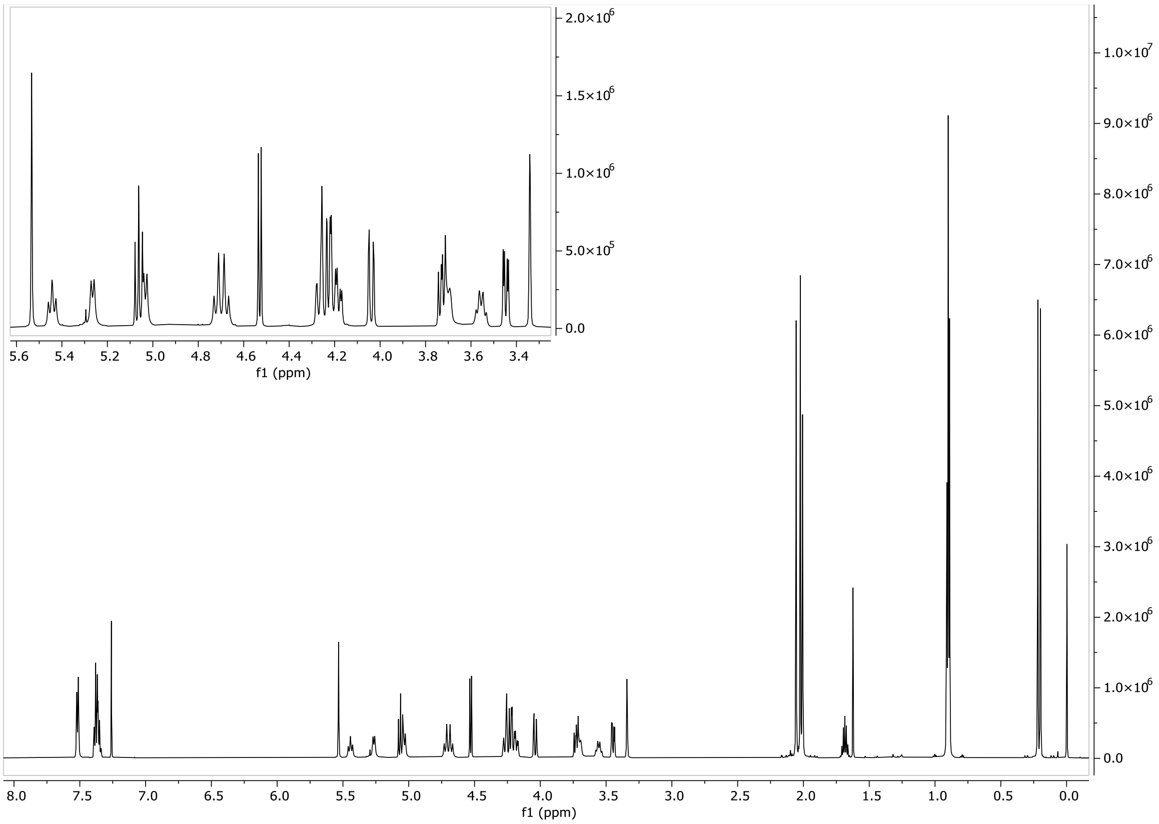


##### HSQC ($CDCl_3$, 600 MHz)

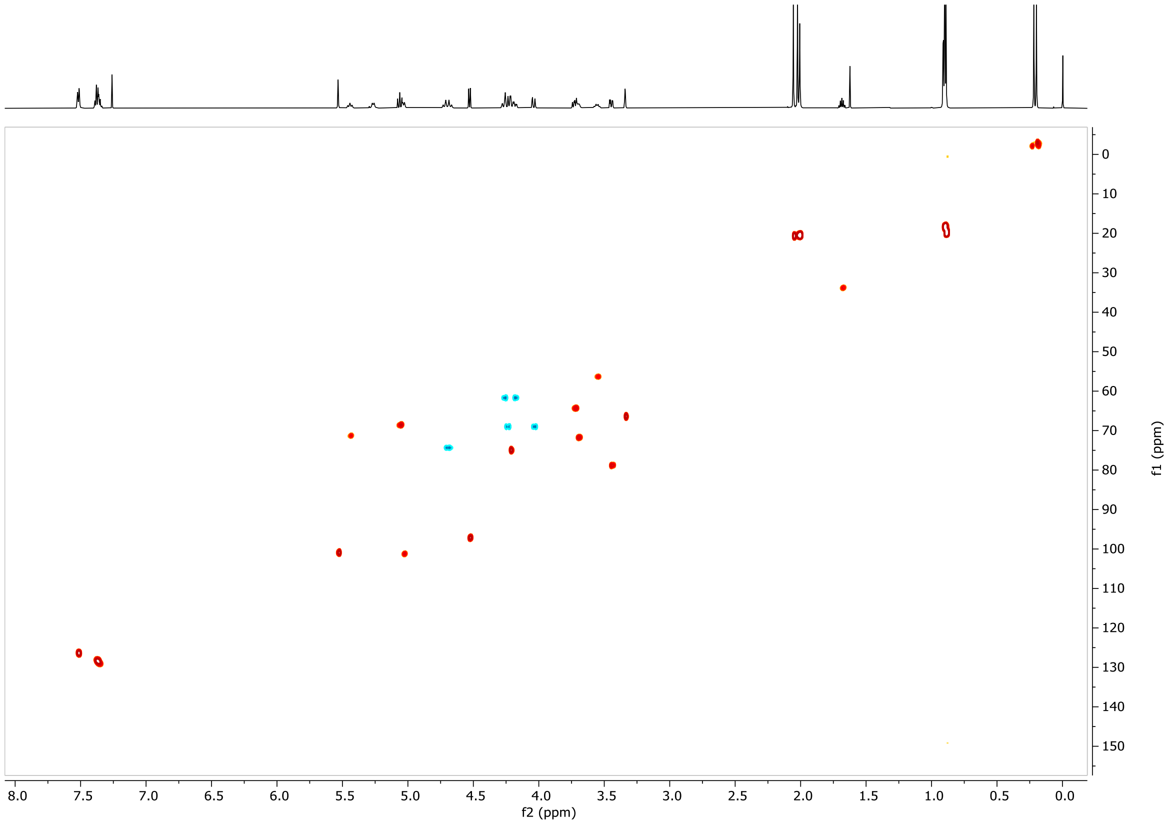

#### COSY ($CDCl_3$, 600 MHz)

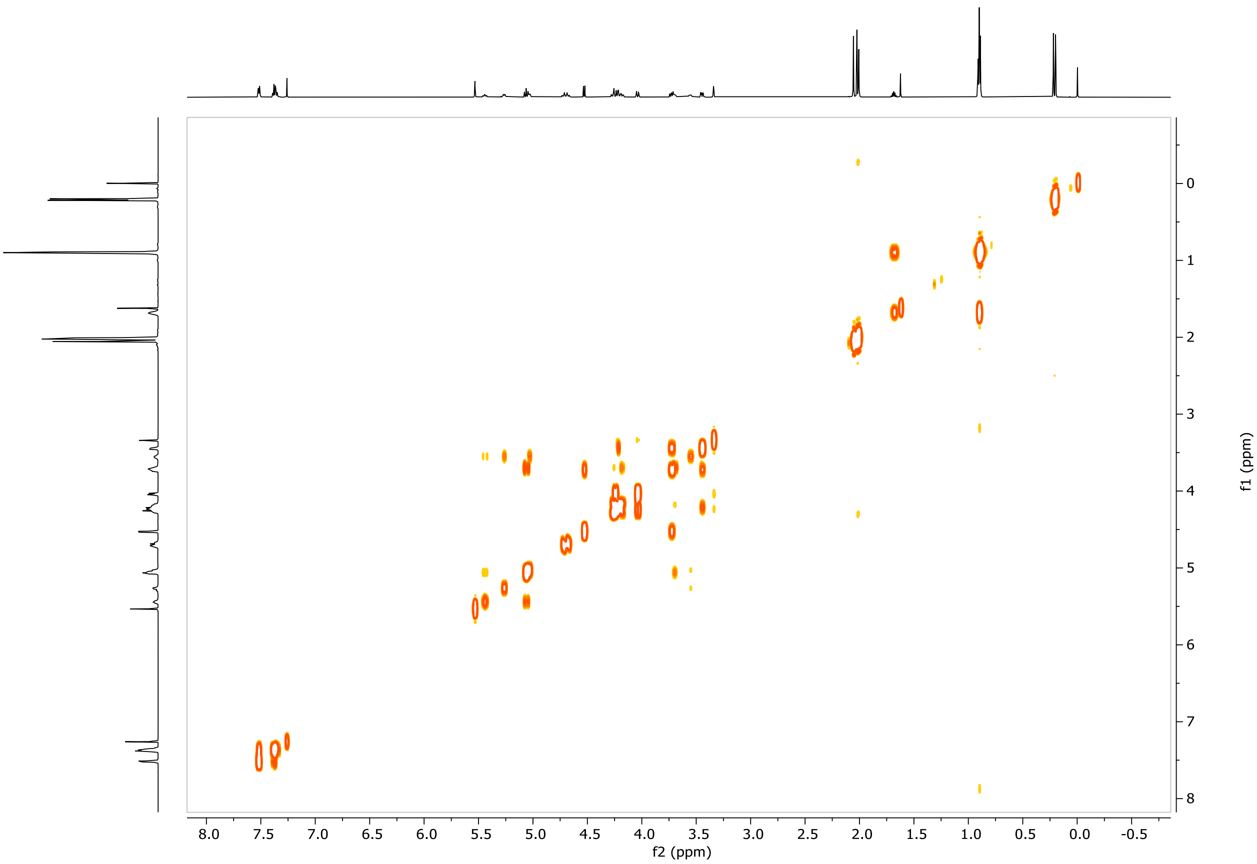


#### 13C-APT ($CDCl_3$, 600 MHz)

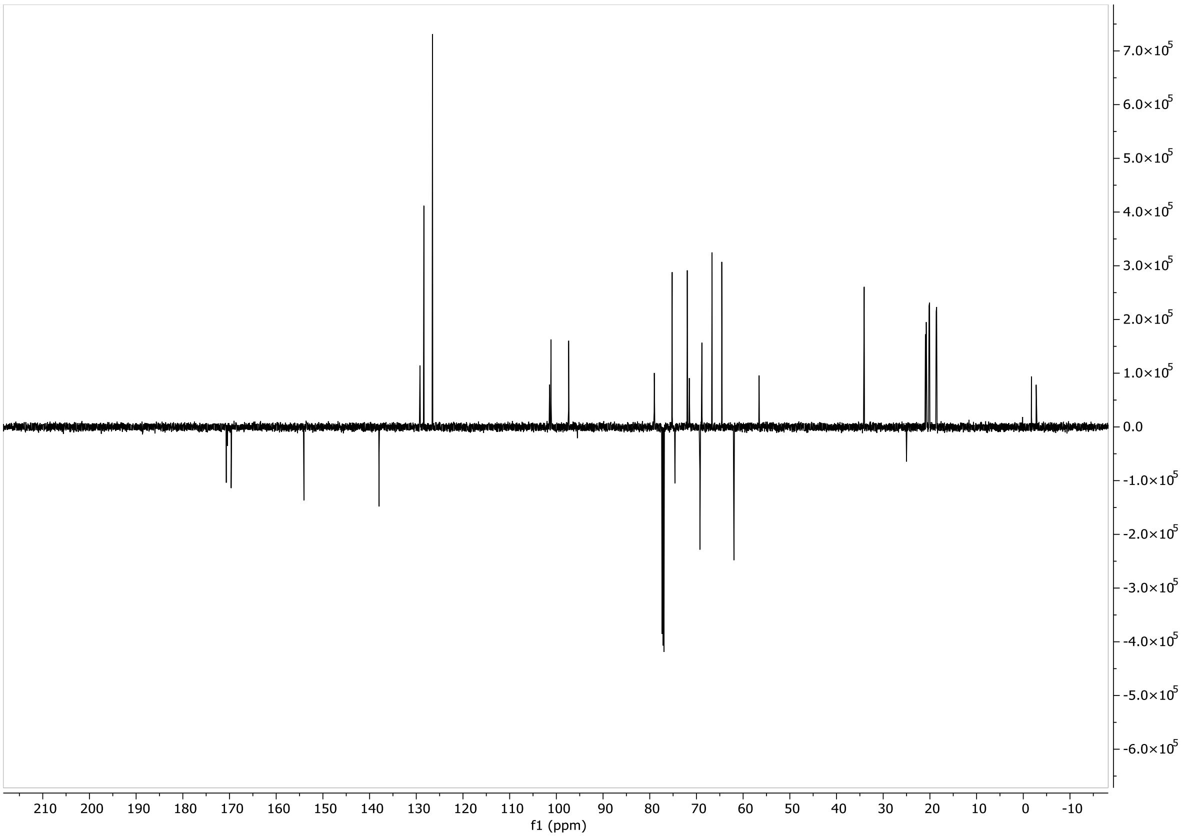

#### Compound S6

#### $^1$H-NMR ($CDCl_3$, 600MHz)

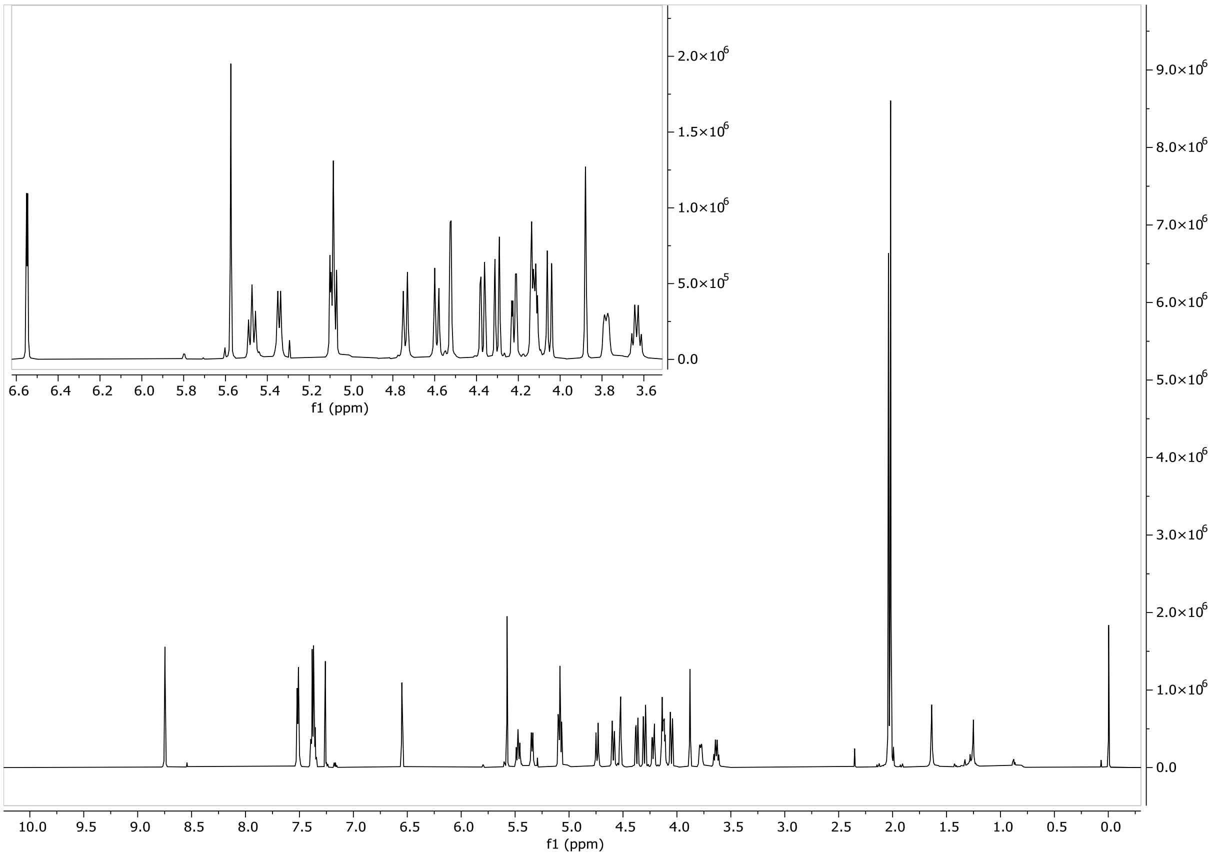


#### HSQC ($CDCl_3$, 600MHz)

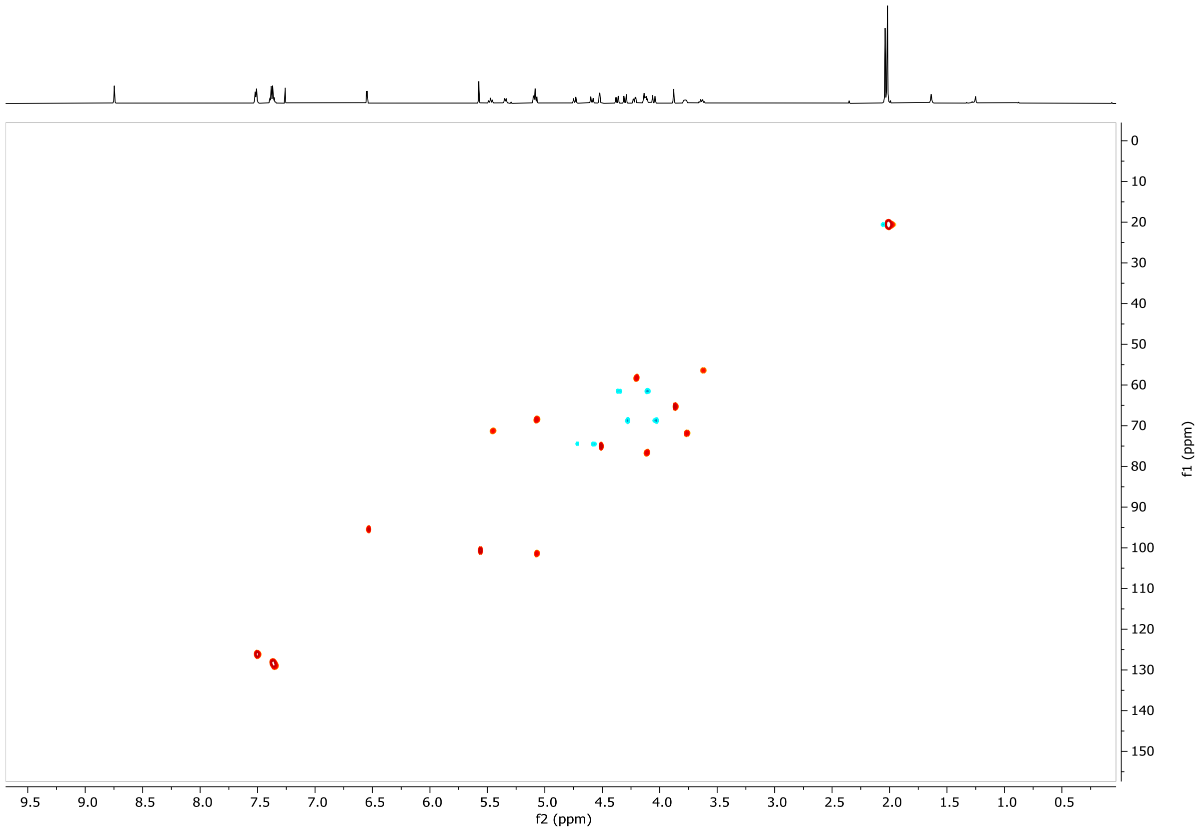

#### COSY ($CDCl_3$, 600MHz)

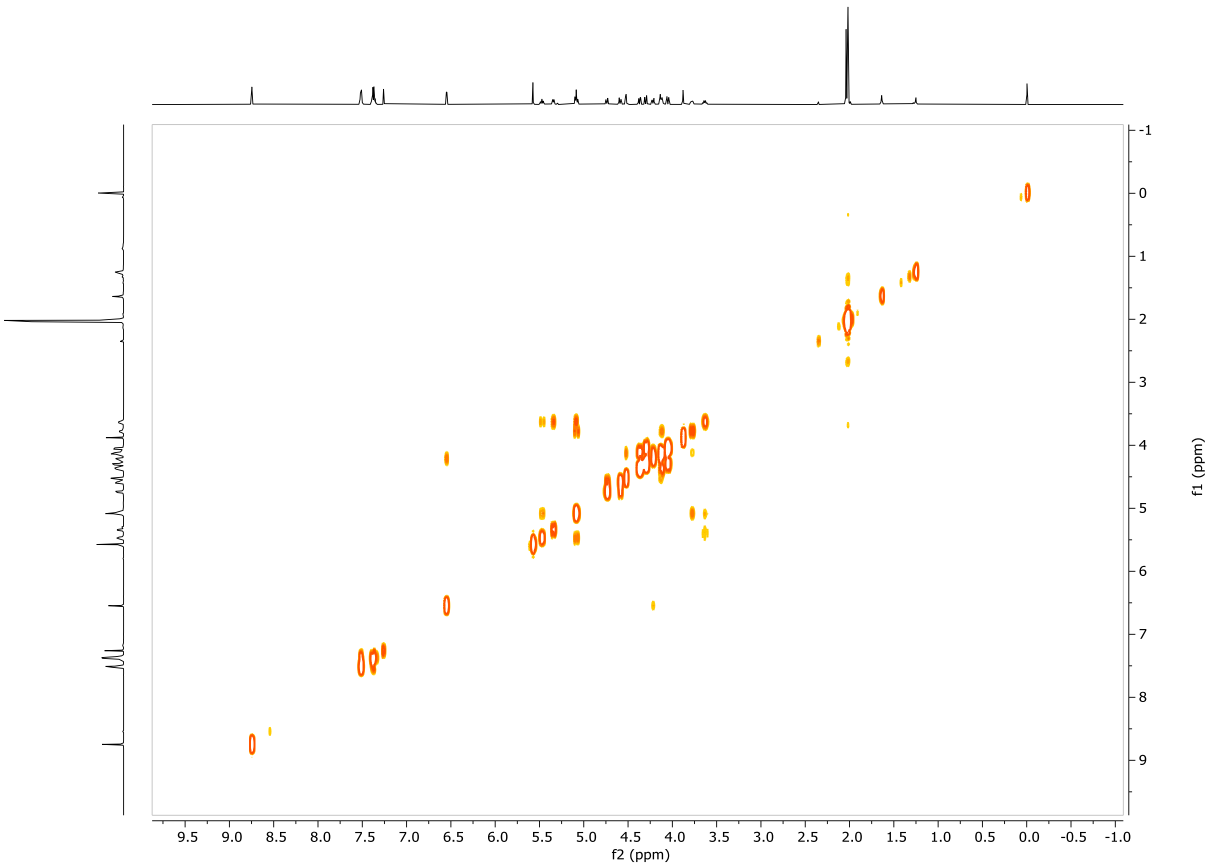


#### $^{13}C$-NMR ($CDCl_3$, 600MHz)

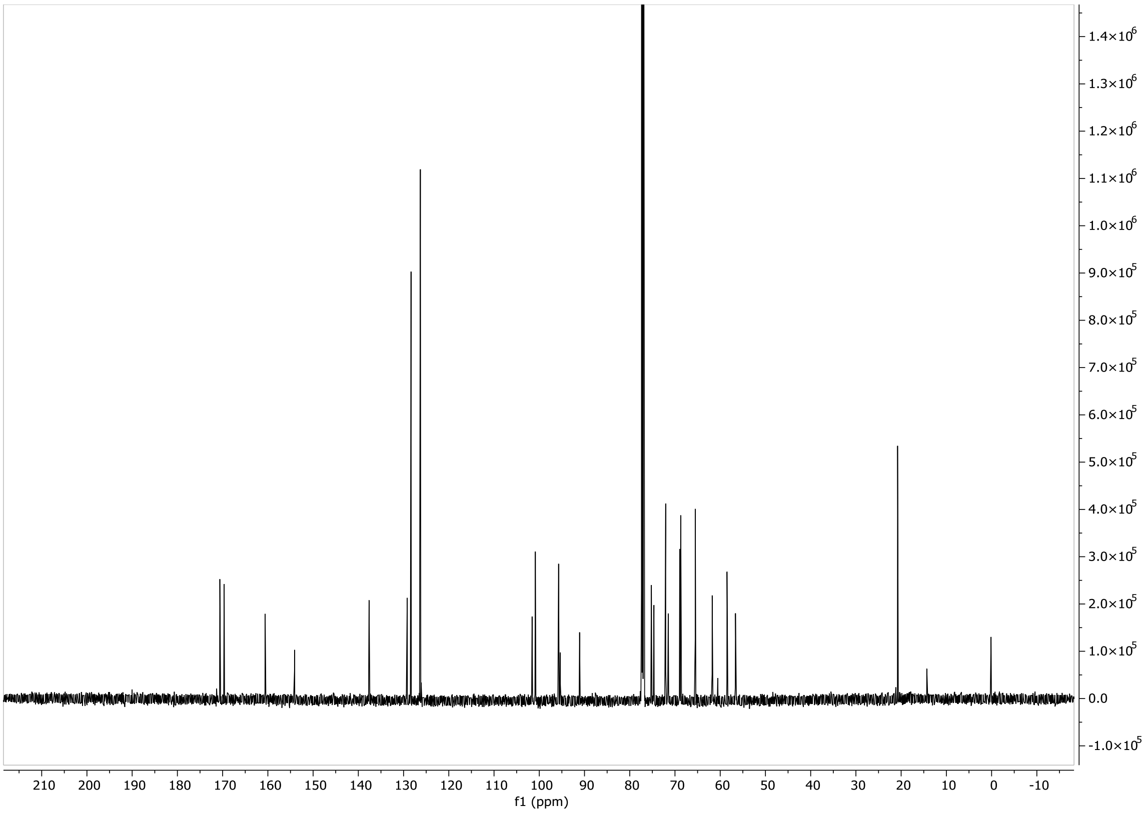

### Compound S7

#### $^1$H-NMR ($CDCl_3$, 600MHz)

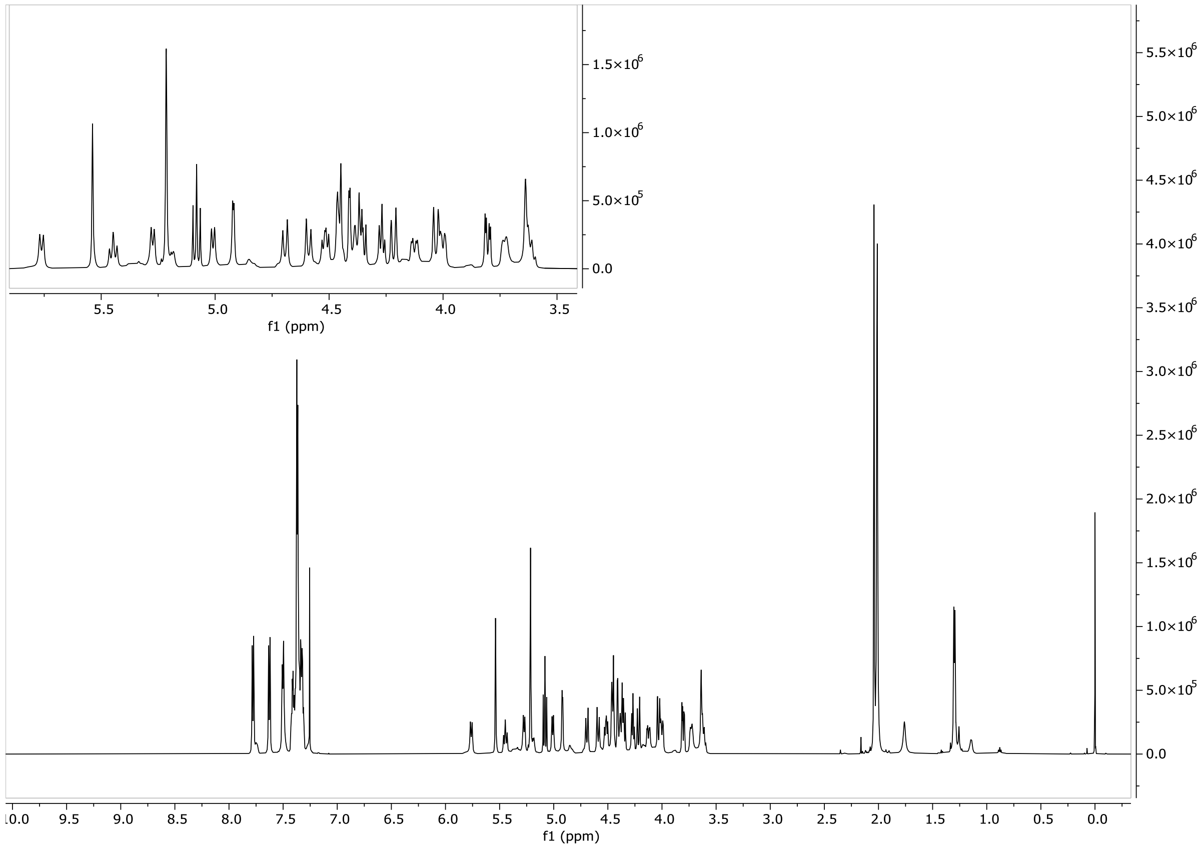


#### HSQC ($CDCl_3$, 600MHz)

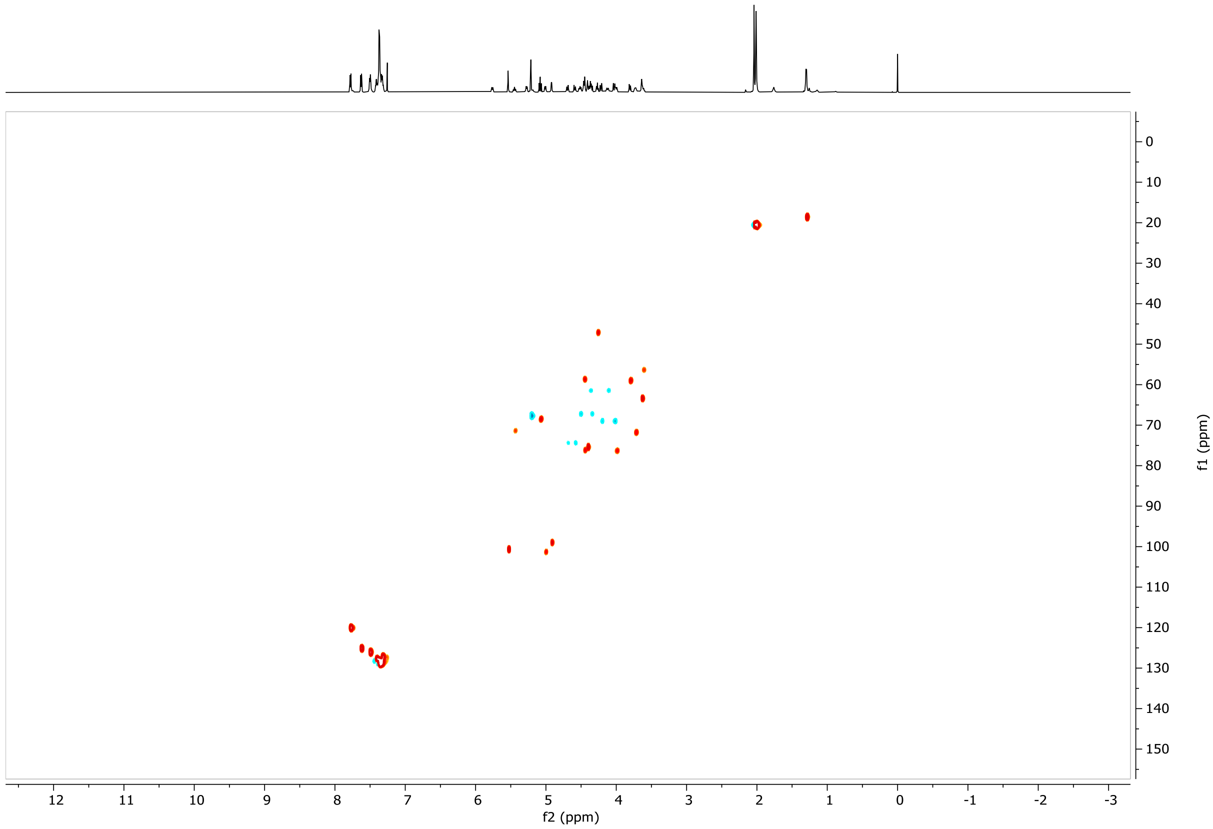

#### COSY ($CDCl_3$, 600MHz)

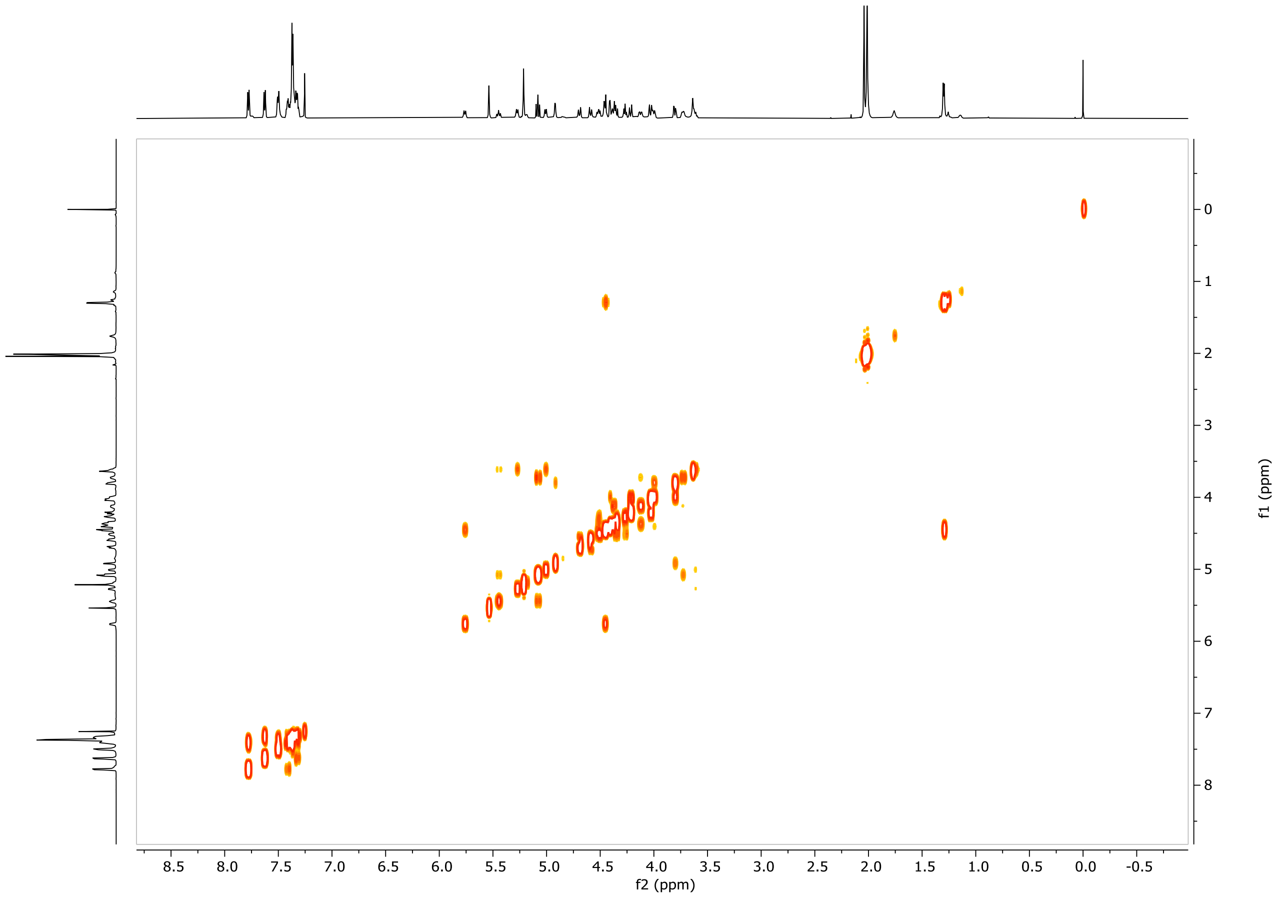


#### $^{13}C$-NMR ($CDCl_3$, 600MHz)

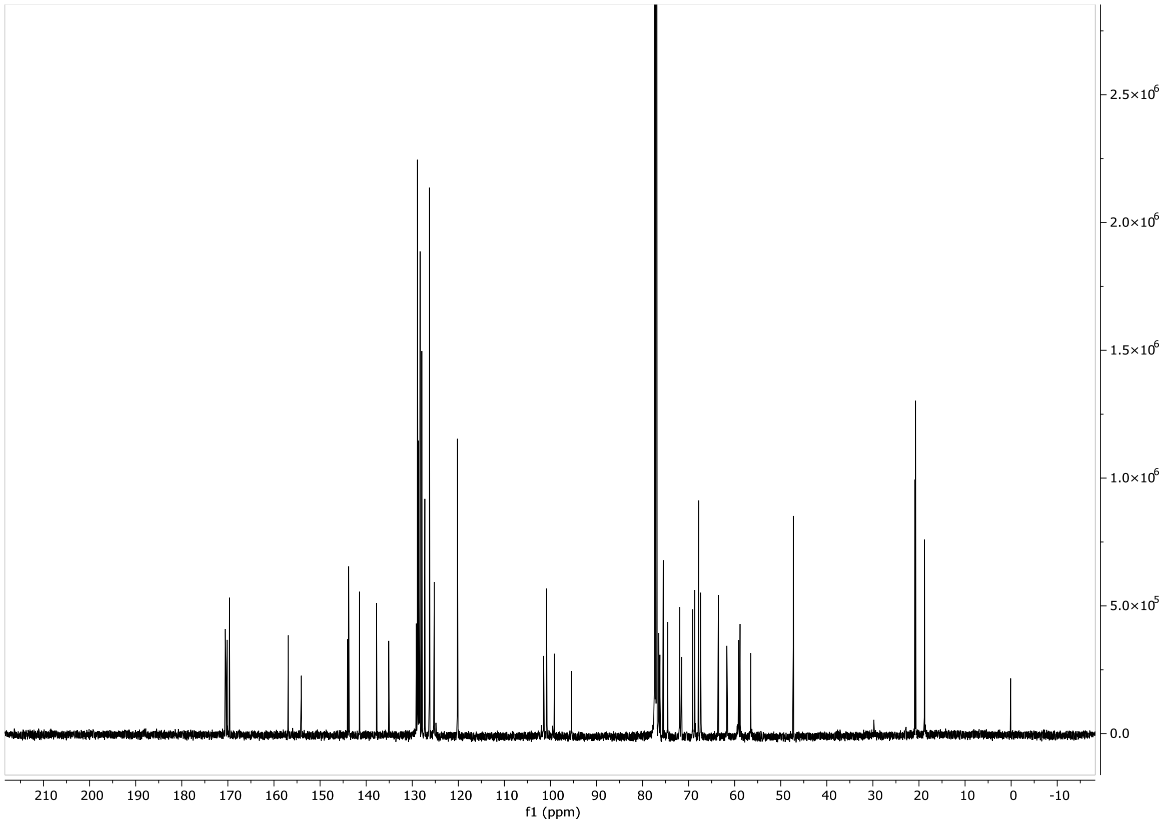

### Compound S8

#### $^1$H-NMR ($CDCl_3$, 600MHz)

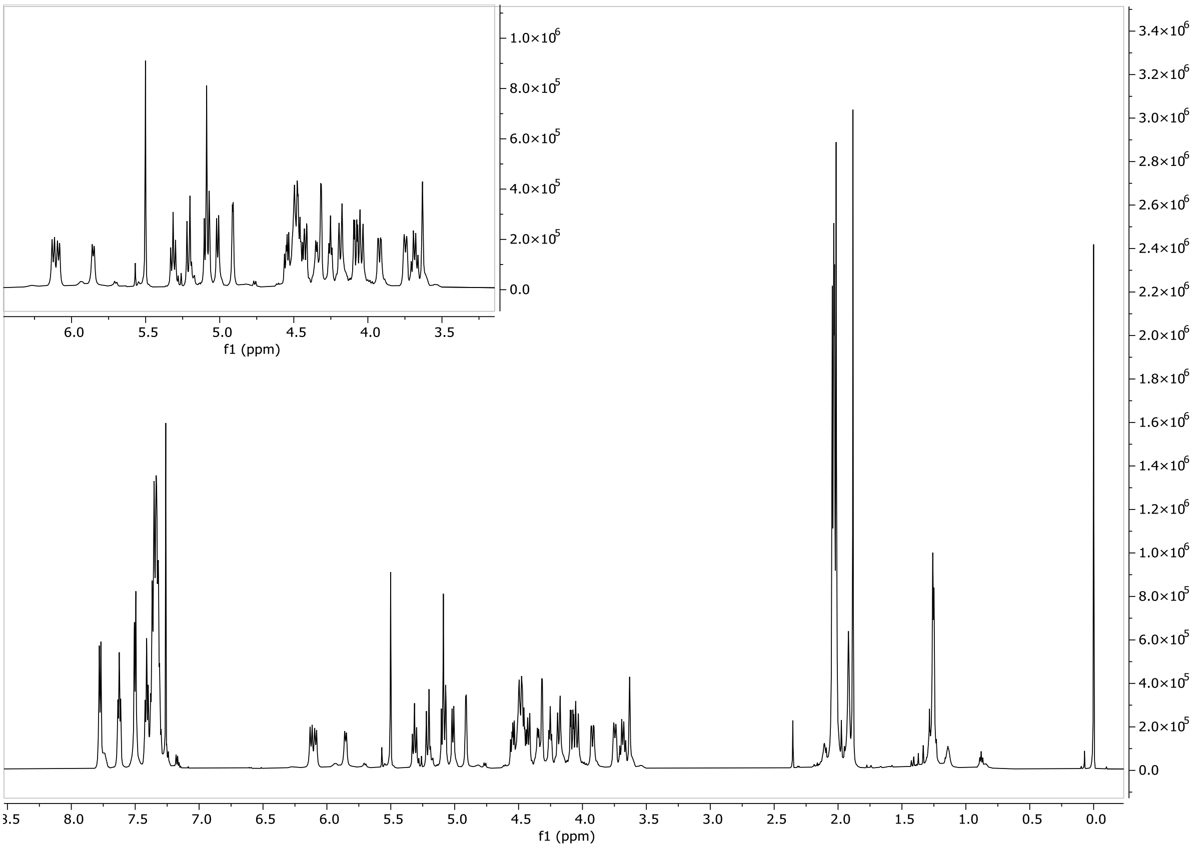


#### HSQC ($CDCl_3$, 600MHz)

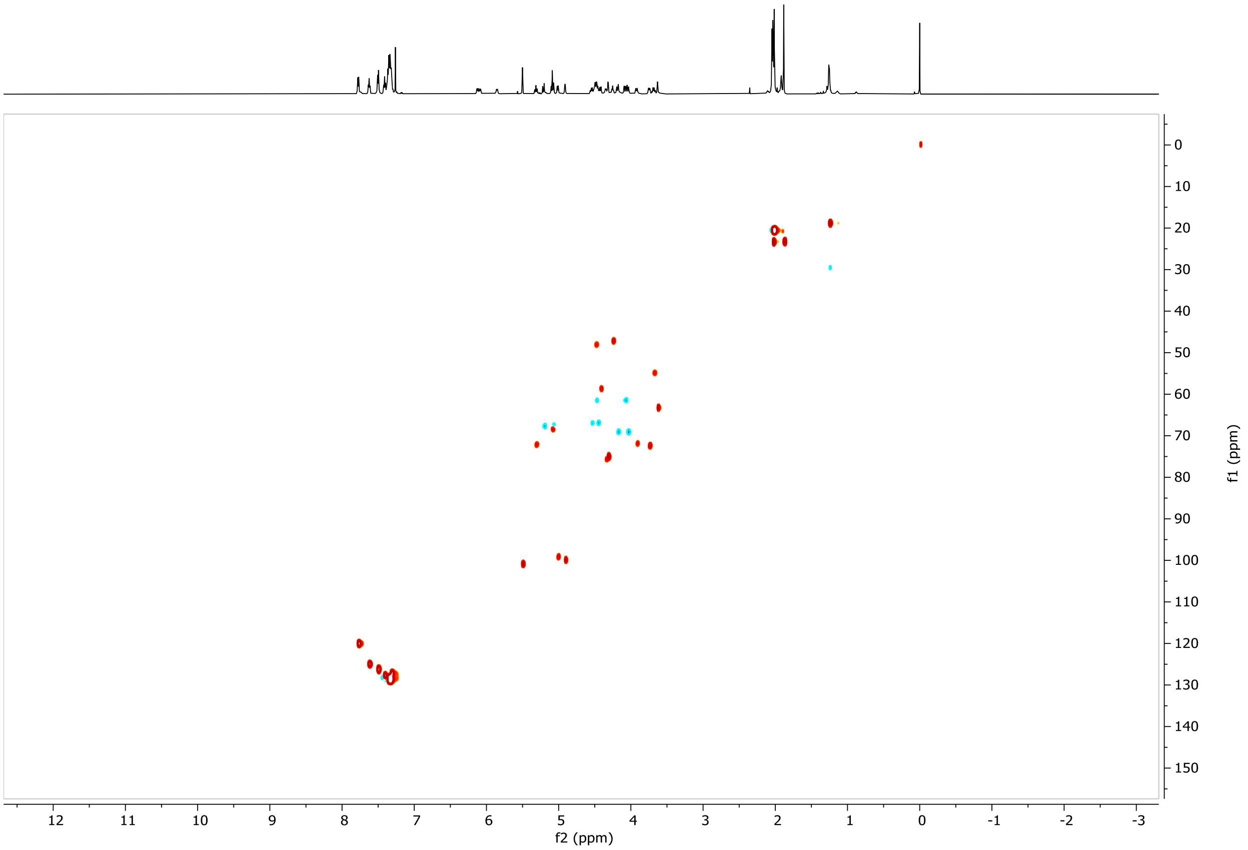

#### COSY ($CDCl_3$, 600MHz)

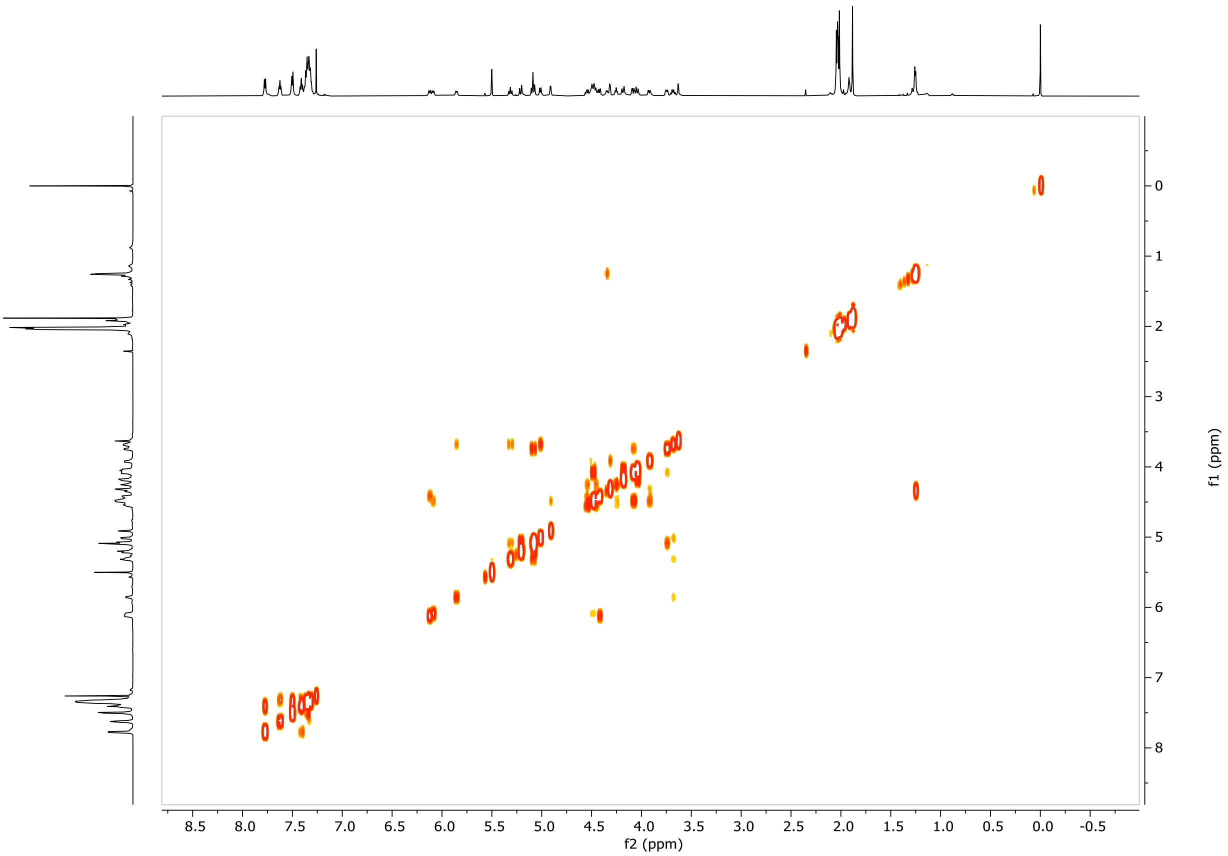


#### $^{13}C$-NMR ($CDCl_3$, 600MHz)

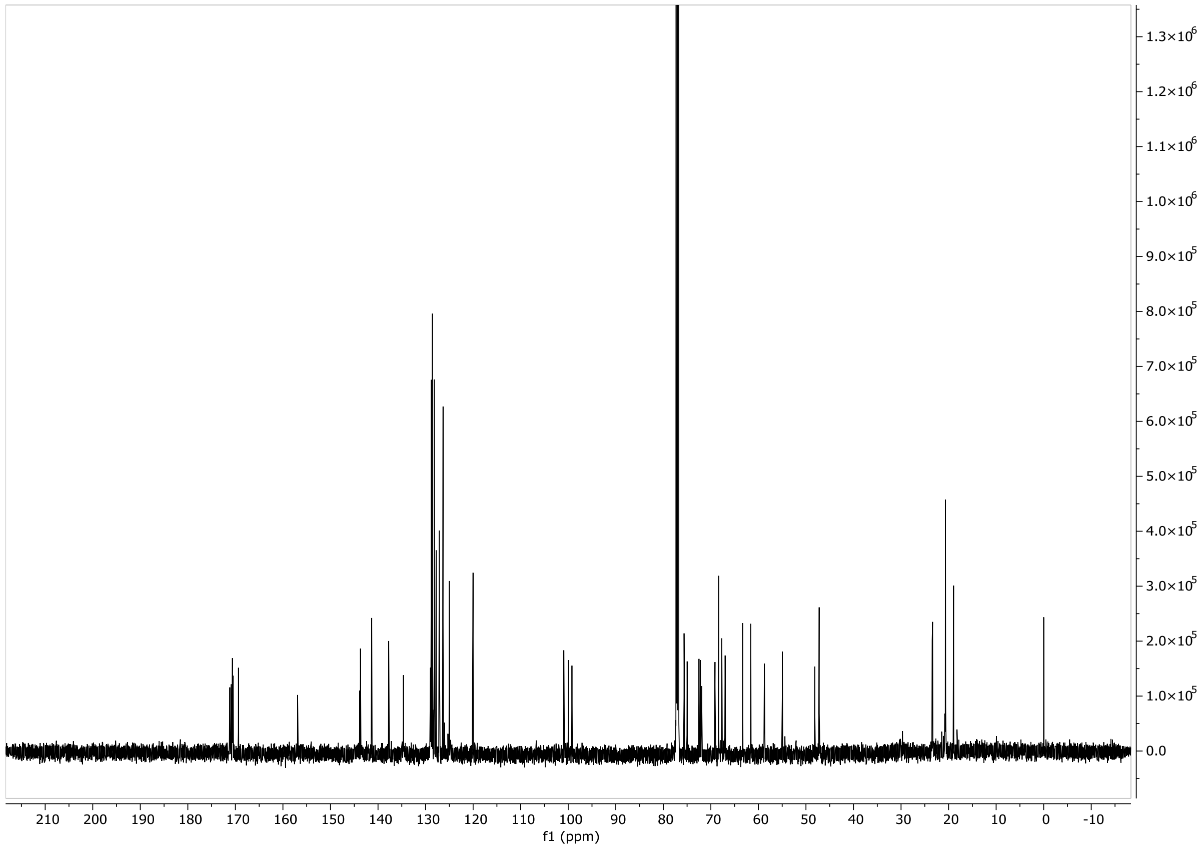

### Compound S9

#### $^1$H-NMR ($CDCl_3$, 600MHz)

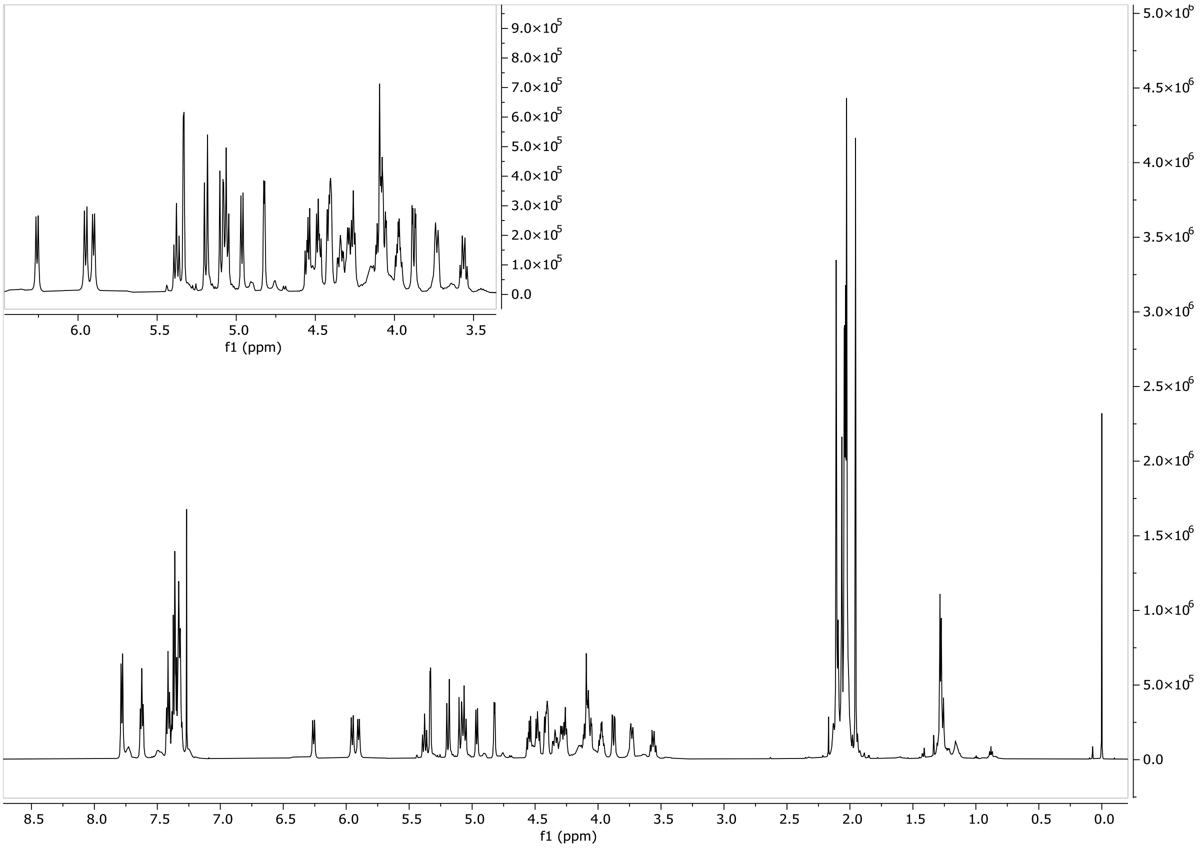


#### HSQC ($CDCl_3$, 600MHz)

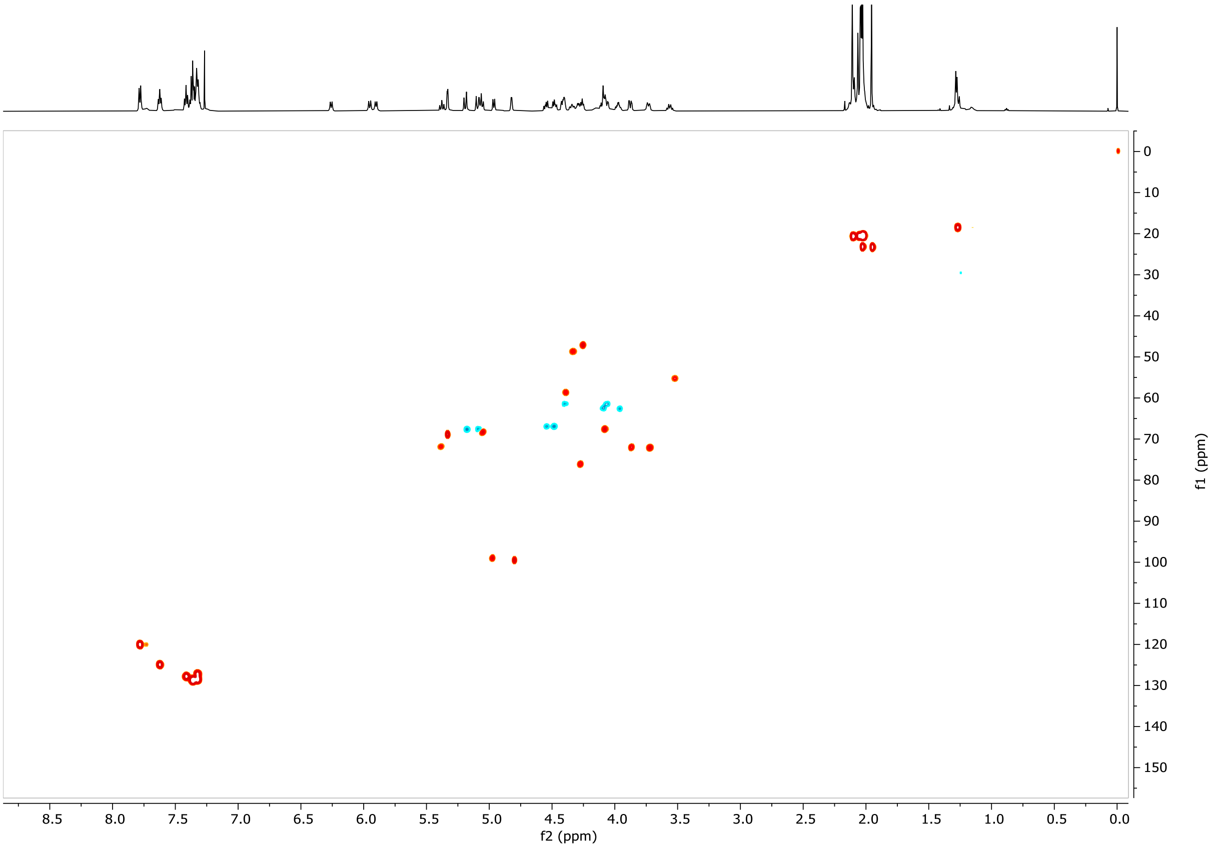

#### COSY ($CDCl_3$, 600MHz)

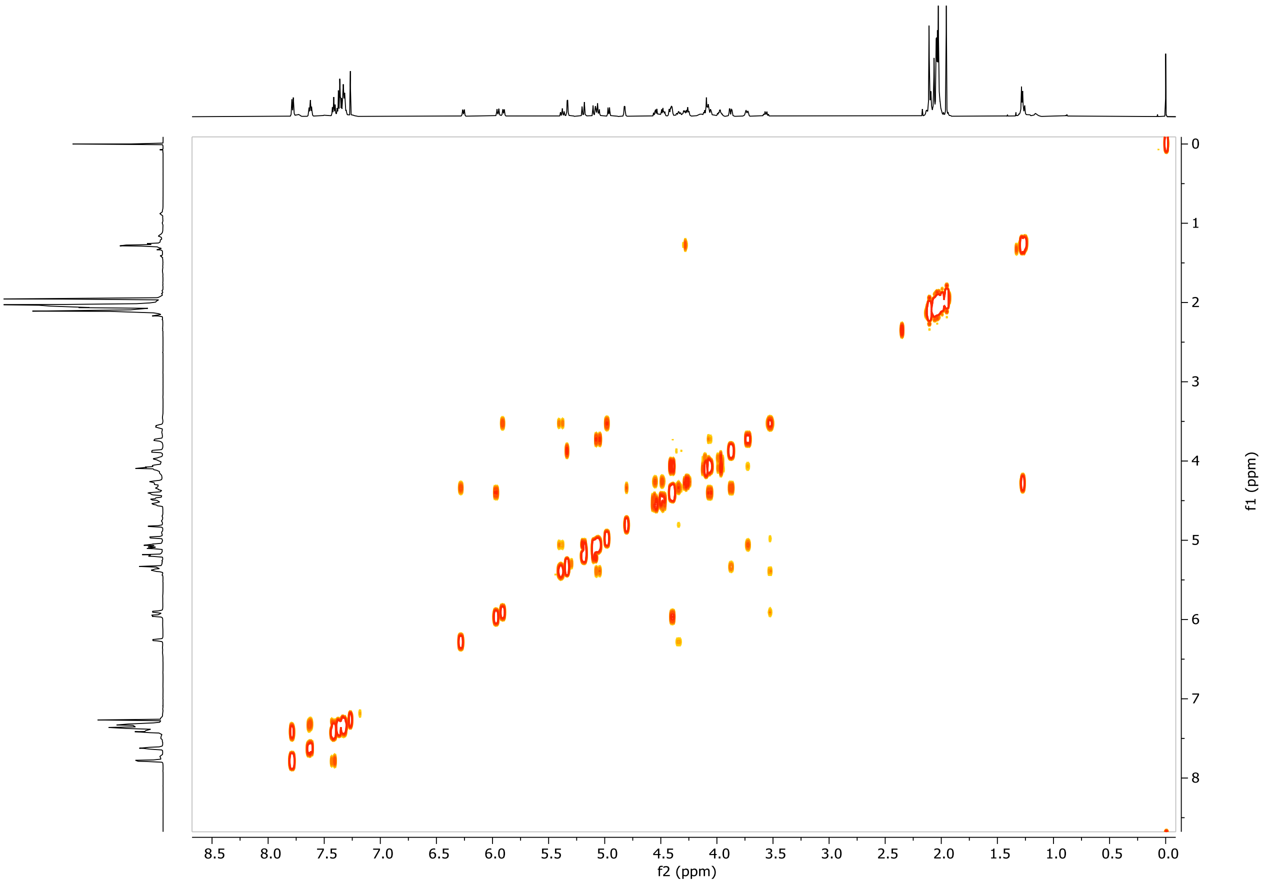


#### $^{13}$C-NMR ($CDCl_3$, 600MHz)

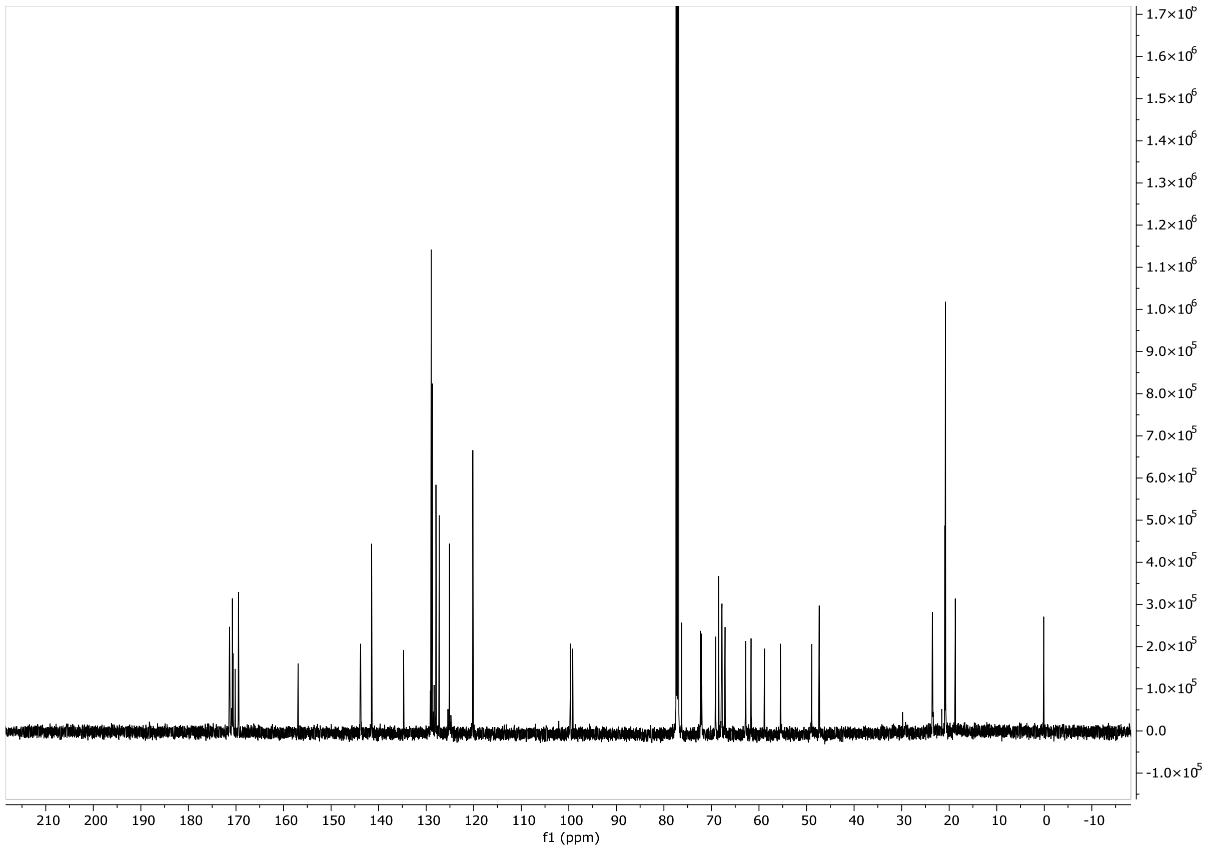

#### Compound S10

#### $^1$H-NMR ($CD_3OD$, 600MHz)

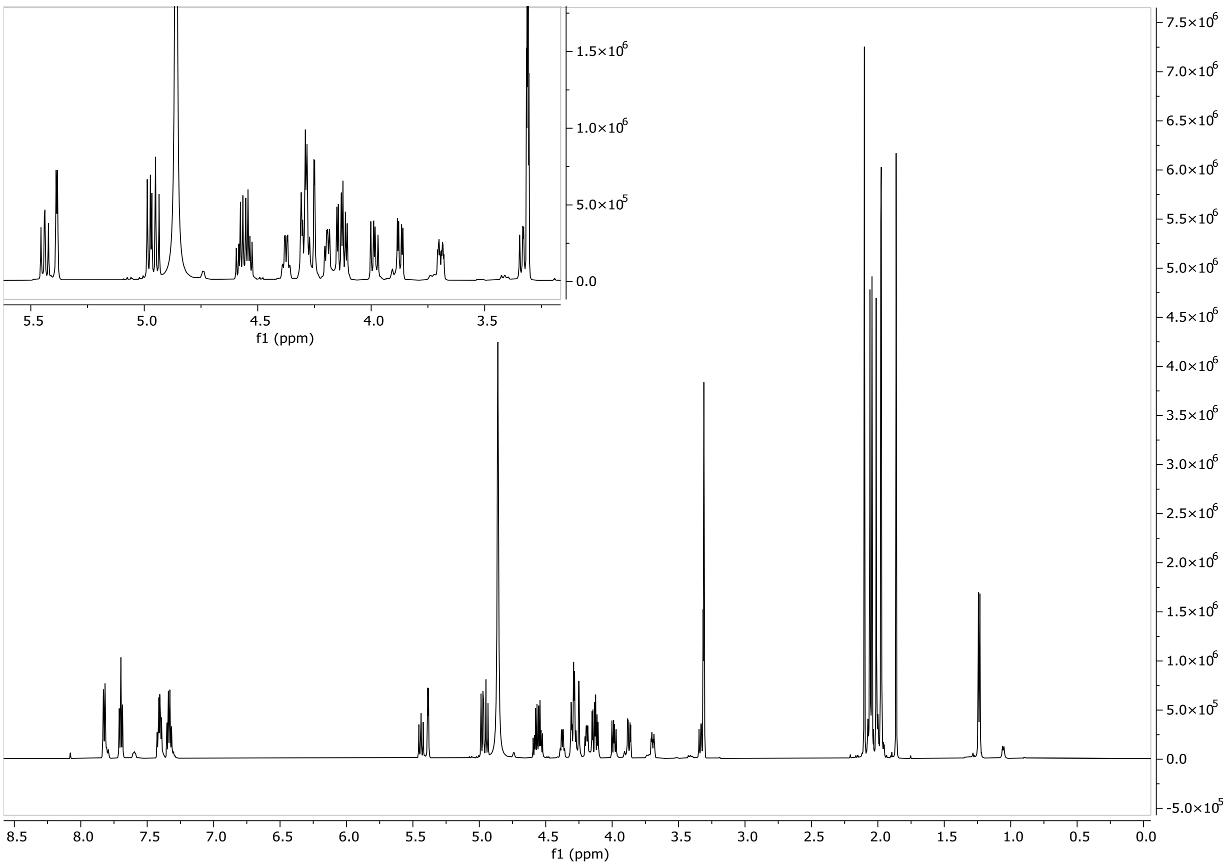


#### HSQC ($CD_3OD$, 600MHz)

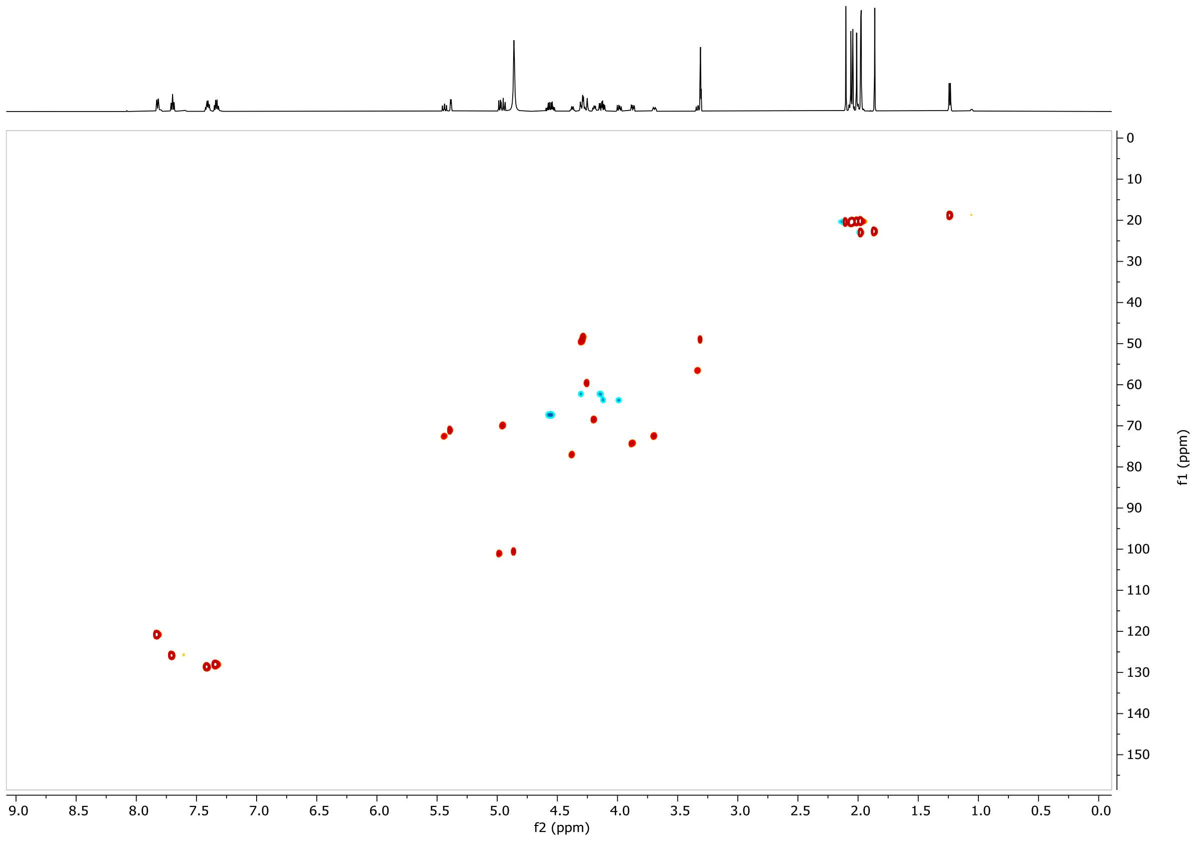

#### COSY ($CD_3OD$, 600MHz)

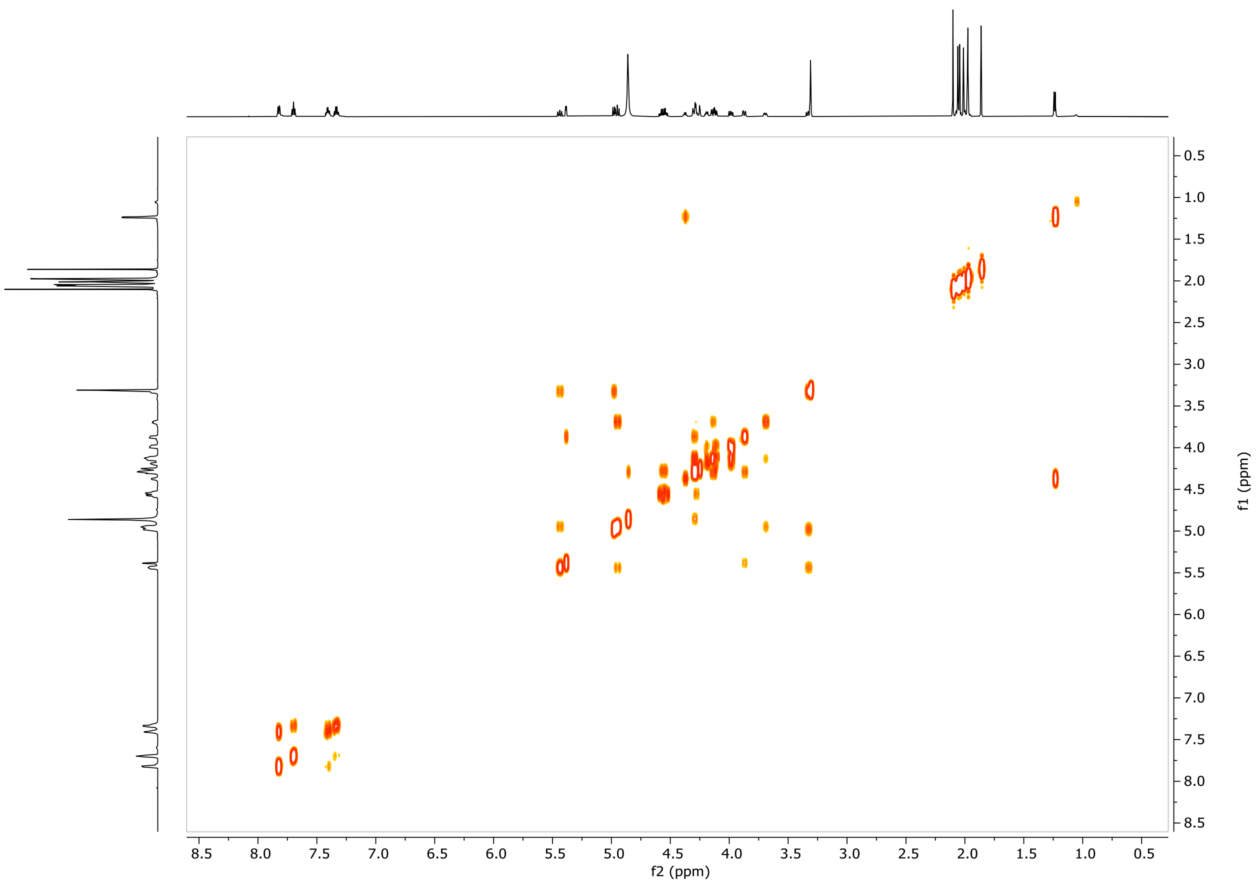


#### $^{13}C$-NMR ($CD_3OD$, 600MHz)

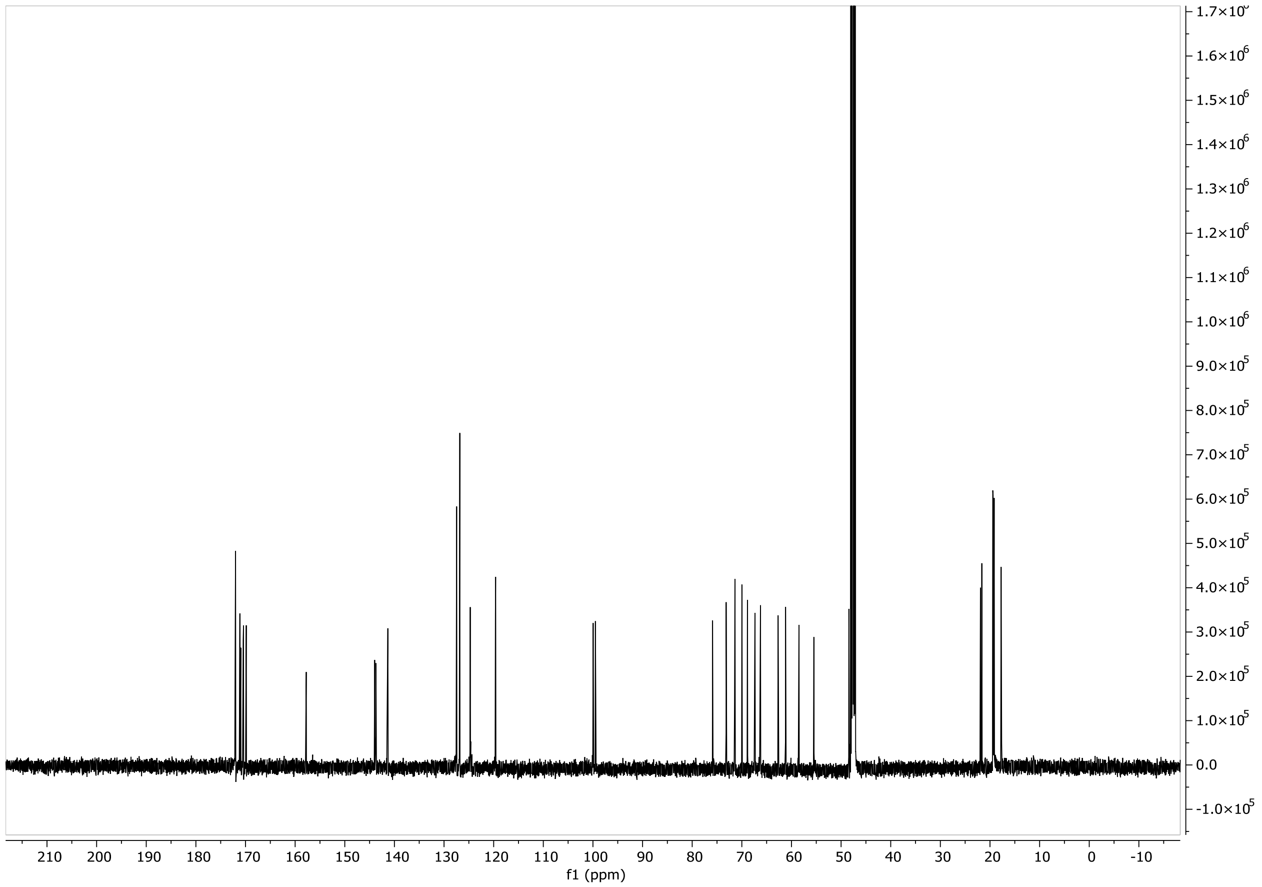

#### NMR and LC-MS spectra of enzymatically prepared products

##### Core1 (2)

##### $^{1}$H-NMR ($D_2O$, 600MHz)

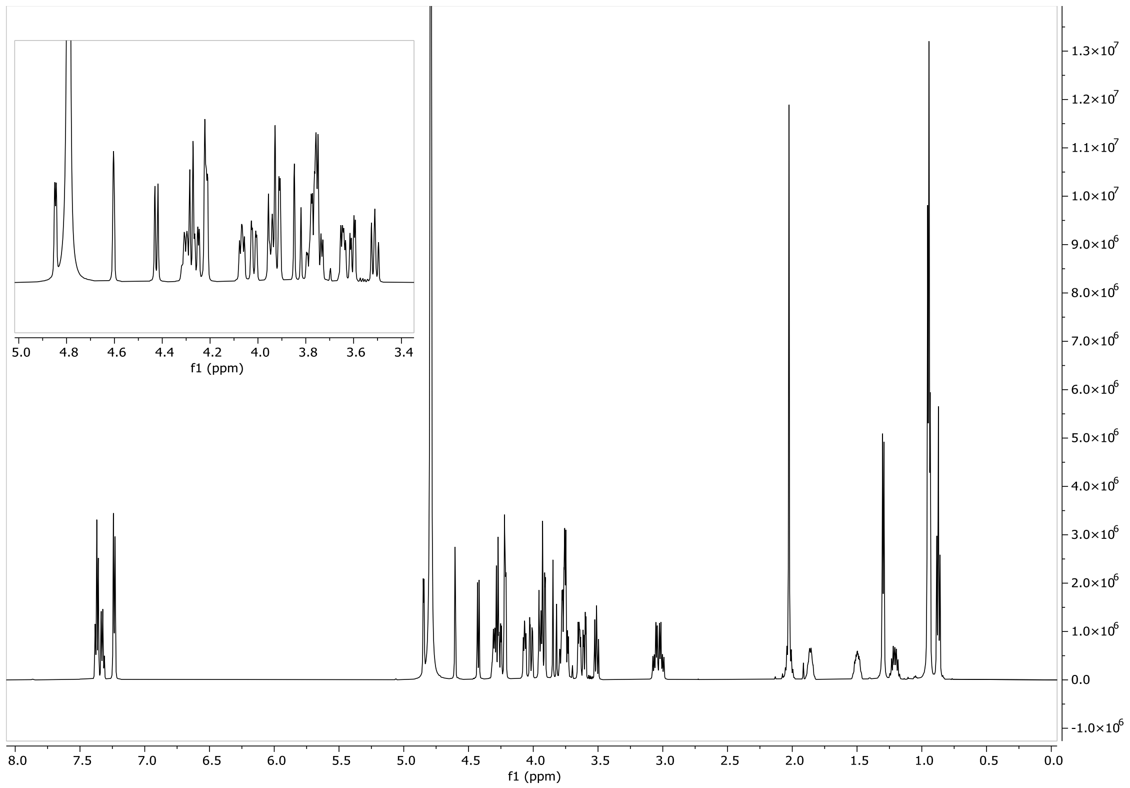


##### HSQC ($D_2O$, 600 MHz)

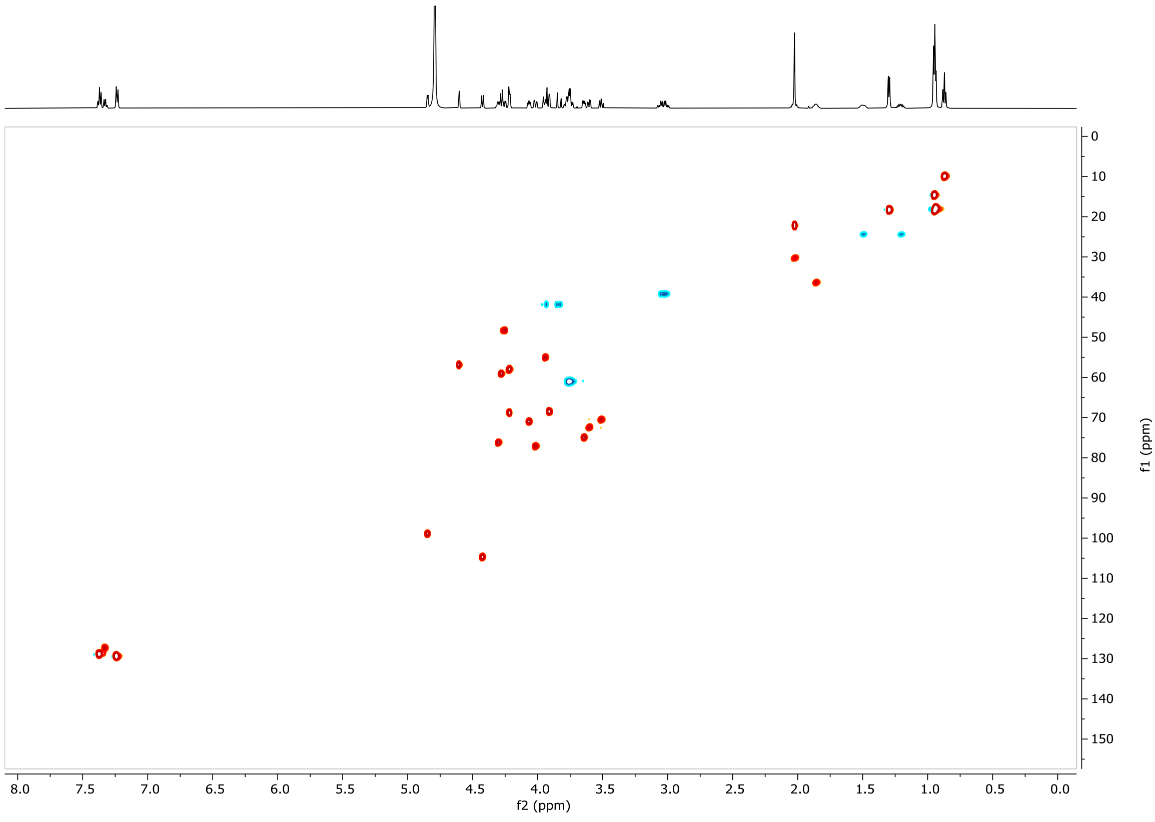

#### LCMS

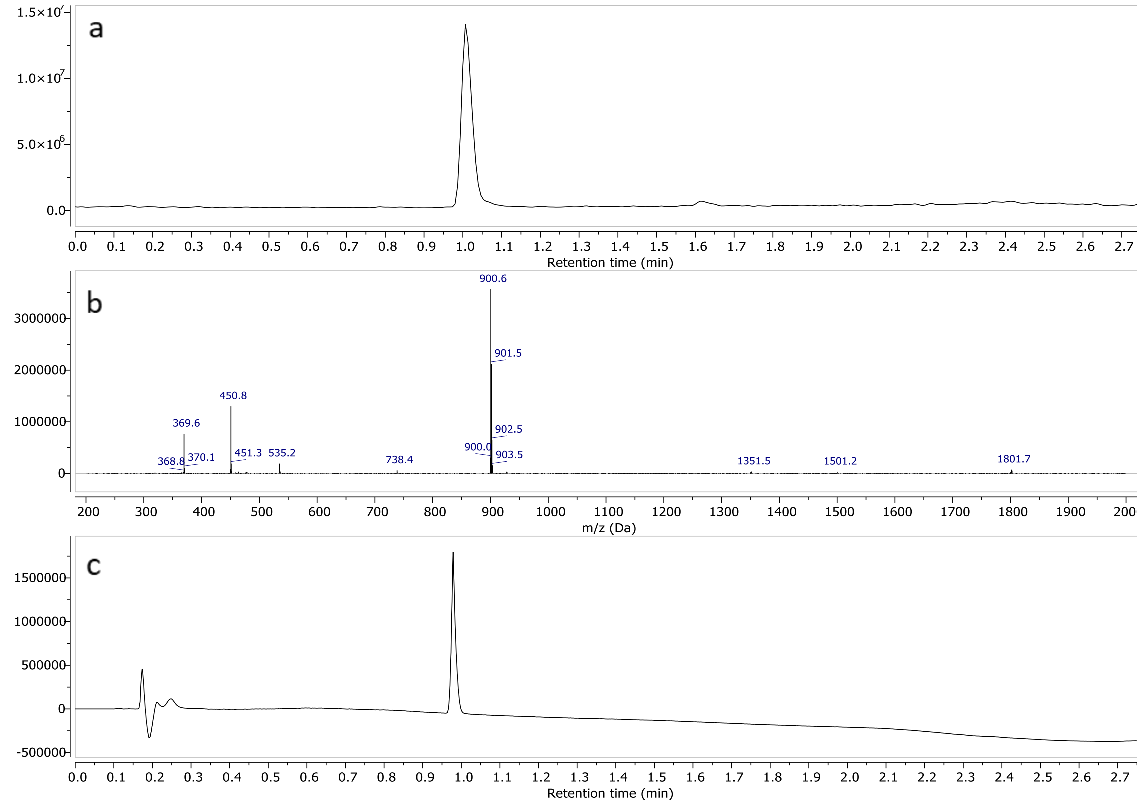


a) TIC – Total Ion Chromatogram (ESI+) b) mass spectrum of **2** (ESI+) c) UV spectrum at 190nm

#### Core 2 (3)

#### $^1$H-NMR ($D_2O$, 600MHz)

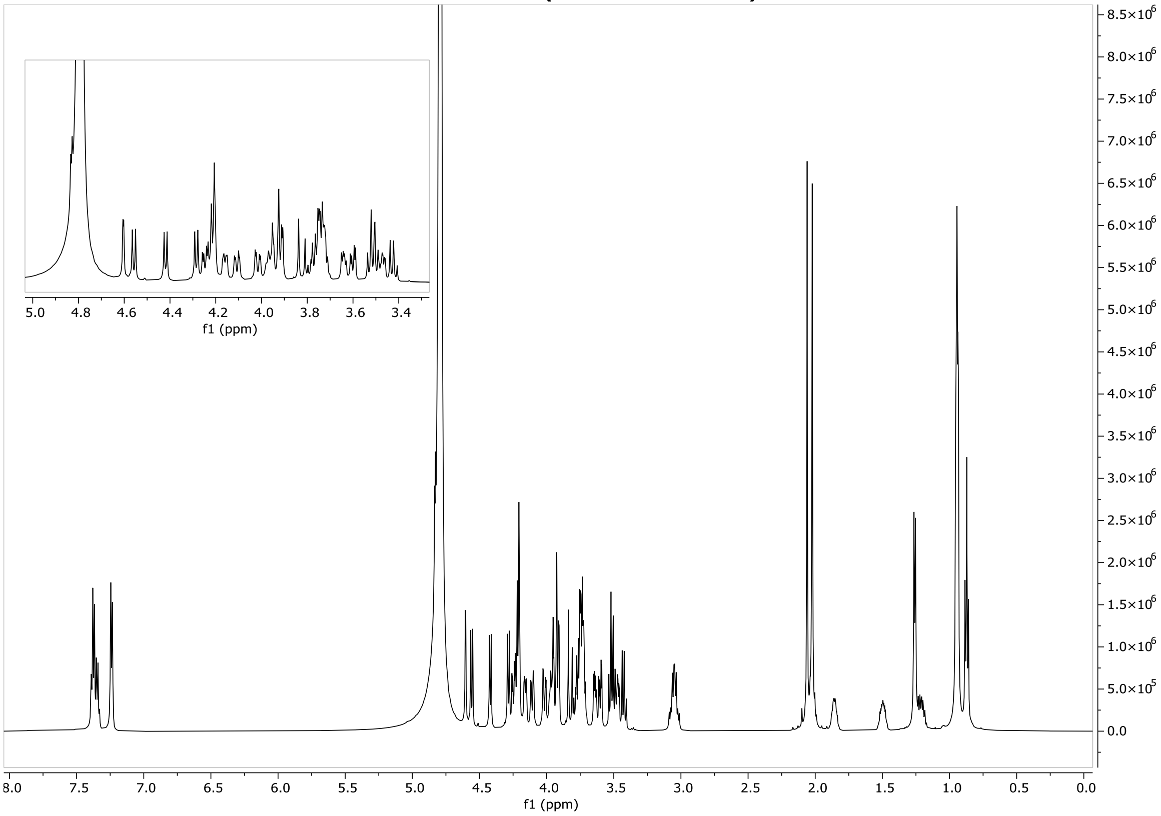

#### HSQC ($D_2O$, 600 MHz)

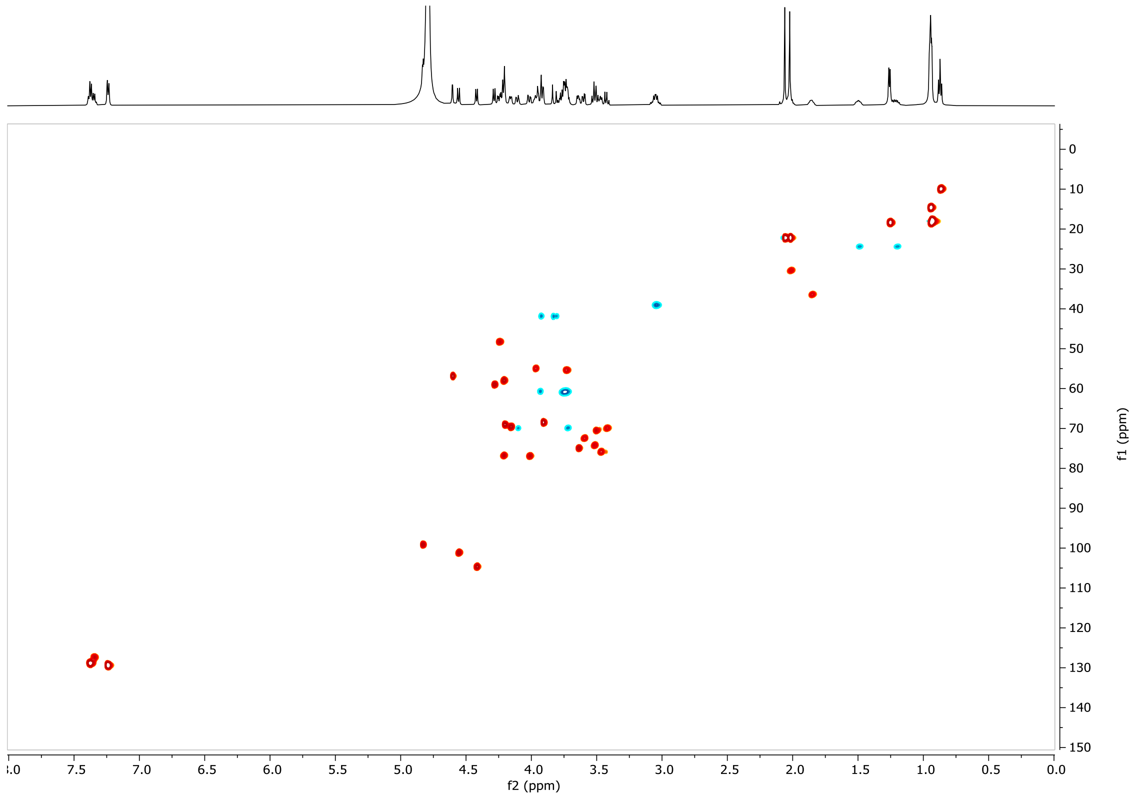


#### LCMS

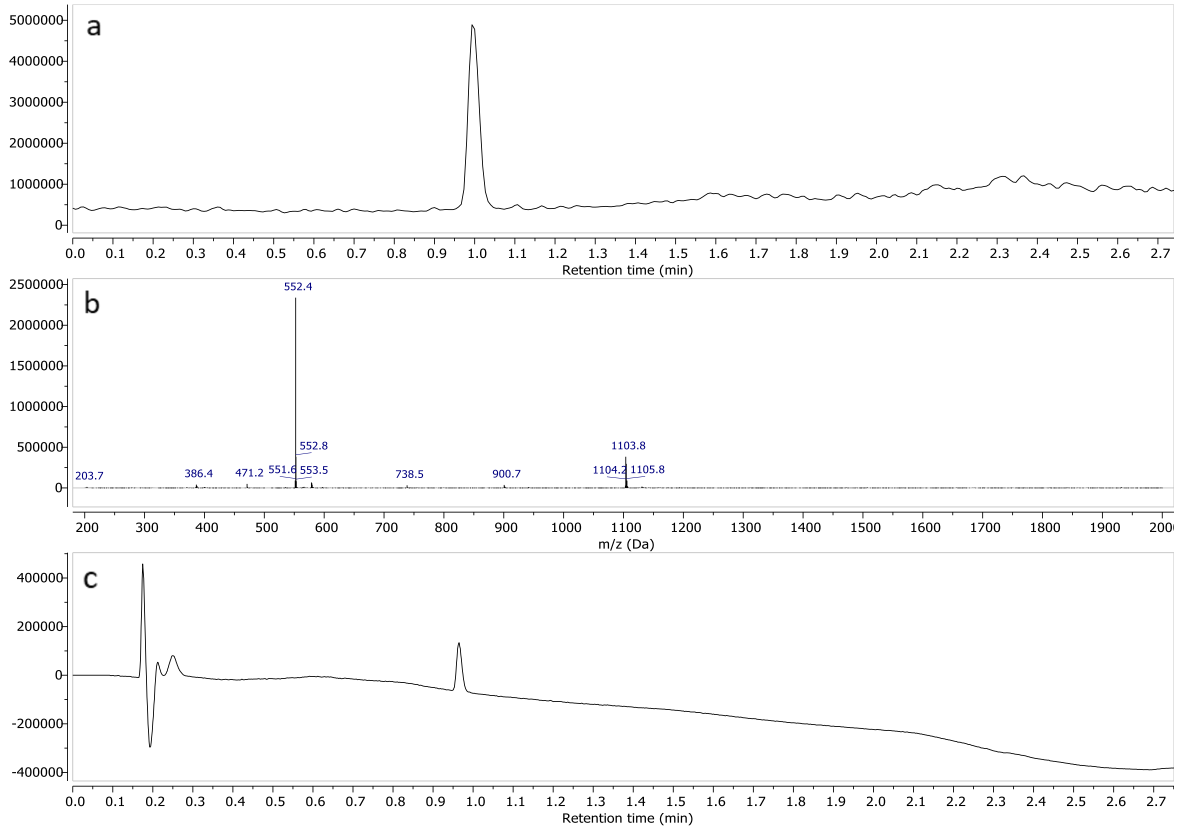


a) TIC – Total Ion Chromatogram (ESI+) b) mass spectrum of **3** (ESI+) c) UV spectrum at 190nm

#### Compound 4

##### $^1$H-NMR ($D_2O$, 600MHz)

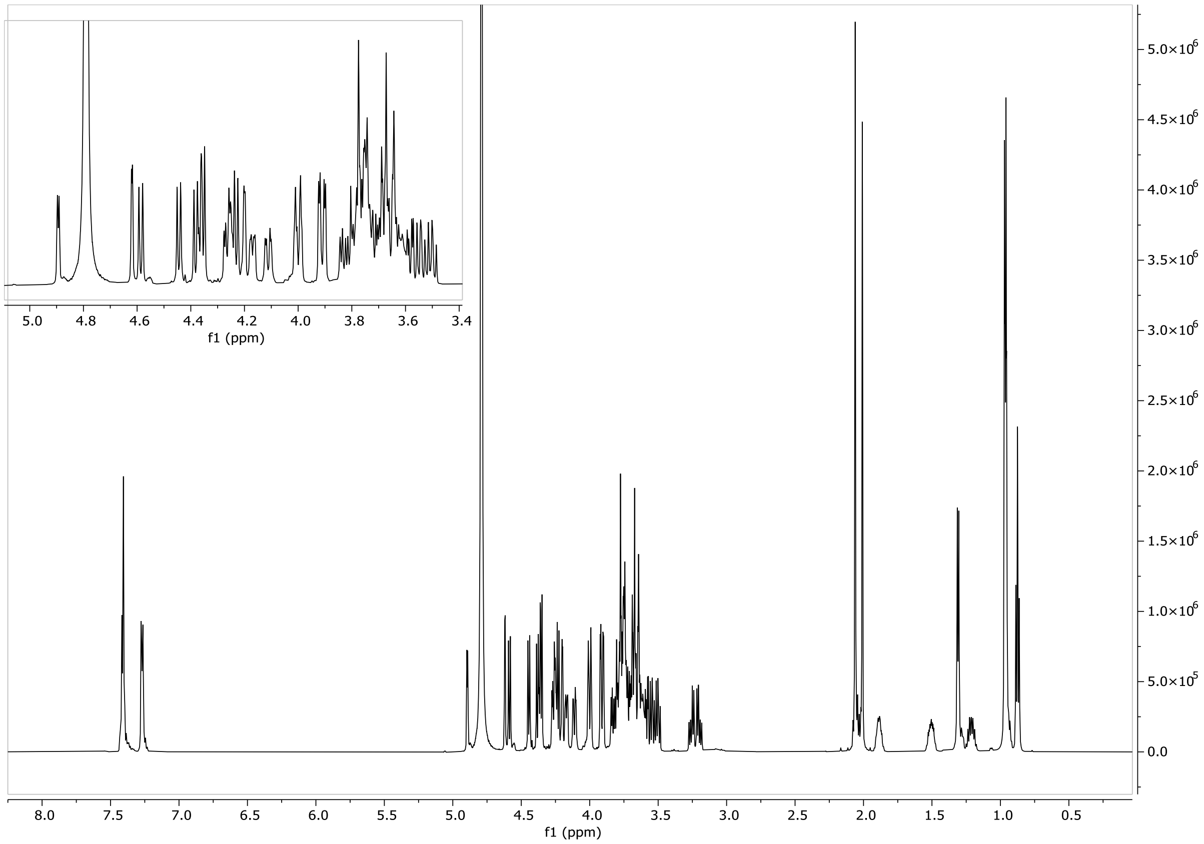


##### HSQC ($D_2O$, 600 MHz)

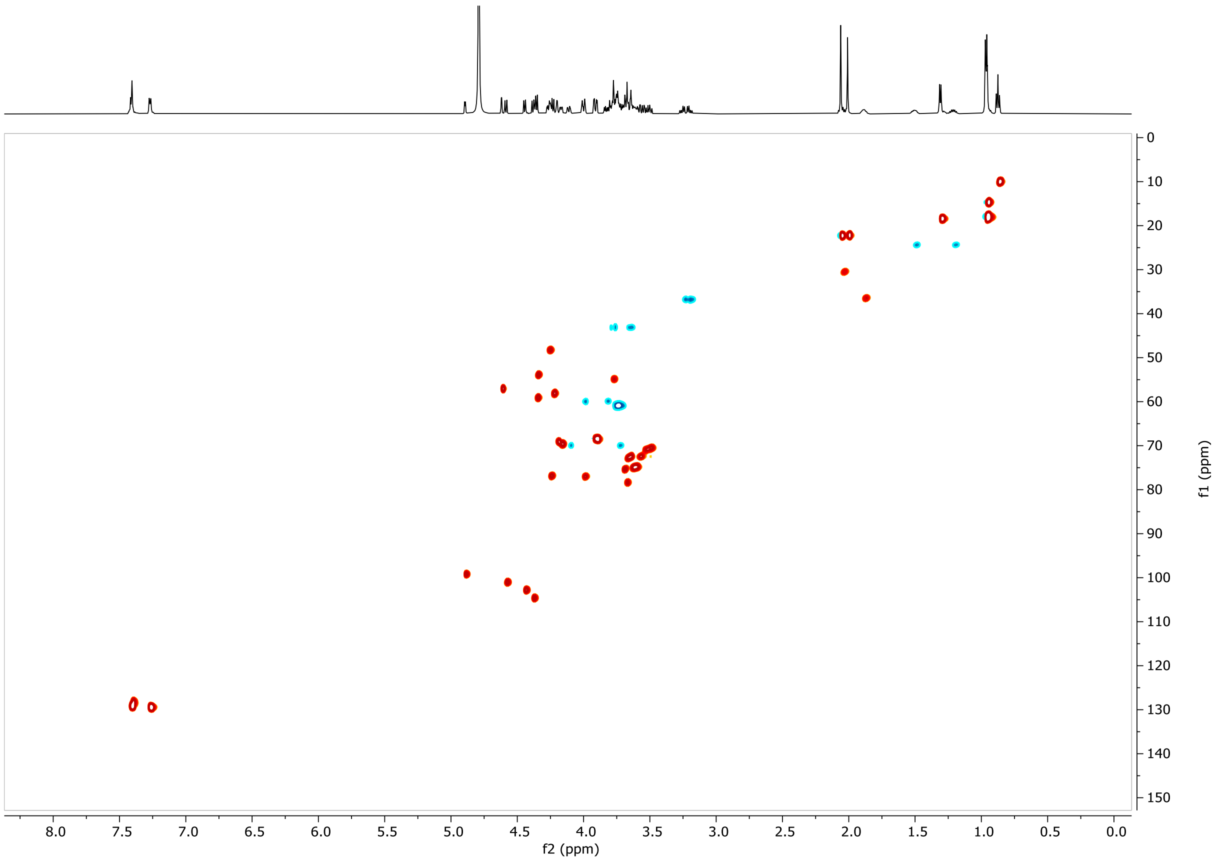

#### LCMS

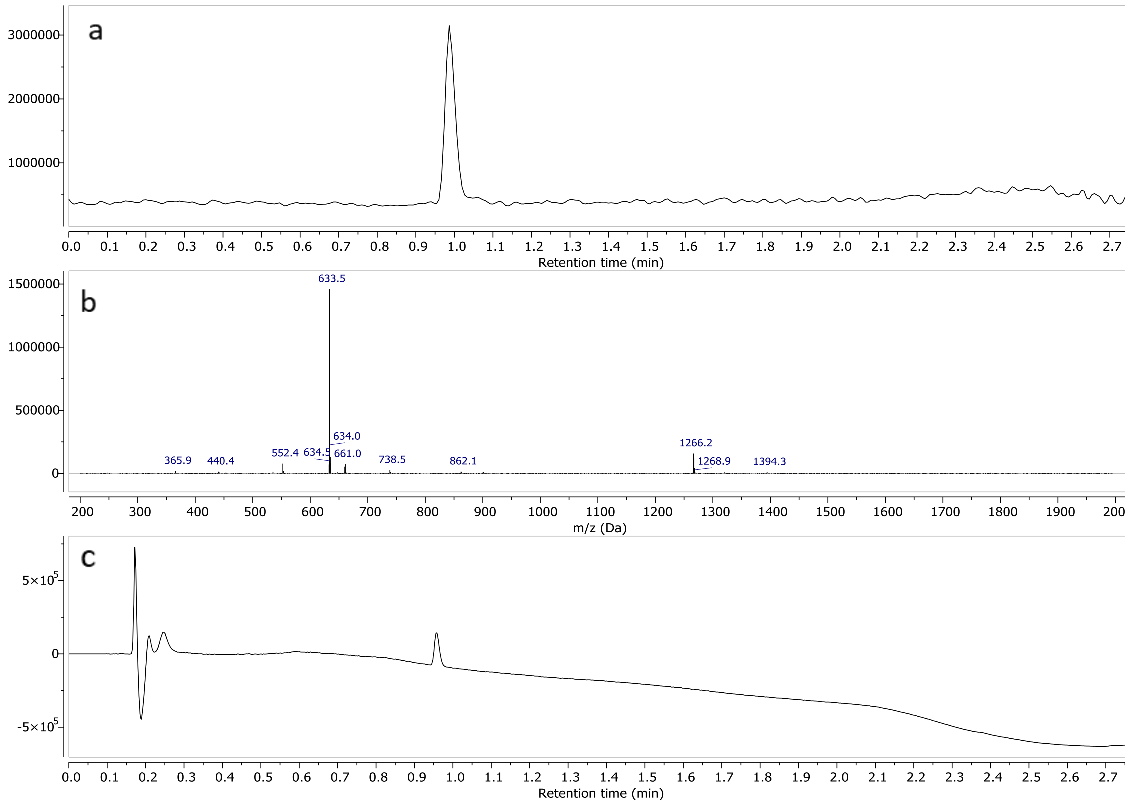


a) TIC – Total Ion Chromatogram (ESI+) b) mass spectrum of **4** (ESI+) c) UV spectrum at 190nm

#### Compound 5

##### [1]H-NMR ($D_2O$, 600MHz)

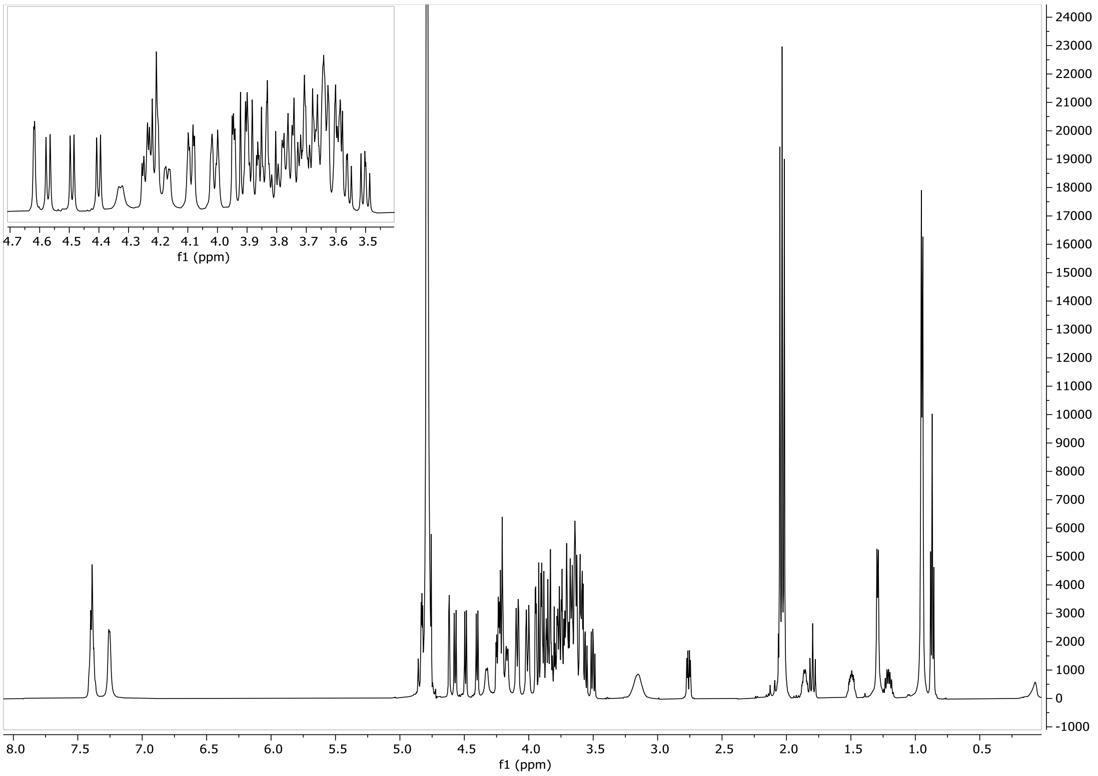

#### HSQC ($D_2O$, 600 MHz)

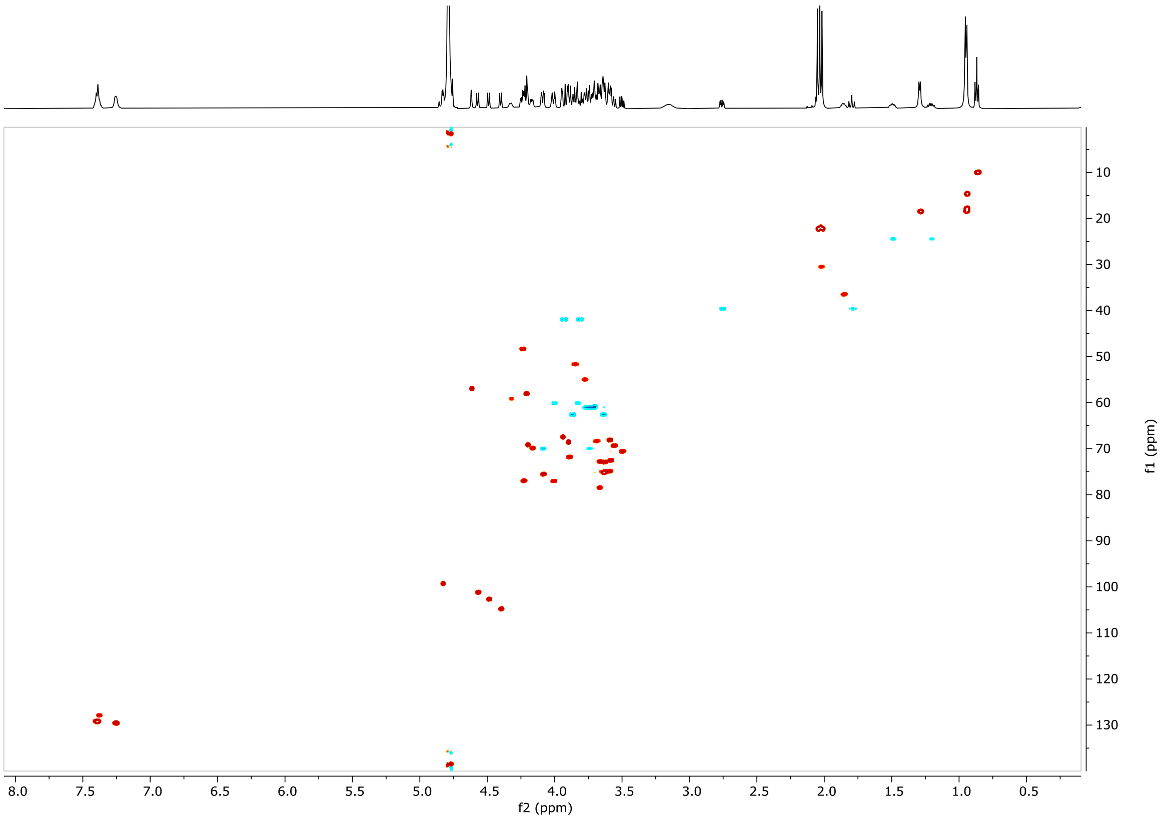


#### LCMS

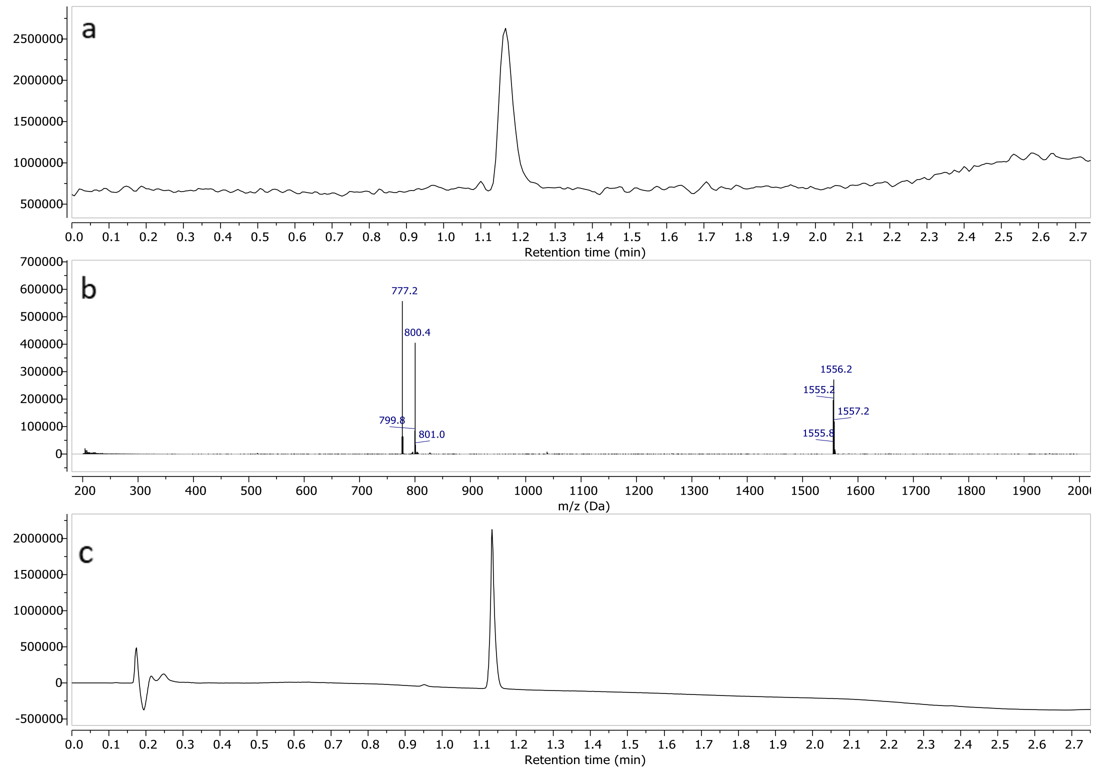


a) TIC – Total Ion Chromatogram (ESI+) b) mass spectrum of **5** (ESI+) c) UV spectrum at 190nm

#### Compound 6

##### $^1$H-NMR ($D_2O$, 600MHz)

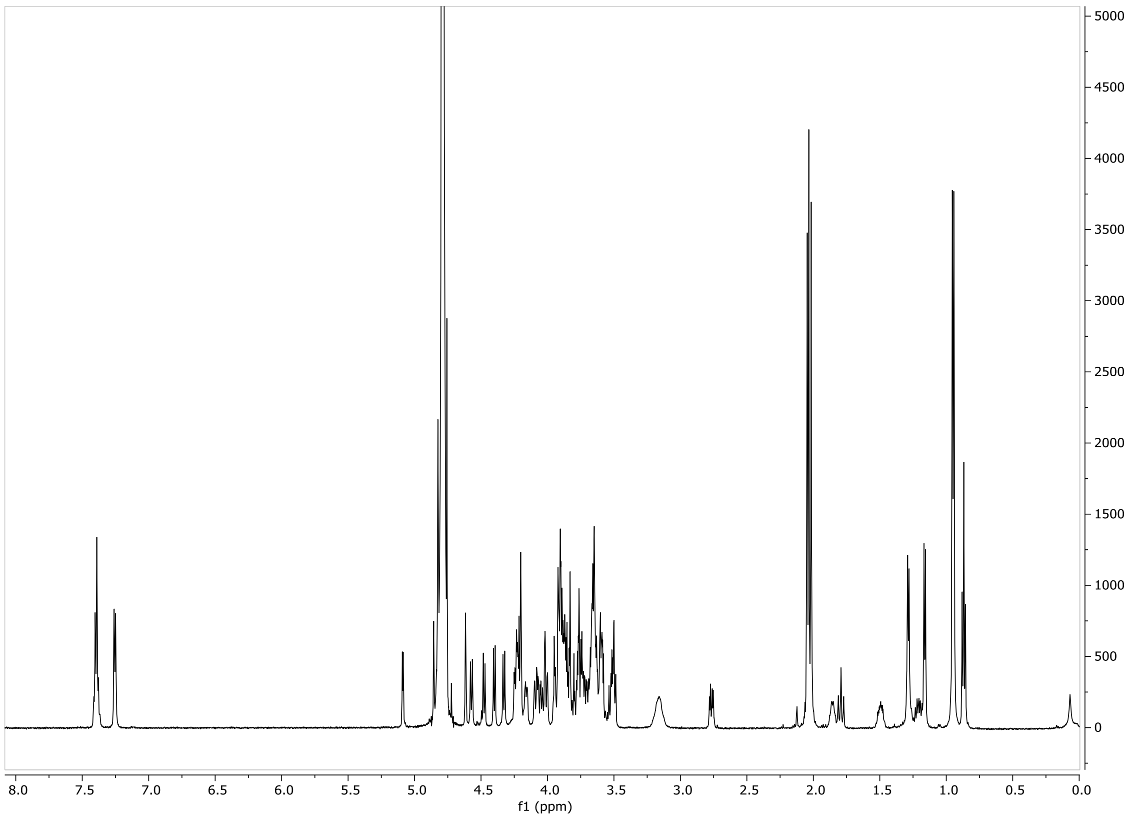


##### HSQC ($D_2O$, 600 MHz)

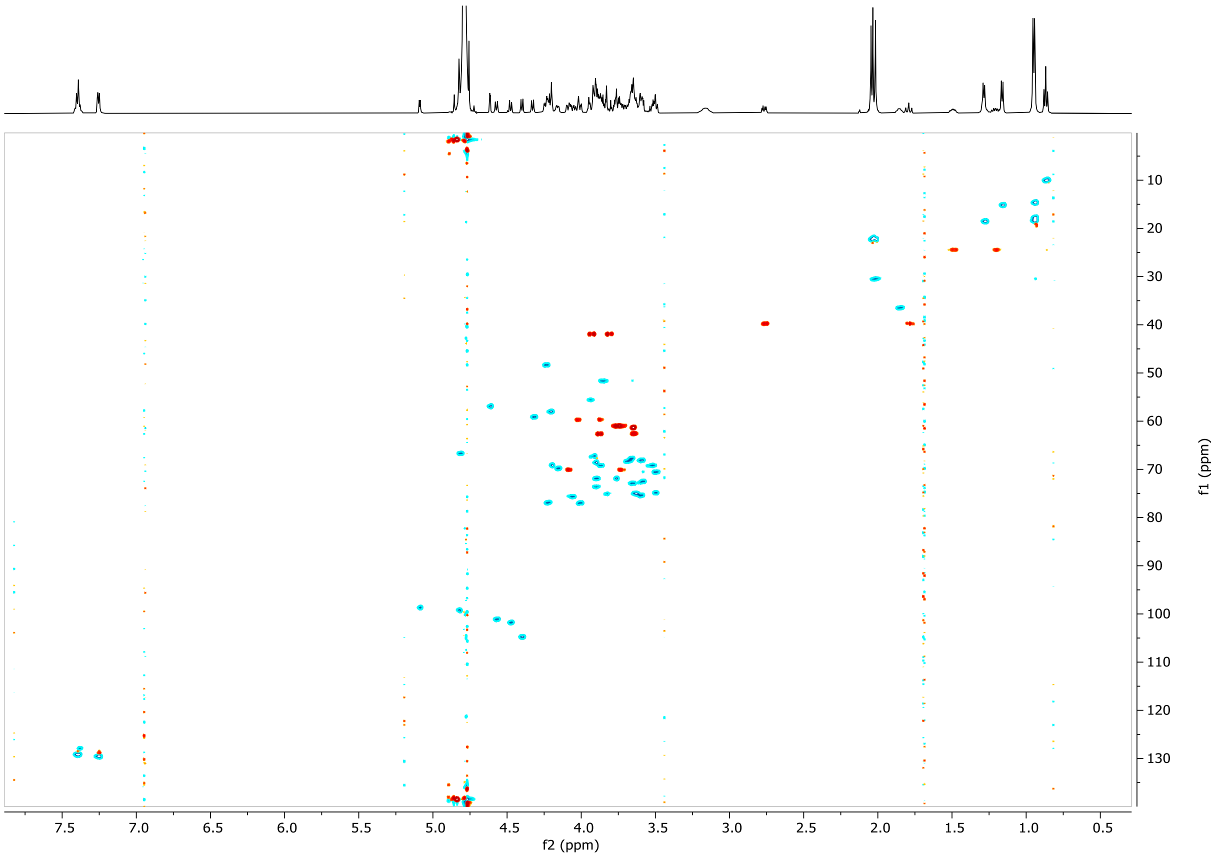

#### LCMS

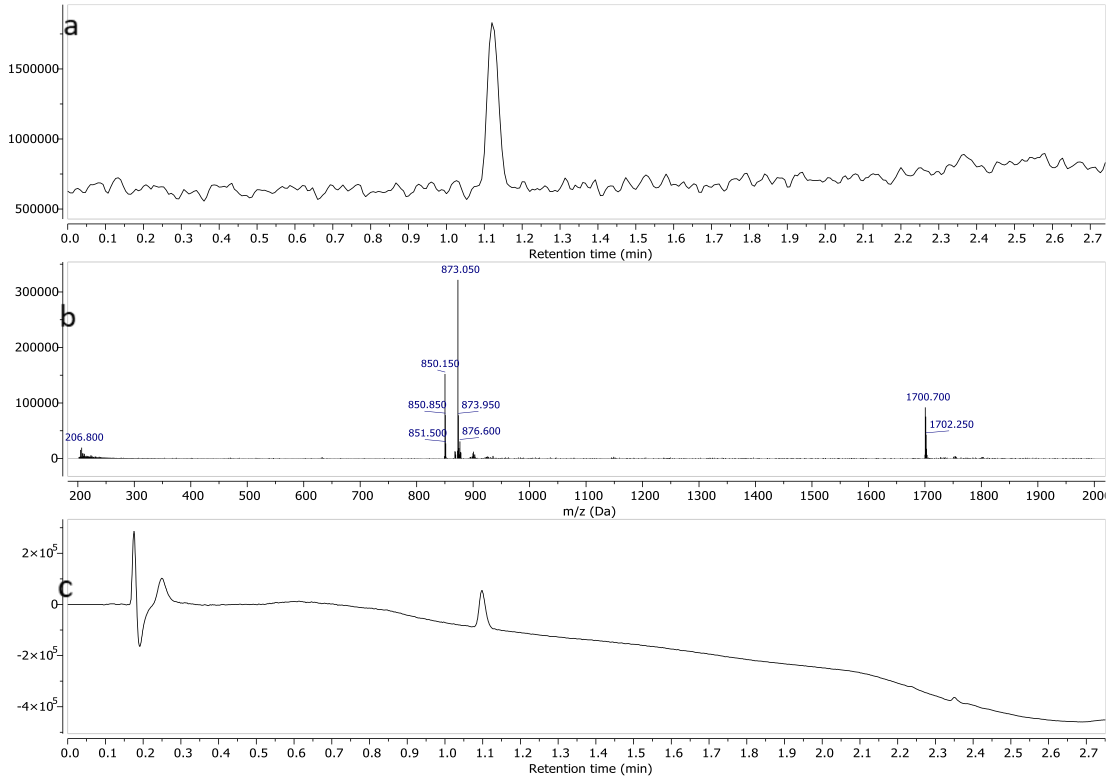


a) TIC – Total Ion Chromatogram (ESI-) b) mass spectrum of **6** (ESI-) c) UV spectrum at 190nm

#### Compound 7

#### $^{1}H$-NMR ($D_2O$, 600MHz)

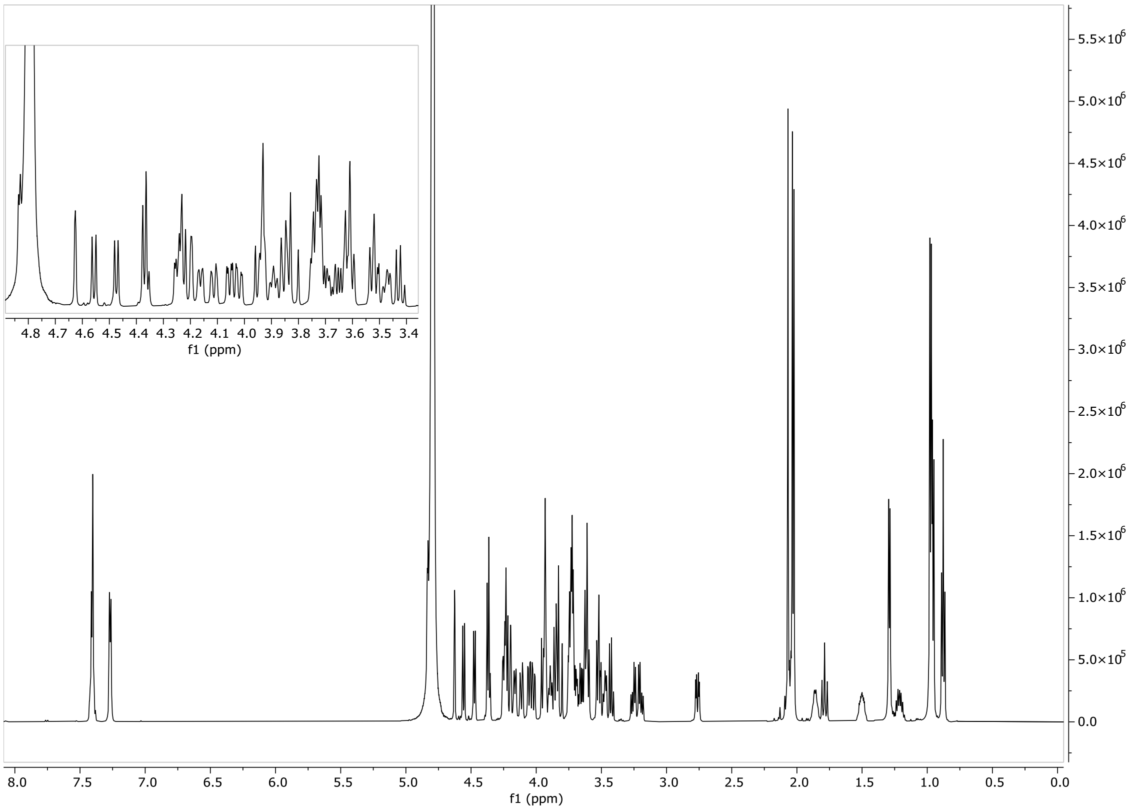

#### HSQC ($D_2O$, 600 MHz)

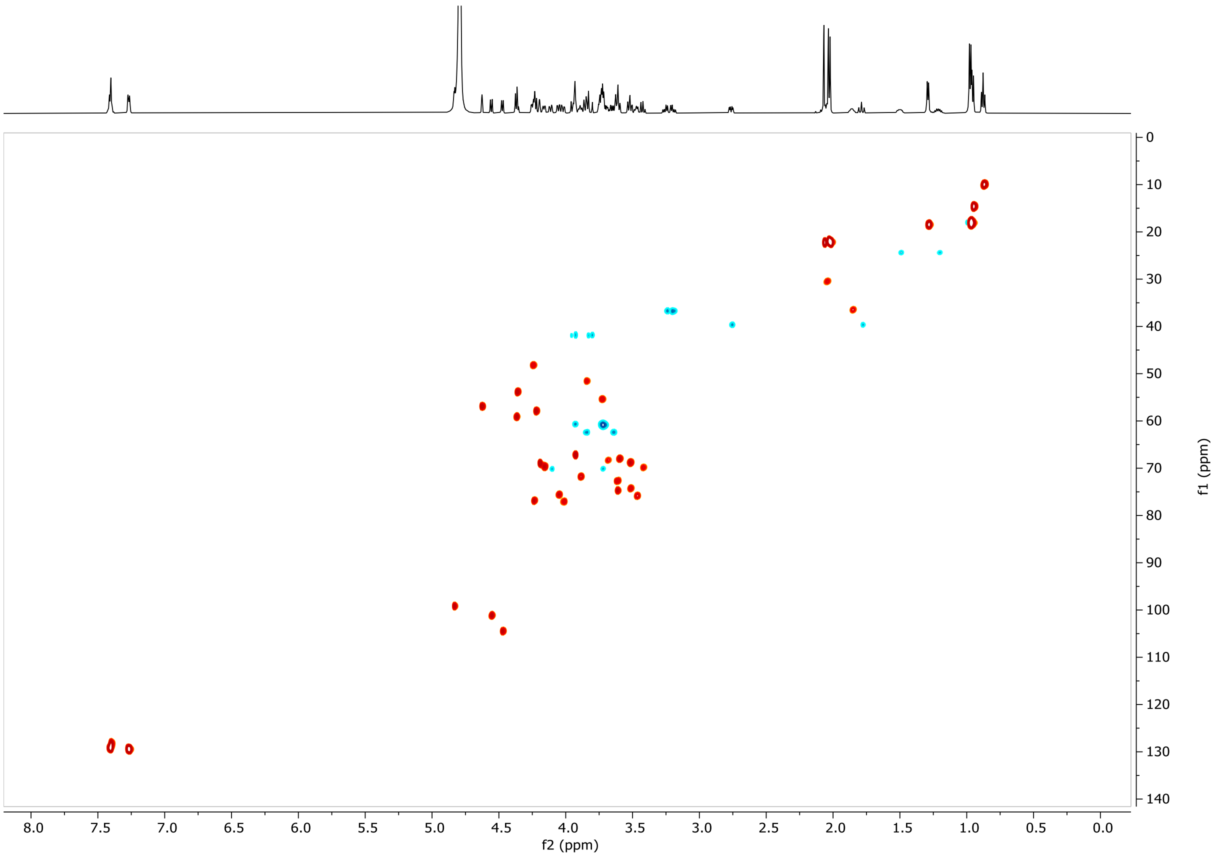


#### LCMS

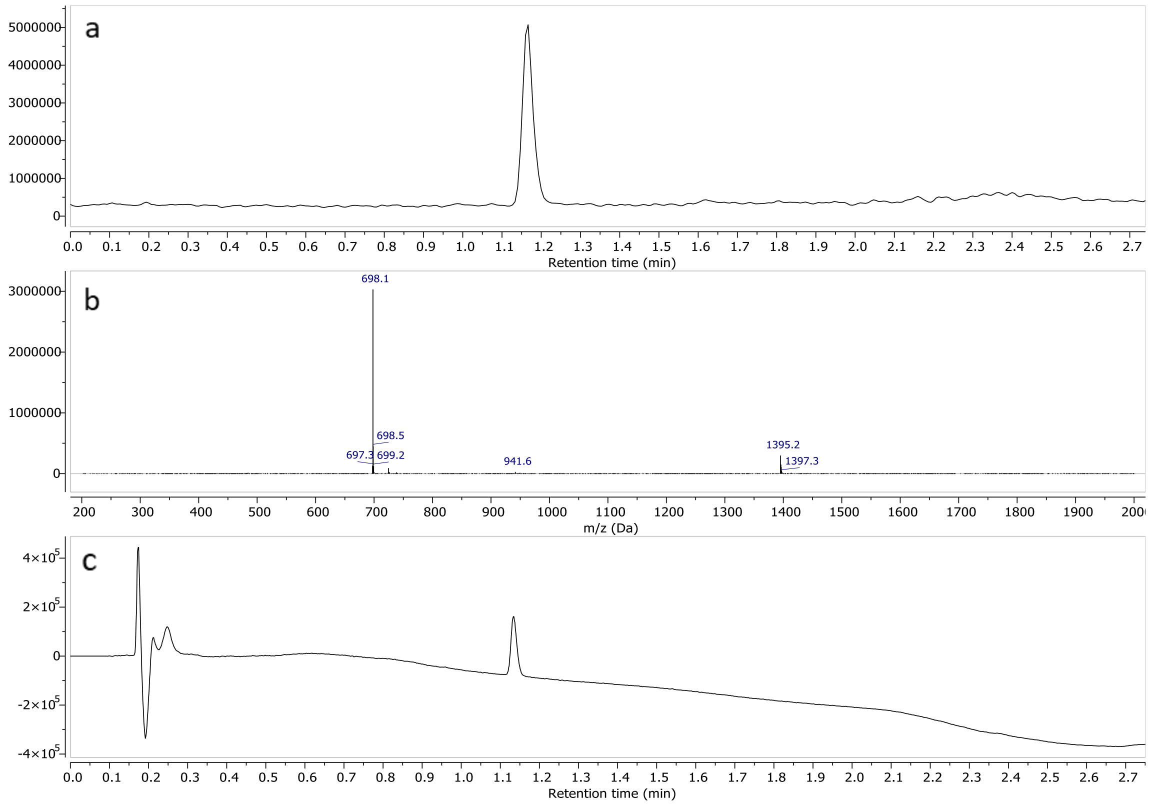


a) TIC – Total Ion Chromatogram (ESI+) b) mass spectrum of **7** (ESI+) c) UV spectrum at 190nm

#### Compound 8

##### $^1$H-NMR ($D_2O$, 600MHz)

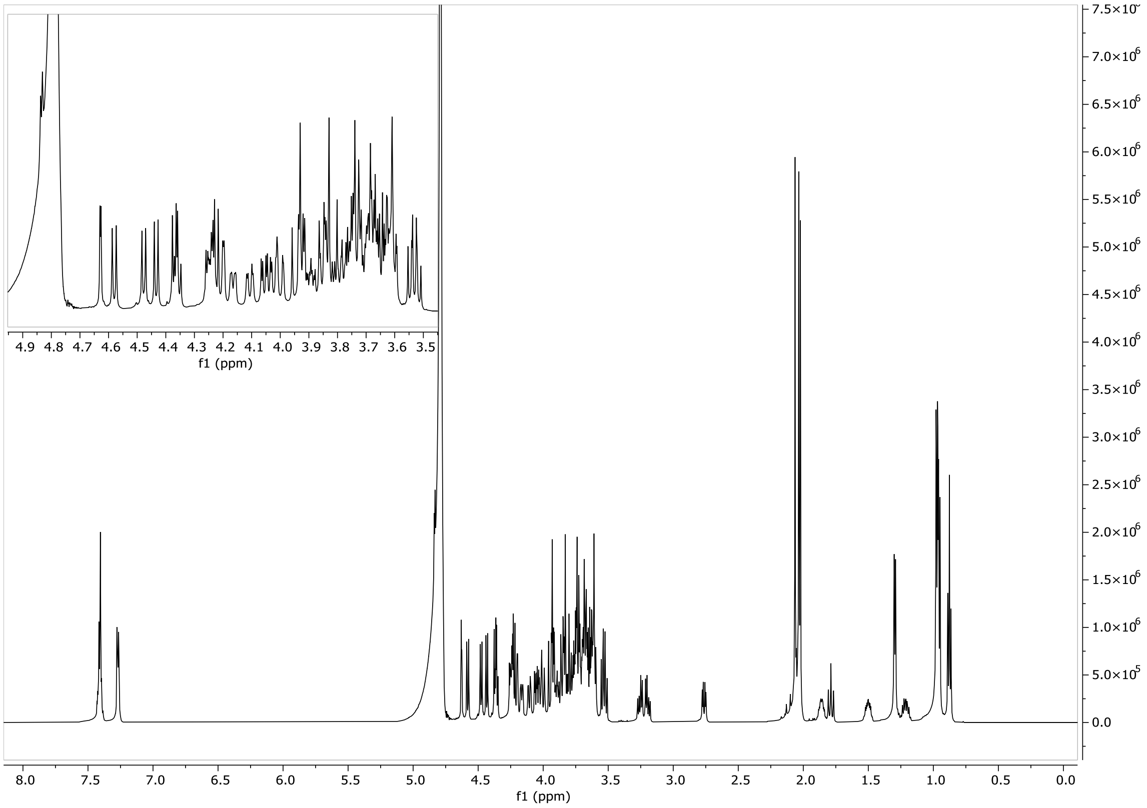


##### HSQC ($D_2O$, 600 MHz)

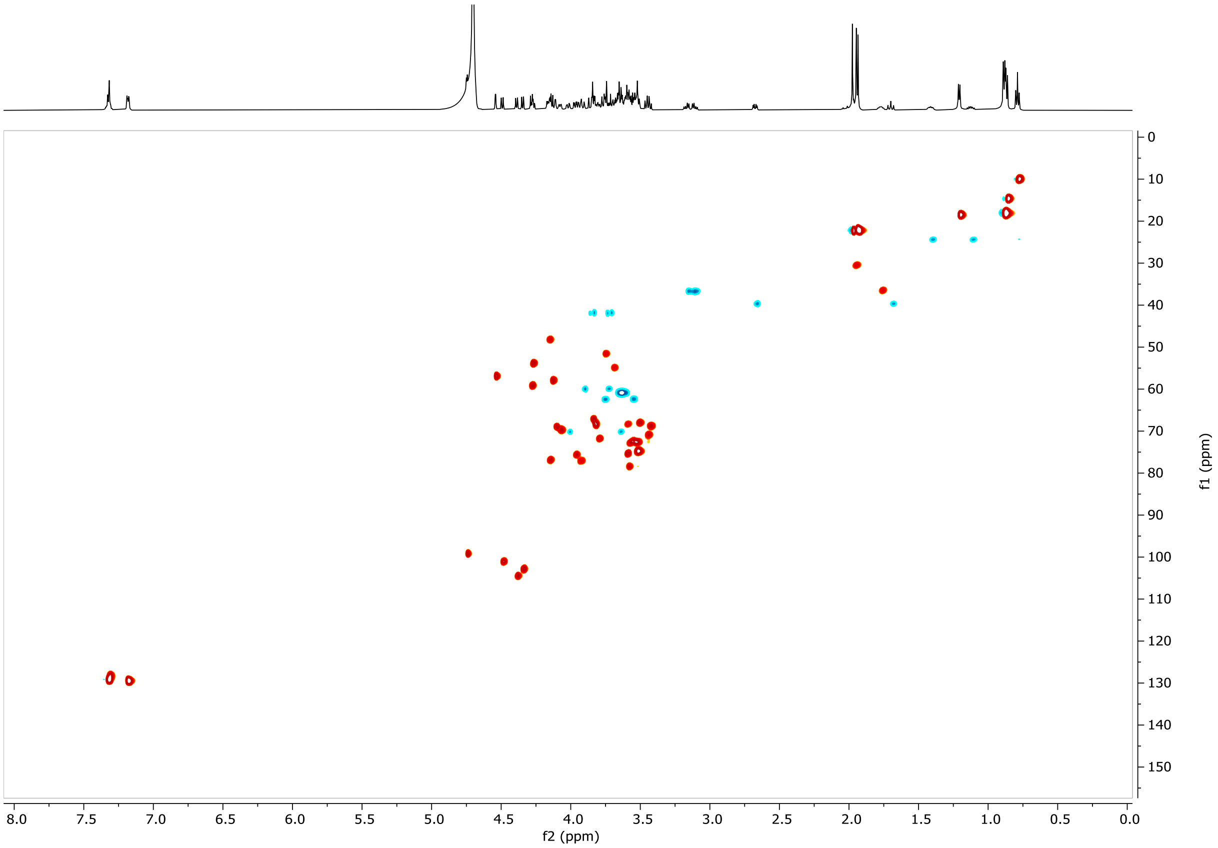

#### LCMS

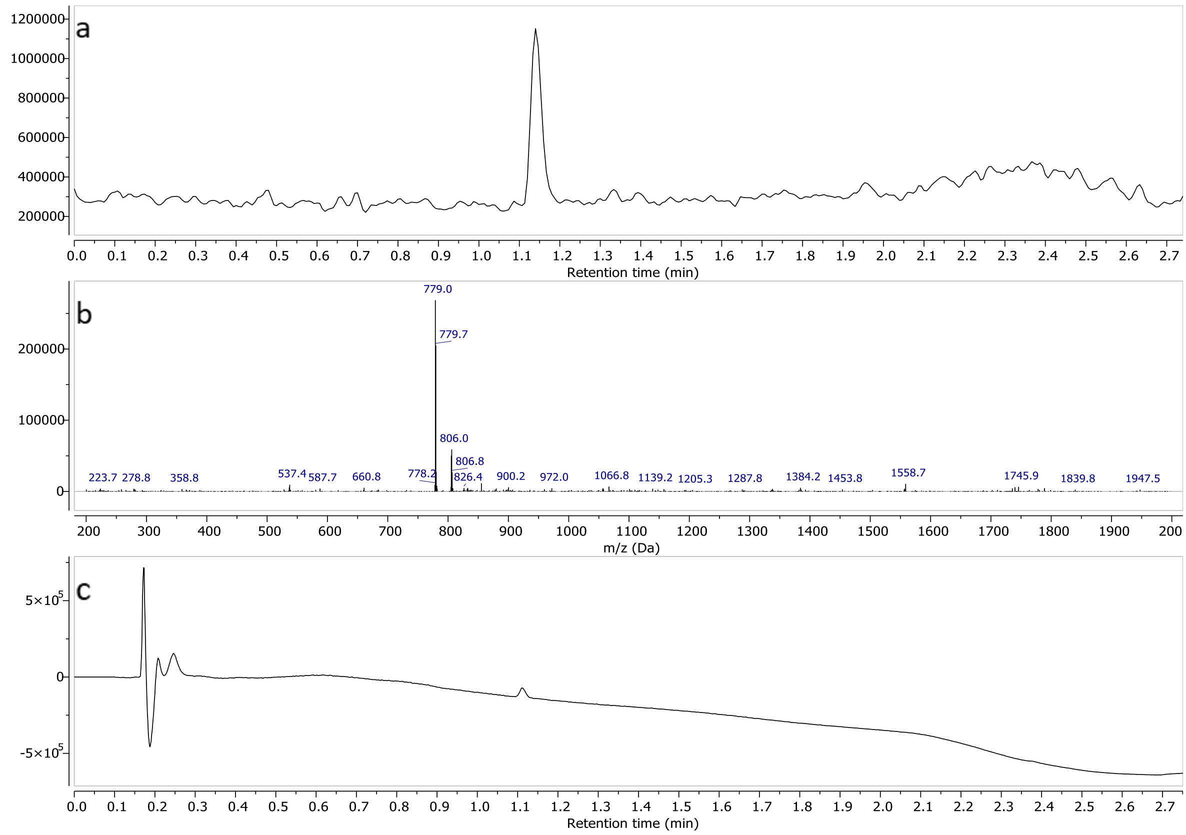


a) TIC – Total Ion Chromatogram (ESI+) b) mass spectrum of **8** (ESI+)c) UV spectrum at 190nm

#### Compound 9

#### $^1$H-NMR ($D_2O$, 600MHz)

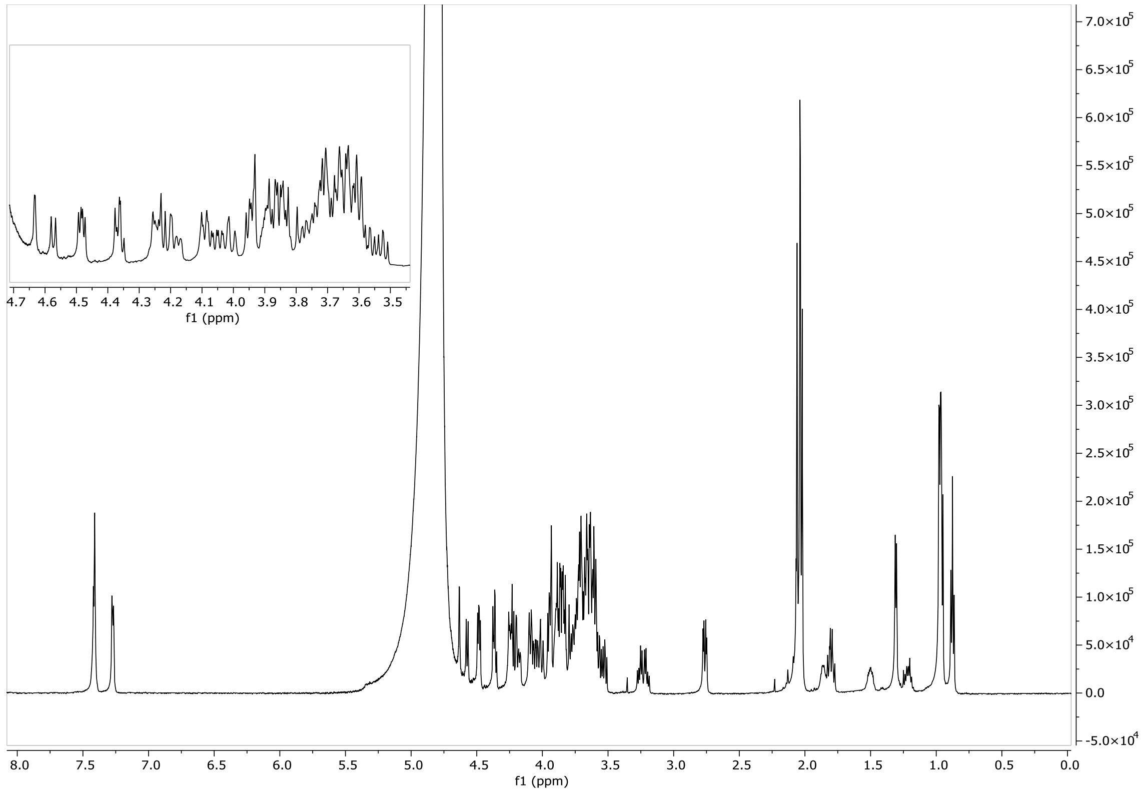

#### HSQC ($D_2O$, 600 MHz)

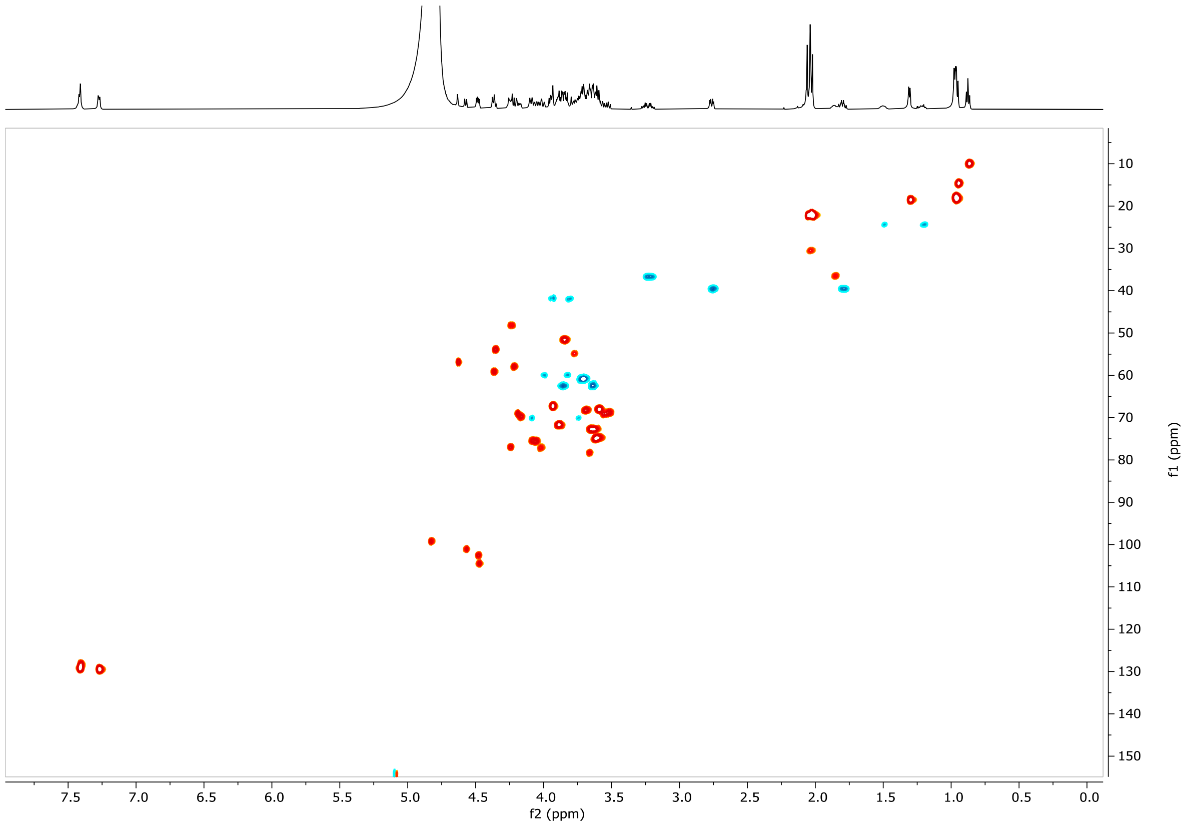


#### LCMS

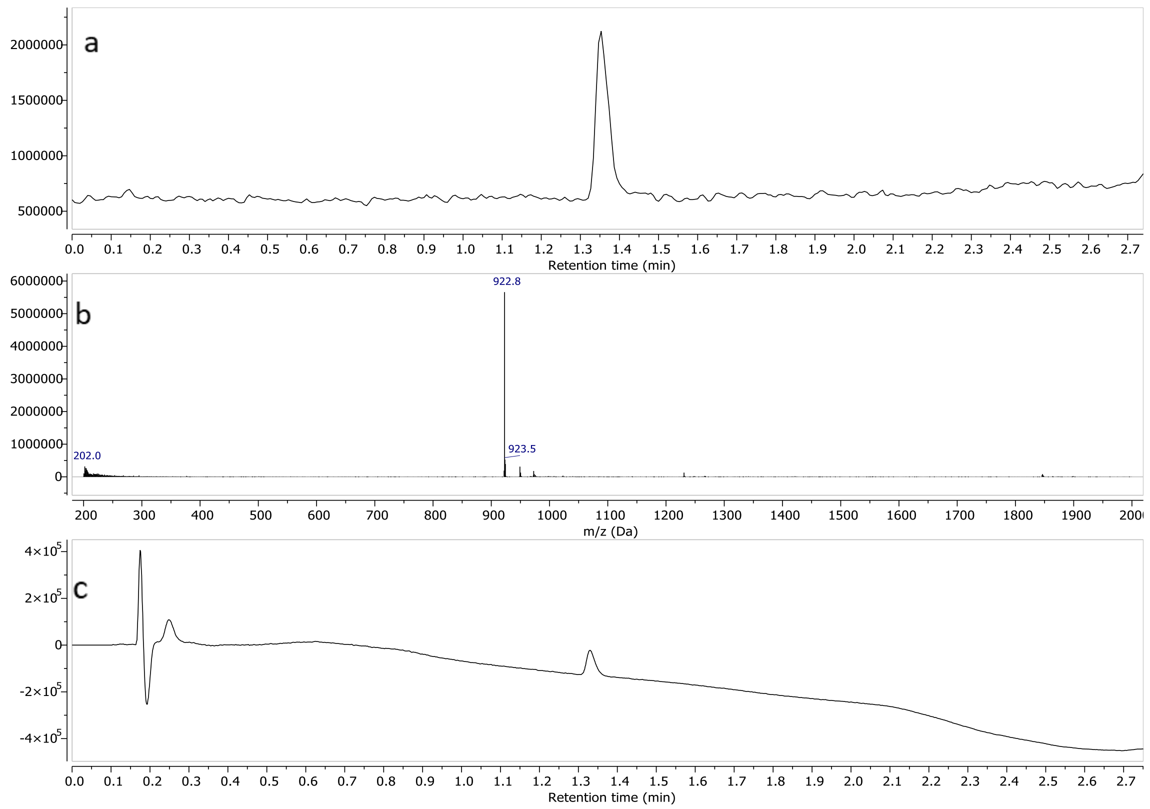


a) TIC – Total Ion Chromatogram (ESI-) b) mass spectrum of **9** (ESI-) c) UV spectrum at 190nm

#### Compound 10

#### $^1$H-NMR ($D_2O$, 600MHz)

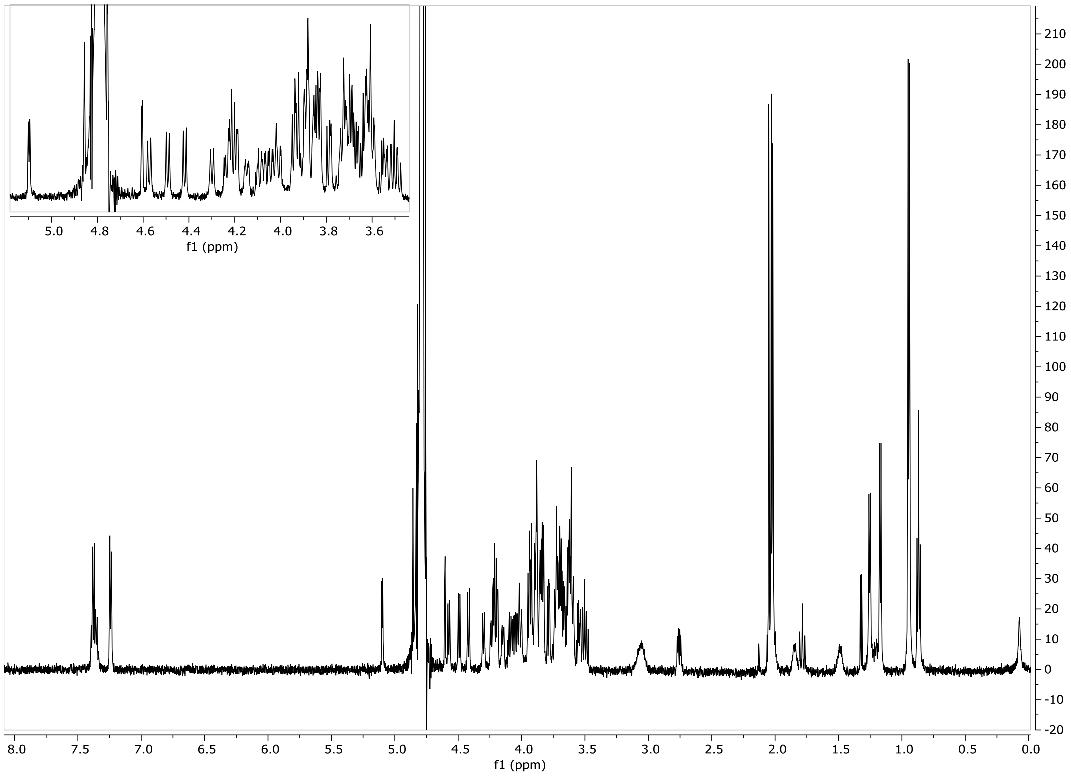


#### HSQC ($D_2O$, 600 MHz)

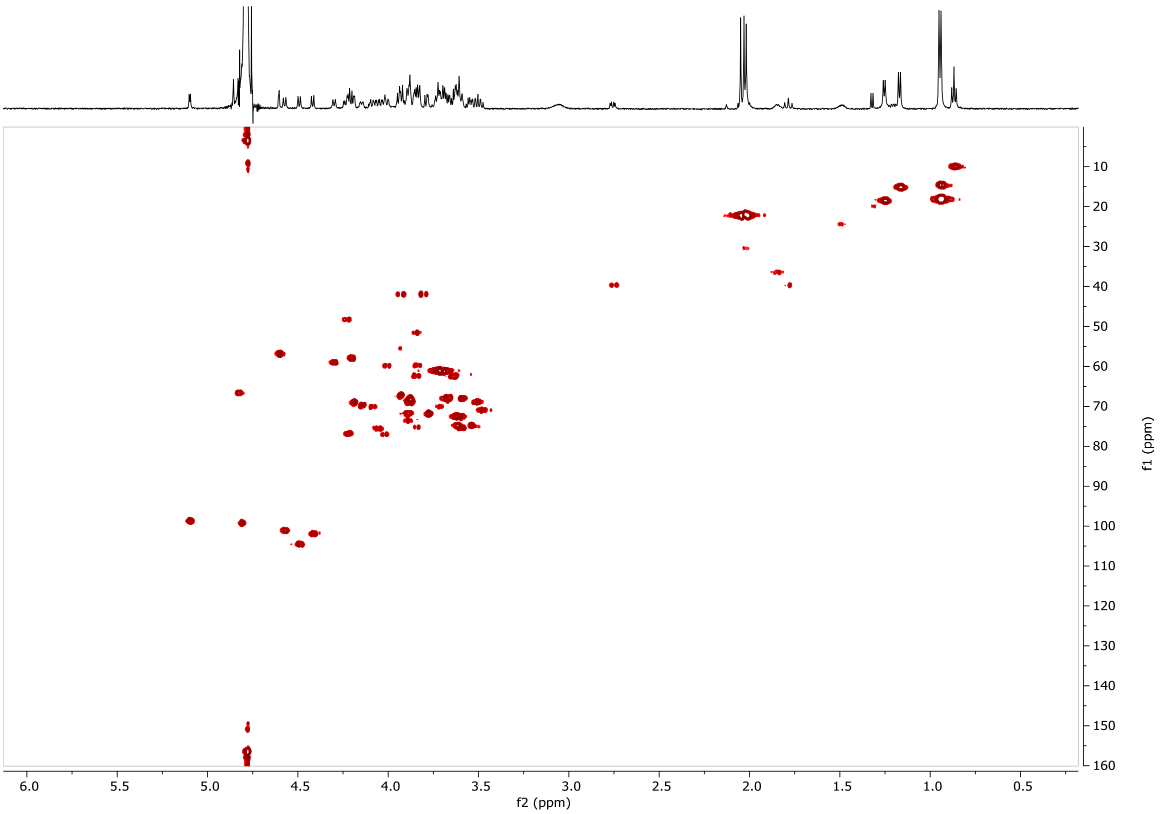


#### LCMS

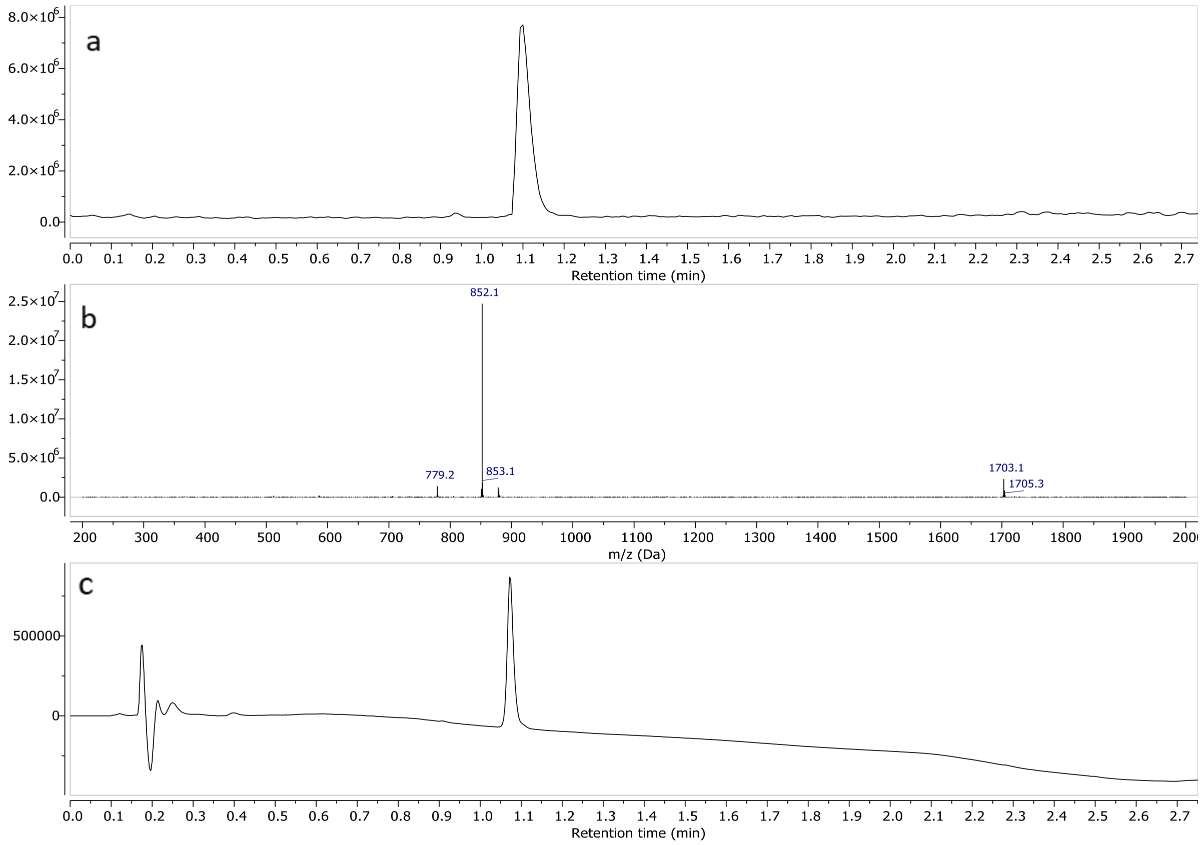


a) TIC – Total Ion Chromatogram (ESI+) b) mass spectrum of **10** (ESI+) c) UV spectrum at 190nm

#### Compound 11

##### $^1$H-NMR ($D_2O$, 600MHz)

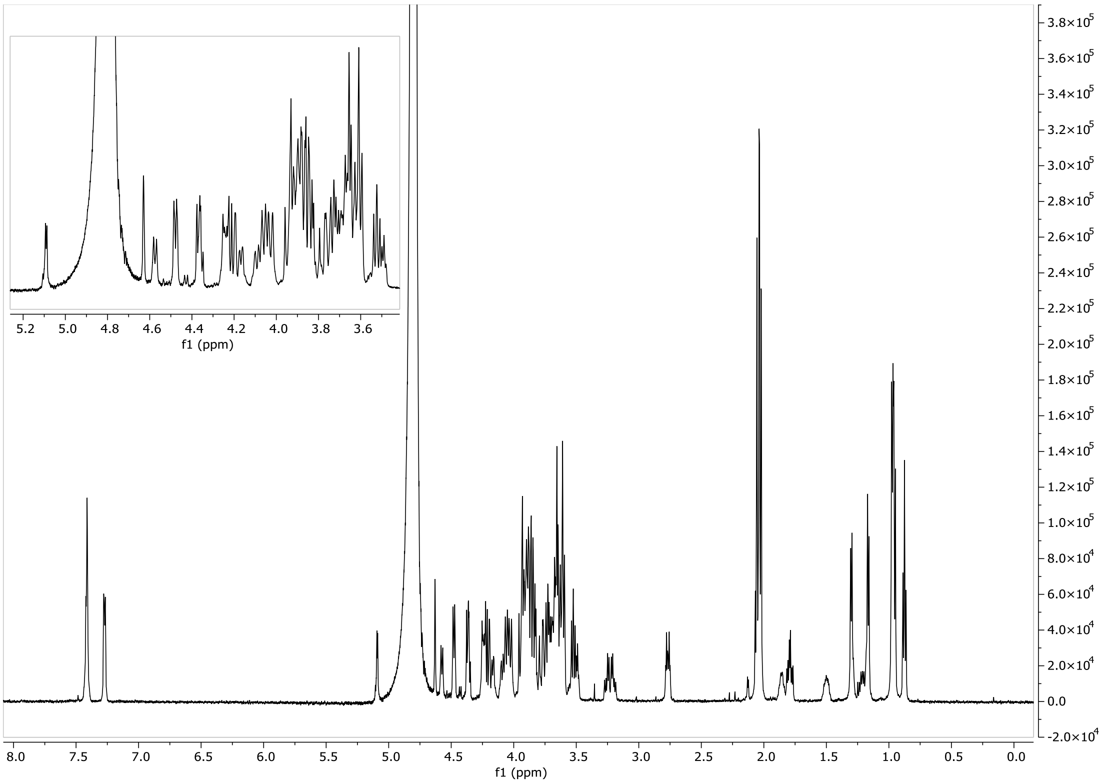

#### HSQC ($D_2O$, 600 MHz)

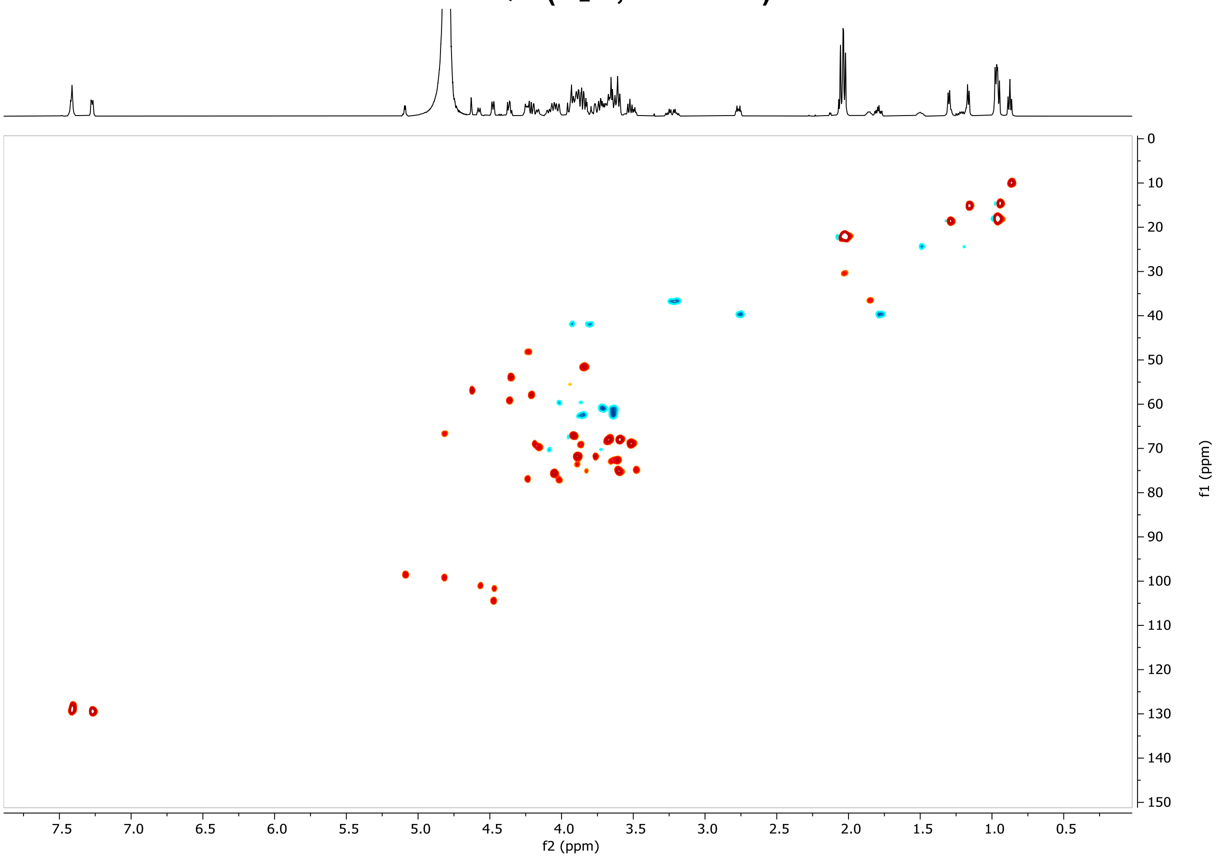


#### LCMS

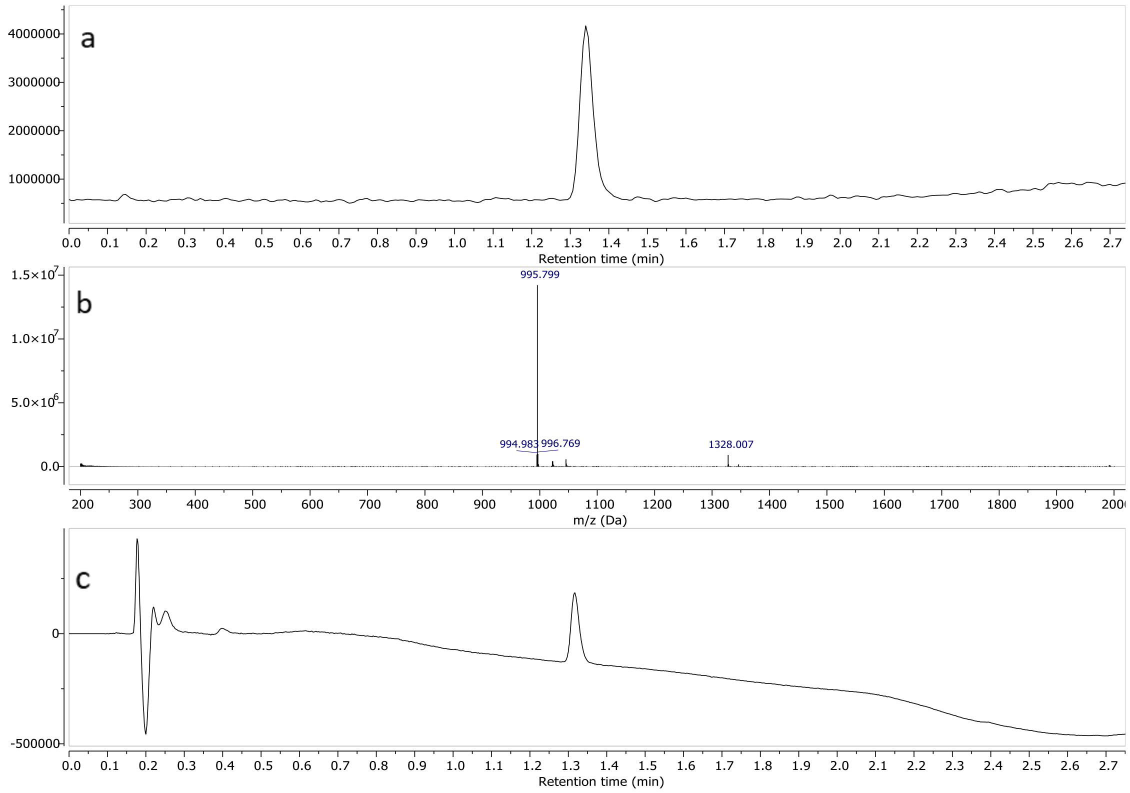


a) TIC – Total Ion Chromatogram (ESI-) b) mass spectrum of **11** (ESI-) c) UV spectrum at 190nm

#### Compound 12

#### $^{1}$H-NMR ($D_2O$, 600MHz)

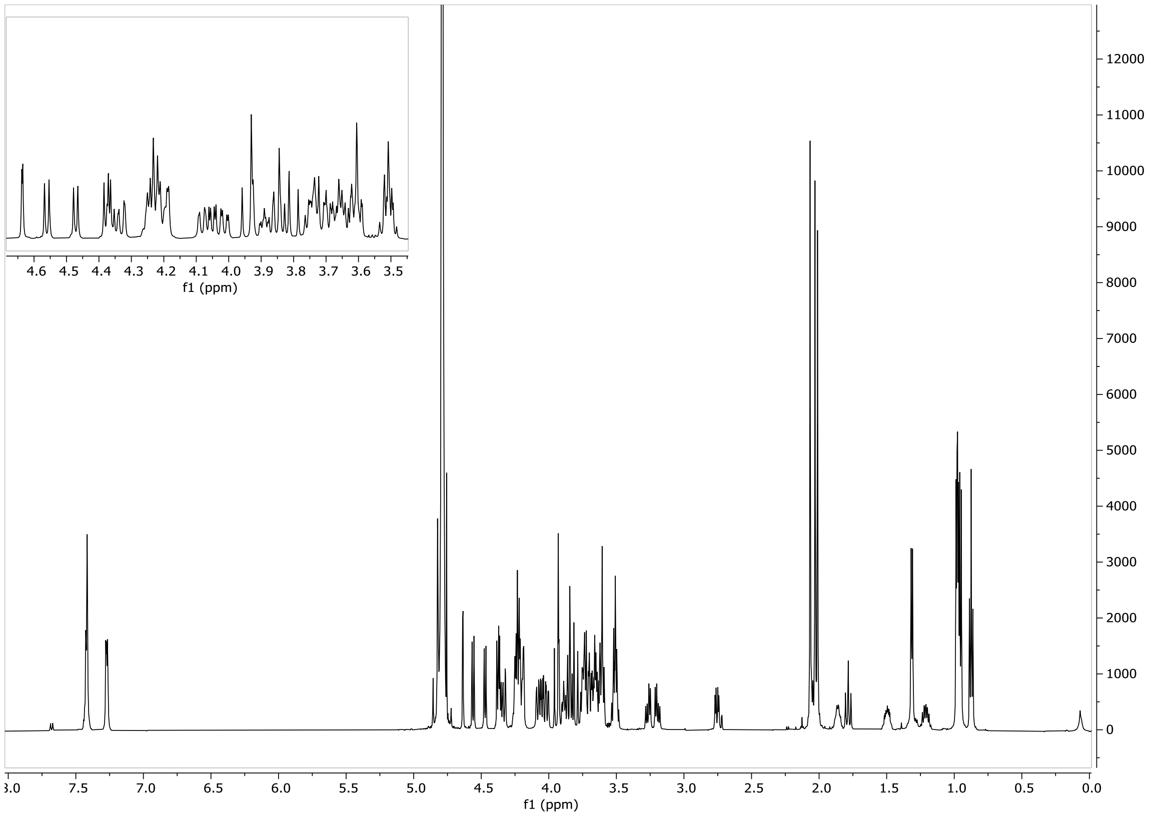


**HSQC ($D_2O$, 600 MHz)-** HSQC contains 2 impurity dots at [4,7;75,0] ppm and [3,56;62,5] ppm which could not be ascribed to a known impurity.

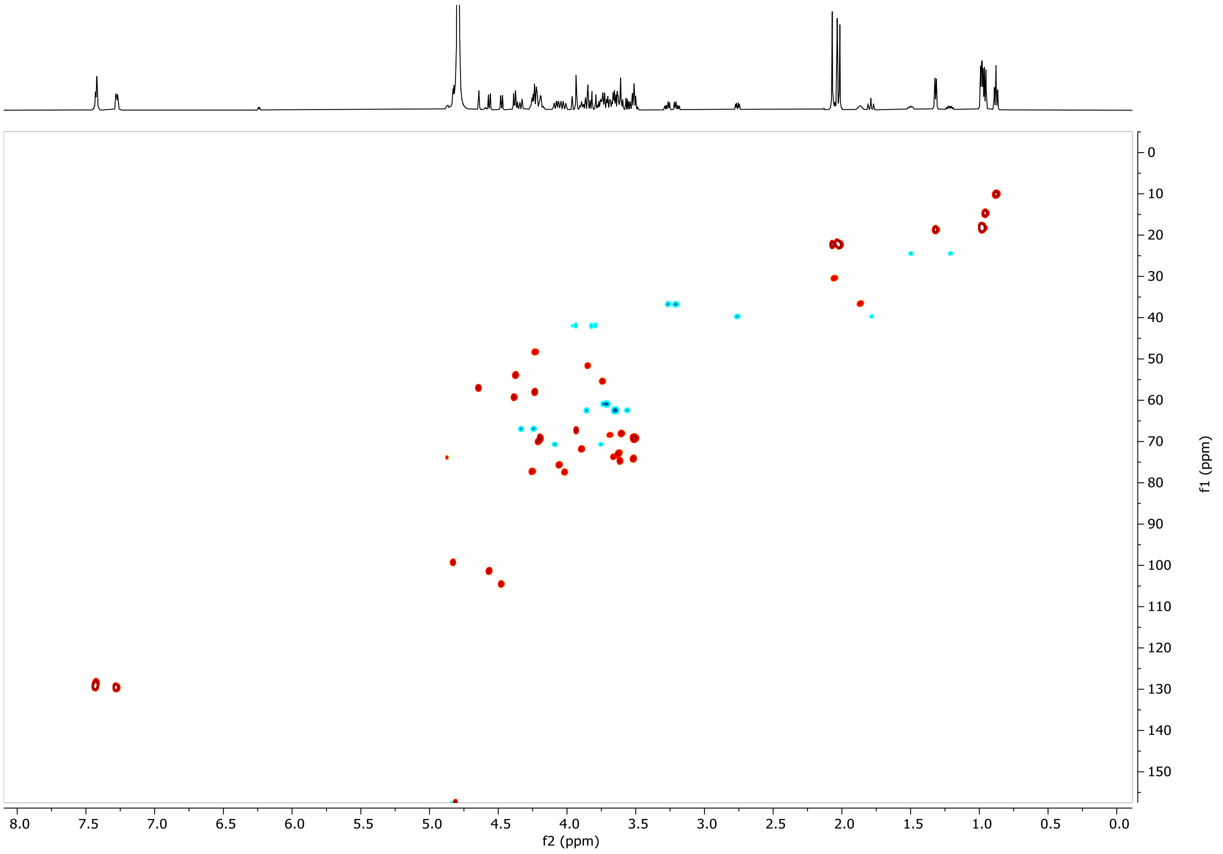

#### LCMS

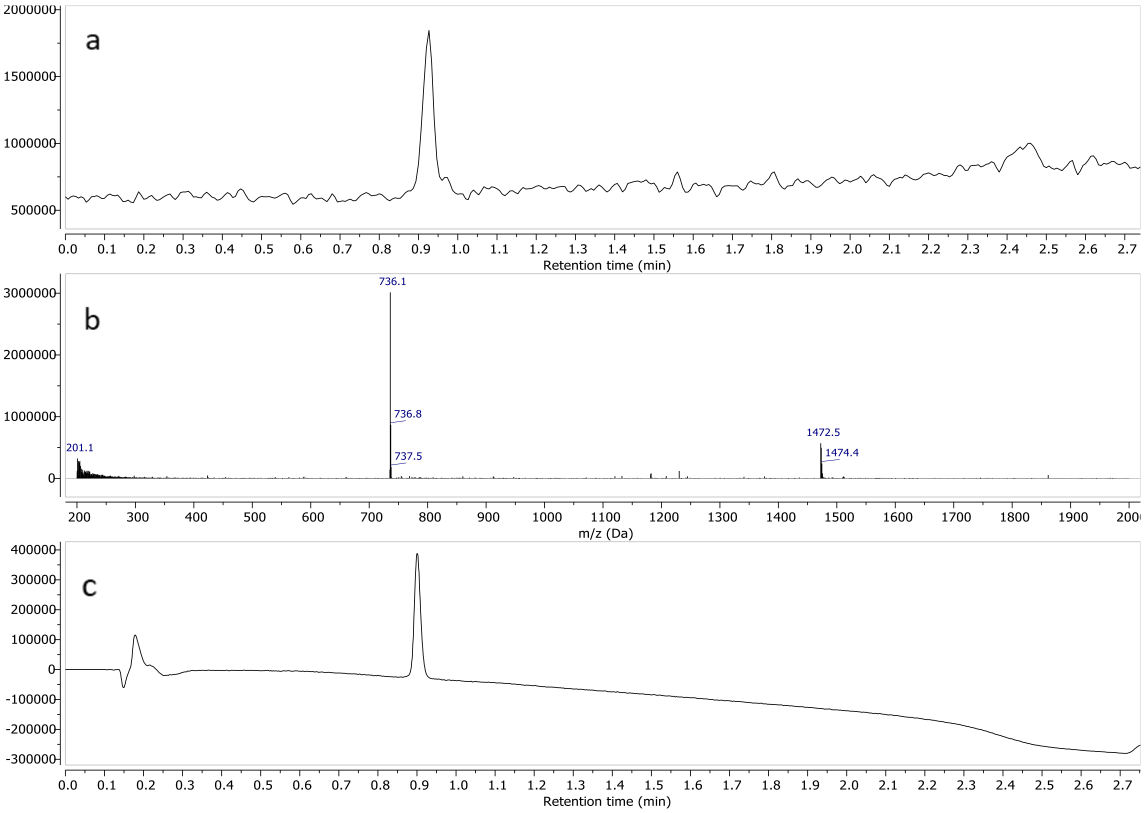


a) TIC – Total Ion Chromatogram (ESI-) b) mass spectrum of **12** (ESI-) c) UV spectrum at 190nm

#### Compound 13

#### $^{1}$H-NMR ($D_2O$, 600MHz)

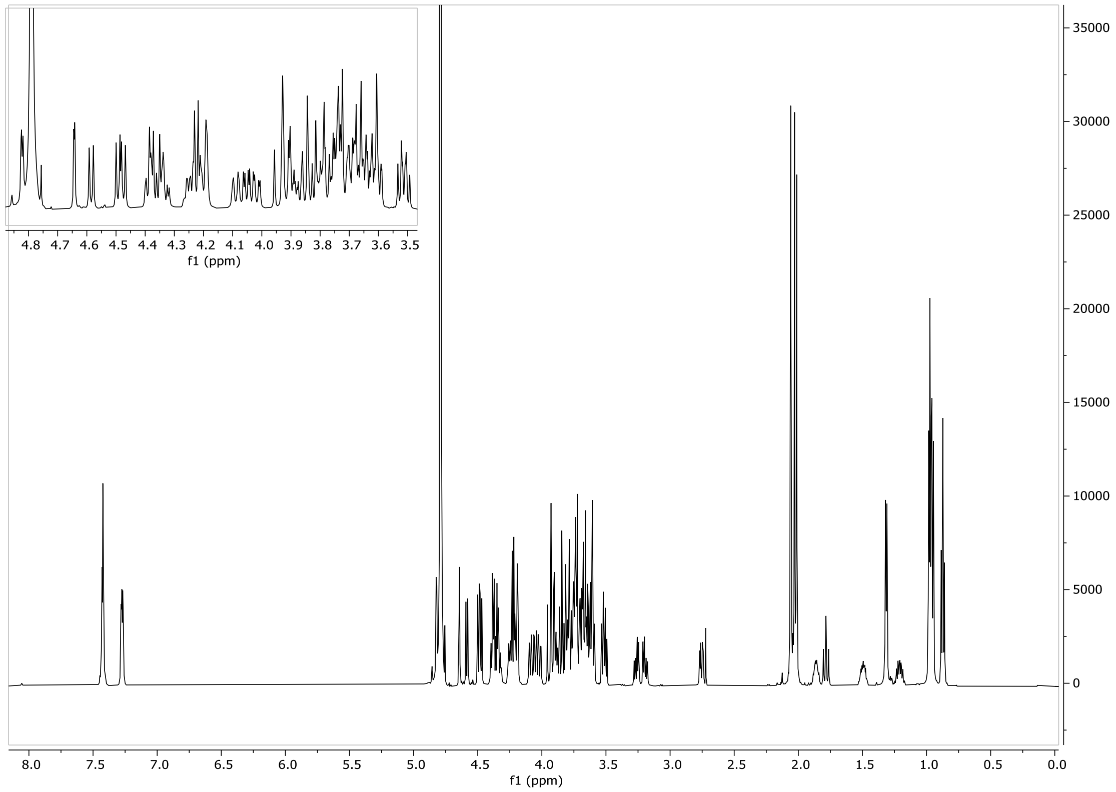

#### HSQC ($D_2O$, 600 MHz)

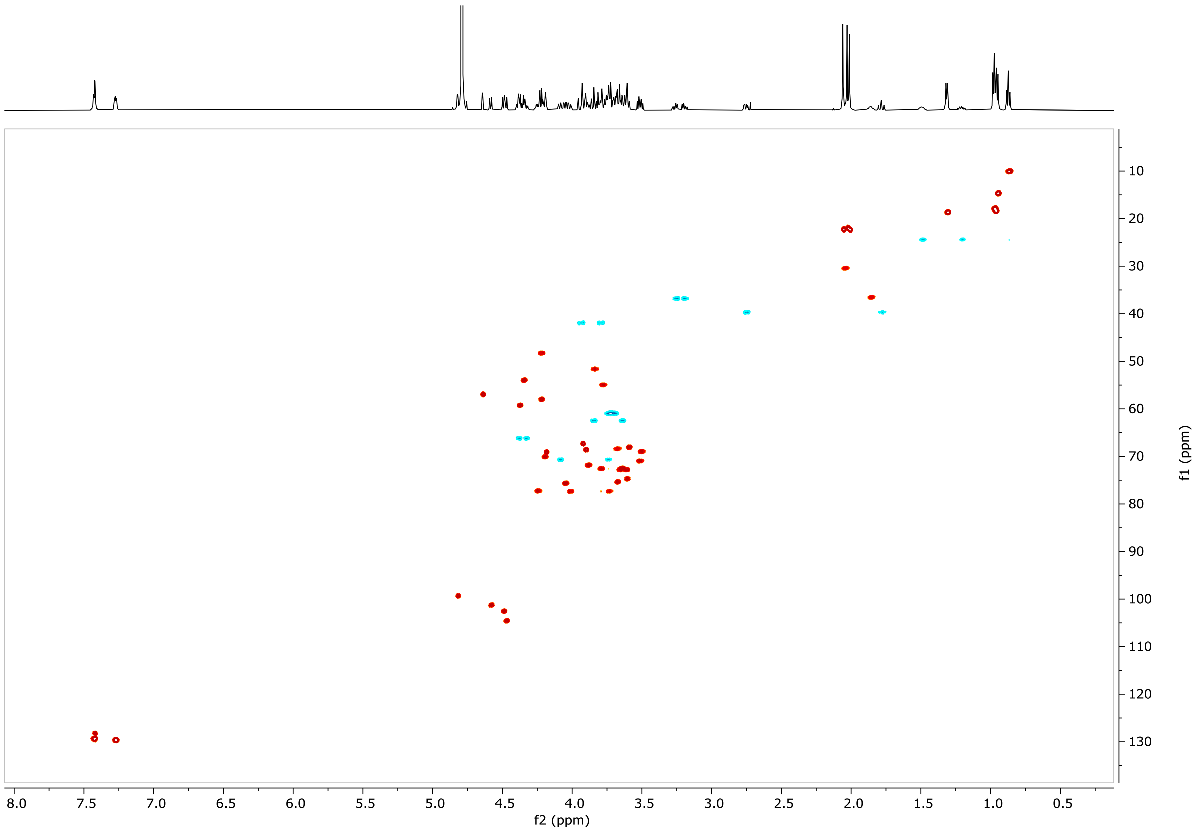


#### LCMS

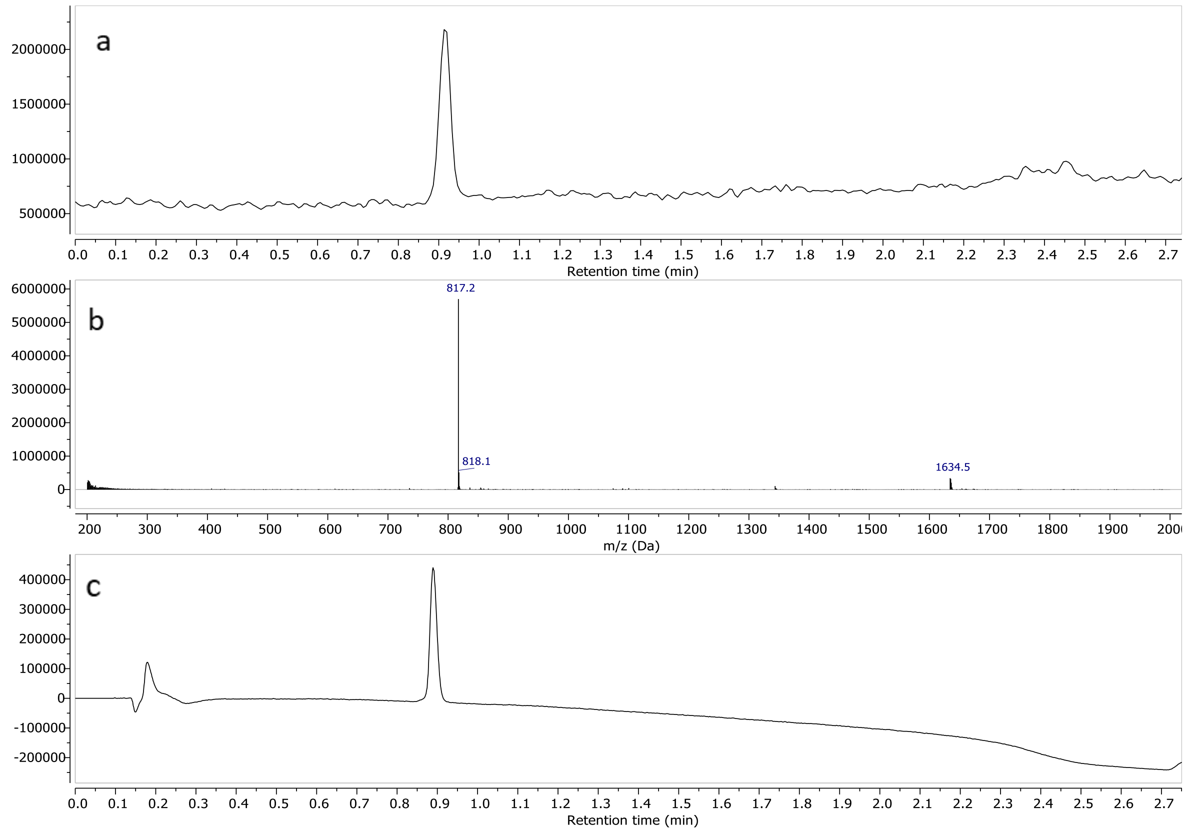


a) TIC – Total Ion Chromatogram (ESI-) b) mass spectrum of **13** (ESI-) c) UV spectrum at 190nm

#### Compound 14

#### $^1$H-NMR ($D_2O$, 600MHz)

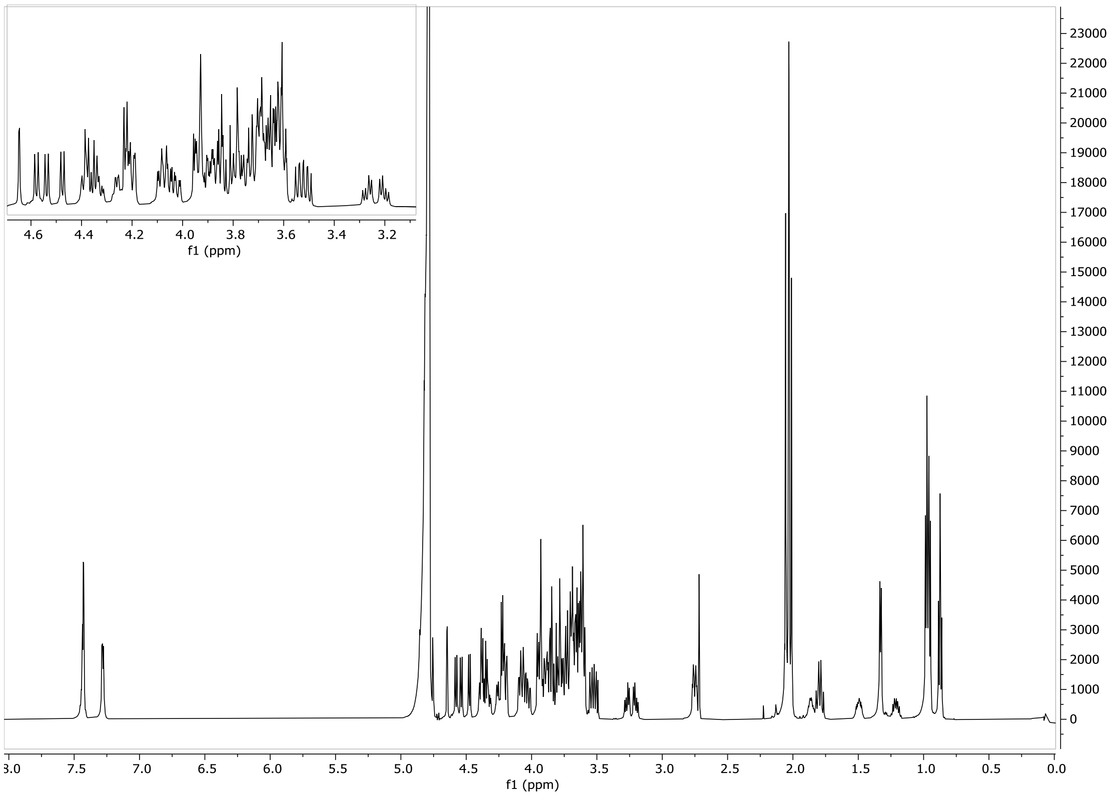


#### HSQC ($D_2O$, 600 MHz)

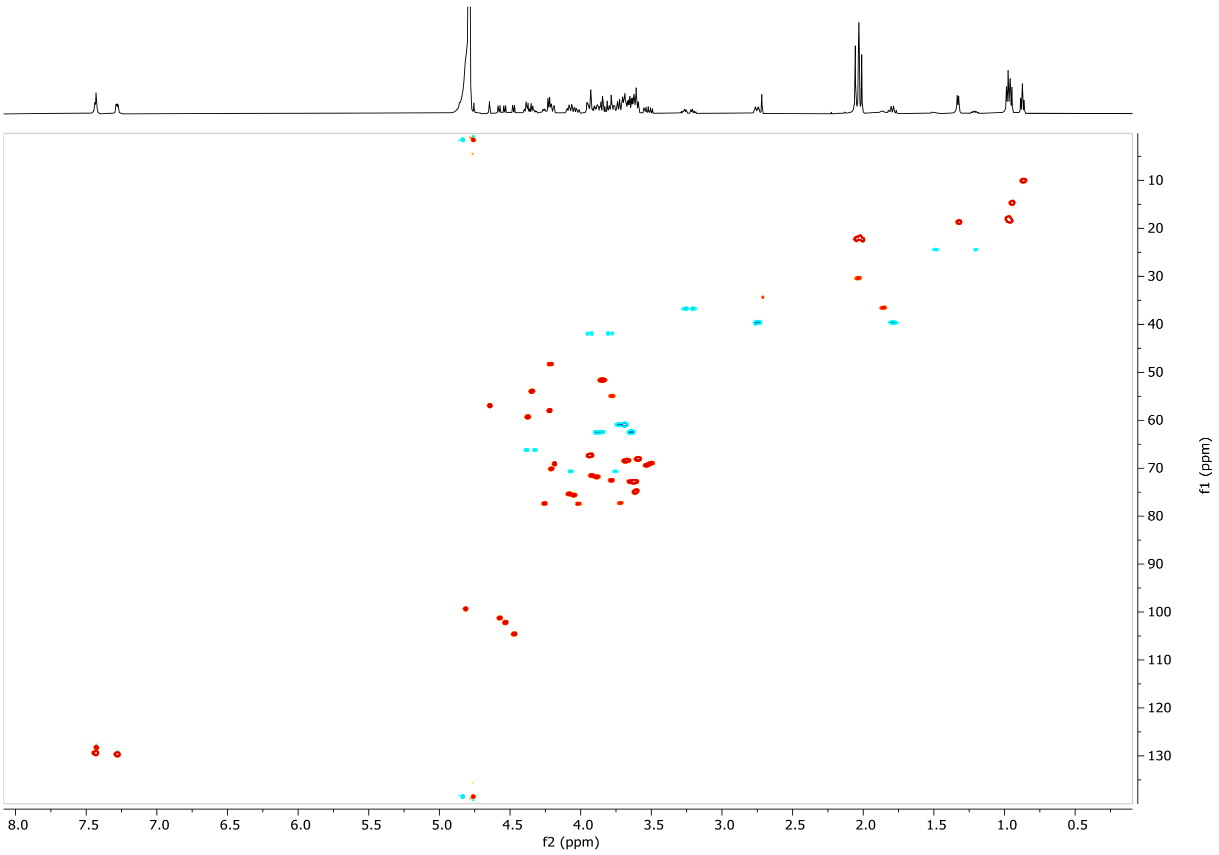

#### LCMS

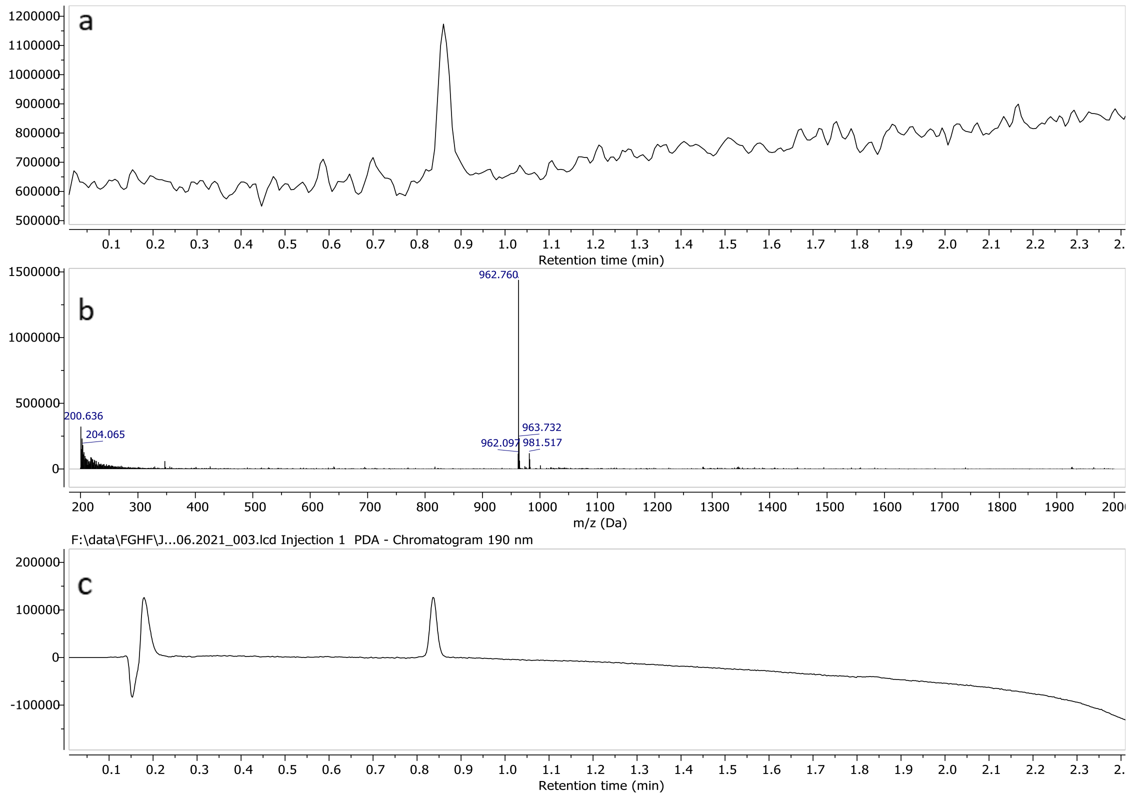


a) TIC – Total Ion Chromatogram (ESI-) b) mass spectrum of **14** ESI-) c) UV spectrum at 190nm

#### Compound 15

#### $^1$H-NMR ($D_2O$, 600MHz)

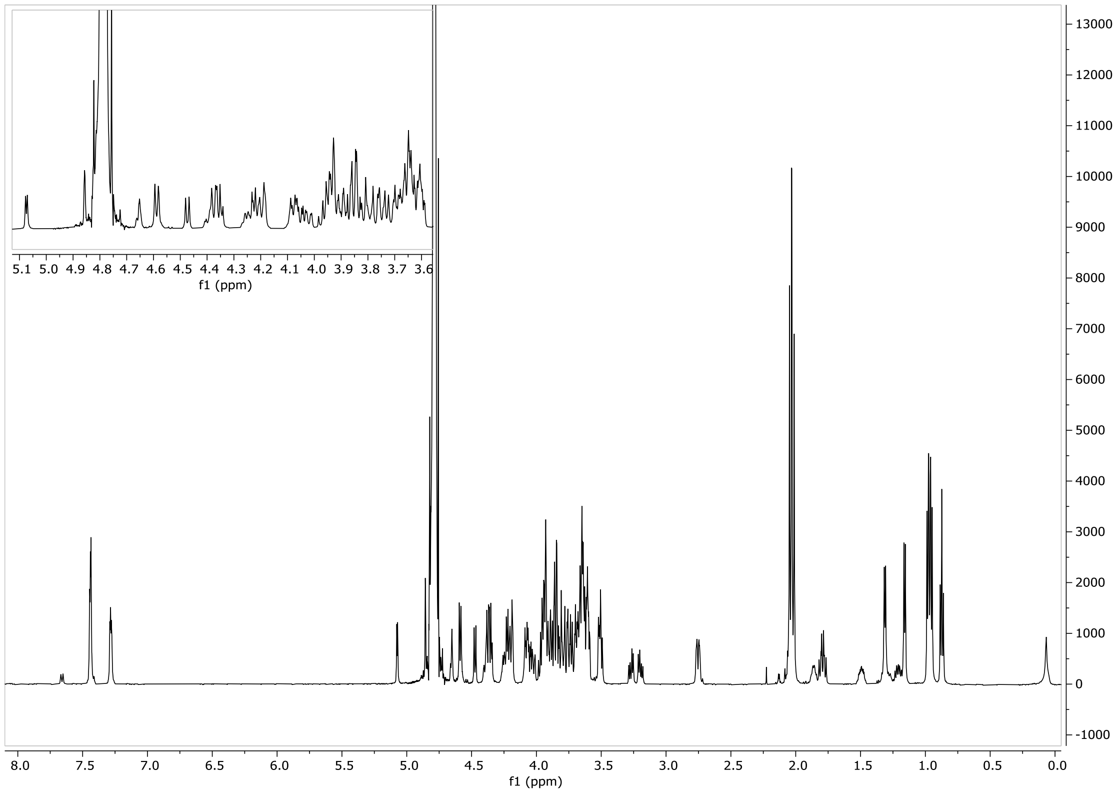

#### HSQC ($D_2O$, 600 MHz)

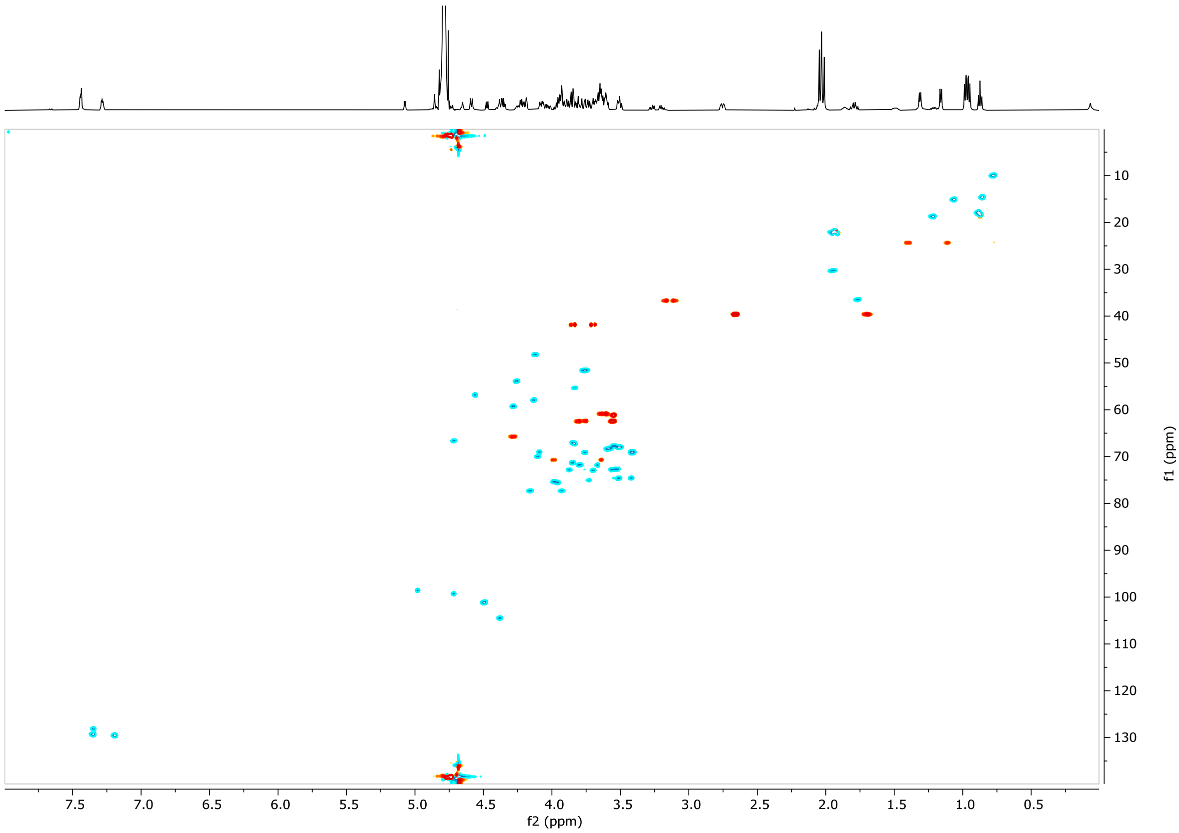


#### LCMS

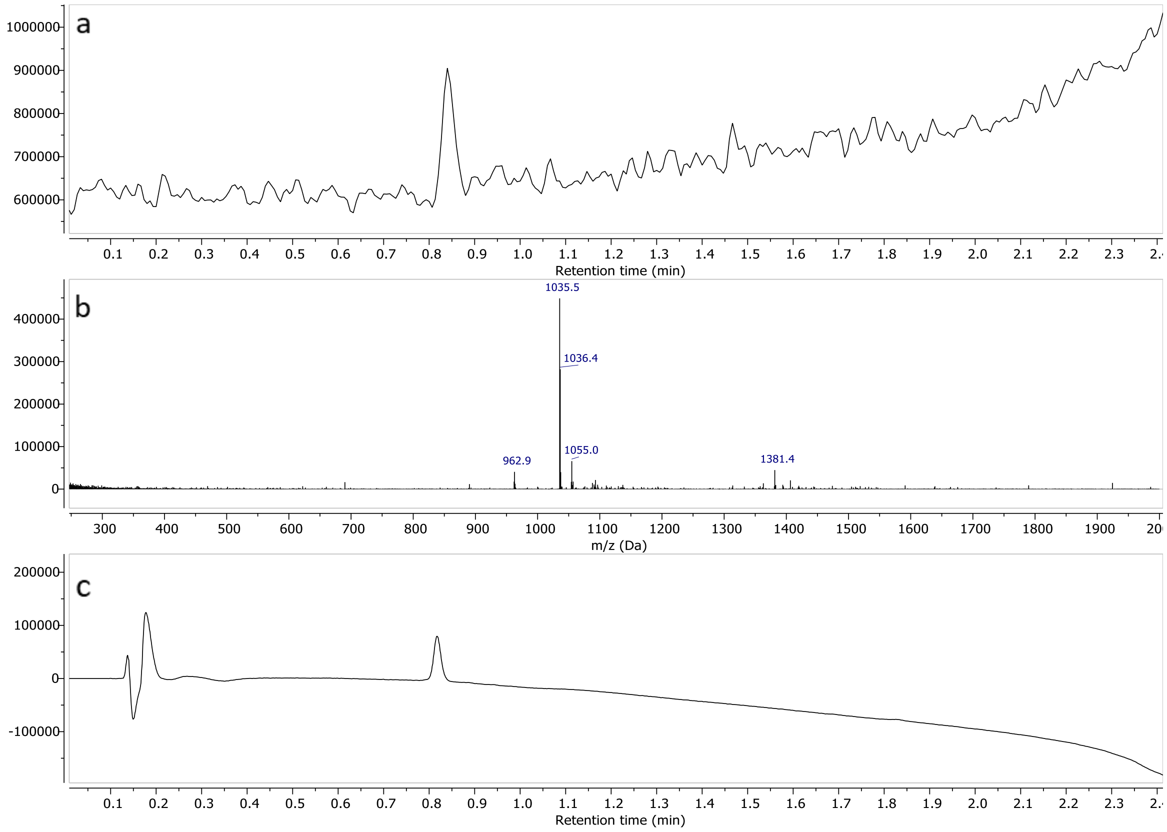


a) TIC – Total Ion Chromatogram (ESI-) b) mass spectrum of **15** (ESI-) c) UV spectrum at 190nm

#### Compound 16

#### $^1$H-NMR ($D_2O$, 600MHz)

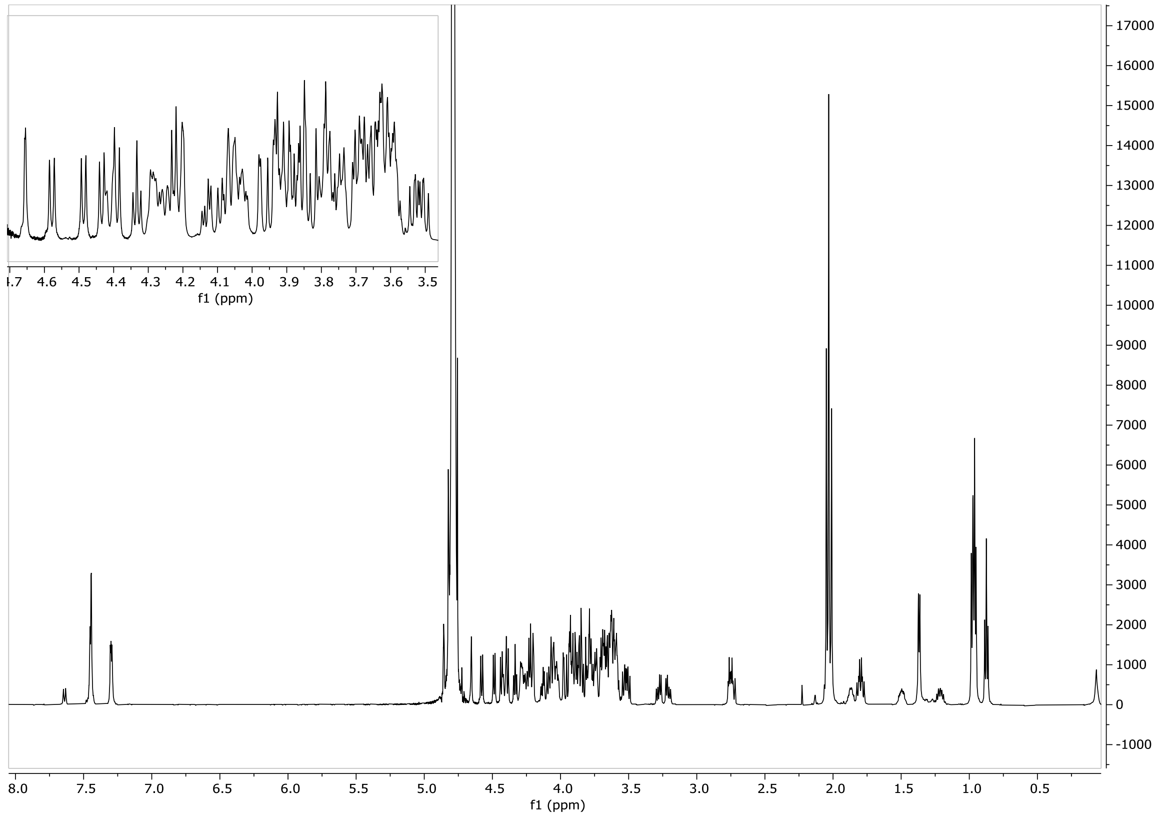


#### HSQC ($D_2O$, 600 MHz)

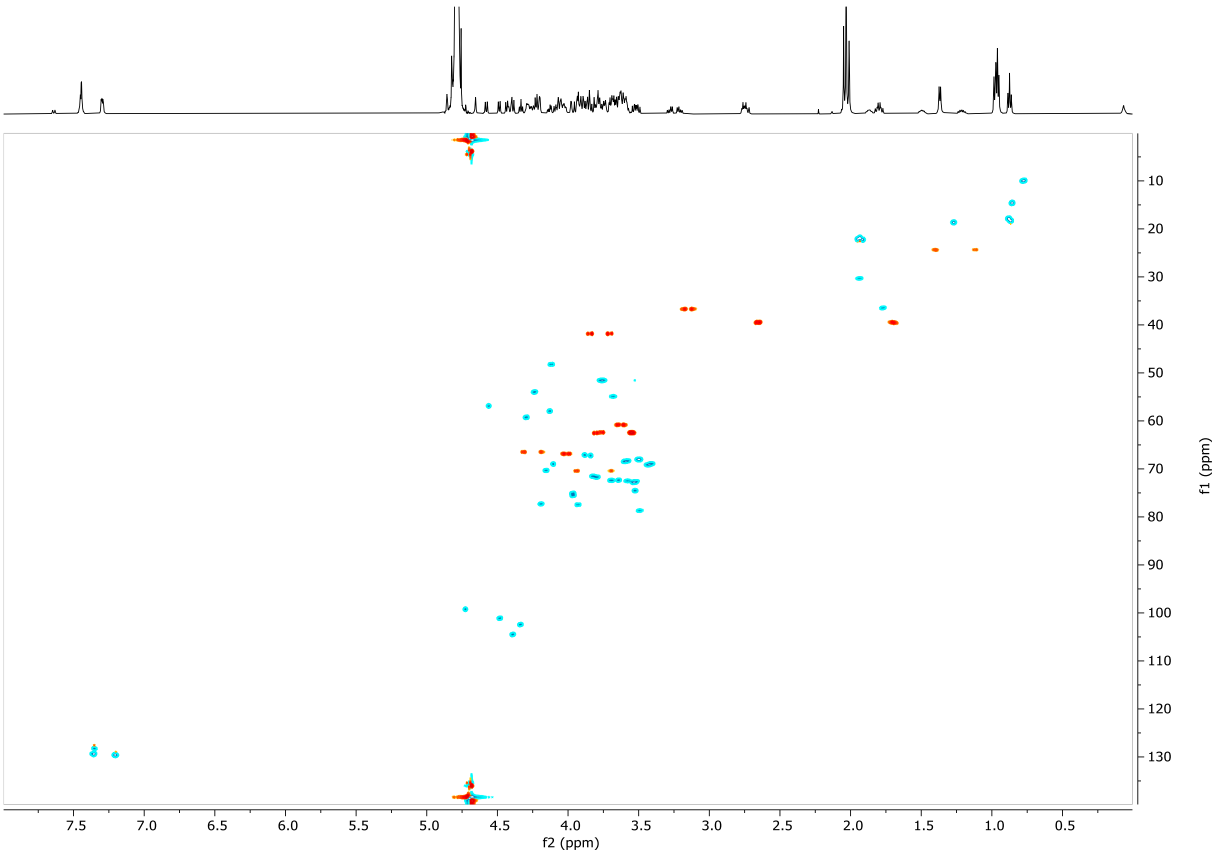

#### LCMS

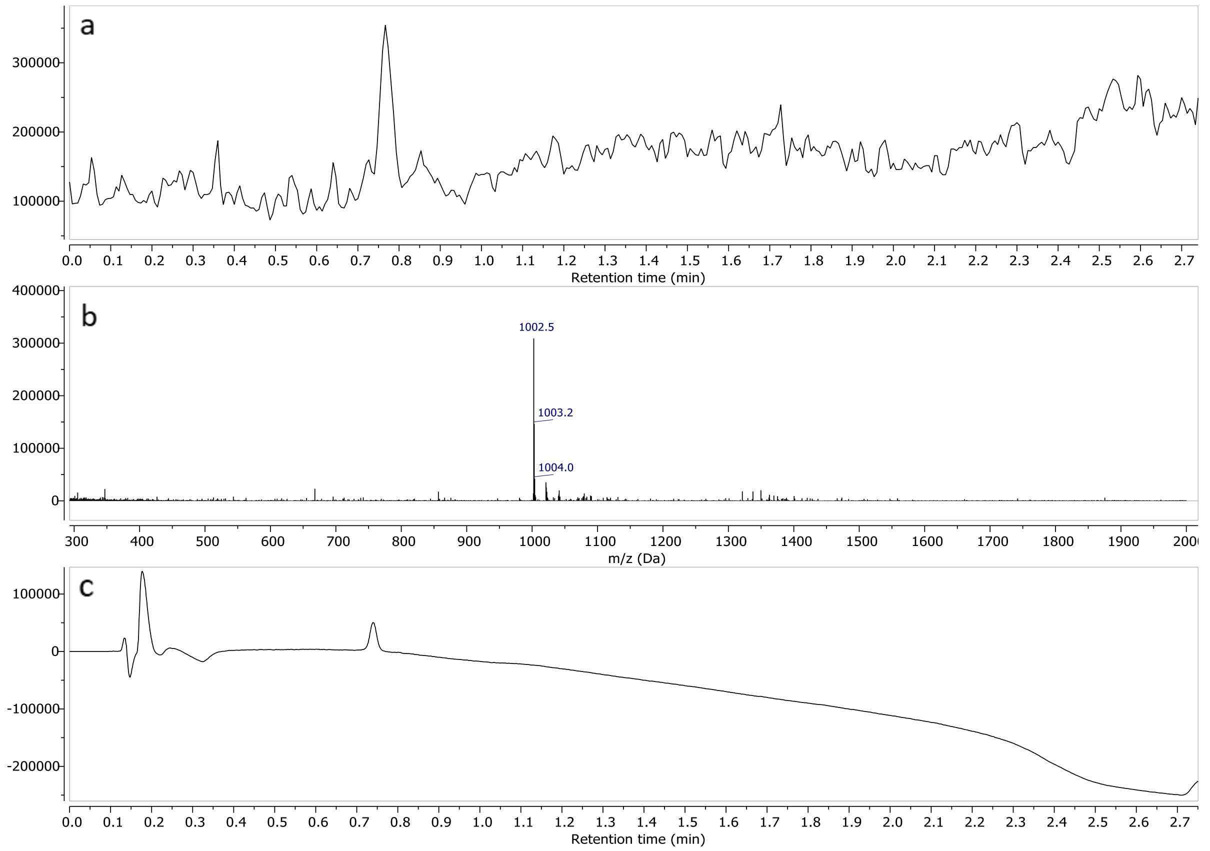


a) TIC – Total Ion Chromatogram b) mass spectrum of **16** c) UV spectrum at 190nm

#### Compound 17

##### $^{1}$H-NMR ($D_2O$, 600MHz)

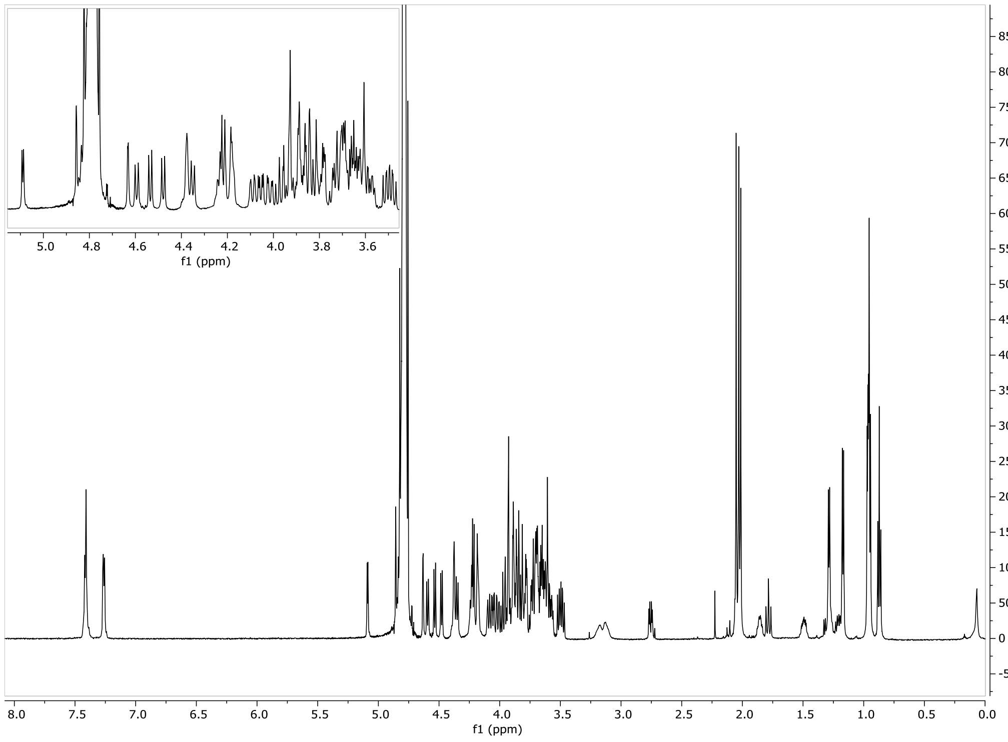

#### HSQC ($D_2O$, 600 MHz)

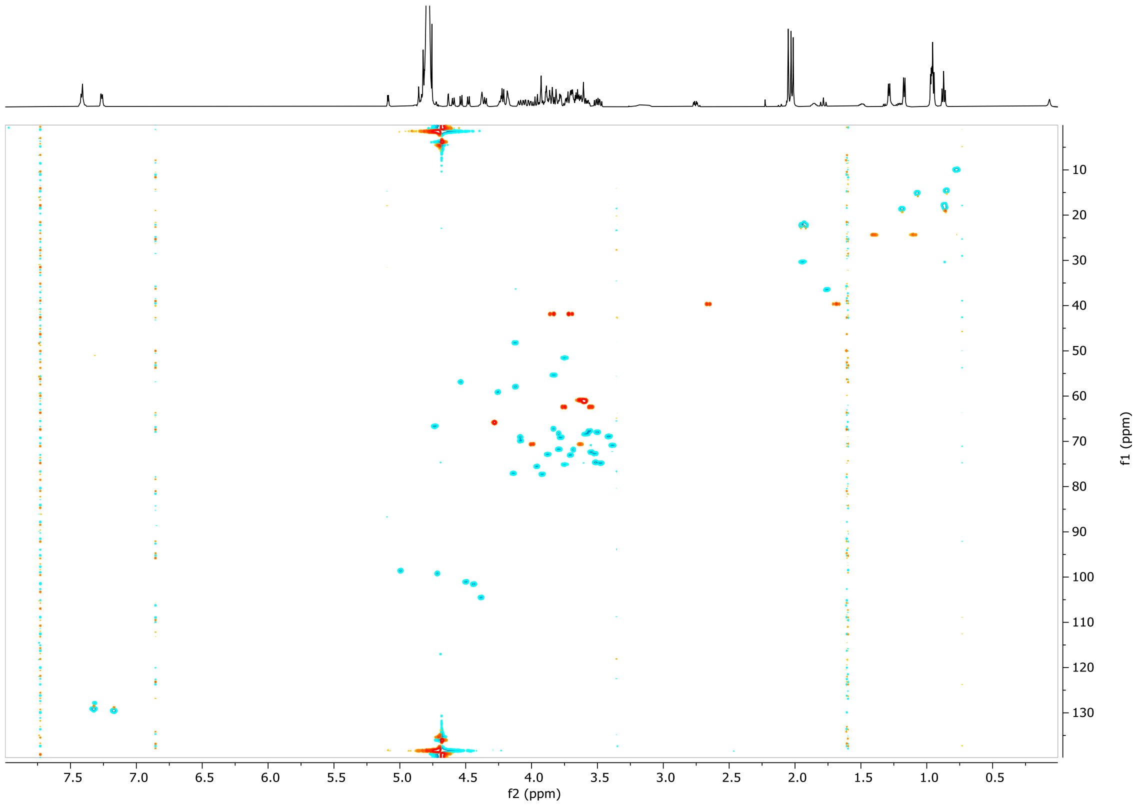


#### LCMS

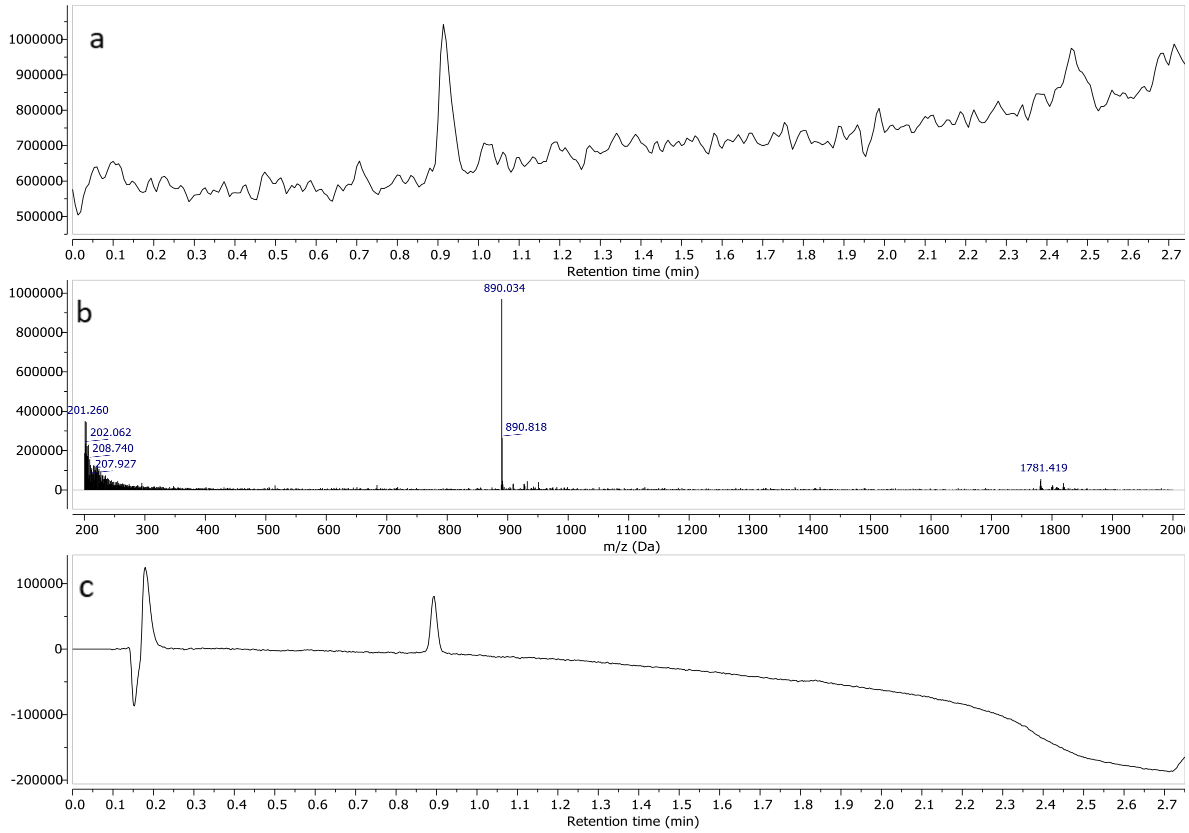


a) TIC – Total Ion Chromatogram (ESI-) b) mass spectrum of **17** (ESI-) c) UV spectrum at 190nm

#### Compound 18

#### $^1$H-NMR ($D_2O$, 600MHz)

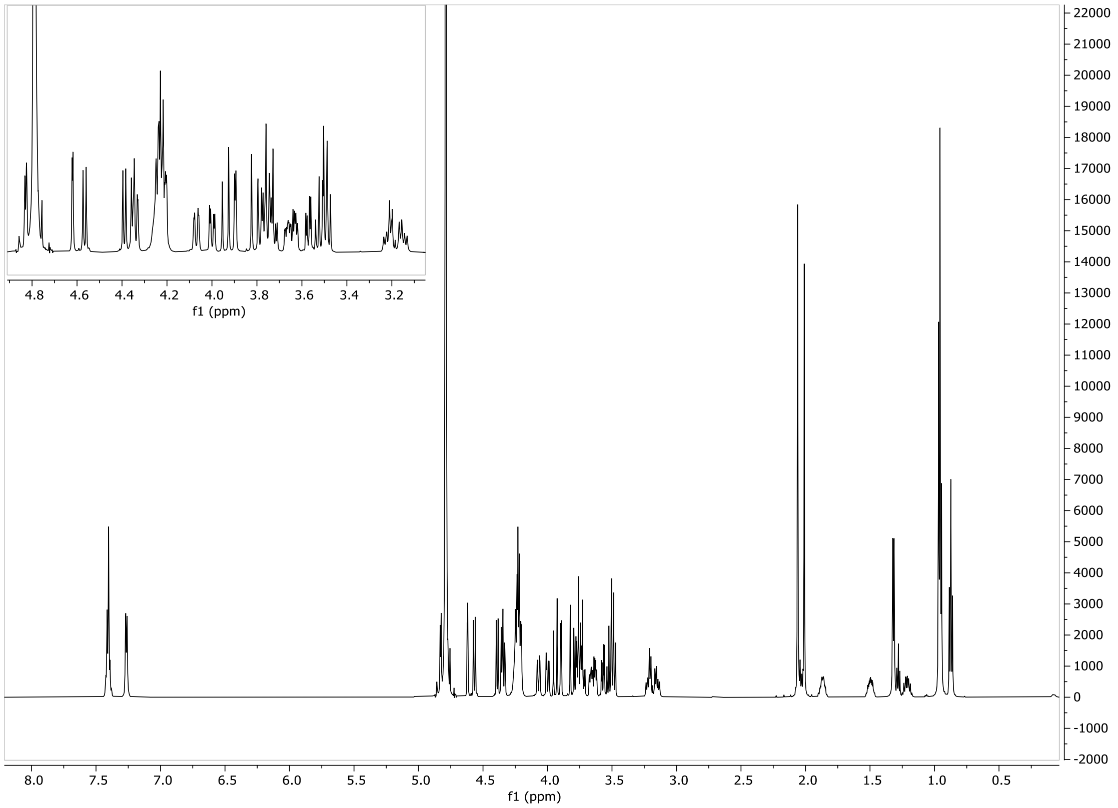


#### HSQC ($D_2O$, 600 MHz)

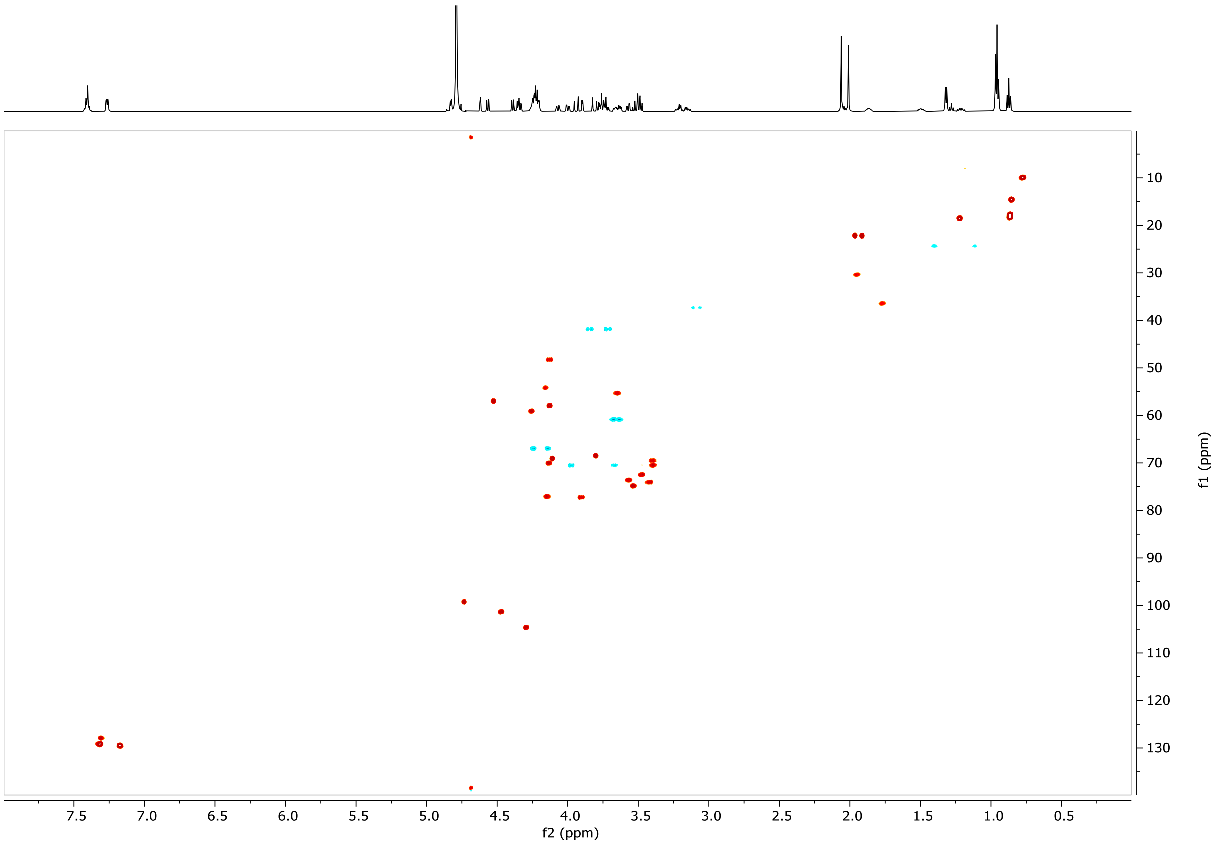


#### LCMS

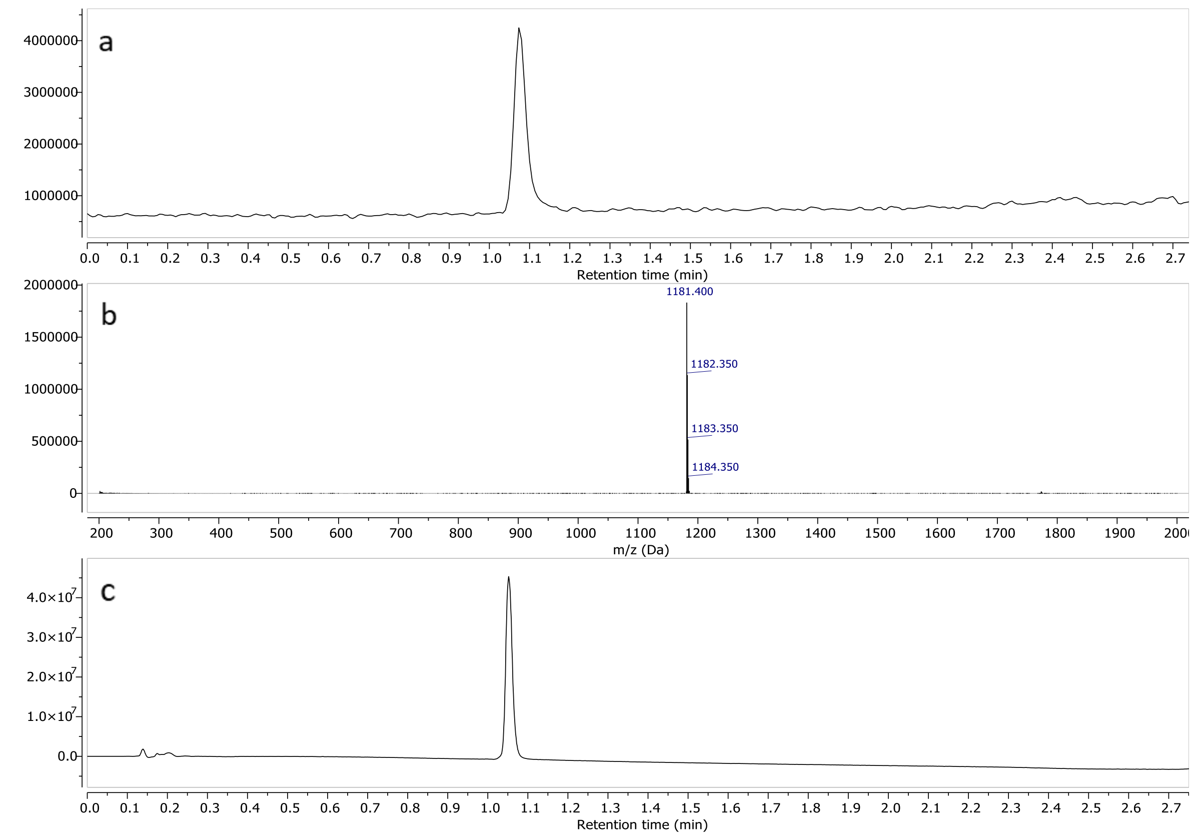


a) TIC – Total Ion Chromatogram (ESI-) b) mass spectrum of **18** (ESI-) c) UV spectrum at 190nm

#### Compound 19

##### $^1$H-NMR ($D_2O$, 600MHz)

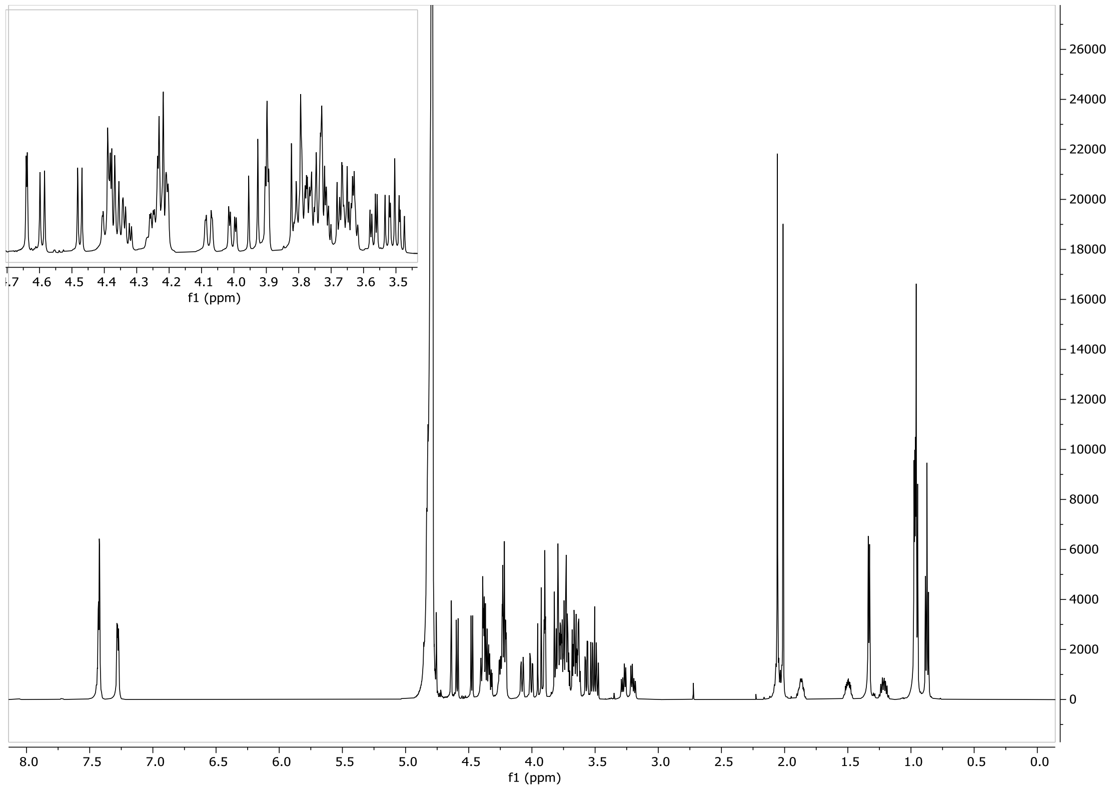

#### HSQC ($D_2O$, 600 MHz)

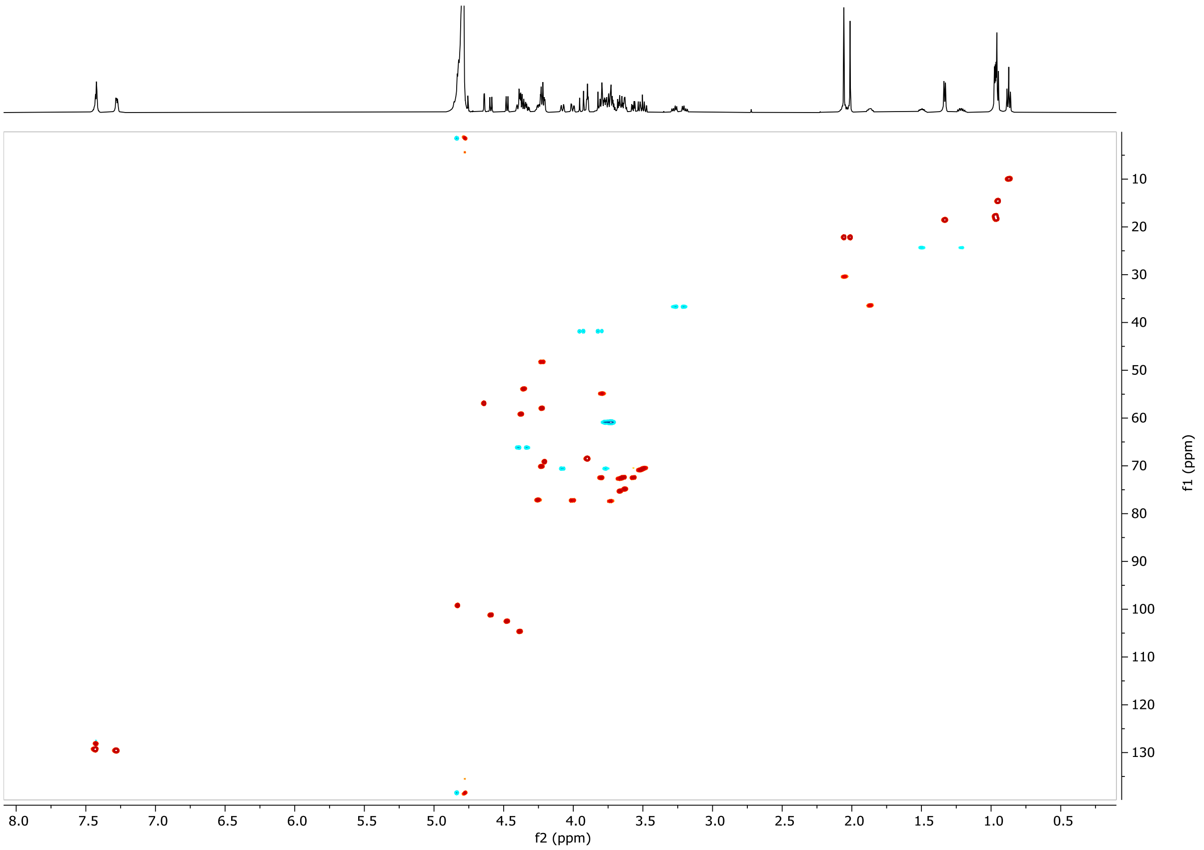


#### LCMS

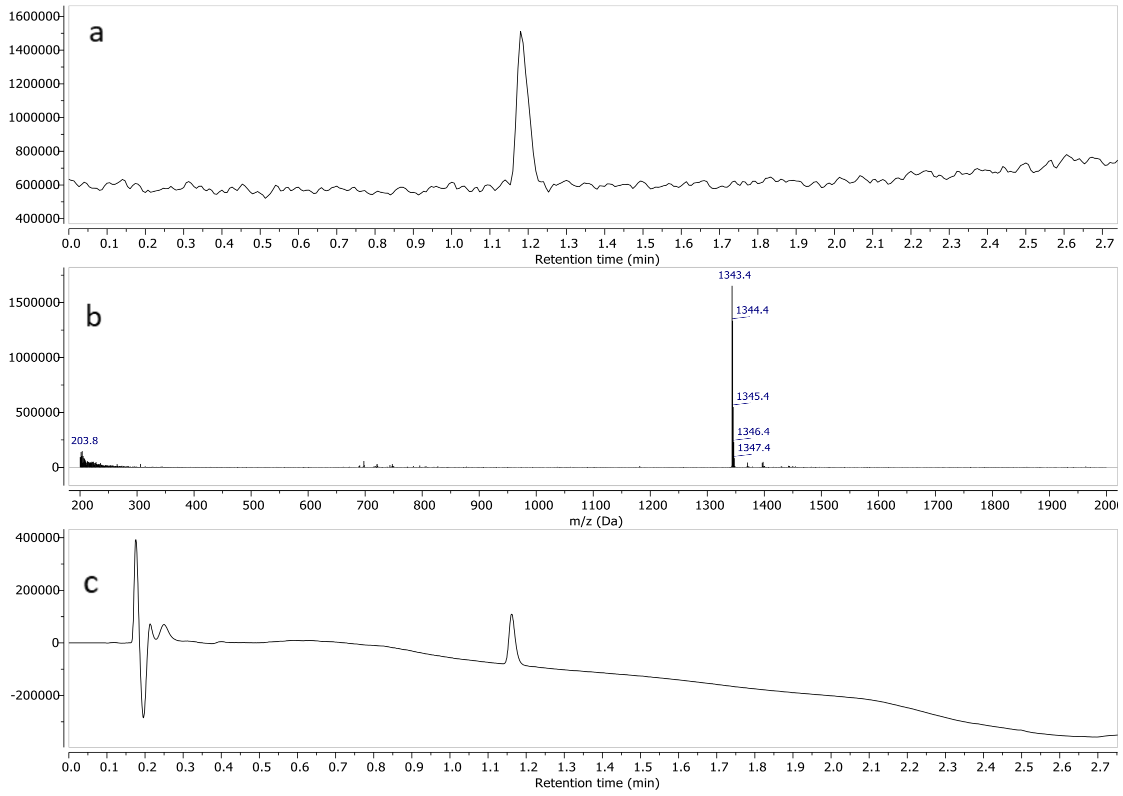


a) TIC – Total Ion Chromatogram (ESI-) b) mass spectrum of **19** (ESI-) c) UV spectrum at 190nm

#### Compound 20

##### $^{1}H$-NMR ($D_2O$, 600MHz)

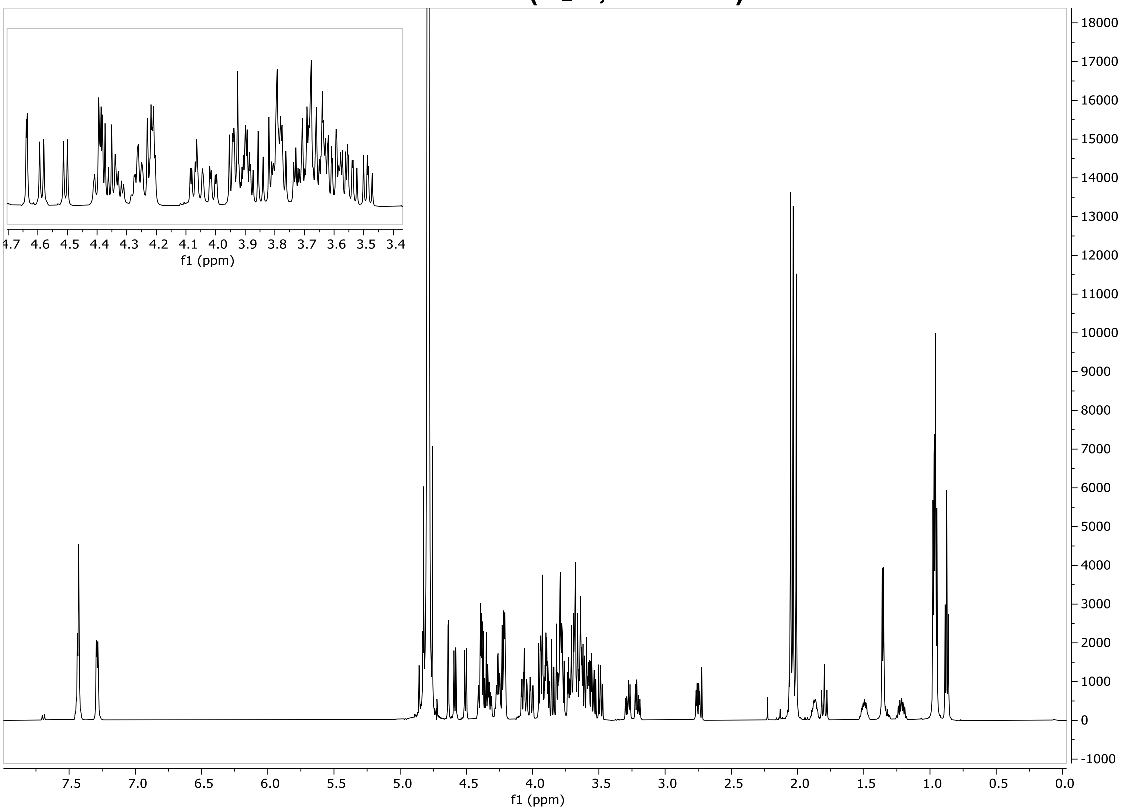


##### HSQC ($D_2O$, 600 MHz)

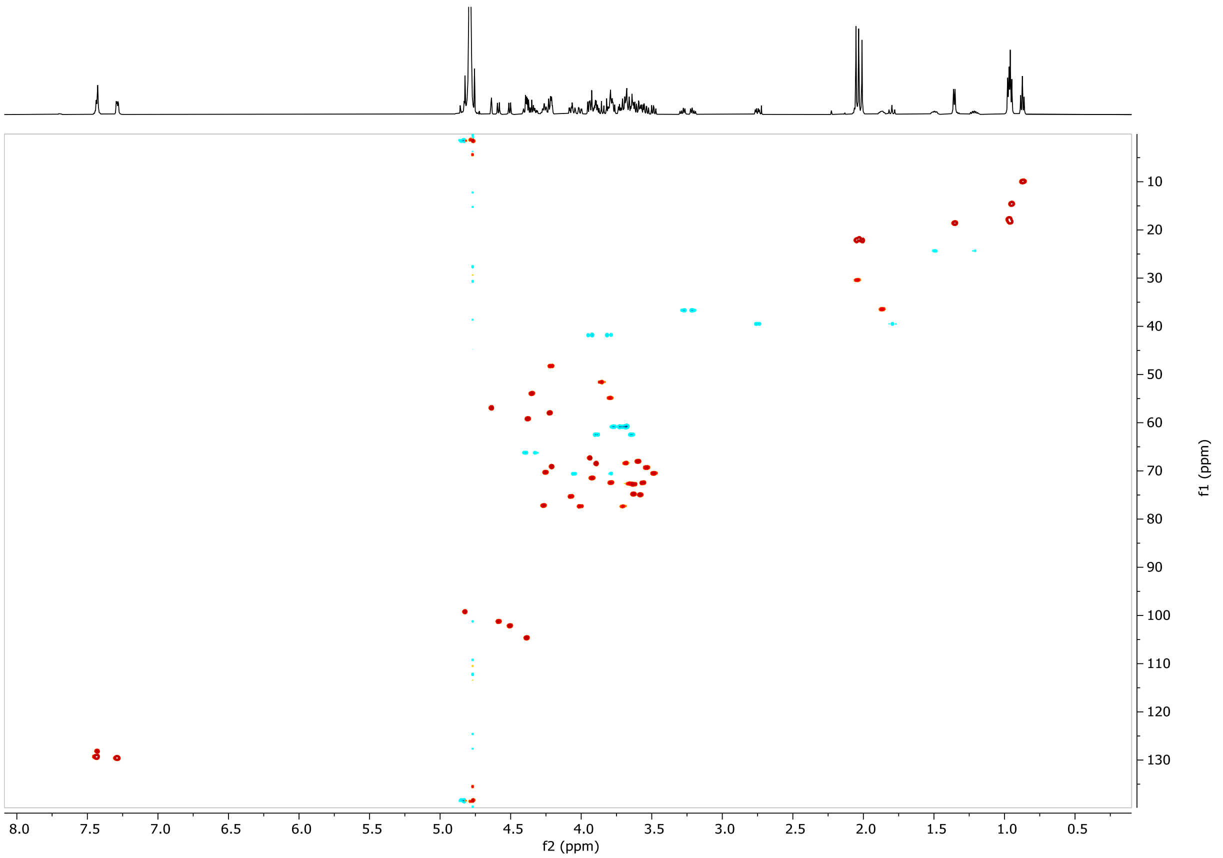


##### LCMS

a

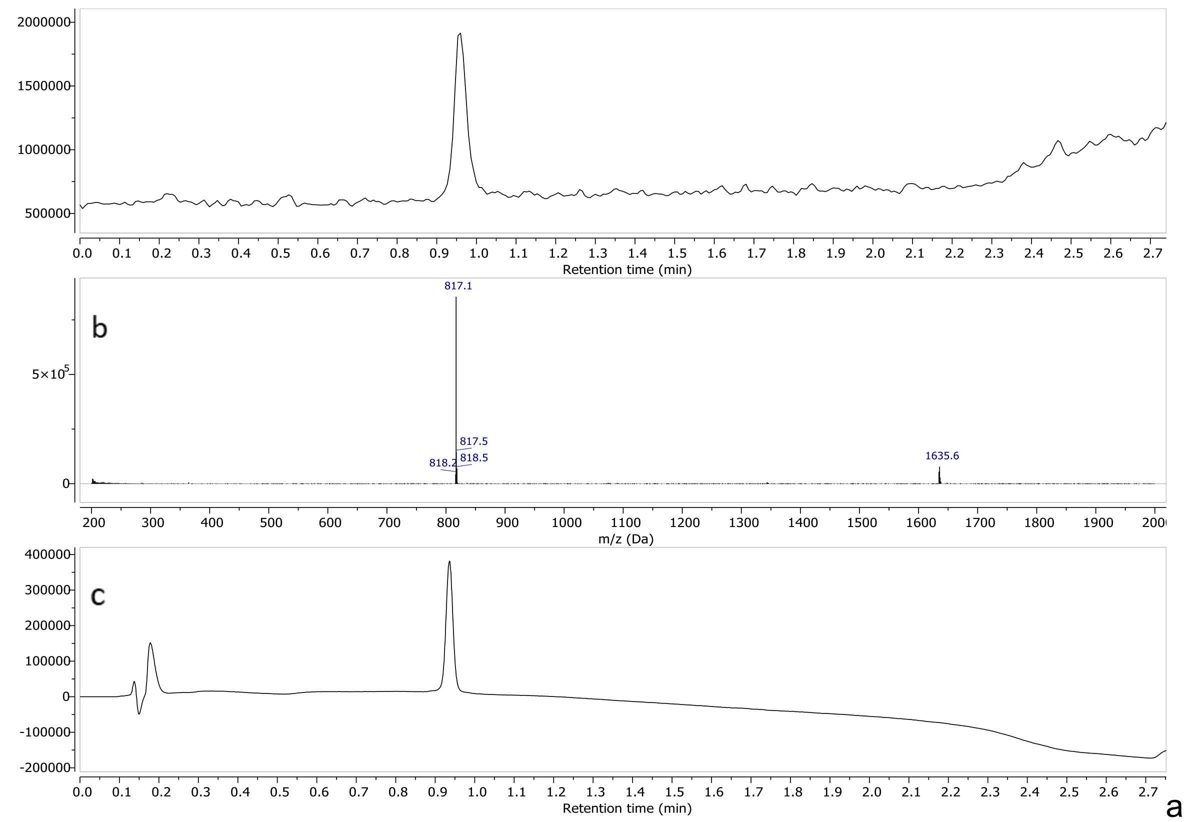


a) TIC – Total Ion Chromatogram (ESI-) b) mass spectrum of **20** (ESI-) c) UV spectrum at 190nm

#### Compound 21

##### $^1$H-NMR ($D_2O$, 600MHz)

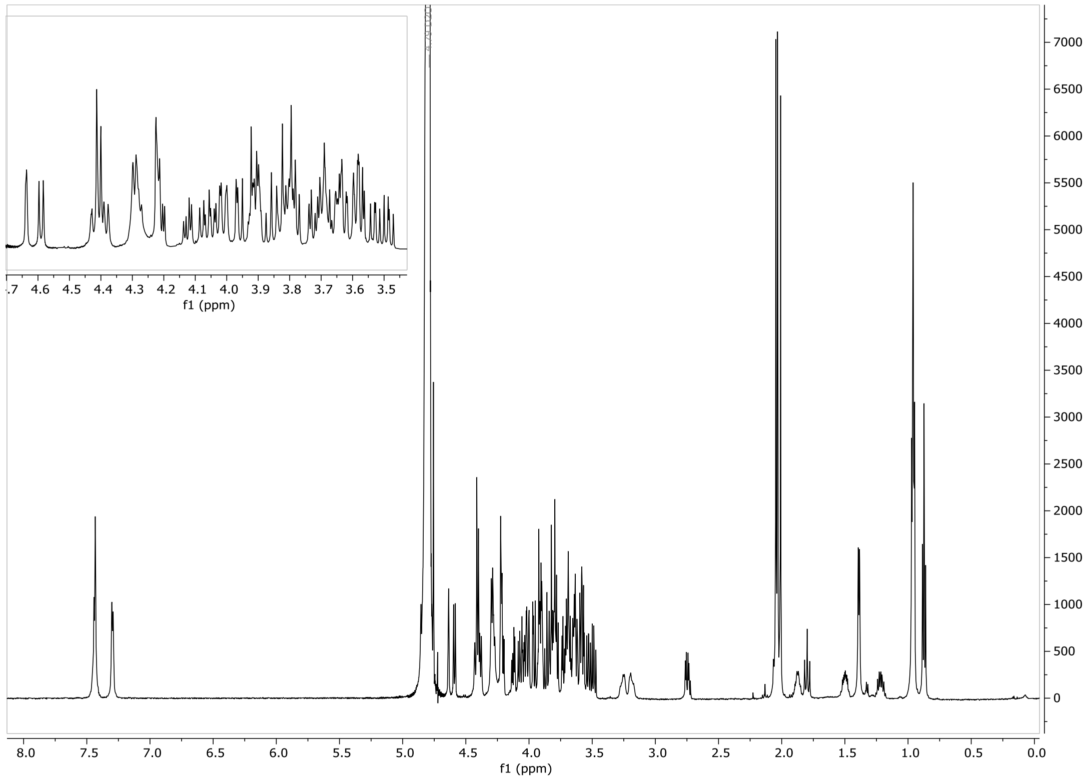

#### HSQC ($D_2O$, 600 MHz)

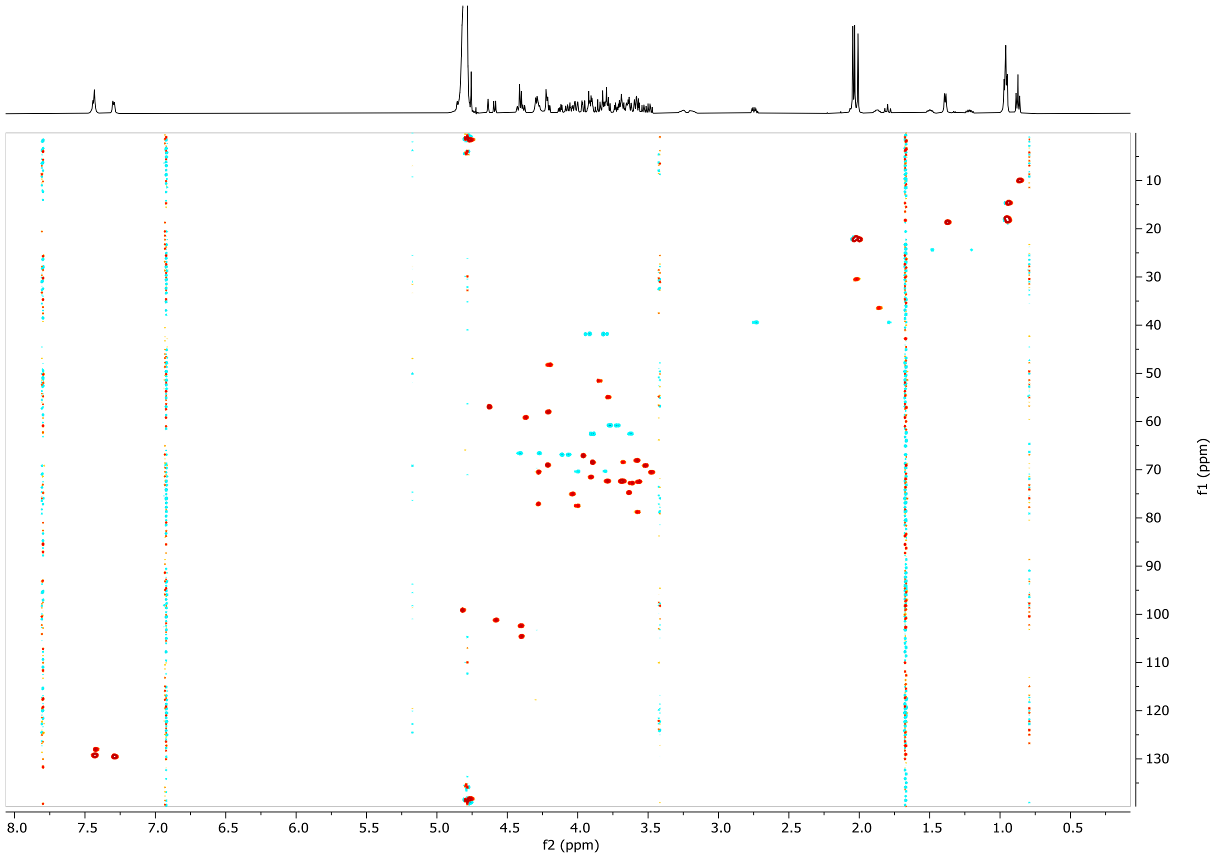


#### LCMS

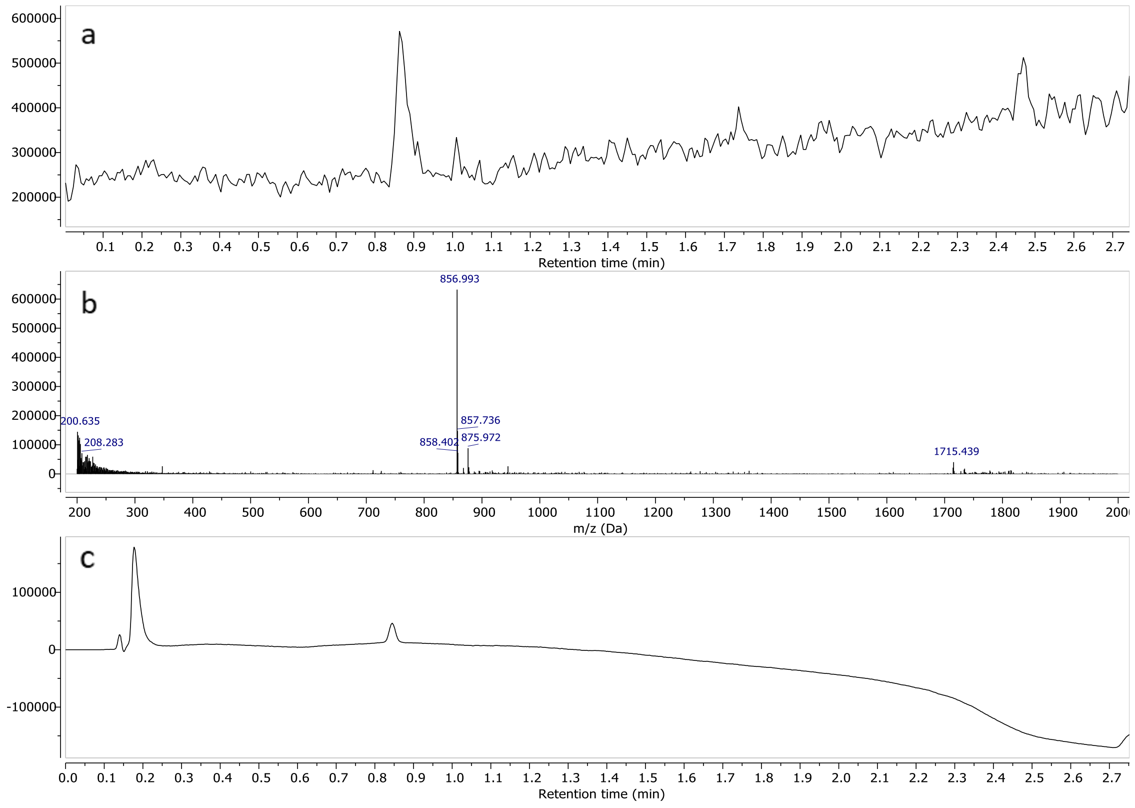


a) TIC – Total Ion Chromatogram (ESI-) b) mass spectrum of **21** (ESI-) c) UV spectrum at 190nm

#### Compound 22

##### $^1$H-NMR ($D_2O$, 600MHz)

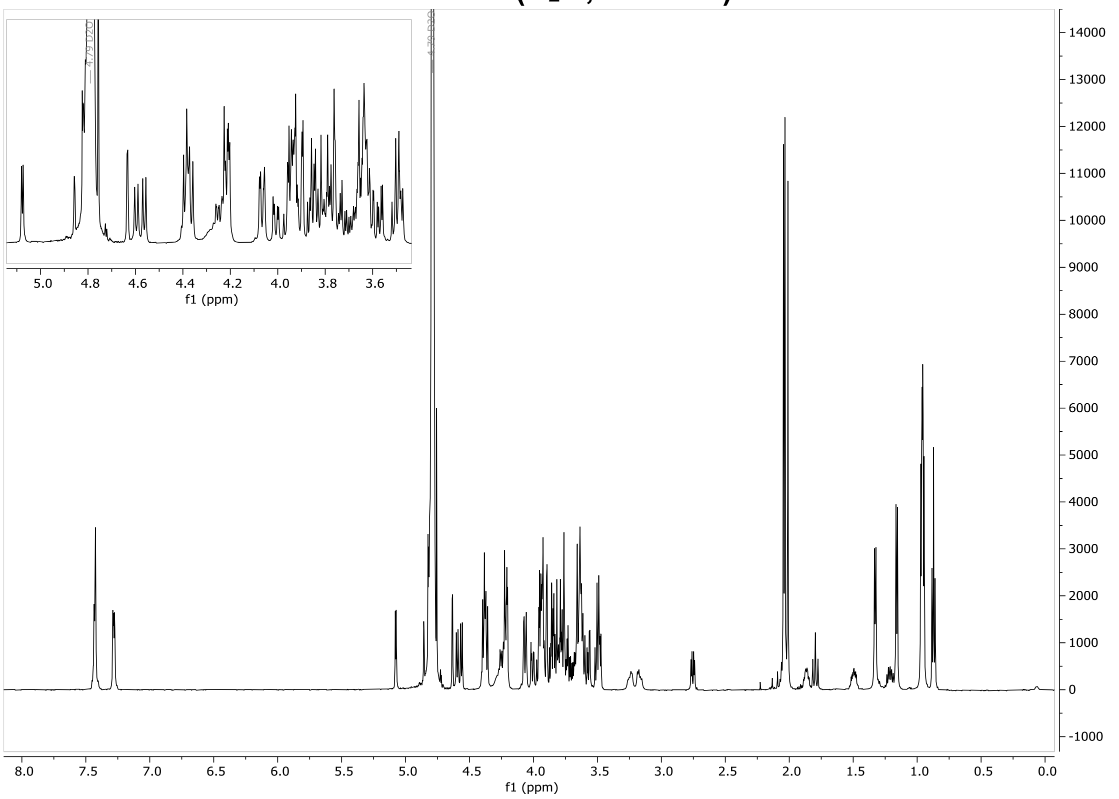


b)

##### c) HSQC ($D_2O$, 600 MHz)

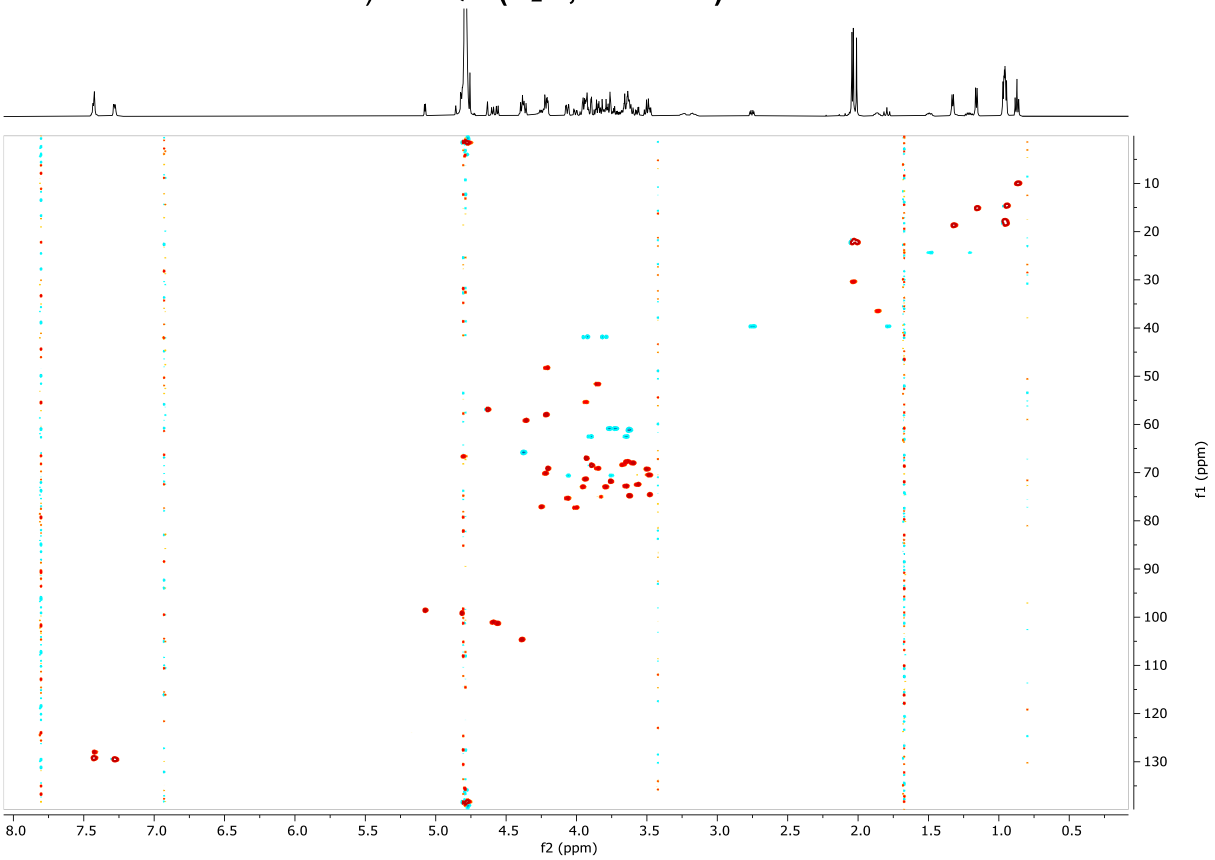


d)

#### f) **LCMS**

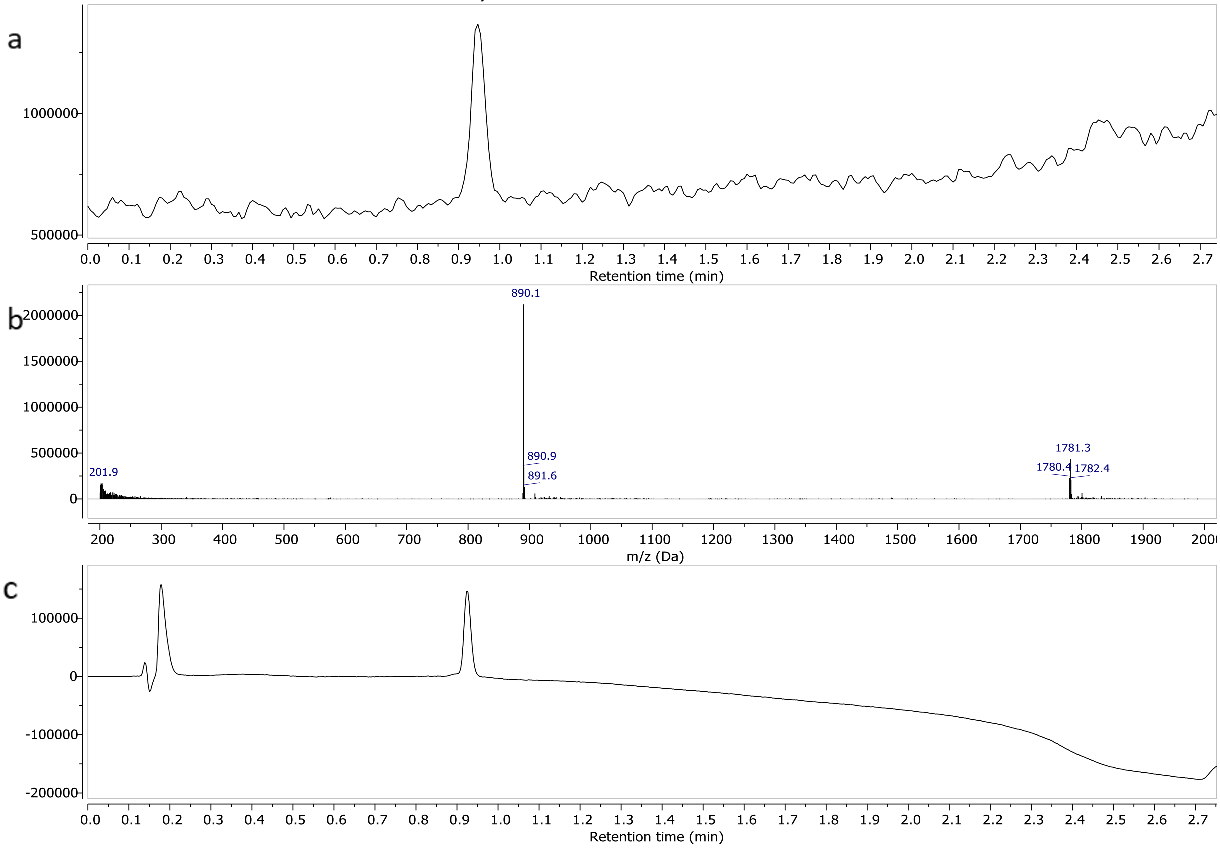


g)

a) TIC – Total Ion Chromatogram (ESI-) b) mass spectrum of **22** (ESI-) c) UV spectrum at 190nm

#### **Core 3 (23)**

##### **$^1$H-NMR ($D_2O$, 600MHz)**

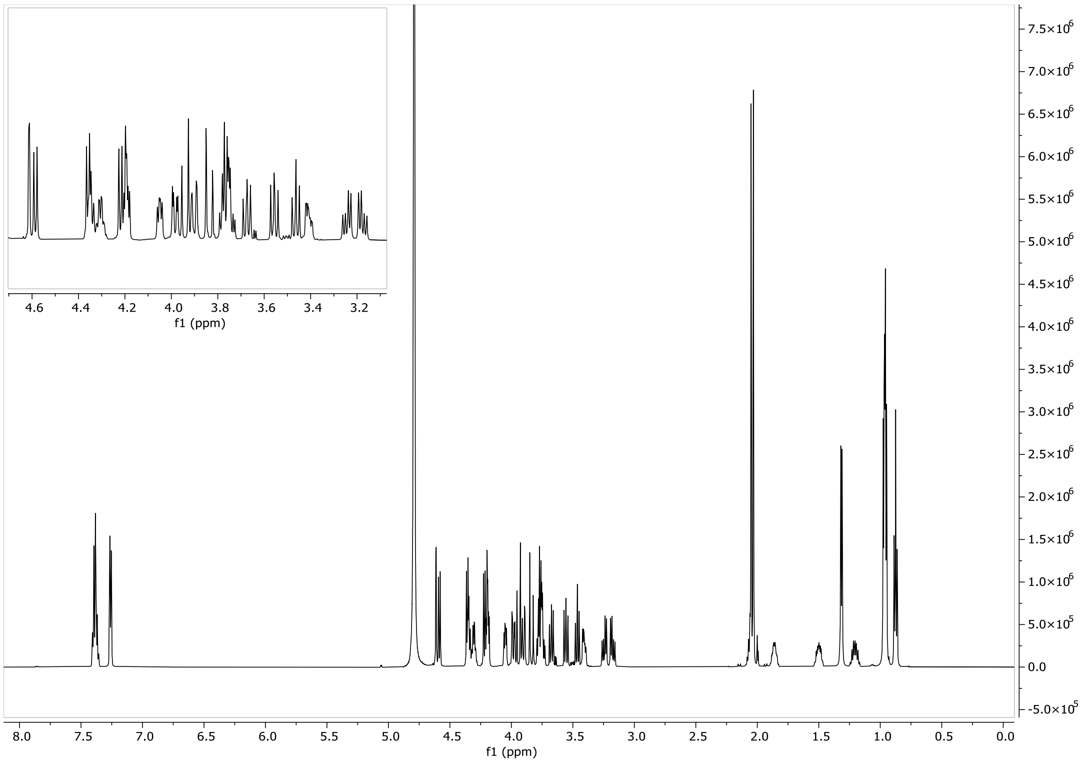

#### Compound 24

#### $^{1}$H-NMR ($D_2O$, 600MHz)

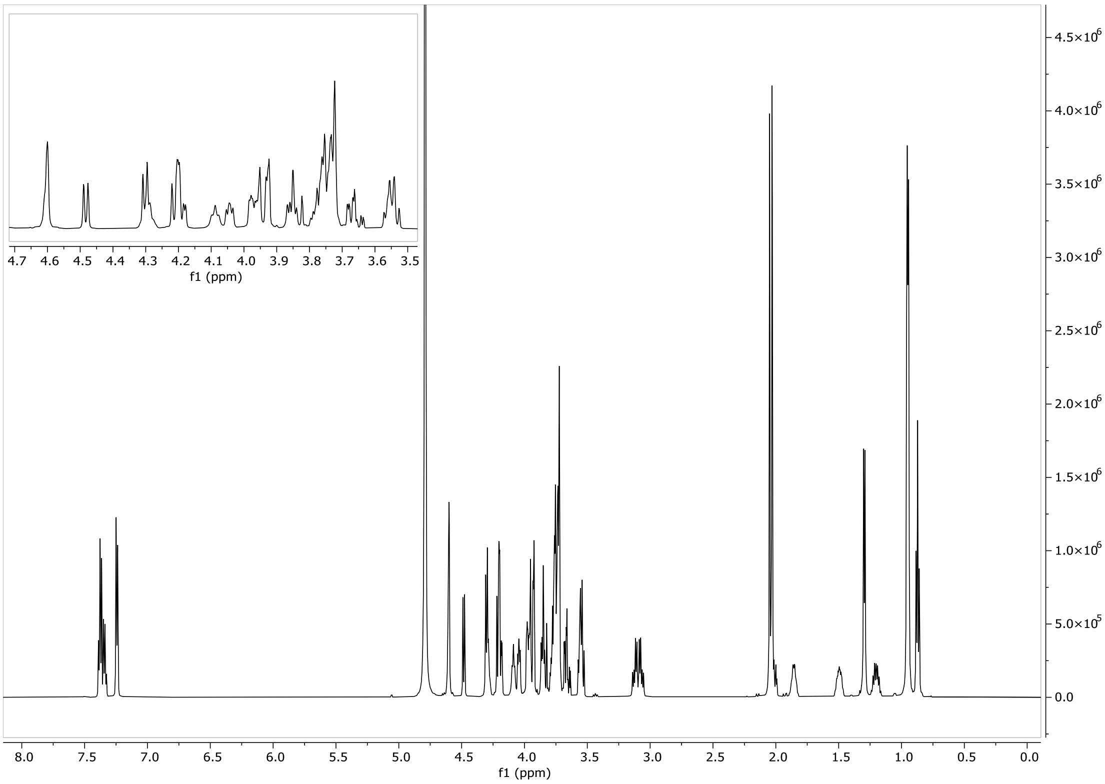


#### HSQC ($D_2O$, 600 MHz)

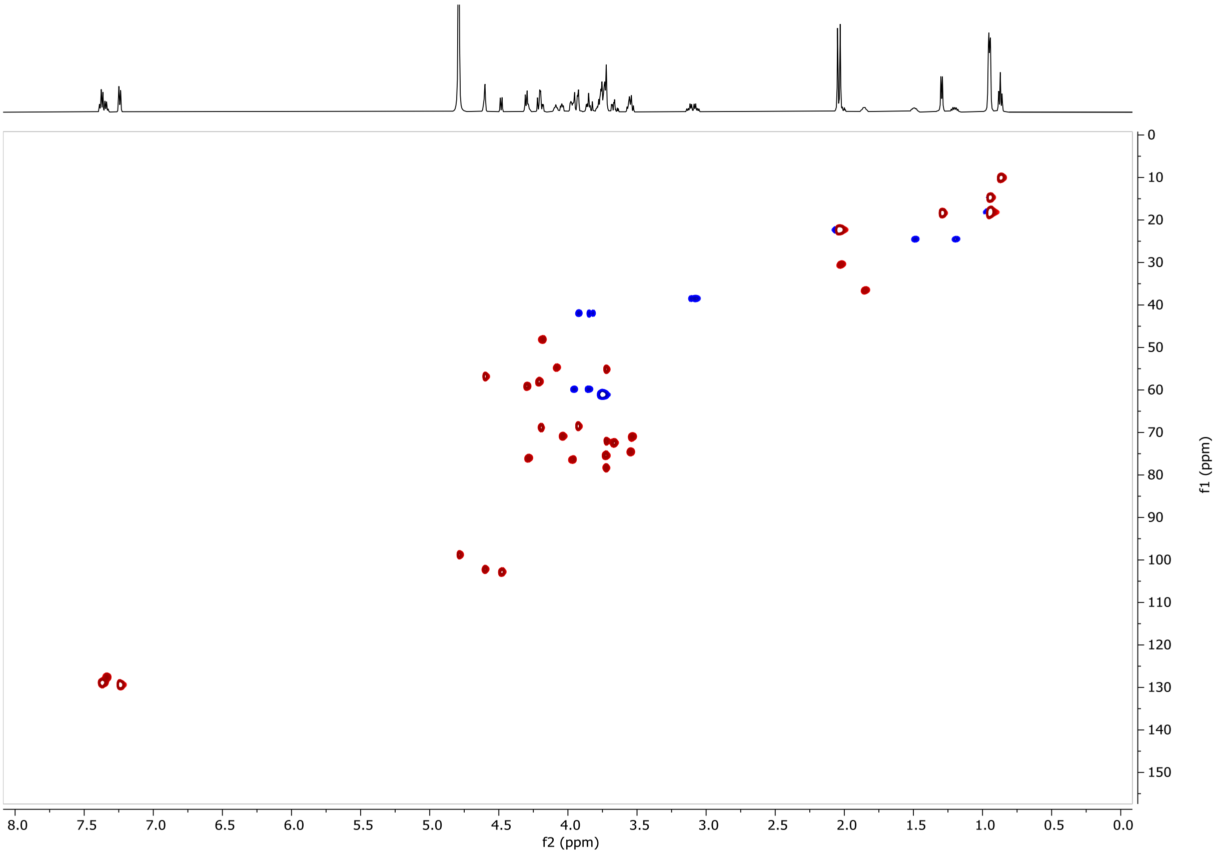

#### LCMS

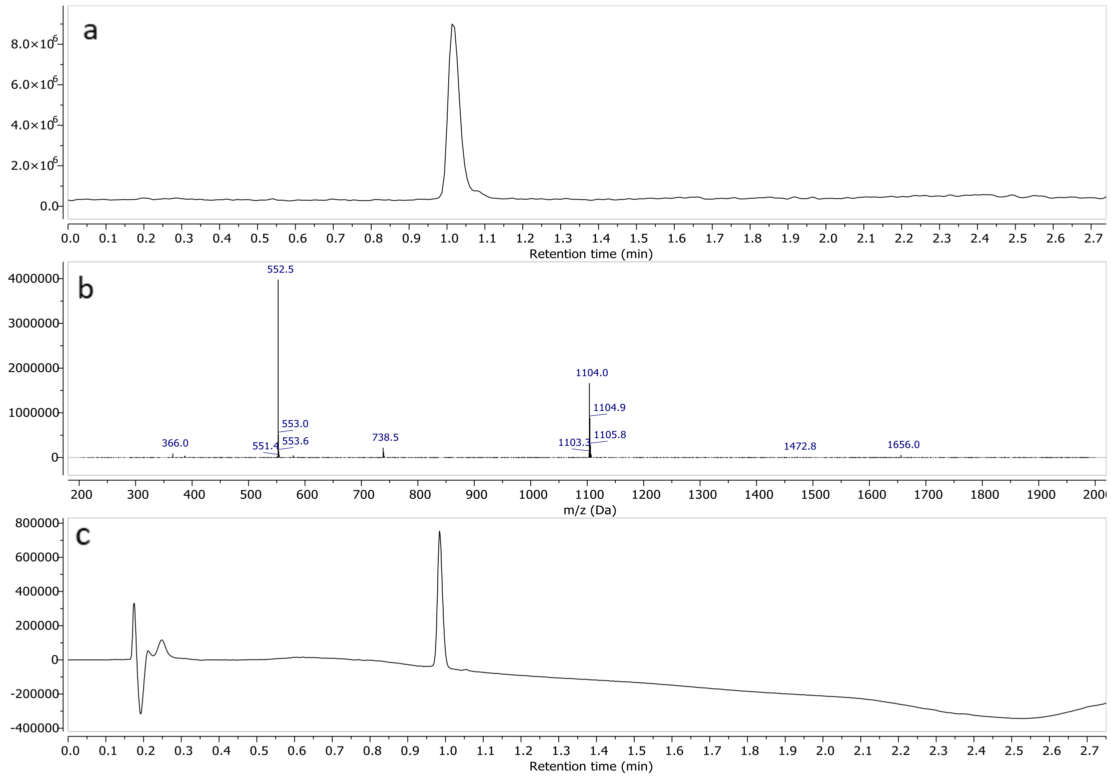


a) TIC – Total Ion Chromatogram (ESI+) b) mass spectrum of **24** (ESI+) c) UV spectrum at 190nm

#### Compound 25

#### $^{1}$H-NMR ($D_2O$, 600MHz)

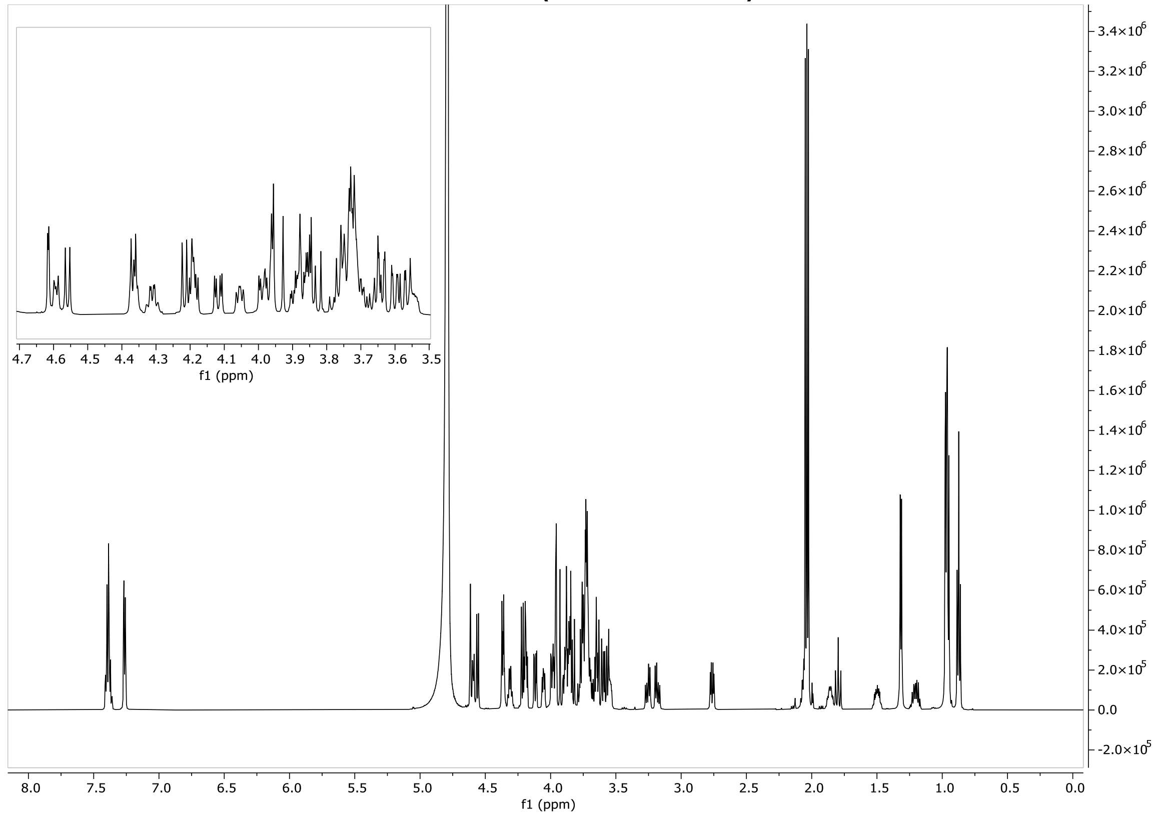

#### HSQC ($D_2O$, 600 MHz)

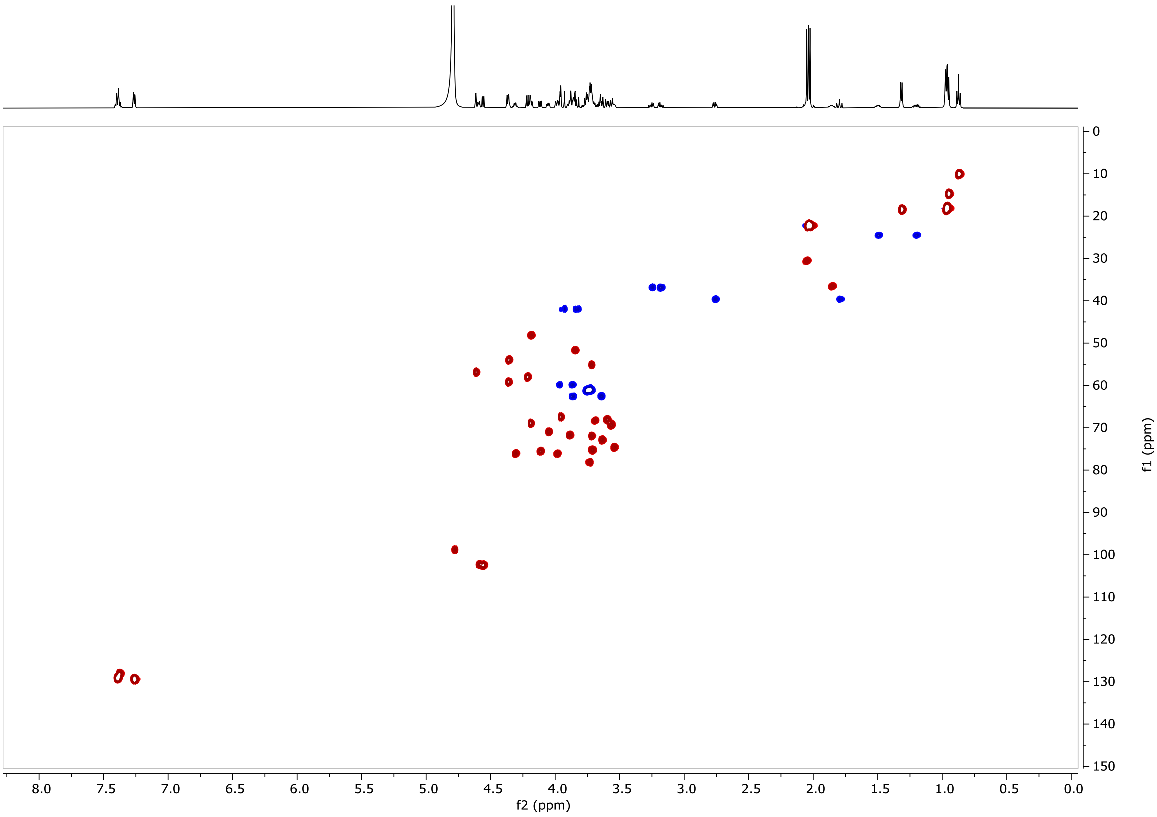


#### LCMS

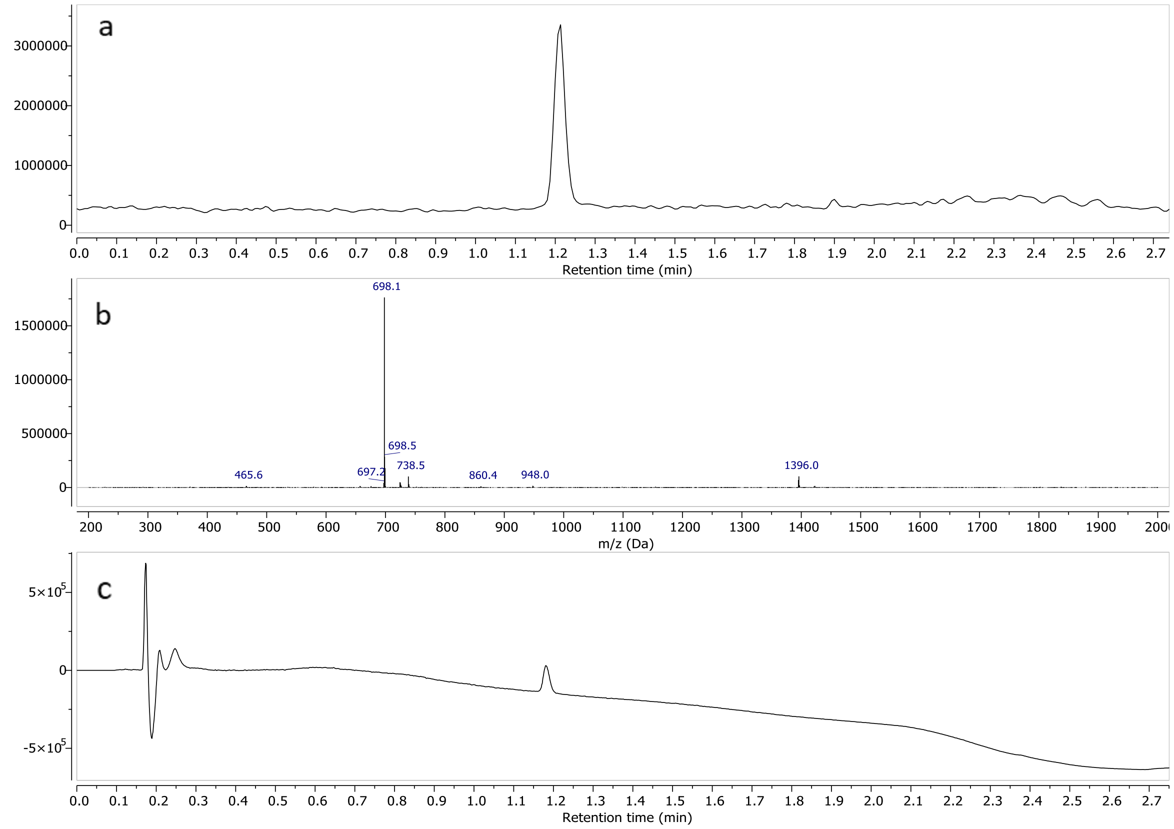


a) TIC – Total Ion Chromatogram (ESI+) b) mass spectrum of **25** (ESI+) c) UV spectrum at 190nm

#### Compound 26

#### $^1$H-NMR ($D_2O$, 600MHz)

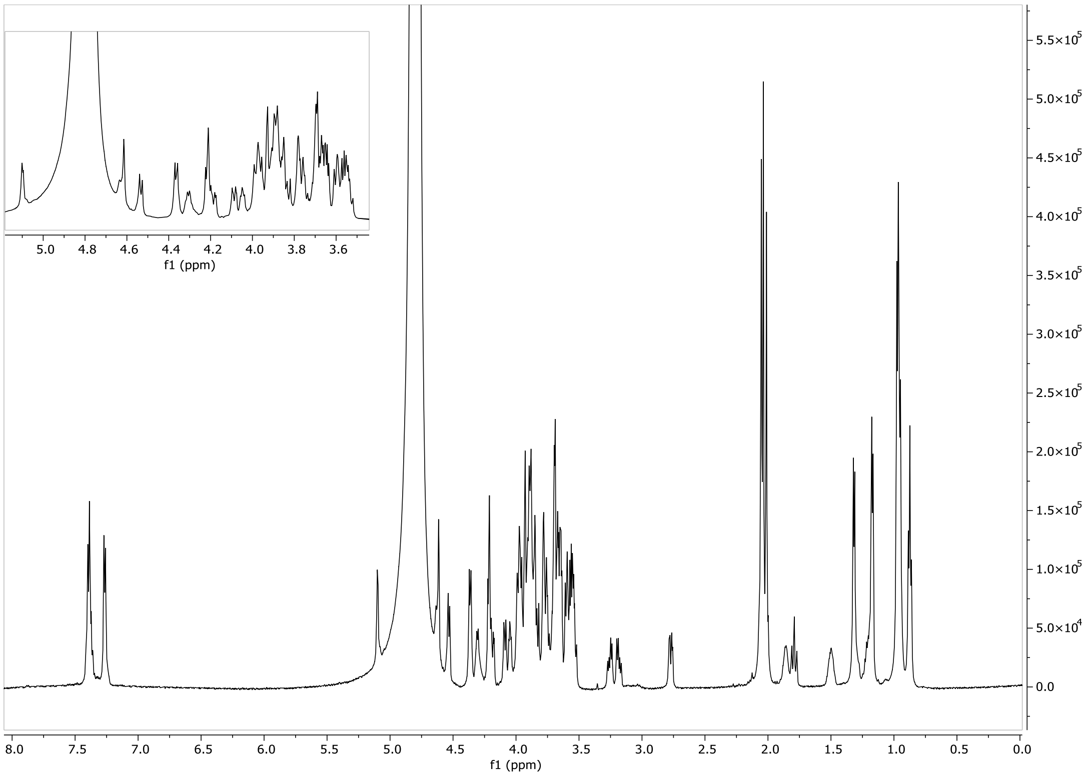


#### HSQC ($D_2O$, 600 MHz)

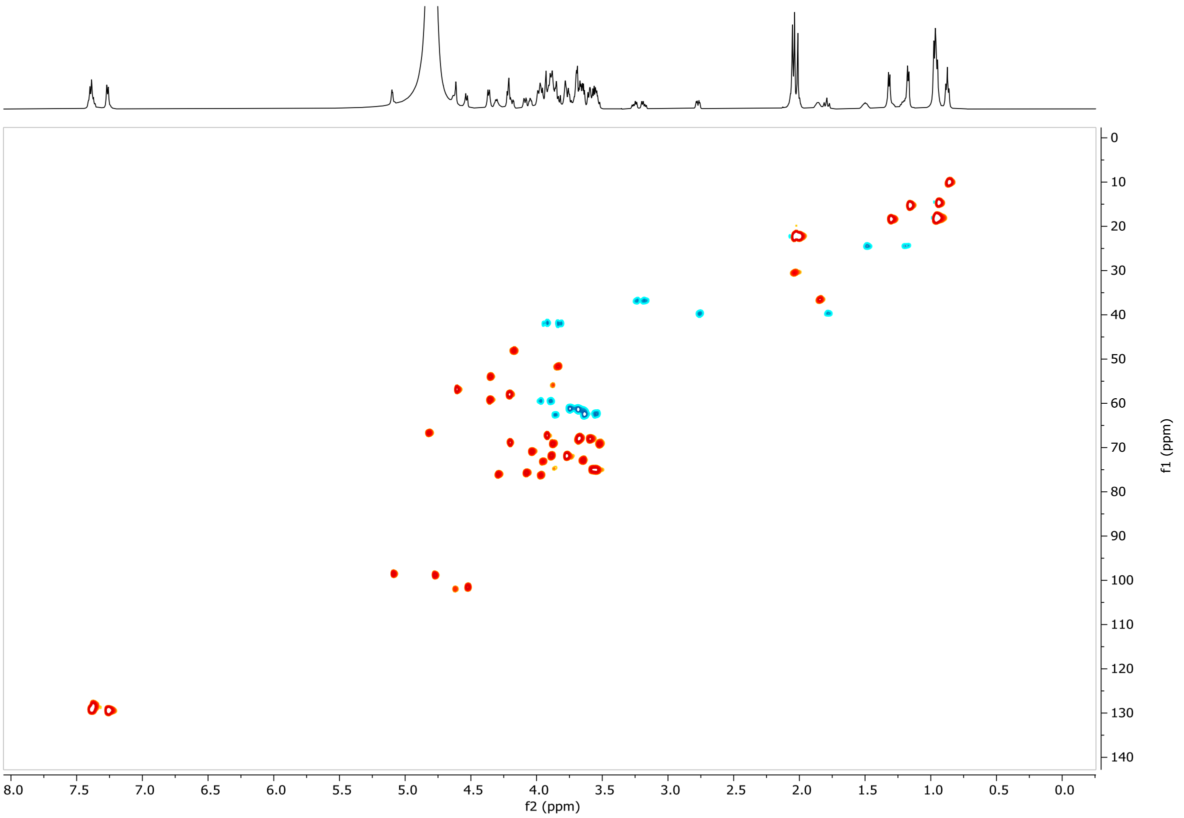

### LCMS

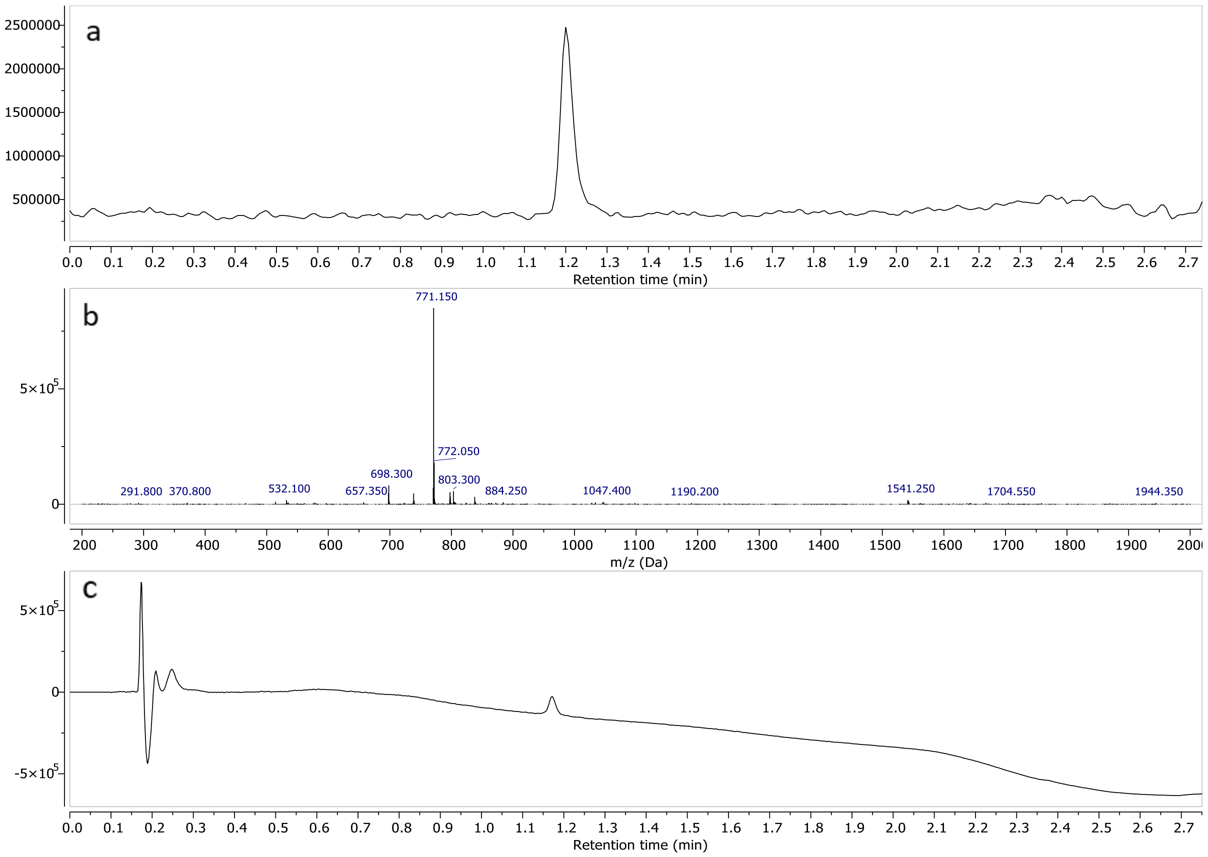


a) TIC – Total Ion Chromatogram (ESI+) b) mass spectrum of **26** (ESI+) c) UV spectrum at 190nm
